# Mutated Tumor Suppressors Follow Oncogenes Profile by the Gene Hypermethylation of Partners in the Protein Interaction Networks

**DOI:** 10.1101/2022.03.28.486156

**Authors:** Somnath Tagore, Milana Frenkel-Morgenstern

## Abstract

As a result of current advances in the analysis of patient sequencing data, many tumors have been characterized in a personalized manner. Such data can also be used to characterize genes that act as either oncogenes or tumor suppressors. These include “defective” tumor suppressor genes which may function as driver oncogenes that play a key role in cancer proliferation due to various genetic alterations, specifically, chromosomal translocations. In this study, we considered protein networks, mutations, methylation data and cancer fusions to classify tumor suppressors that may convert into oncogenes. Moreover, we developed a novel network-based parameter called the ‘preferential attachment score’ to categorize genes as oncogenes and/or tumor suppressors. Such classification was achieved using a naïve Bayes computation approach. We used an ABC-MCMC method for selecting features for training our classification algorithm. We then performed a survey of tumor suppressors and oncogenes from the perspective of somatic mutations and network properties for 691 TCGA cases. For comparative purposes, we chose currently well-established methods, such as MutSigCV, OncodriveCLUST, Oncodrive-FM, 20/20+, ActiveDriver, MuSiC, TUSON, OncodriveFML, and found that our algorithm outperformed these other tolls, with 93.3% efficiency. Based on 691 TCGA cohorts, we found that tumor suppressors presented the highest mutation frequency in most tumor types, relative to oncogenes. Using protein-protein interaction data, we found that essential proteins, tumor suppressors and oncogenes had higher degrees of connectivity and betweenness centrality, relative to normal proteins. Similarly, tumor suppressors and oncogenes had lower clustering coefficients, as well as shortest path distances (FDR < 0.05). Finally, most mutated tumor suppressors integrate hyper-methylated partners in the protein interaction networks of 3091 fusions, following the patterns of oncogenes (43%). Thus, these results further characterize cancer oncogenes and tumor suppressors in the context of deep analysis of cancer network alterations.

**Availability:** Source scripts are available at https://github.com/somnathtagore/NBC and the resource is available at http://ontum.md.biu.ac.il/index.html

## 1 Introduction

### 1.1 Background

Since the first cancer genome was sequenced in 2008, various large-scale studies analyzing multiple tumor types have been conducted (***Ley et al., 2008; Jaratlerdsiri et al., 2018; Deng et al., 2018)***. These analyses also generated catalogs of tumor-specific mutations, such as single nucleotide variants. Such mutation data can further be used to identify and characterize genes that act as either oncogenes (OG) or tumor suppressor genes (TS). Numerous studies have demonstrated genetic alternations involving the gain-of-function of OGs, together with the loss-of-function of TSs which determine cell cycle processes that control tumor formation and development. Furthermore, mutation rates in tumors vary significantly in various cancer types (***Lawrence et al., 2014; Agajanian et al., 2018)***. Therefore, genes with particular patterns of non-random mutations are of interest for further study using specific computational strategies to generating optimal solutions and identify genes that act as are cancer drivers or passengers. Various genomics-based and other experimental efforts have served to refine the compendium of cancer driver genes based on their clinical relevance (***Garraway et al., 2013***). However, many TS and OG remain uncharacterized, as do those TS that may convert into driver oncogenes. Moreover, down-sampling analysis of nearly 5,000 tumor genomes predicted the presence of several driver genes that mutate at low frequencies (***Zhao et al., 2016***).

Protein-protein interaction (PPI) network analysis based on computational methods have also been used to identify disease-specific genes, as well as cancer sub-types and sub-networks (***Ideker et al., 2002; Hofree et al., 2013***). Likewise, as more genetic and genomic data on cancer becomes available, the list of OGs and TSs has been expanded through molecular, cellular, genomic, and computational studies, including non-coding RNA genes (***Wrzeszczynski et al., 2011***). Considering the gain-of-function of OG mutations and loss-of-function of TS mutations, TSs and OGs may also be involved in the regulation of cellular functions in a Yin-Yang manner (***Sun et al., 2011***). Furthermore, OG mutations are usually dominant, such that one mutant copy is enough to impact cellular activity. In contrast, TS mutations tend to be recessive, such that they should follow the famous Knudson’s ‘two-hit hypothesis’, stating that both copies of a tumor suppressor gene must mutate so as to cause a loss-of-function.

At the same time, a growing body of evidence has shown that even partial inactivation of TSs could critically contribute to tumorigenesis (***Berger et al., 2011; Lawrence et al., 2018***). Additionally, functions can be switched between OGs and TSs, depending on the situation. Current therapeutic applications have shown that targeting OGs and their related pathways offers a promising strategy for developing novel drugs, including antibodies and small synthetic molecules (***Osborne et al., 2004***). Therefore, further understanding of OGs and TSs in the terms of networks will provide novel insight into their actions in tumorigenesis.

Some TSs acts as OGs in some cancers. These include epithelial cadherin (E-cadherin), known for its ability to keep cancer cells glued together in a tumor, preventing them from breaking off and metastasizing, yet which can also function as an oncogene in some cancers (***Vergara et al., 2015***). In breast cancer, loss of E-cadherin is generally considered a harbinger of metastasis. However, it has also been found that most breast cancer that had spread retain E-cadherin expression (***Vergara et al., 2015***). Likewise, Smurf2 has also been reported as serving a dual role in cancer by functioning as both a tumor promoter and suppressor, controlling the stability of proteins that play major roles in cell cycle progression, proliferation, differentiation and metastasis. Aberrant Smurf2 expression occurs in breast, esophageal, pancreatic and renal cancers (***Blank et al., 2012***). Ovarian tumors also have been paradoxically found to produce increasing amounts of E-cadherin as they grow (***Anderson et al., 2010***). Karyotyping and epidemiological analyses of mammary tumors at various stages suggest that breast carcinomas become increasingly aggressive through the stepwise accumulation of genetic changes The majority of genetic changes found in human breast cancer can be categorized as either gain-of-function mutations in proto-oncogenes, which stimulate cell growth, division and survival, or loss-of-function mutations in tumor suppressor genes that normally help prevent unrestrained cellular growth and promote DNA repair and cell cycle checkpoint activation. Epigenetic de-regulation also contributes to the abnormal expression of these genes. For example, genes that encode enzymes involved in histone modification are mutated in primary renal cell carcinomas (***van Haaften et al., 2009***). In addition, the involvement of non-coding RNAs in tumorigenesis and tumor metastasis has been recently documented (***Croce, 2009; Shimono et al., 2009***). Non-coding RNAs can act as oncogenes or tumor suppressor genes, depending on the context.

Numerous genomic and experimental efforts have sought to refine the compendium of cancer driver genes, given their clinical relevance for cancer (***Barretina et al., 2012; Gonzalez-Perez et al., 2013; Ding et al., 2014***). However, in spite of immense efforts, evidence suggests that many uncharacterized TSGs and OGs exist. Perhaps most notably, down-sampling analysis of nearly 5,000 tumor genomes predicted the existence of hundreds of elusive driver genes mutated at intermediate and low frequencies (***Lawrence et al., 2014; Halliday et al., 2018***). As mutations do not occur evenly across the genome, mutation frequency is not perfectly correlated with driver gene potency (***Lawrence et al., 2014***). Thus, infrequently mutated driver genes can potentially have strong phenotypes. In fact, there are sequenced tumors that lack only a single mutation in characterized driver genes (***Imielinski et al., 2012***). Several computational approaches have been employed to detect infrequently mutated or rare driver genes. Analysis of mutation patterns, rather than frequency, circumvents sample size issues to some extent (***Schroeder et al., 2014; Vandin et al., 2912***), although drivers with atypical mutational patterns may be missed by such strategies. Alternatively, dimensionality reduction from genes to gene clusters or pathways can be used to address statistical power limitations, such as the cost of bias resulting from incomplete knowledge of protein networks (***Leiserson et al., 2015; Ciriello et al., 2012; Wang et al., 2018***). Finally, pan-cancer project findings can be used to examine similarities and differences among the genomic and cellular alterations found across diverse tumor types, thus increasing sample size.

In this study, we used naïve Bayes computation (NBC) to classify cancer genes as OGs, TSs or drivers/passengers. We addressed cancer mutations, as well as the cancer fusions collected in the ChiTaRS-3.1 database (***Gorohovski et al., 2017***). Moreover, we used the protein-protein interaction data generated by our previously developed tool ChiPPI (***Frenkel-Morgenstern et al., 2017***). ChiTaRS-3.1 contains comprehensive information about fusion genes and transcripts. Similarly, our customized Chimeric Protein-Protein-Interactions (ChiPPI) tool uses domain-domain co-occurrence scores to identify interactors of fusion. Gene fusions are essential diagnostic biomarkers that play important roles in various hematological disorders, as well as in sarcomas, along with solid tumors (***Mitelman et al., 2007***). In cancer, fusions are mostly produced by chromosomal translocations (***Frenkel-Morgenstern et al., 2012***). Moreover, information concerning fusion interactions is essential for analyzing their dynamics. This is particularly true in humans, where the complexity of protein-protein interactions (PPI) networks is baffling, given the ~25,000 protein-coding genes and the ≥100,000 cellular components that serve as nodes in the interactome.

In the present study, we used TCGA data comprising 25 cancer types with 3,091 fusions for the final testing of our method of TS and OG classification within cancers of interest. We initially classiy the cancer types into three broad categories, namely, leukemia and lymphoma (LL), sarcoma (SC) and solid tumors (ST). We also performed a survey of tumor suppressors and oncogenes from the perspectives of somatic mutations and network properties for 691 cohort cases in the TCGA dataset. In addition, we compared our method with well-established methods, such MutSigCV, OncodriveCLUST, Oncodrive-FM, 20/20+, ActiveDriver, MuSiC, TUSON, OncodriveFML, and found that our algorithm scored better score than these methods, with an overall accuracy 92.5% in identifying TSs and 94.2% in identifying OGs. ***Figure 1*** illustrates an overview of our methodology (discussed later) and ***Figure 2*** provides a more technical background of our algorithm (discussed later).

**Figure 1:**
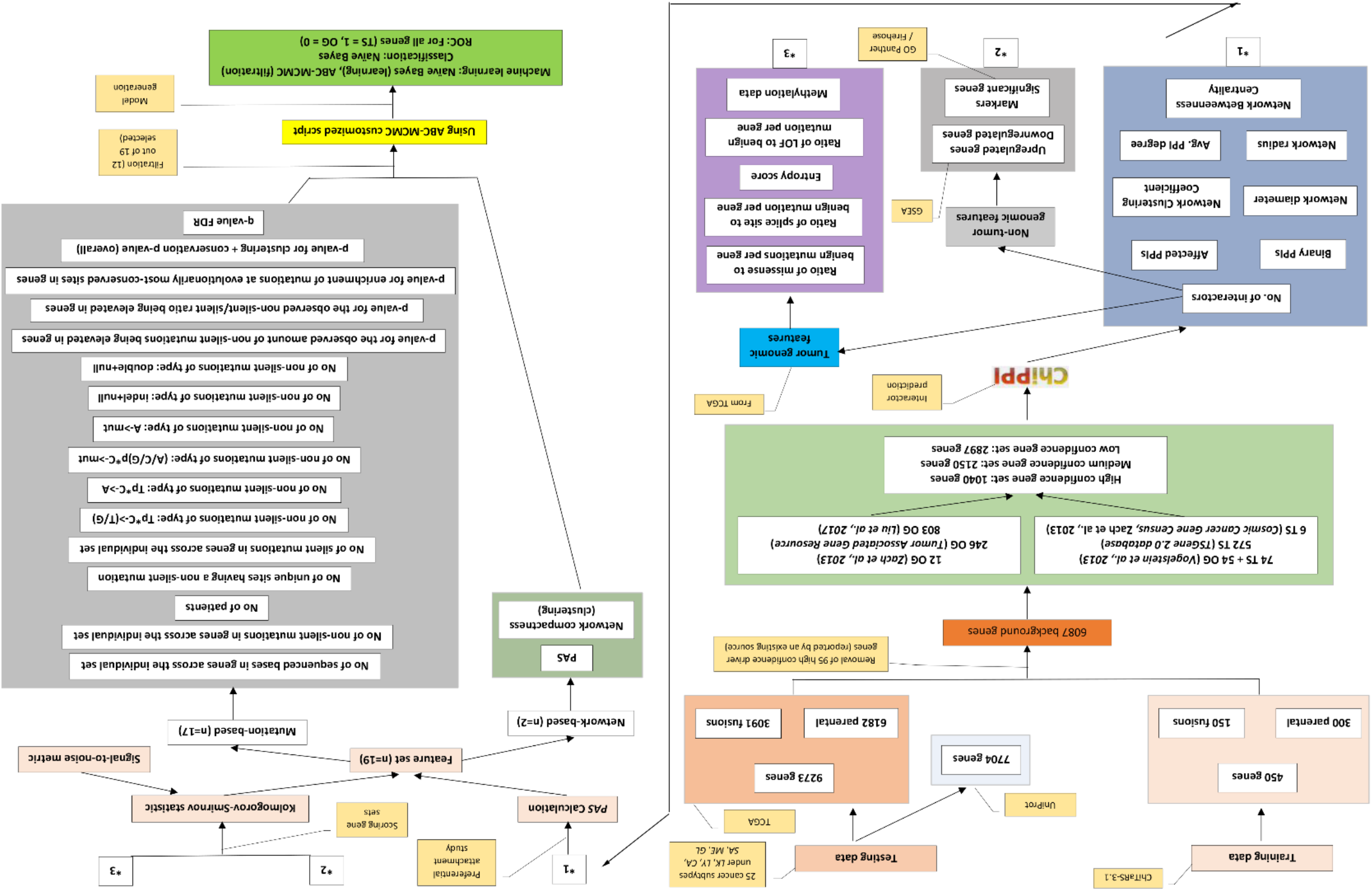
Algorithm of ABC-MCMC. The process initiates with a training phase, followed by testing, background gene selection, high-, medium-, low-confidence gene set definition, ChiPPI, PAS calculation, tumor and non-tumor feature assessement, scoring, filtration and final feature selection.

**Figure 2:**
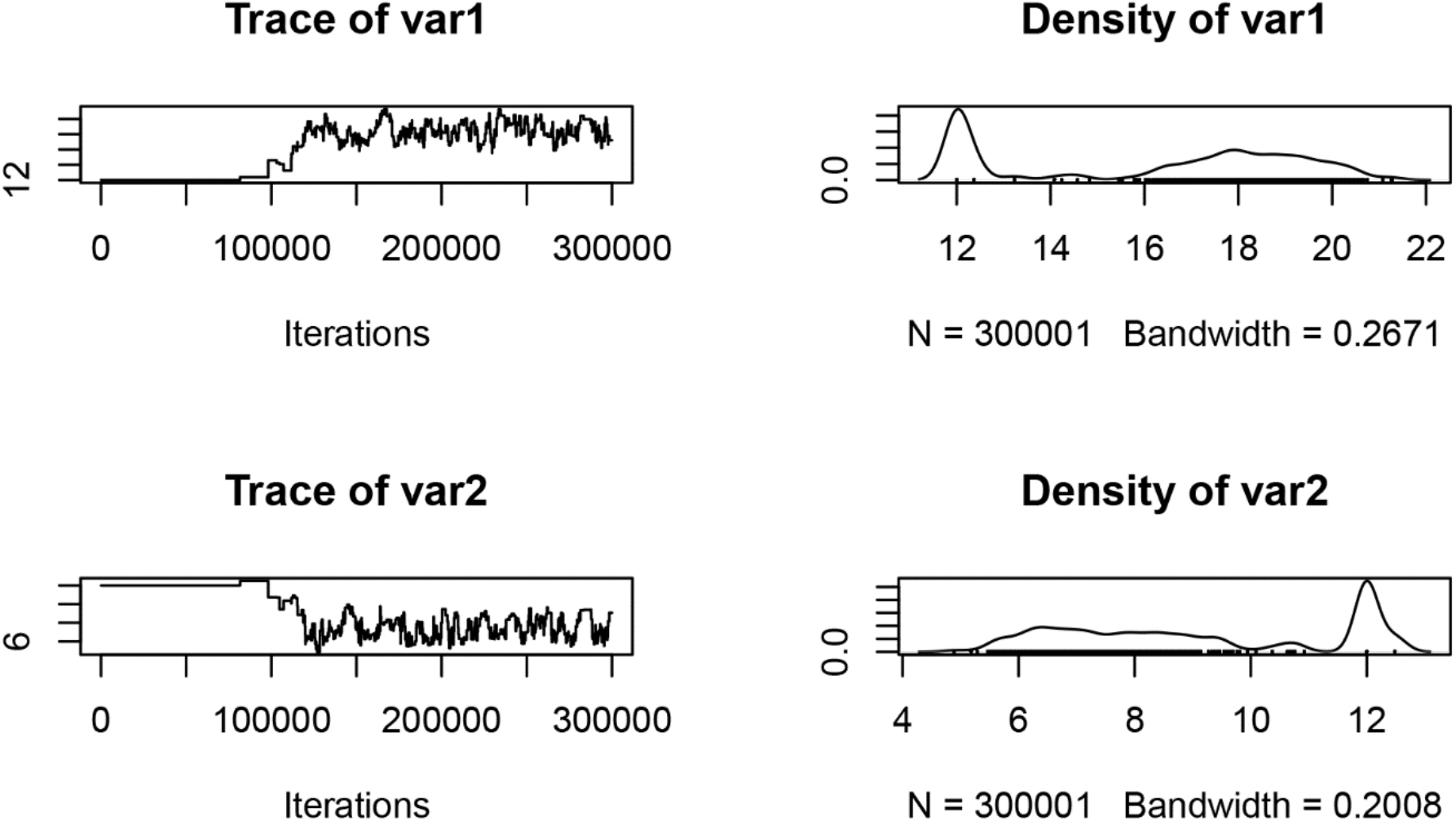
Trace and density charts corresponding to feature values: Number of non-silent mutations in genes across the individual set and number of unique sites having a non-silent mutation. The other selected features are: number of non-silent mutations of type Tp*C->(T/G); number of non-silent mutations of type Tp*C->(T/G); number of non-silent mutations of type Tp*C->A; number of non-silent mutations of type (A/C/G)p*C->mut; number of non-silent mutations of type A->mut; number of non-silent mutations of type: indel+null; p-value (overall); p-value for the observed amount of non-silent mutations being elevated in genes; p-value for the observed non-silent/silent ratio being elevated in genes; and p-value for enrichment of mutations at evolutionarily most-conserved sites in genes.

## 2 Results

### 2.1 Analysis of cancer-specific TS and OG clusters shows that cancer mutations in OGs are found in highly connected networks

Cancer-centric TS and OG clusters were generated using NBC for bladder urothelial carcinoma (BLCA), breast invasive carcinoma (BRCA), cervical squamous cell carcinoma and endocervical adenocarcinoma (CESC), colon adenocarcinoma (COAD), lymphoid neoplasm diffuse large B-cell lymphoma (DLBC), glioblastoma multiforme (GBM), head and neck squamous cell carcinoma (HNSC), kidney chromophobe (KICH), kidney renal clear cell carcinoma (KIRC), kidney renal papillary cell carcinoma (KIRP), brain lower grade glioma (LGG), liver hepatocellular carcinoma (LIHC), lung squamous cell carcinoma (LUSC), ovarian serous cystadenocarcinoma (OV), pancreatic cancer (PACA), prostate adenocarcinoma (PRAD), sarcoma (SARC), skin cutaneous melanoma (SKCM), stomach adenocarcinoma (STAD), thyroid carcinoma (THCA), and uterine corpus endometrial carcinoma (UCEC), respectively. These data contain potential overlaps, with the total number of unique genes being 16,977 for all cancers. Using NBC, we obtained a group of probability “models” with parameters that were benchmarked corresponding to biological target. This, on one hand, requires identification of a suitable parameter *k* and corresponding clustering of gene sets into non-overlapping gene subsets {*s*_1_, *s*_2_,…, *s*_*k*_}, represented by partition *S* of the gene set. On the other hand, the probability model structure captures a division of gene sets into biologically meaningful gene sub-sets, which are here defined as non-informative and informative, depending on whether or not they provide information about the clustering matrix. Thus, given the completely different 12 features of gene sets for the 21 cancers sets, NBC achieved high efficiency.

As an example, ***Figure 3*** illustrates some important OGs (e.g., ARG2, CD28, ATM) and TSs (e.g., HERC5, ATP5O) from DLBC (L**Y**). It can be seen that NBC was able to segregate these gene sets into Tumor-suppressors (TSs), Oncogenes (OGs) and Drivers (DRs)/Passengers (Pas), based on the 12 features considered (listed in the Methods section). Oncogenes drive abnormal cell proliferation as a consequence of genetic alterations that either increase gene expression or lead to uncontrolled activity of the oncogene-encoded proteins. Tumor suppressor genes represent the opposite side of cell growth control, normally acting to inhibit cell proliferation and tumor development. Mutations that provide a selective growth advantage, and thus promote cancer development, are termed driver mutations, and those that do not are termed passenger mutations. Likewise, taking a literature-based approach (***Tagore et al., 2018***), we collected 3,190 proteins belonging to OGs and/or TSs in Leukemia (LK), Lymphoma (LY), Melanoma (ME), Glioblastoma (GL), Sarcoma (SC) and Carcinoma (CA). Out of these, in Leukemia (LK), Lymphoma (LY), Melanoma (ME) and Glioblastoma (GL) (272 OGs, 238 TSs) (Table S3, Suppl. data), Sarcoma (SC) (385 OGs, 329 TSs) (Table S4, Suppl. data) and Carcinoma (CA) (299 OGs, 200 TSs) (Table S5, Suppl. data) are known from the literature (***Deng et al., 2018***). Furthermore, the number of DRs/PAs were found to be 114 (LK, LY, ME, and GL) (Table S6, Suppl. data), 312 (SC) (Table S7, Suppl. data) and 409 (CA) (Table S8, Suppl. data), respectively. NBC predicted novel OGs and TSs as follows: 70 (OGs), 45 (TSs) in LK, LY, ME, and GL; 200 (OGs), 112 (TSs) in SC; and 209 (OGs), 200 (TSs) in CA from previously unclassified data, using the 12 selected features (see Methods section). Cancer-centric clusters for other cancers are also presented (Figures S1-S15, Suppl. Data).

**Figure 3:**
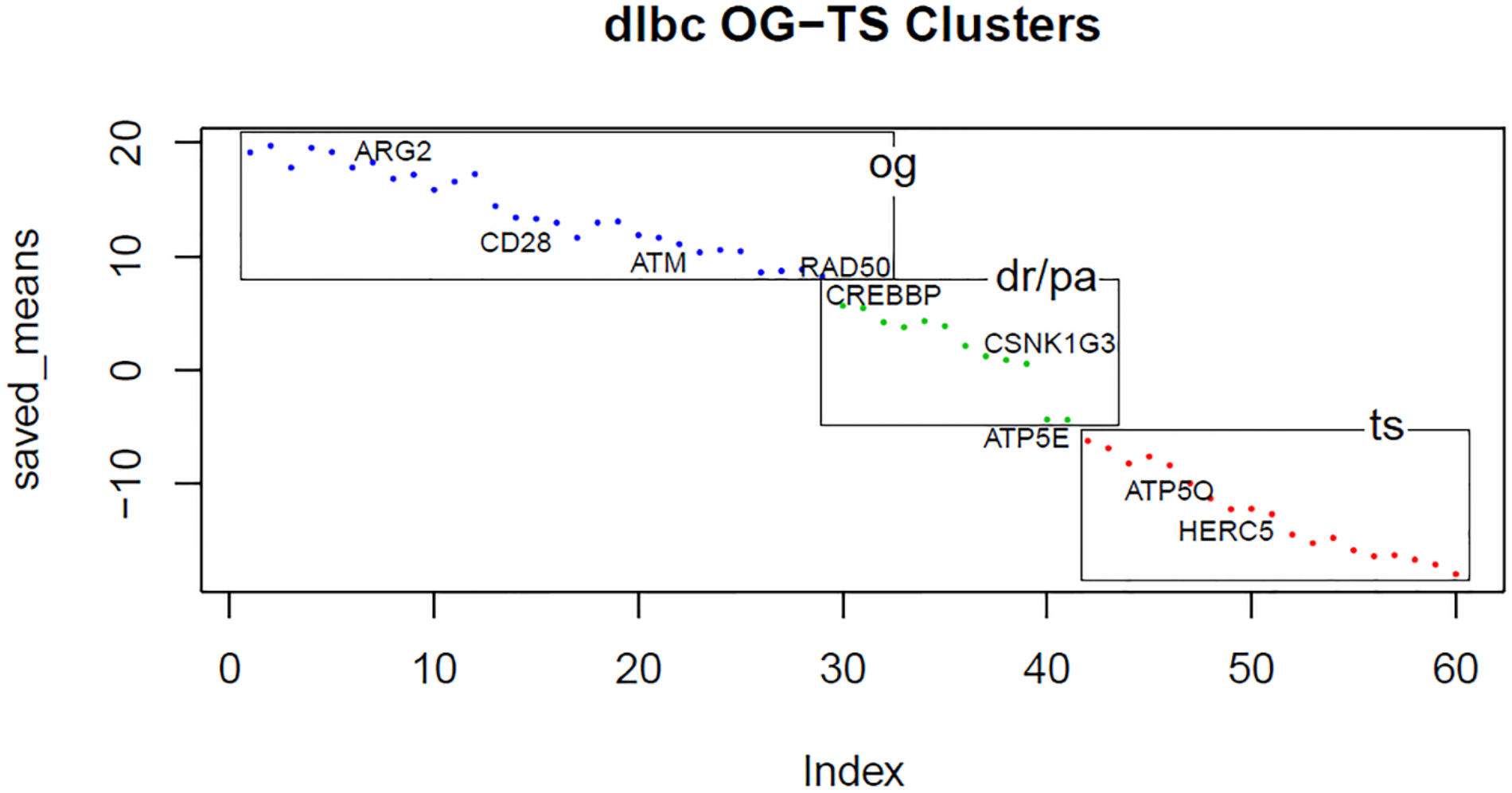
Some important OGs and TSs from DLBC (LY): ARG2, CD28, ATM, RAD50 (OG); CREBBP, CSNK1G3, ATP5E (DR/PA); ATP5Q, HERC5 (TS). It can be observed that OGs and TSs form separate clusters, whereas DR/PA proteins lie in-between them in a different group.

We further developed a novel network parameter called the preferential attachment score (PAS), which we incorporated during the clustering procedure as follows. We use *essential communities’* for identifying recurrent modules that participate in major PPIs using *preferential attachment*. This is defined as the attachment specificity of an incoming protein to an essential community. We found that the PAS could be used to separate OGs from TSs, with values ranging from 0.2 to 11.8 reflecting OGs and from 60.8 to 308.3 reflecting TSs (Table S10, Suppl. data). We found great variance in PAS scores <u>i</u>n LK, LY, ME, and GL, scores ranged from 1.01 - 9.93 (OGs) and 80.3-240.5 (TSs), in SC from 0.6-11.8 (OGs) and 60.8-308.3 (TSs), and in CA from 0.2-10.8 (OGs) and 80.3-200.5 (TSs), respectively. We found that from the 6,087 proteins initially considered, the highest number of OGs and TSs classified belonged to CA, followed by ST and LK, LY, ME, and GL. We got similar results considering DRs/PAs. These results indicate that due to the lower number of non-clustered proteins belonging to SC and CA, more unclassified functions are prevalent. This results in more OGs and TSs than already classified in LK, LY, ME, and GL. These results indicate that PAS can be used as a good parameter for segregating OGs and TSs based on network and mutational features.

### 2.2 LK and LY contain high numbers of communities due to the compact architectures of their networks

Our goal was to understand whether network communities have any role to play in the architecture underlying a network. For this purpose, we defined communities as those whose size (*m*) was >3 and the number of interactions for each participant in the community was *m*-1. We assumed that more clustered proteins would have higher numbers of interactions. We found that communities existed in LK, LY, ME, GL, SC and CA that have an average PPI score that was less than that of the higher PPI-scoring clusters. These results indicate that communities in all major groups of cancers make major contributions in determining the overall architecture of the networks. For example, in LK, LY, ME, and GL, we could identify communities for 44 fusion PPI networks, whereas in five cases, communities could not be found, due to their less compact PPI architecture. Similarly, in SC, we have identified communities for 46 fusion PPI networks, while in two cases, communities were not found (CIC-DUX4, NR6A1-TRHDE). In CA, we identified communities for 35 fusion PPI networks.

We used this connectivity association to distinguish between interactions in LK, LY, ME, GL, SC and CA PPI networks based on PAS values to reveal community attachment vertices. For example, in LK, LY, ME, and GL, we detected community attachment vertices that included KMT2A, HDAC2, SMARCC1, POLR2A, SMARCA2, CREBBP, and SIN3A (KMT2A-MLLT10). All these proteins contribute to forming communities and thus help in defining overall network structure. We also observed that of 50 fusions in LK, LY, ME, and GL, some common attachment vertices were KMT2A, CREBBP, SIN3A, HDAC2, NTRK1, TP53, SMARCA2, HDAC1 and EP300. Similarly, in SC, we found the attachment vertices NUP153, VCP, KPNB1, ELAVL1, CUL3, NTRK1, C1QBP, and EIF4B (PAPPA-NUP107). Finally, in CA, the attachment vertices CTBP1, CTBP2, KAT2B, CTBP1, CTBP2, CDC23, KAT2B in BCAS4-BCAS3 were noted (Table S11 in Suppl. data). These results indicate that the number of communities in LK and LY were elevated due to the compact architectures of their networks.

We also found that the number of communities were higher in LK, LY, ME, and GL than in SC and CA. This could be because of the presence of higher numbers of connected clusters in LK, LY, ME, and GL, as compared to SC and CA, where the number of proteins with fewer numbers of interactors was greater. Due to the higher number of communities in LK, LY, ME, and GL, open links were fewer, resulting in lower occurrences of lesser-interacting proteins. Furthermore, in SC, the number of open links was augmented, resulting in more occurrence of lesser-interacting proteins that tend to attach to these communities. Finally, in CA, the number of communities was significantly reduced, due to their smaller sizes. These results indicate that the structure of a PPI network depends on the existence of community-based modules, which further increase overall compactness.

### 2.3 Differential gene expression analysis across 25 cancers shows that genes are down-regulated during metastasis

Our goal here was to identify the types of proteins that show altered patterns of encoding gene expression in cancer. For this purpose, we performed gene-set enrichment analysis (GSEA) on all genes across 25 cancers and identified the top 30 up-regulated, as well as marker genes. The number of OGs that were up-regulated were comparatively higher in 13 cancers (DLBC, SARC, BRCA, COAD, HNSC, KIRC, LIHC, LUSC, OV, PACA, STAD, THCA, and SKCM), with the highest numbers being seen in SARC (20%), LUSC (20%), PACA (17%), and THCA (20%). Likewise, up-regulated TSs were found in three cancers, KICH (13%), UCEC (13%) and LGG (10%), whereas the occurrence of up-regulated OGs and TSs were equal in four cancers (BLCA, KIRP, PRAD and GBM). Figure 4 illustrates a heatmap depicted up-regulated genes across DLBC (LY). It can be seen that the top 30 up-regulated genes are correlated based on their *fc* and *log2fc* values. These results indicate that the number of up-regulated OGs was higher in major cancer types, relative to down-regulated OGs.

**Figure 4:**
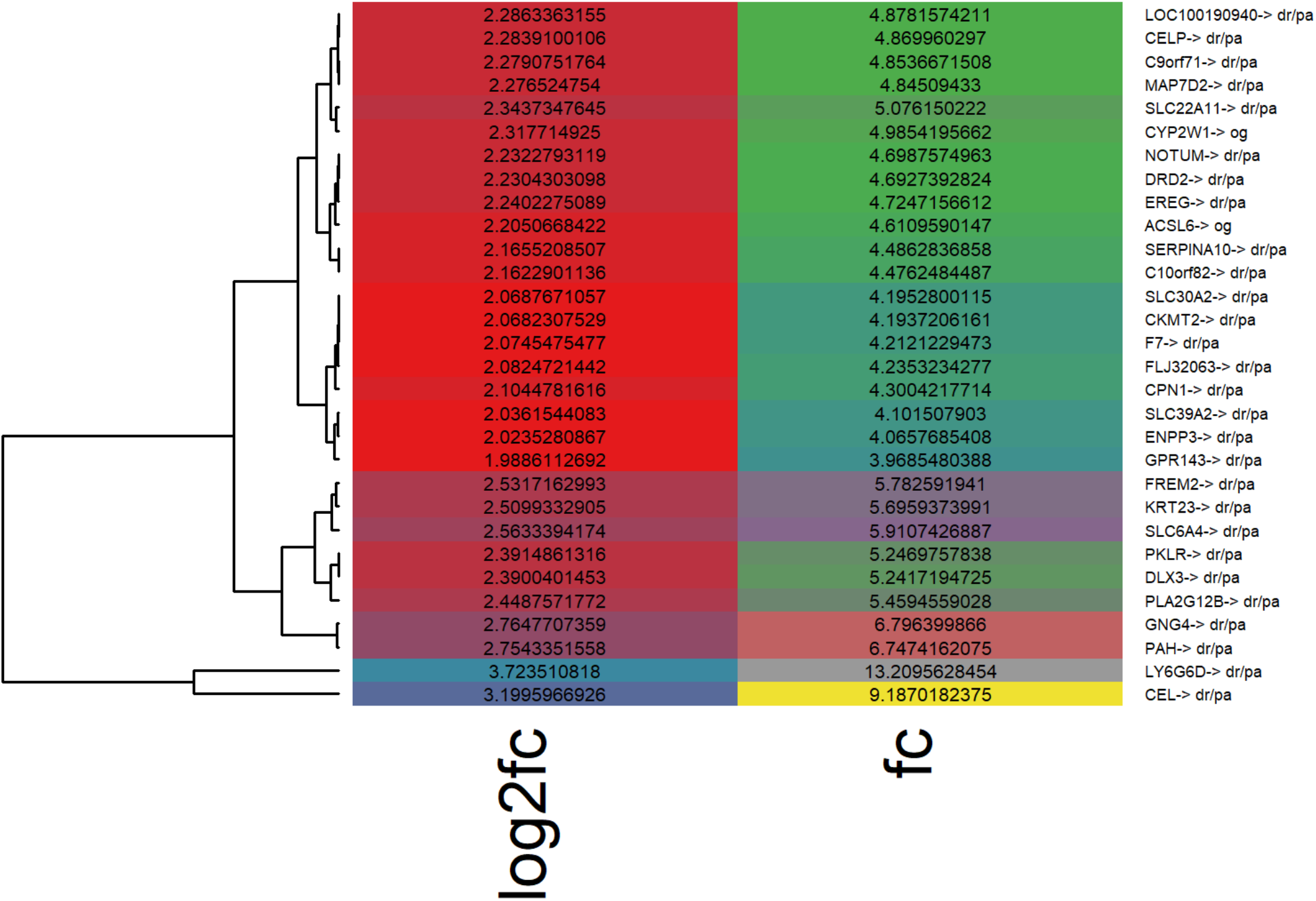
Heatmap for upregulated genes across DLBC (LY), namely, CYP2W1, ACSL6 (OG); FREM2, CEL (DR/PA). The top 30 up-regulated genes are correlated based on their fc and log2fc values. Here, LY6G6D is highly up-regulated, followed by CEL, GNG4, PAH, and SLC6A4.

Moreover, Figures S16-S36 (in Suppl. data) illustrate heatmaps for biomarkers across 21 cancer types. We found that the distribution of OGs was comparatively higher in 14 cancers (DLBC, SARC, BLCA, BRCA, COAD, HNSC, KICH, KIRC, KIRP, PACA, PRAD, STAD, SKCM, and GBM), with the highest levels being seen in COAD (17%), KICH (13%), SKCM (13%), and GBM (17%). Likewise, TSs markers were found in LIHC (6%), THCA (6%), and UCEC (6%), whereas the occurrence of marker OGs and TSs were equal in CESC, LUSC and OV.

Similarly, Figure 5 illustrates the heatmap for down-regulated genes across DLBC (LY). The number of OGs that were down-regulated were comparatively more in 13 cancers (DLBC, SARC, BLCA, BRCA, COAD, KICH, KIRC, KIRP, LIHC, PRAD, STAD, UCEC, and SKCM), with highest values being seen in BLCA (13%), PRAD (17%), and STAD (23%). Likewise, down-regulated TSs were found in HNSC (10%), LUSC (10%), THCA (6%) and GMB (6%), whereas the occurrence of down-regulated OGs and TSs were equal in CESC, OV, and LGG.

**Figure 5:**
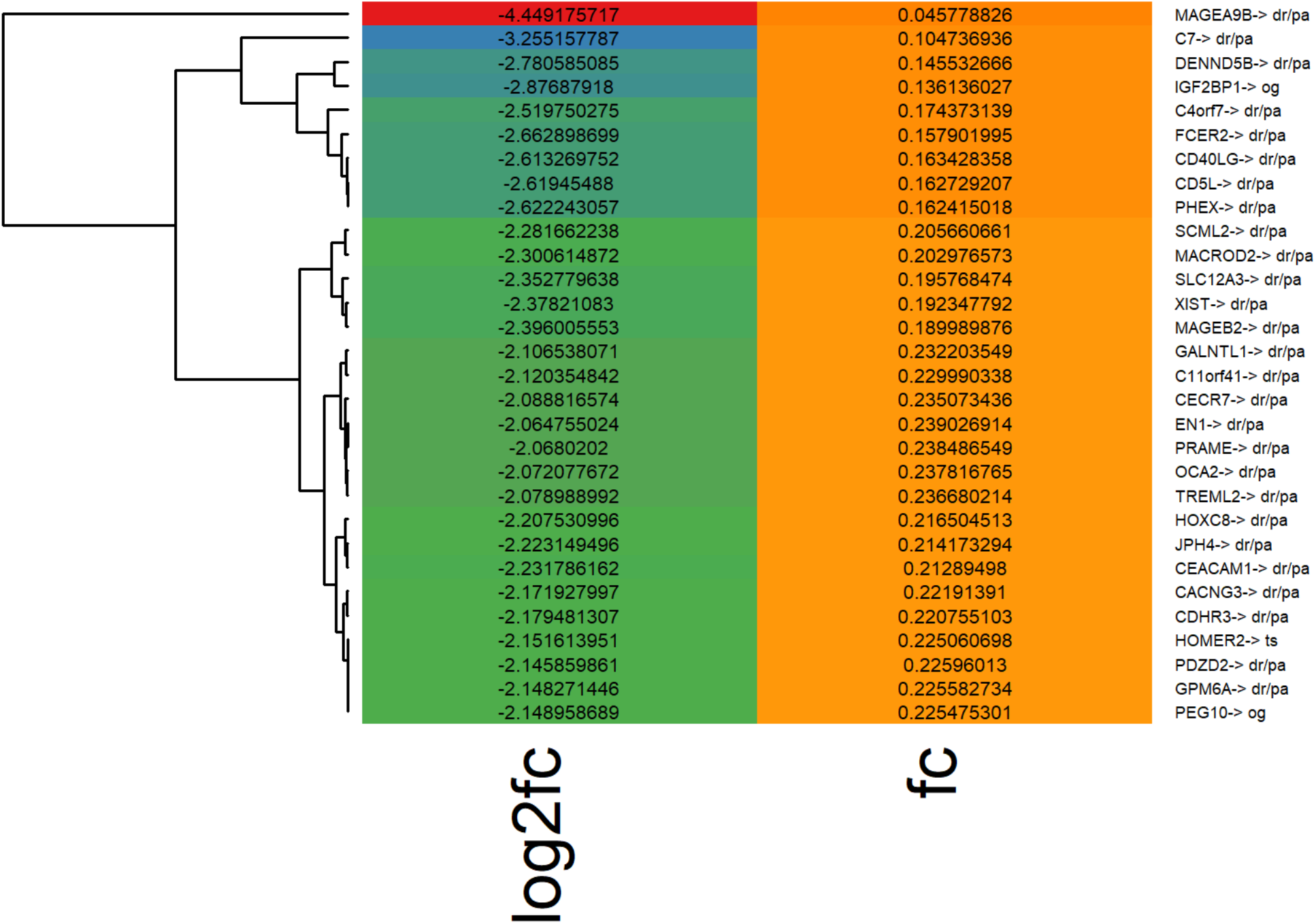
Heatmap for down-regulated genes across DLBC (LY), namely, IGF2BP1, PEG10 (OG); HOMER2 (TS); TREML2 (DR/PA). Here, MAGEA9B is highly down-regulated, followed by C7, IGF2BP1, DENND5B, and FCER2.

Finally, Figures S37-S57 (Suppl. data) illustrate heatmaps for significant genes across 21 cancers. We found that the distribution of OGs was comparatively wider in 12 cancers (DLBC, BLCA, COAD, KIRC, KIRP, LIHC, LUSC, PACA, PRAD, STAD, THCA, UCEC, and SKCM), with the widest distribution being seen in COAD (27%). Likewise, significant TS genes were found in SARC (6%), OV (10%), and LGG (17%), whereas the occurrences of OGs and TSs were equal in BRCA, HNSC, and GBM.

In all 21 cancers considered, we also identified *ARGFXP2, EHMT1, BCLAF1* and *DPF2* as being down-regulated. Similarly, we found only *AMBRA1* to be up-regulated in all cancers, while 188 common marker genes and 100 significant genes were also detected in all cancer types (Figures S58-S60, Suppl. Data). These results indicate that overall, differential gene expression analysis across 25 cancers showed that genes are down-regulated during metastasis. One major reason for this could be the overall compactness of the network, which increases the stress imparted onto individual genes.

### 2.4 Enriched pathway analysis shows that some pathways down-regulated in LK, LY were enriched in CA

For positive cross-verification, we examined how many pathways of the top 30 according to GSEA appear in the top 30 pathways previously studied. For negative cross-verification, we checked how many pathways of the top 30 according to GSEA were ranked below the top 100, listed according to the literature. Genes such as *PRL, C20orf103, RET, GGA2, ICSBP1, ITGA7, DAP, IRAK1*, and *PPARG* were over-expressed almost exclusively in MYST3-CREBBP in CLLE-ES (LK). Among those genes, *PRL* and *RET* have been reported as being involved in leukemogenesis (***Gerlo et al., 2005***). Interestingly, *STAT5A* and *STAT5B* were found to be significantly under-expressed in our MYST3-CREBBP, suggesting a negative regulatory effect between the prolactin and STAT5 proteins. GSEA results also showed that there were 21 and 31 up-regulated and 10 and 8 down-regulated pathways in LK and LY, respectively, whereas 53 up-regulated and 12 down-regulated pathways were noted in ME and finally, 27 up-regulated and 10 down-regulated pathways were detected in GL (Figures S61-S64, Suppl. data). In SC, 42 pathways were up-regulated and 10 were down-regulated (Figure S65, Suppl. data). In CA, genes assigned to the cell cycle, DNA replication, the spliceosome, proteasomes, mismatch repair, the p53 signaling pathway, nucleotide excision repair were up-regulated (Figures S66-S69, Suppl. data) (***Khetchoumian et al., 2008***). Additionally, the results indicated that down-regulated pathways in LK and LY were all enriched in CA (specifically, olfactory transduction, the renin angiotensin system and neuroactive ligand receptor interactions).

### 2.5 TSs and OGs have higher degrees of betweenness and lower clustering coefficients and shortest-path lengths than other proteins

The average degree of TSs was 87.92, significantly higher than that measured for other proteins (41.57; p = 3.76 × 10^−18^). Similarly, the average degree of the OGs was 76.49, also significantly higher than that measured for other proteins (45.21; p = 2.87 × 10^−14^). However, other than these network-based features, we did not observe any significant differences between TS and OG protein (p = 0.325). Figure 6 shows the degree and betweenness distributions for the top 30 gene sets for TSs, OGs and other proteins. The results for betweenness were consistent with those for degree. These observations indicate that TSs and OGs had the highest degree and betweenness in the human PPI network, as compared to other proteins. It has been observed that prominent cancer drivers have higher degrees in interaction networks (***Jonsson et al., 2006***). Likewise, the average clustering coefficient of TSs was 0.088, which was significantly lower than the values of other proteins (0.118; p = 0.015). Similarly, we found that the average clustering coefficient of OG proteins was 0.118, which was significantly lower than that for other proteins (p = 0.182). Figure 6 shows the distribution of clustering coefficient values, and the average value of each protein set. Finally, the average shortest-path distance of the TSs was 2.892, which was significantly shorter than that of other proteins (3.774; p = 3.03 × 10^−14^). Interestingly, the average shortest-path distance of TSs (2.892) was slightly lower than that of OGs (3.116; p = 0.040). Finally, Figure 6 also shows the distribution of shortest path distance values, and the average value of each protein set. These results indicate that TSs and OGs have higher degrees and betweenness and lower clustering coefficients and shortest-path lengths, as compared to normal proteins.

**Figure 6:**
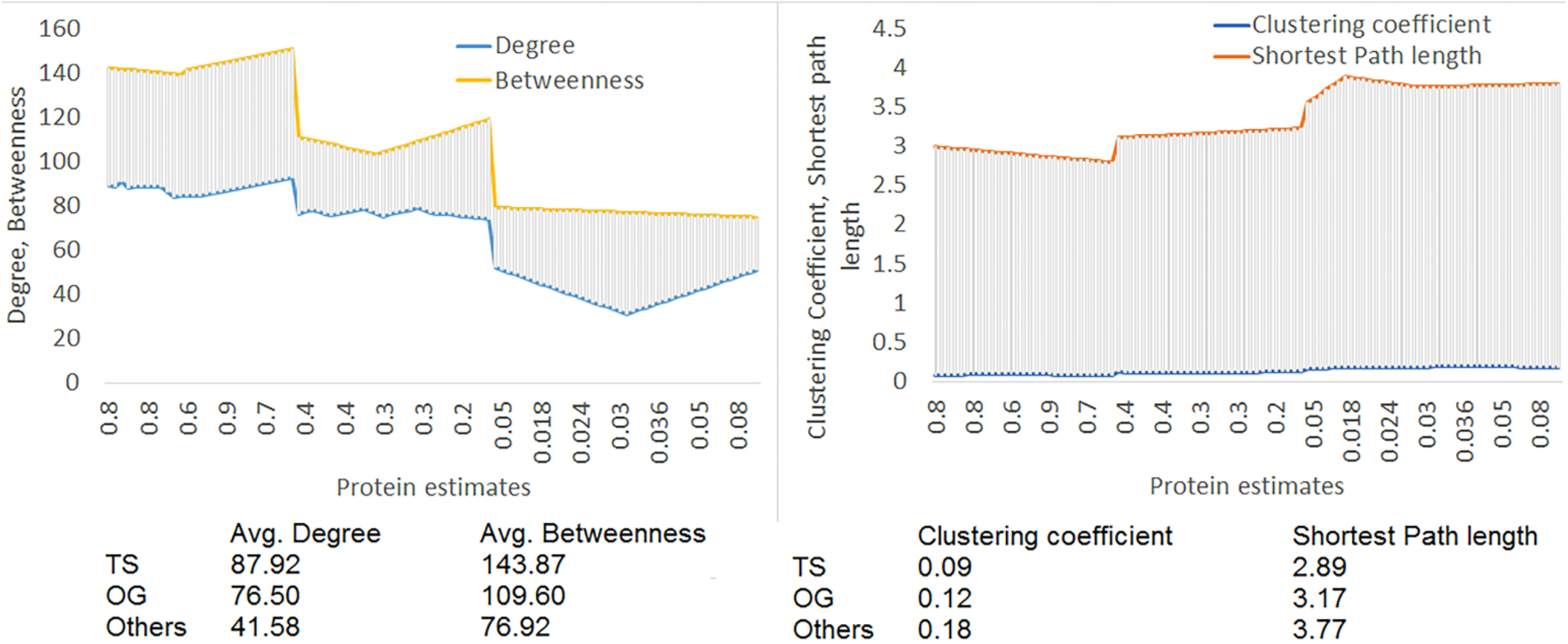
The degree and betweenness, clustering coefficients and shortest path distance distributions for top 30 gene sets for TSs, OGs and other proteins. The average degree and betweenness was higher in TSs, followed by OGs and others, whereas the average clustering coefficient and average shortest path length was lowest in TSs.

### 2.6 Hyper-methylation of TSs and their oncogenic behavior

Aberrant TS patterns of TS promoter methylation are known to be a frequent and occur early in carcinogenesis, while several tumor-specific alterations in DNA methylation have been found in the circulation of patients with different types of cancer (***Hattori et al., 2018***). In our study, we found that certain genes, namely, *ARHGEF12, BCR, CBL, CDKN1B, DMBT1, DNMT1, DNMT3A, ECT2, ETS2, ETV6, EZH2, FLT3, FOXL2, FOXO1, FOXO3, FOXO4, FUS, GLI1, HDAC1, HOPX, IDH1, KLF4, LHX4, LITAF, MAP3K8, MIR106A, MIR107, MIR125B1, MIR146A, MIR150, MIR155, MIR203A, MIR20A, MIR223, MXI1, NCOA4, NOTCH1, NOTCH2, NOTCH3, NPM1, NR4A3, PAX5, PHB, PML, PTPN11, RARB, RASSF1, RB1, RHOA, RUNX1, SALL4, SIRT1, SKIL, SPI1, SUZ12, TCF3, WDR11, WHSC1L1, WT1, YAP1* and *ZBTB16* showed oncogenic potential, mainly acting as TSs. Previously, it was noted that hyper-methylation of CpG islands located in the promoter regions of tumor suppressor genes is an important mechanism of gene inactivation, such as with the *Rb* gene, where the first discovery of methylation in a CpG island of a TS in a human cancer was made (***Esteller, 2002***). These results indicate that TSs show oncogenic behavior when hyper-methylated, as also described in the literature (***Hattori et al., 2018***).

### 2.7 Assessment studies using receiver operator characteristic (ROC) analysis

We used ROC analysis to test the performance of our method. In particular, we considered separating OGs and TGs from the DR/PA category. We obtained the best results for ROC of 0.9 and 0.89 for OGs and TSs, respectively. Known oncogenes, including *TRAF7* and *ALK*, were missed using previously published tools at the P < 0.05 cutoff but were easily detected by our method (Figure 7). The sensitivity of our method was highest for OG and TS classification in LK, LY, ME, and GL, followed by SC and CA (Figure 7). This was because of the existence of more unclassified and unknown data for CA than for SC or LK, LY, ME, and GL. These results indicate that sensitivity could be improved further with more training and once more test sets are considered.

**Figure 7:**
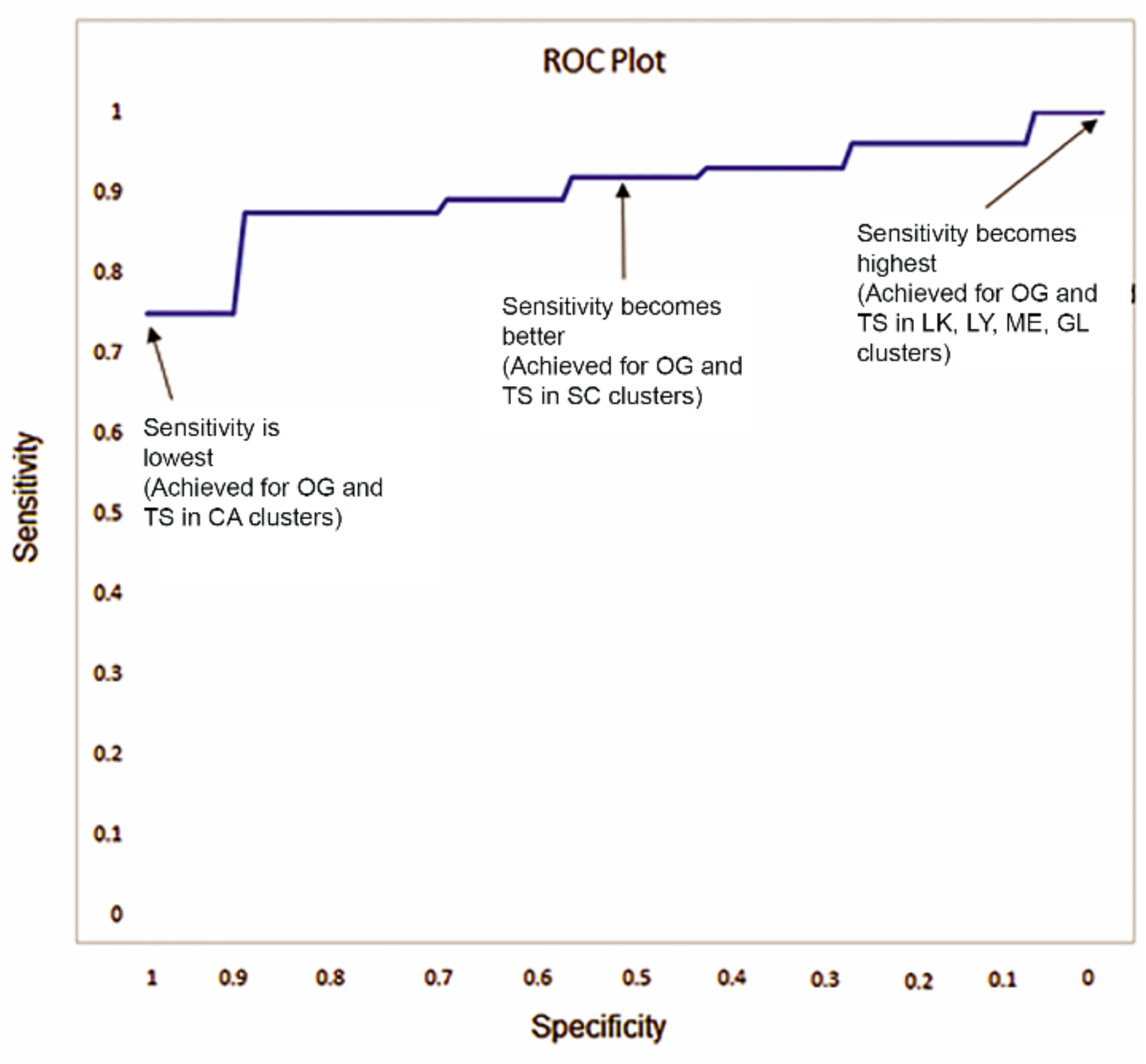
ROC plot of the NBC method illustrates that the sensitivity is highest for LK, LY, ME and GL clusters. We obtained the best results for ROC were 0.9 and 0.89 for OGs and TSs, respectively.

### 2.8 Existing algorithms and resources for studying cancer drivers

We performed complementary tests to evaluate the performance of NBC with respect to Random Forest 5 (RF5) (***Kumar et al., 2015***), Deep Learning Model (***Tavanaei et al., 2017***), OG/TS from ovarian cancer (***Wrzeszczynski et al., 2011***), MutSigCV (***Lawrence et al., 2014***), OncodriveCLUST (***Tamborero et al., 2013***), Oncodrive-FM (***Gonzalez-Perez et al., 2012***), 20/20+ (***Tokheim et al., 2016***), ActiveDriver (***Reimand et al., 2013***), MuSiC (***Dees, et al., 2012***), TUSON (***Davoli et al., 2013***) and OncodriveFML (***Mularoni et al., 2016***), each having different performances in classification tasks. The methodology of each of these algorithms differs from one another. Yet, none of these methods, except for RF5, separates likely oncogenes and tumor suppressors. Here, we discuss some critical observations related to these differences. The Random Forest method (***Kumar et al., 2015***) performs classification based on detecting deviations from expected patient distribution with Chi-square statistics, identifying truncation rates, calculating the probability of unaltered residues, and detecting deviation from expected cancer distribution. The Deep Learning Model (***Tavanaei et al., 2017***) uses a convolutional neural network to classify the feature map sets extracted from tertiary protein structures, wherein each feature map set represents some biological features, such as surface *C*_*α*_ atoms, residues based on polarity, hydrophobicity, or hydrophilicity, to name a few, associated with the atomic coordinates appearing on the outer surface of protein’s three-dimensional structure. ***Wrzeszczynski et al.**, 2011* used epigenetic and genomic features to examine possible OGs and TSs in primary ovarian tumors, addressing methylation and expression data for each gene under amplified or deleted copy number conditions, finally classifying them as OGs with low methylation and elevated expression or as TSs with lower copy number variation and hyper-methylation. MutSigCV (***Lawrence et al., 2014***) considers a list of mutations from a sample set, builds a model of the background mutation processes during tumor formation, and analyzes the mutations of each gene to identify genes that were mutated more often than expected by chance, given the background model. Oncodrive-CLUST (***Tamborero et al., 2013***) considers single-nucleotide protein-affecting mutations, wherein positions with some mutations above a background rate threshold are identified as potentially meaningful cluster seeds, followed by a grouping of these positions to form clusters. Oncodrive-fm (***Gonzalez-Perez et al., 2012***) detects likely driver genes and pathways in cancer through the analysis of functional mutations, prioritizing genes or pathways that show a bias toward the accumulation of functional somatic variants. The 20/20+ method (***Tokheim et al., 2016***) extends the ratiometric 20/20 rule, evaluates the proportion of inactivating mutations and recurrent missense mutations in a particular gene by automated integration of multiple features of positive selection. ActiveDriver (***Reimand et al., 2013***) identifies significantly mutated signaling regions in proteins, based on generalized linear regression and tests a pair of hypotheses for a given gene and its phospho-site region, using information on pSNV position within a phospho-site, protein structured and unstructured regions and cancer type-specific mutation rates. MuSiC (***Dees, et al., 2012***) separates significant events which are likely drivers for disease from passenger mutations present in mutational discovery sets using a variety of statistical methods. TUSON (***Davoli et al., 2013***) takes a parameterized approach to derive an overall significance and ranking for each gene, based on the calculation of a combined p-value (and q-value). Finally, OncodriveFML (***Mularoni et al., 2016***) estimates the accumulated functional impact bias of tumor somatic mutations in genomic regions of interest, both coding and non-coding, based on a local simulation of the mutational processes affecting it.

#### 2.8.1 Identifying gene overlaps among NBC and currently used methods

We identified the overlap of predicted OGs and TSs with respect to all known (1,450) and potential (15,527) cancer drivers, from the overall set of 16,977 genes for all 25 cancers. In this case, we also considered 1,450 driver genes that have known amplifications, deletions, frameshifts, and missense and nonsense mutations. We found that the OGs and TSs predicted by NBC MutsigCV and 20/20+ were enriched for driver genes and as part of TCGA (Table S12, Suppl. Data; Figure 8a). NBC, MutsigCV and 20/20+ predicted higher numbers of OGs and TSs, as compared to other methods from the training dataset. We next considered some smaller set of 1,450 driver genes, as supported elsewhere (***Zhao et al., 2016; Kim et al., 2018; Kumar et. al., 2015***), and found that the NBC results improved considerable (Figure 8b). We assumed that OGs or TSs predicted by at least two or more methods could be considered as potential drivers (***Tamborero et. al., 2013; Zhu et. al., 2015; Pavel et. al., 2016***).

**Figure 8:**
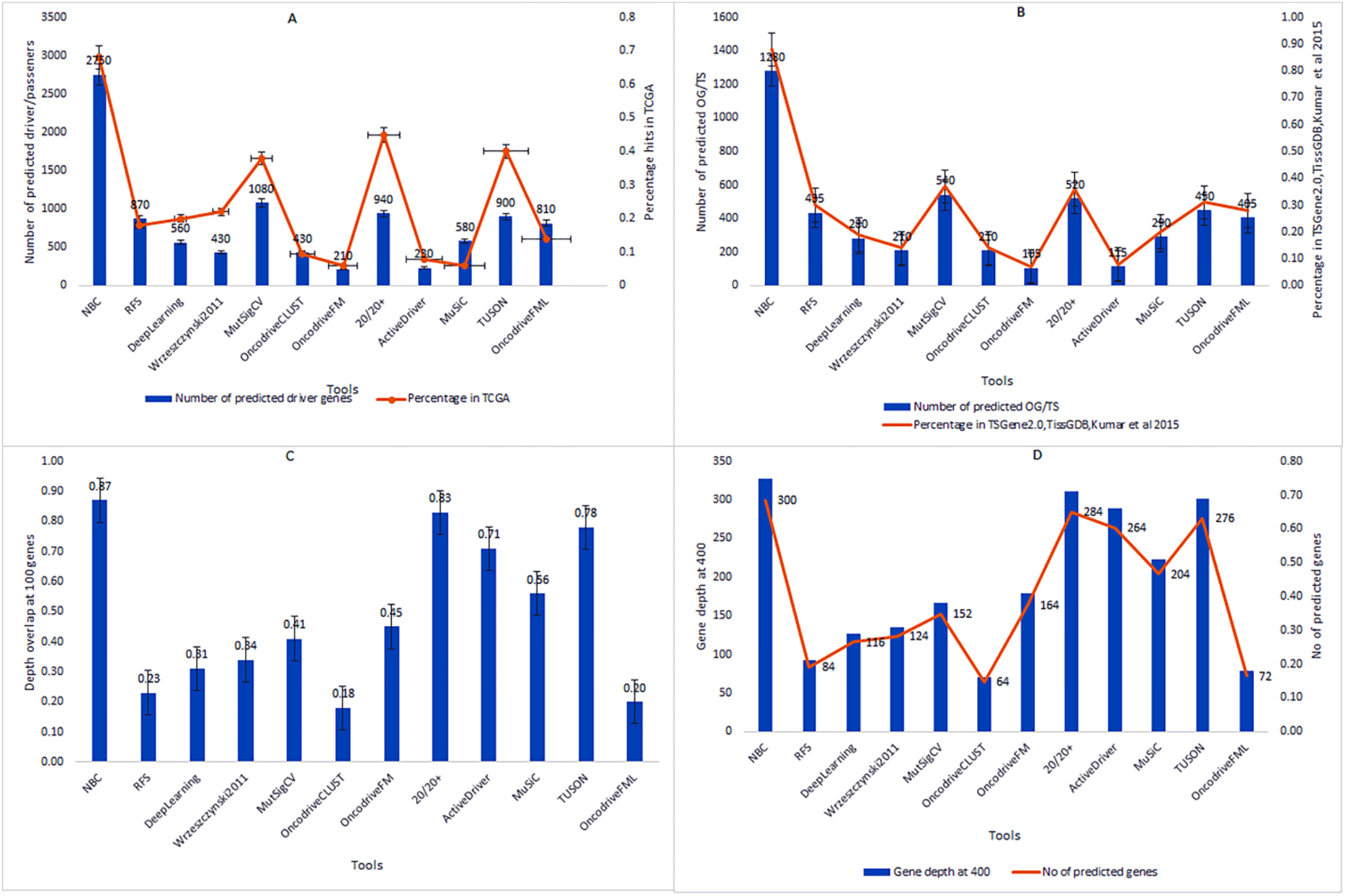
Depth overlap and driver gene predictions. (a) The OGs and TSs predicted by NBC, MutsigCV and 20/20+ were enriched for driver genes and part of TCGA. NBC, MutsigCV and 20/20+ predicted higher number of OGs and TSs, as compared to other methods from the training dataset. (b) For a smaller set of 1,450 driver gene (Zhao et al., 2016; Kim et al., 2018; Kumar et. al., 2015), the results of NBC improved considerably. (c) Examining the overlap of depth at 100 genes identified NBC, 20/20+, TUSON and ActiveDriver as having the highest consistency, defined by the Benjamini-Hochberg method. (d) The consistency of most methods decreased when gene depth increased beyond 400.

We also ranked the methods based on predicted gene overlaps from the *TSGene* (*Kumar et al 2015)* and found that NBC had made a substantial higher number of predictions, followed by MutsigCV and 20/20+ (***Figure 10a***). We also compared the methods based on whether the predicted OGs and TSs were unique for any one method or more and found that approximately 75% of OGs/TSs identified by NBC, MuSiC, OncodriveCLUST, and OncodriveFM were not identified by any of the other methods. Likewise, the overall prediction rate of unique OGs and TSs identified by NBC, MuSiC, OncodriveCLUST, OncodriveFM, and 20/20+ was found to be 93.05, 87, 72.25, 79.75, and 89, respectively. We also did literature-based analysis to validate these results and found accuracies of 91.2 (NBC), 88 (MuSiC), 71.3 (OncodriveCLUST), 75.2 (OncodriveFM), and 86.8 (20/20+), respectively (Table S13).

#### 2.8.2 Predicting OGs and TSs from drivers/passengers

Currently, no gold standard exists for addressing and estimating the accuracy and variability of predictions. We tested the 12 methods listed above on 25 repetitions of a random two-way split for all 15,527 genes, while maintaining sample proportions in the 25 cancer types. We, furthermore, ranked gene lists using p-values. For fair comparison, we considered those methods predicting that many drivers would be less likely to have consistent rankings than those predicting that only a few would. We used repeated measure ANOVA or Friedman tests to understand gene overlaps during random split. We chose the Friedman test, as the gene sets had been measured in different cancer types, were randomly chosen from all human protein-coding genes and were not necessarily normally distributed. Examining the overlap of depth of 100 genes revealed NBC, 20/20+, TUSON and ActiveDriver as having the highest consistency, as defined by the Benjamini-Hochberg method (Figure 8c). We also observed that the consistency of most methods decreased when gene depth increased beyond 400 (Figure 8d). For understanding the background mutation rate in genes and its potential influence on factors such as gene expression, we analyzed expected false-positive driver gene predictions. We set a critical value 0.1 for OG/TS prediction, based on the number of mutations required for a gene to be different from the background. We also modeled when the genes actually had mutation rates that varied around the critical value, and estimated false positives using the Bonferroni correction. We thus compared the number of mutations required to meet the critical value with different background mutation rates and different levels of variability for a sample size of 15,527 genes (Figure 9a). We observed that the number of false positives increased as the number of samples increased, whereas at the lower background mutation rates of 0.7 mutations per MB, both the number of false positives and the number of samples remained low (Figure 9b). For an intermediate background mutation rate of 5 mutations per MB, ~1,882 false positives were expected from 15,527 samples, whereas for a high background mutation rate (12 mutations per MB), both medium and high unexplained variability produced higher numbers of expected false positives (Figure 9b).

**Figure 9:**
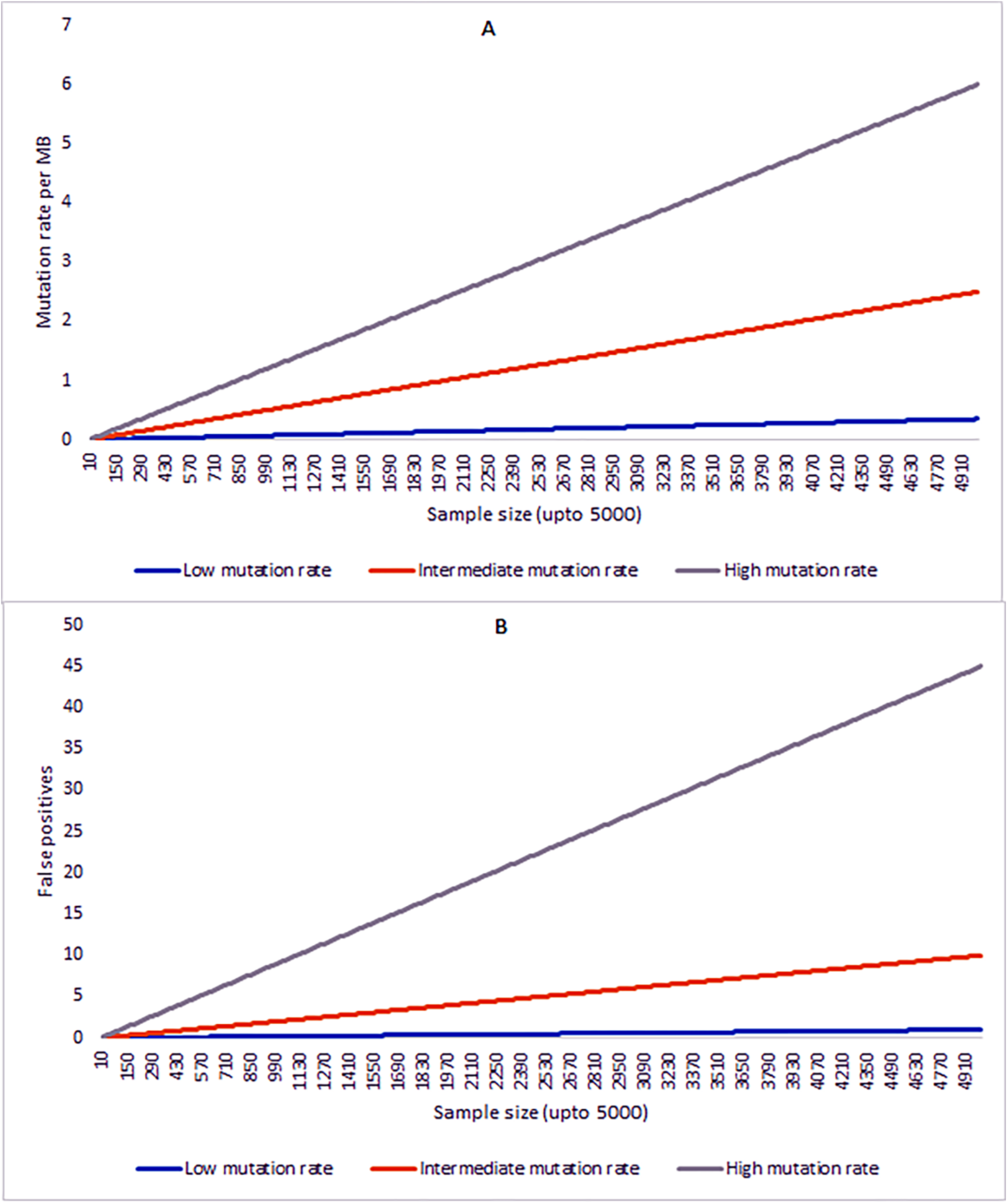
Background mutation studies. (a) The number of mutations required to meet the critical value with different background mutation rates and different levels of variability for the sample size of 15,527 genes. (b) The number of false positives increased as the number of samples increased, whereas at the lower background mutation rates of 0.7 mutations per MB, these remained low. (c) For an intermediate background mutation rate of 5 mutations per MB, ~1,882 false positives were expected from 15,527 samples, whereas for a high background mutation rate (12 mutations per MB), both medium and high unexplained variability produced higher numbesr of expected false positives.

**Figure 10:**
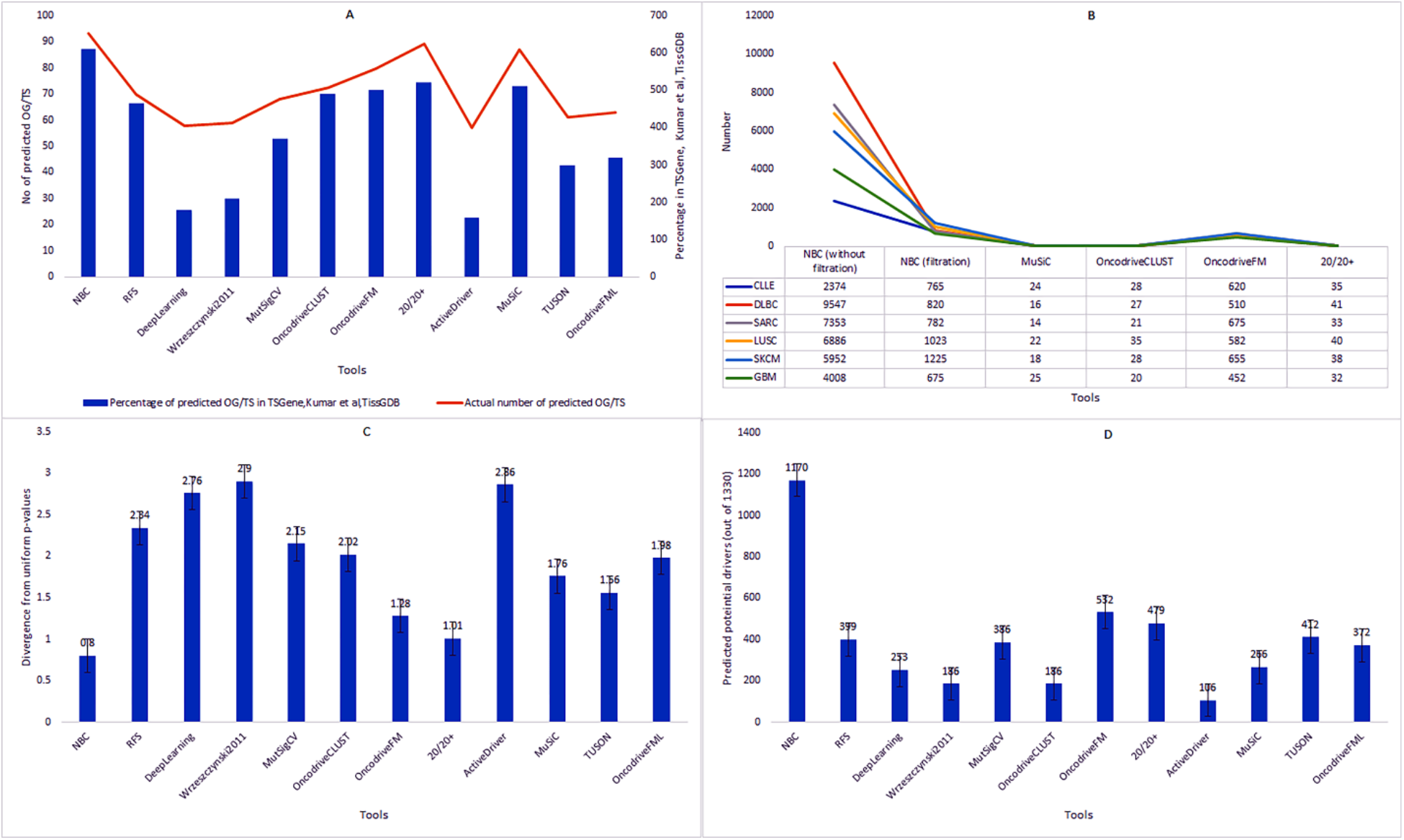
Ranking of methods. (a) Based on predicted gene overlaps from the TSGene, Kumar et al. (2015), and TissGDB data, we found that NBC had listed substantially higher number of predictions, followed by MutsigCV and 20/20+. (b) After filtering out common or overlapping genes, the final statistics was 765 (CLLE), 820 (DLBC), 782 (SARC), 1,023 (LUSC), 1,225 (SKCM), and 675 (GBM). SARC had the fewest predicted drivers/passengers (q ≤ 0.1), as predicted from CLLE. (c) For the current dataset of 15,527 genes, NBC, OncodriveFM and 20/20+ showed the least difference between observed and theoretical p-values. (d) For gene depth evaluation, rank depth was 1,330 genes (based on the highest number in SKCM) as the number of driver genes in a single cancer type. The most consistent methods were NBC, OncodriveFM and 20/20+, whereas the least consistent were MuSiC and OncodriveCLUST.

#### 2.8.3 Evaluation of genes based on cancer type

We evaluated the performance of NBC with various the other methods based on an analysis of 25 specific cancer types. The parameters used for evaluating performances were the number of driver/passenger genes predicted from the entire dataset and overall log-fold change. For this purpose, we considered six cancers from LK, LY, SA, CA, ME and GL, namely, CLLE (LK), DLBC (LY), SARC (SA), LUSC (CA), SKCM (ME), and GBM (GL), respectively. The number of predicted cancer-specific driver/passenger genes by NBC were 2,374 (CLLE), 9,547 (DLBC), 7,353 (SARC), 6,886 (LUSC), 5,952 (SKCM), and 4,008 (GBM), with multiple overlapping genes in these cancers. After filtering out common or overlapping genes, the final statistics were 765 (CLLE), 820 (DLBC), 782 (SARC), 1,023 (LUSC), 1,225 (SKCM), and 675 (GBM). SARC had the fewest predicted drivers/passengers (q ≤ 0.1), as predicted by CLLE (MuSiC = 24, OncodriveCLUST = 28, OncodriveFM = 620, 20/20+ = 35), DLBC (MuSiC = 16, OncodriveCLUST = 27, OncodriveFM = 510, 20/20+ = 41), SARC (MuSiC = 14, OncodriveCLUST = 21, OncodriveFM = 675, 20/20+ = 33), LUSC (MuSiC = 22, OncodriveCLUST = 35, OncodriveFM = 582, 20/20+ = 40), SKCM (MuSiC = 18, OncodriveCLUST = 28, OncodriveFM = 655, 20/20+ = 38), and GBM (MuSiC = 25, OncodriveCLUST = 20, OncodriveFM = 452, 20/20+ = 32), respectively (Figure 10b). Likewise, for the current dataset of 15,527 genes, NBC, OncodriveFM and 20/20+ showed the least differences between observed and theoretical P values (Figure 10c). For gene depth evaluation, we used a rank depth of 1,330 genes (based on the highest number in SKCM) as the number of driver genes in a single cancer type. The most consistent methods were NBC, OncodriveFM and 20/20+ (***Figure 10d***), whereas the least consistent were MuSiC and OncodriveCLUST. Overall, NBC predicted 6,087 potential cancer drivers for all cancer types from a total of 15,527 protein-coding genes.

#### 2.8.4 Overall accuracy of classification

Finally, we hypothesized that our model would be able to separate all three gene classes, namely, TSs, OGs and DRs/PAs from one another. Training was performed with five-fold cross-validation, with results averaged over 100 repetitions. Given the lack of agreement among these various methods, we compared p-values reported by each method to those expected theoretically. Such comparisons are often used in statistics and can indicate invalid assumptions or inappropriate heuristics. Theoretically, the p-value distribution should be approximately uniform after likely driver genes are removed. Therefore, we removed all genes predicted to be drivers by at least three methods after Benjamini–Hochberg multiple-testing correction (q ≤ 0.1). For quantifying the differences between the observed and expected p-values, we used log2-fold change. NBC and 20/20+ had log2-fold change values that were five-fold lower than the other methods. We also compared observed and theoretical p-values with quantile–quantile plots, which provide a detailed view of p-value behavior. NBC p-values had by far the best agreement with theoretical expectations across the entire range of supported values. For a significant p ≤ 0.05, OncodriveClust, OncodriveFM, OncodriveFML, ActiveDriver, and MuSiC substantially under-estimated p-values, whereas MutsigCV and 20/20+ substantially over-estimated them. Thus, NBC achieved better scores than all of the other methods considered (Table S14). When assessed for performance on each task separately, NBC was significantly better or not significantly different from the best individual tests. Therefore, our proposed NBC method has performed better than its counterparts in detecting TSs (0.925) and OGs (0.942).

## 3 Methods

### 3.1 Training datasets

We constructed a list of high-confidence protein-coding genes covering mutations and methylation-pattern gene sets. We used 25 different cancer types, including highly frequent cancers in both men and women. The list primarily includes partner interactors of fusions, especially those resulting from chromosomal rearrangements. The training data included 450 genes (for 150 fusions) from the ChiTaRS-3.1 database (***Gorohovski et al., 2017***). Since, our main analysis depends on classifying the interactor genes of fusions, we used ChiPPI (***Frenkel-Morgenstern et al., 2017***) for identifying the interactors. The chimeric protein–protein interaction (ChiPPI) algorithm (***Frenkel-Morgenstern et al., 2017***) uses a ‘domain–domain co-occurrence’ score to calculate PPI likelihood. This score is based on previous observations regarding the preferences of domain–domain interactions and co-occurrence methods. The final score for a fusion protein and its interactors combines all the preserved domains of the two parental proteins of the fusion. The PPI network for each fusion is built by combining all interactors of both parental proteins, as well as removing ‘missing’ interactors, lost due to deletion of parental protein domains. with 50 fusions belonging to each category (Table S9, Suppl. data). Figure 1 **NOW?** illustrates an overview of our methodology.

### 3.2 Background and test gene-set selection

We selected 9,273 genes (6,182 parental genes) to be used in the study of 25 cancer sub-types and 3,091 fusions from TCGA based on six broad cancer categories, namely, leukemia (LK), specifically, chronic lymphocytic leukemia – ES (CLLE-ES);, lymphoma (LY), specifically, lymphoid neoplasm diffuse large B-cell lymphoma – US (DLBC-US), and malignant lymphoma DE (MALY-DE), Sarcoma (SA), specifically, sarcoma – US (SARC-US), carcinoma (CA), specifically, bladder urothelial cancer – US (BLCA-US), breast cancer – US (BRCA-US), Cervical squamous cell carcinoma – US (CESC-US), colon adenocarcinoma – US (COAD-US), head and neck squamous cell carcinoma – US (HNSC-US), kidney Chromophobe – US (KICH-US), kidney renal clear cell carcinoma – US (KIRC-US), kidney renal papillary cell carcinoma – US (KIRP), liver hepatocellular carcinoma – US (LIHC-US), liver cancer – JP (LIRI-JP), lung squamous Cell Carcinoma – US (LUSC-US), ovarian Cancer – AU (OV-AU), pancreatic cancer endocrine neoplasms – AU (PACA-AU), prostate adenocarcinoma – US (PRAD-US), renal cell cancer - EU/FR (RECA-EU), gastric adenocarcinoma – US (STAD-US), head and neck thyroid Carcinoma US (THCA-US), uterine corpus endometrial carcinoma – US (UCEC), melanoma (ME), specifically, skin cutaneous melanoma – US (SKCM-US); and Glioblastoma (GL), specifically, brain glioblastoma multiforme – US (GBM-US), and brain lower grade glioma – US (LGG-US). We also selected 7,704 protein-coding genes from UniProt.

Of these 25 cancer gene sets, we included our analysis for 21 and excluded 4 (i.e., CLLE, MALY, LIRI, and RECA), due to unavailability of suitable mutational and methylation data. Our NBC approach requires a set of genes with ideally very limited cancer evidence to be used as background examples. Finally, we removed the 95 high-confidence driver genes that were reported in at least one source from the above. This resulted in 6,087 background genes used for classifier training.

### 3.3 High-, medium- and low-confidence gene sets

We initiated our methodology by including 71 TS and 54 OG mutation driver genes, as reported (***Vogelstein et al., 2013***). We also included 572 newly reported TSs from TSGene 2.0 (***Zhao et al., 2016***) and also supplemented these TSs with six genes, namely, *CDKN1B*, *FANCD2, FAT1*, *IKZF2, MYCN, TSC2* and *PARK*, as labeled by the Cosmic Cancer Gene Census (CGC) (***Futreal et al., 2004***) and by ***Zack et al. (2013)***. For OGs, we considered 12 genes, namely, *SOX2*, *CCNE1*, *BCL2L1*, *E2F3*, *CDK4*, *CDK6*, *NEDD9*, *IGF1R*, *PAX8*, *MCL1*, *ZNF217* and *TERT* (***Zack et al., 2013***). We then added 246 OGs as reported at the Tumor Associated Gene (TAG) resource. Fourth, we supplemented the testing data with 803 human OGs that included 698 protein-coding genes and 105 non-coding OGs (***Liu et al., 2017***). Next, we removed *MYCN* from our list as it may serve as both a TS and OG, according to the literature. Thus, we did not use *MYCN* as a driver gene in the high-confidence set. Finally, the resulting high-confidence set consisted of 495 TSs and 545 OGs. We also created medium- and low-confidence driver gene sets to determine NBC efficiency beyond high-confidence drivers (*n* = 1040). We also assumed that these gene sets may contain high false positive data. The medium-confidence set (*n* = 2,150) included all genes present in at least one of the following genomics-based sources: ***Zack et al. (2013)***, ***Lawrence et al. (2014)***, HotNet2 (***Leiserson et al., 2015***), OncoDriveClust (***Tamborero et al., 2013***), and ActiveDriver (***Reimand et al., 2013***). Finally, the low-confidence gene set (*n* = 2,897) includes genes that were not part of the high- or low-confidence sets.

### 3.4 Extracting features from the high-, medium- and low-confidence gene sets

ChiPPI (***Frenkel-Morgenstern et al., 2017***) was used to retrieve PPI data for all 6,091 fusions. PPI statistics, such as number of interactors, binary PPIs, affected PPIs, network diameter, radius, average degree of PPI, network average clustering coefficients and betweenness centrality, were calculated using ChiPPI. We also considered essentiality features for all genes. For instance, information with respect to coding sequence was determined as the longest isoform within the gene, as retrieved from UniProt (***Pundir et al., 2017***), whereas Ensembl (***Zerbino et al., 2018***) was used to retrieve the GC percent per gene. All data related to mutation and methylation were considered from TCGA for all 21 cancer types.

### 3.5 Tumor- and non-tumor-derived genomic features

For tumor data, we considered feature scores, such as ratio of missense mutations to benign mutations per gene, ratio of splice site mutations to benign mutation per gene, entropy score, and ratio of loss-of-function mutations to benign mutations per gene. These features were extracted from TCGA data. For non-tumor genomic features, we considered enrichment analysis, up-regulated and down-regulated genes (using GSEA) (***Subramanian et al., 2005***), enriched, and significant genes (using GO Panther) (***Mi et al., 2017***). For identifying significantly enriched pathways, we again considered GSEA. The Kolmogorov-Smirnov statistic was used for scoring gene sets, whereas a signal-to-noise metric was used to rank them. This signal-to-noise metric could be expressed as the amount of biological signal, relative to the amount of noise, and depends not only on the amount of noise, which could be a direct measure of quality, but also on the amount of signal. As an example, for a study *x*, if the data matrix is 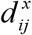, the Pearson correlation, 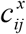, between genes *i* and *j* is described by Eq. 1,

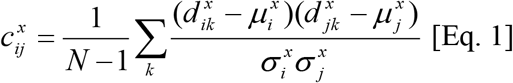

where 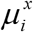 is the mean, 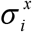 is the standard deviation and *c*_*ij*_ is the median of those correlations across all studies. Likewise, the signal-to-ratio, *λ*_*x*_, is described by Eq. 2,

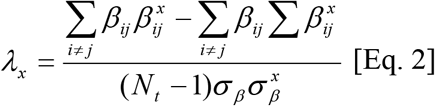

where *β*_*ij*_ is the inverse hyperbolic tangent for *c*_*ij*_, 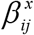, is the inverse hyperbolic tangent for 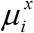, *N*_*t*_ is the total number of gene-gene correlations, *σ*_*β*_ is the standard deviation of gene-gene correlations for a median matrix and 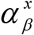 is the standard deviation of gene-gene correlations for *x*. Finally, using methylation data from TCGA, we extracted marker gene information.

### 3.6 Final feature selection and filtration

Filtration of most relevant features was performed using our customized R script, based on the ABC-MCMC algorithm. Approximate Bayesian computation (ABC) has been widely used in the scientific community to analyze population demography, growth rates, and time of divergence in the last decade. ABC-based methods approximate the likelihood function by simulations, the outcomes of which are compared with observed data. With the ABC rejection algorithm, a set of parameter points is first sampled from the prior distribution (***Csilléry et al., 2010***). Thus, ABC methods generate samples from a distribution which is not the true posterior distribution of interest, but rather a distribution which is hoped to be close to the real posterior distribution of interest. Likewise, there is a close connection between *likelihood free* Markov Chain Monte-Carlo (MCMC) methods and ABC. Consider the case of a perfectly observed system, so that there is no latent variable layer. Then, there are model parameters *θ*, described by a prior, *π*(*θ*), and a forwards-simulation model for data *x*, defined by *π*(*x* | *θ*). Thus, a simple algorithm for simulating from the desired posterior *π*(*θ* | *x*) can be, *first*, simulate from the joint distribution *π*(*θ* | *x*) by simulating *θ*^*^ ~ *π*(*θ*) and then *x*^*^ ~ *π*(*x* | *θ*^*^). This gives a sample (*θ*^*^, *x*^*^)from the joint distribution.

A simple rejection algorithm which rejects the proposed pair, unless *x* matches the true data *x*, clearly gives a sample from the required posterior distribution. The final feature set consisted of two network-based features (PAS and network compactness value) and 17 mutation-based features (number of sequenced bases in genes across the individual set, number of non-silent mutations in genes across the individual set, number of patients with at least one non-silent mutation, number of unique sites having a non-silent mutation, number of silent mutations in genes across the individual set, number of non-silent mutations of type Tp*C->(T/G), number of non-silent mutations of type Tp*C->A, number of non-silent mutations of type (A/C/G)p*C->mut, numbero of non-silent mutations of type A->mut, number of non-silent mutations of type indel+null, number of non-silent mutations of type double+null, p-value for the observed amount of non-silent mutations being elevated in genes, p-value for the observed non-silent/silent ratio being elevated in genes, p-value for enrichment of mutations at evolutionarily most-conserved sites in genes, p-value for clustering + conservation p-value (overall), q-value FDR).

A *community* is defined as a cluster of proteins that have a high average clustering coefficient and which is very compact. In our study, we identified *essential* and *non-essential communities*. We defined *essential communities* as those whose size is less than three and the number of interactions for each participant in the community generates a *clique*. Similarly, *non-essential communities* are those whose size is equal to or more than three, but the number of interactions for each participant in the *community* generates a *clique*. We used *essential communities* for further analysis as they participate in more PPIs for identifying whether proteins tend to interact with them in PPI networks using *preferential attachment*. The PAS for a protein is defined as the attachment specificity of an incoming protein to an essential community (given as Eq. 3).

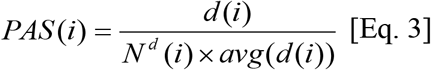

The ABC-MCMC algorithm identified ten possible feature combinations among the list of 19 possible features that could be further used for classification (Figure 2 **NOW?**). In Figure 2, we discuss the trace and density charts corresponding to feature values. The trace plot shows the values the parameter took during the runtime of the chain, whereas the density plot shows histogram of the values in the trace plot. It can be seen that the feature values are well distributed and follow a skewed Gaussian curve, thus enabling us to select these features for training. Thus, for training we concentrated on 12 features out of 19 (illustrated in Figure 2).

### 3.7 Model generation

#### Machine Learning

***NBC*** employs a naïve Bayes method for learning (***Friedman et al., 1997***), and an ABC-MCMC method for feature filtration (***Csilléry et al., 2010***).

#### Classification

Classification of genes as TSs, OGs, or drivers/passengers was performed using the naïve Bayes method. The naive Bayes method uses models that assign class labels to problem instances, represented as vectors of feature values, where the class labels are drawn from some finite set. Naïve Bayes classifiers assume that the value of a feature is independent of the value of any other feature, given the class variable. The advantage of the Naïve Bayes model is that it only requires a small number of training data to estimate the parameters necessary for classification. For each gene, the posterior probability of being a TS, OG, or driver/passenger gene was computed according to the feature set.

#### ROC step

ROC curves were calculated using the training dataset for 6,087 genes. TS labels were set to 1 and OG labels to 0. The detailed algorithmic descriptions are present in Mathematical Description of Methods (in Suppl. data).

## Discussion

The biggest application of next generation sequencing-based technologies is identifying novel cancer genes. This is a two-stage process, namely, *stage 1*, where cancer genes are separated from genes with passenger mutations; *stage 2*, where cancer genes are differentiated into likely tumor suppressors and oncogenes. Both stages are crucial because mechanism-specific predictions are needed to guide downstream analyses and experiments. Further, various oncogenic signaling pathways may act as drivers for tumorigenesis and can also induce events such as chromosomal instability. This is important since they disrupt the events required for accurate chromosome segregation during mitosis by decreasing the rate of correction of k-MT attachment errors and/or increasing the rate of formation of those errors through extra centrosomes or disruption of centromere geometry. Likewise, tumor suppressors combine their known roles in cell cycle progression, growth, and differentiation with the induction of genomic instability is not necessarily new and a substantial body of evidence supports.

Most importantly, it is essential to predict cancer driver genes based on patient genomics data, such as The Cancer Genome Atlas (TCGA). Recently, many studies have taken place to look for cancer driver genes with greater than expected background mutational rates, thus identifying a ranked list of candidate genes based on a small collection of genetic features such as somatic mutations and copy number alterations. But, in terms of distinguishing between tumor suppressor and oncogenes, only handful of approaches exist, such as TUSON and the 20/20+ machine-learning method.

In this work, we use a naïve Bayes computation approach to classify cancer genes as OGs, TSs or drivers/passengers for 21 cancer types with 3,091 fusions. Based on the mutation data from TCGA, we found that TSs had the highest mutation frequency. Based on the human PPI network, we found that TSs and OGs had similar global network topological characteristics. Likewise, the TSs and OGs tended to interact with each other. Integration of mutation frequency with TS and OG networks provided insight that TSs and OGs might jointly contribute to cancer development. We also found that from the initial 6,087 genes, the highest number of OGs and TSs classified belongs to CA, followed by ST and LK, LY, ME, and GL. Likewise, in CA, genes for the cell cycle, DNA replication, spliceosome, proteasomes, mismatch repair, p53 signaling pathway, nucleotide excision repair and other ten pathways were up-regulated Additionally, some down-regulated pathways in LK and LY were all enriched in CA. However, three pathways were down-regulated in CA but up-regulated in the LK and LY. TS and OG proteins had the highest degree and betweenness in the human PPI network, as compared to other proteins. Also, for all 21 cancer types, we identified *ARGFXP2*, *EHMT1*, *BCLAF1* and *DPF2* as being down-regulated. Similarly, we found only *AMBRA1* to be up-regulated in all cancer, as well as 188 common marker genes and 100 significant genes. These results indicate that most mutated tumor suppressors that integrate hyper-methylated partners in protein interaction networks follow the patterns of oncogenes. In summary, this study comprehensively investigated TSs and OGs from the perspective of networks, providing novel insight into their roles in cancer development and treatment.

## Acknowledgements

We acknowledge Dr. Dorith Raviv-Shay for proofreading the manuscript. We thank Dr. Michael Blank for viable discussions on TSs and OGs.

## Author Contributions

S.T. conducted the experiments; S.T. and M.F.-M. performed the analysis. S.T. and M.F.-M. designed the experiments and wrote the paper.

## References

Agajanian, S. et al. (2018) Machine Learning Classification and Structure-Functional Analysis of Cancer Mutations Reveal Unique Dynamic and Network Signatures of Driver Sites in Oncogenes and Tumor Suppressor Genes. J Chem Inf Model. 2018 Sep 25. doi: 10.1021/acs.jcim.8b00414.

Akula, N. et al. (2011) A network-based approach to prioritize results from genome-wide association studies. PLoS One 6(9): e24220.

Anderson, K. et al. (2010) Misregulated E-Cadherin Expression Associated with an Aggressive Brain Tumor Phenotype. PLoS ONE 5(10): e13665.

Barretina, J. et al. (2012) The Cancer Cell Line Encyclopedia enables predictive modelling of anticancer drug sensitivity. Nature 483: 603–607.

Berger, A.H. et al. (2011) A continuum model for tumour suppression. Nature 476(7359): 163–169.

Blank, M. et al. (2012) A tumor suppressor function of Smurf2 associated with controlling chromatin landscape and genome stability through RNF20. Nat Med. 18: 227–234.

Ciriello, G. et al. (2012) Mutual exclusivity analysis identifies oncogenic network modules. Genome Res. 22: 398–406.

Croce, C.M. (2009) Causes and consequences of microRNA dysregulation in cancer. Nat Rev Genet 10: 704–714.

Csilléry, K. et al. (2010) Approximate Bayesian Computation (ABC) in practice. Trends Ecol Evol 25, 410–418.

Davoli, T., et al. (2013) Cumulative haploinsufficiency and triplosensitivity drive aneuploidy patterns and shape the cancer genome. Cell 155(4): 948–962.

Dees, N.D. et al. (2012) MuSiC: Identifying mutational significance in cancer genomes. Genome Res 22(8): 1589–1598.

Deng, Y. et al. (2018) A pan-cancer atlas of cancer hallmark-associated candidate driver lncRNAs. Mol Oncol. 2018 Sep 14. doi: 10.1002/1878-0261.12381.

Ding, L. et al. (2014) Expanding the computational toolbox for mining cancer genomes. Nat. Rev. Genet. 15: 556–570.

Esteller, M. (2002) CpG island hypermethylation and tumor suppressor genes: a booming present, a brighter future. Oncogene 21: 5427–5440.

Frenkel-Morgenstern, M. et al. (2012) Chimeras taking shape: potential functions of proteins encoded by chimeric RNA transcripts. Genome Res. 22(7): 1231–42.

Frenkel-Morgenstern, M. et al. (2017) ChiPPI: A Novel Method for Mapping Chimeric Protein-Protein Interactions Uncovers Selection Principles of Protein Fusion Events in Cancer. Nucleic Acids Res. 45(12): 7094–7105.

Friedman, N. et al. (1997) Bayesian Network Classifiers. Machine Learning 29(2-3): 131–163.

Futreal, P. A. et al. (2004) A census of human cancer genes. Nat. Rev. Cancer 4: 177–183.

Garraway, L.A. et al. (2013) Lessons from the cancer genome. Cell 153(1): 17–37.

Gerlo, S. et al. (2005) Multiple, PKA-dependent and PKA-independent signals are involved in cAMP-induced PRL expression in the eosinophilic cell line Eol-1. Cell Signal 17: 901–9.

Gonzalez-Perez, A. et al. (2013) Computational approaches to identify functional genetic variants in cancer genomes. Nat. Methods 10: 723–729.

Gorohovski, A. et al. (2017) ChiTaRS-3.1-the enhanced chimeric transcripts and RNA-seq database matched with protein-protein interactions. Nucleic Acids Res. 45(D1): D790–D795.

Halliday, B.J. et al. (2018) Germline mutations and somatic inactivation of TRIM28 in Wilms tumour. PLoS Genet. 14(6): e1007399.

Hattori et al. (2018) Analysis of DNA Methylation in Tissues Exposed to Inflammation. Methods Mol Biol. 1725: 185–199.

Hofree, M. et al. (2013) Network-based stratification of tumor mutations. Nat Methods 10(11): 1108–1115.

Ideker, T. et al. (2002) Discovering regulatory and signalling circuits in molecular interaction networks. Bioinformatics 18(Suppl 1): S233–S240.

Imielinski, M. et al. (2012) Mapping the hallmarks of lung adenocarcinoma with massively parallel sequencing. Cell 150: 1107–1120.

Jaratlerdsiri, W. et al. (2018) Whole Genome Sequencing Reveals Elevated Tumor Mutational Burden and Initiating Driver Mutations in African Men with Treatment-Naive, High-Risk Prostate Cancer. Cancer Res. 2018 Sep 14. pii: canres.0254.2018.

Jonsson, P.F. et al. (2006) Global topological features of cancer proteins in the human interactome. Bioinformatics 22(18): 2291–7.

Khetchoumian K. et al. (2008) Trim24 (Tif1 alpha): an essential ‘brake’ for retinoic acid-induced transcription to prevent liver cancer. Cell Cycle 7, 3647–3652.

Kim, P. et al. (2018) TissGDB: tissue-specific gene database in cancer. Nucleic Acids Res. 46(D1): D1031–D1038.

Kumar, R.D. et al. (2015) Statistically identifying tumor suppressors and oncogenes from pan-cancer genome-sequencing data. Bioinformatics 31(22): 3561–8.

Lawrence, M. S. et al. (2014) Discovery and saturation analysis of cancer genes across 21 tumour types. Nature 505: 495–501.

Lawrence, B. et al. (2018) Recurrent loss of heterozygosity correlates with clinical outcome in pancreatic neuroendocrine cancer. NPJ Genom Med. 20; 3:18.

D. M. et al. (2015) Pan-cancer network analysis identifies combinations of rare somatic mutations across pathways and protein complexes. Nat. Genet. 47: 106–114.

Ley, T.J. et al. (2008) DNA sequencing of a cytogenetically normal acute myeloid leukaemia genome. Nature 456(7218): 66–72.

Liu, Y. et al. (2017) ONGene: A literature-based database for human oncogenes. Journal of Genetics and Genomics. 44(2): 119–121.

Mi, H. et al. (2016) PANTHER version 11: expanded annotation data from Gene Ontology and Reactome pathways, and data analysis tool enhancements. Nucleic Acids Res. 45(D1): D183–D189.

Mitelman, F. et al. (2007) The impact of translocations and gene fusions on cancer causation. Nat Rev Cancer. 7(4): 233–45.

Mularoni, L. et al. (2016) OncodriveFML: A general framework to identify coding and non-coding regions with cancer driver mutations. Genome Biol 17(1): 128.

Osborne, C. et al. (2004) Oncogenes and tumor suppressor genes in breast cancer: potential diagnostic and therapeutic applications. Oncologist 16(4): 361–377.

Pavel, A.B. et. al. (2016) Identifying cancer type specific oncogenes and tumor suppressors using limited size data. J Bioinform Comput Biol. 14(6): 1650031.

Pundir et al. (2017) UniProt Protein Knowledgebase. Methods Mol Biol. 1558: 41–55.

Reimand, J. et al. (2013) Systematic analysis of somatic mutations in phosphorylation signaling predicts novel cancer drivers. Mol. Syst. Biol. 9: 637.

Schroeder, M.P. et al. (2014) OncodriveROLE classifies cancer driver genes in loss of function and activating mode of action. Bioinformatics 30(17): i549–55.

Shimono, Y. et al. (2009) Down-regulation of miRNA-200c links breast cancer stem cells with normal stem cells. Cell 138: 592–603.

Subramanian, A. et al. (2005) Gene set enrichment analysis: a knowledge-based approach for interpreting genome-wide expression profiles. Proc Natl Acad Sci U S A. 102(43): 15545–50.

Sun, W. et al. (2011) Anticancer activity of the PR domain of tumor suppressor RIZ1. Int J Med Sci 8(2): 161–167.

Tagore, S. et al. (2018) A Comprehensive Approach Characterizing Fusion Proteins and Their Interactions Using Biomedical Literature. bioRxiv 371088; doi: https://doi.org/10.1101/371088.

Tamborero, D. et al. (2013) OncodriveCLUST: exploiting the positional clustering of somatic mutations to identify cancer genes. Bioinformatics 29(18): 2238–44.

Tavanaei, A. et al. (2017) A deep learning model for predicting tumor suppressor genes and oncogenes from PDB structure. 2017 IEEE International Conference on Bioinformatics and Biomedicine (BIBM), Kansas City, MO, 2017, pp. 613–617.

Tokheim, C.J., et al. (2016) Evaluating the evaluation of cancer driver genes. Proc Natl Acad Sci U S A. 113(50): 14330–14335.

Vandin, F. et al. (2012) De novo discovery of mutated driver pathways in cancer. Genome Res. 22(2): 375–85.

van Haaften, G. et al. (2009) Somatic mutations of the histone H3K27 demethylase gene UTX in human cancer. Nat Genet 41: 521–523.

Vergara, D. et al. (2015) Proteomics analysis of E-cadherin knockdown in epithelial breast cancer cells. J Biotechnol. 202: 3–11.

Vogelstein, B. et al. (2013) Cancer genome landscapes. Science 339(6127): 1546–1558.

Wang, K. et al. (2018) Dissecting cancer heterogeneity based on dimension reduction of transcriptomic profiles using extreme learning machines. PLoS One. 13(9): e0203824.

Wrzeszczynski, K.O. et al. (2011) Identification of tumor suppressors and oncogenes from genomic and epigenetic features in ovarian cancer. PLoS One 6(12): e28503.

Zack, T. I. et al. (2013) Pan-cancer patterns of somatic copy number alteration. Nat. Genet. 45: 1134–1140.

Zerbino et al. (2018) Ensembl 2018. Nucleic Acids Res. 46(D1): D754–D761.

Zhao, M. et al. (2016) TSGene 2.0: an updated literature-based knowledgebase for tumor suppressor genes. Nucleic Acids Res. 44(D1): D1023–31.

Zhu, K. et. al. (2015) Oncogenes and tumor suppressor genes: comparative genomics and network perspectives. BMC Genomics. 16(7): S8.

