## Supplementary data for "Mutated Tumor Suppressors Follow Oncogenes Profile by the Gene Hypermethylation of Partners in the Protein Interaction Networks"

1. Figure S1: Cluster for BLCA
2. Figure S2: Cluster for CESC
3. Figure S3: Cluster for COAD
4. Figure S4: Cluster for HNSC
5. Figure S5: Cluster for KICH
6. Figure S6: Cluster for KIRC
7. Figure S7: Cluster for KIRC
8. Figure S8: Cluster for LGG
9. Figure S9: Cluster for LIHC
10. Figure S10: Cluster for LUSC
11. Figure S11: Cluster for OV
12. Figure S12: Cluster for PRAD
13. Figure S13: Cluster for STAD
14. Figure S14: Cluster for THCA
15. Figure S15: Cluster for UCEC
16. Figure S16: Marker genes heatmap for BLCA
17. Figure S17: Marker genes heatmap for BRCA
18. Figure S18: Marker genes heatmap for CESC
19. Figure S19: Marker genes heatmap for COAD
20. Figure S20: Marker genes heatmap for DLBC
21. Figure S21: Marker genes heatmap for GBM
22. Figure S22: Marker genes heatmap for HNSC
23. Figure S23: Marker genes heatmap for KICH
24. Figure S24: Marker genes heatmap for KIRC
25. Figure S25: Marker genes heatmap for KIRP
26. Figure S26: Marker genes heatmap for LGG
27. Figure S27: Marker genes heatmap for LIHC
28. Figure S28: Marker genes heatmap for LUSC
29. Figure S29: Marker genes heatmap for OV
30. Figure S30: Marker genes heatmap for PACA
31. Figure S31: Marker genes heatmap for PRAD
32. Figure S32: Marker genes heatmap for SARC
33. Figure S33: Marker genes heatmap for SKCM
34. Figure S34: Marker genes heatmap for STAD
35. Figure S35: Marker genes heatmap for THCA
36. Figure S36: Marker genes heatmap for UCEC
37. Figure S37: Significant genes heatmap for BLCA
38. Figure S38: Significant genes heatmap for BRCA
39. Figure S39: Significant genes heatmap for CESC

40. Figure S40: Significant genes heatmap for COAD
41. Figure S41: Significant genes heatmap for DLBC
42. Figure S42: Significant genes heatmap for GBM
43. Figure S43: Significant genes heatmap for HNSC
44. Figure S44: Significant genes heatmap for KICH
45. Figure S45: Significant genes heatmap for KIRC
46. Figure S46: Significant genes heatmap for KIRP
47. Figure S47: Significant genes heatmap for LGG
48. Figure S48: Significant genes heatmap for LIHC
49. Figure S49: Significant genes heatmap for LUSC
50. Figure S50: Significant genes heatmap for OV
51. Figure S51: Significant genes heatmap for PACA
52. Figure S52: Significant genes heatmap for PRAD
53. Figure S53: Significant genes heatmap for SARC
54. Figure S54: Significant genes heatmap for SKCM
55. Figure S55: Significant genes heatmap for STAD
56. Figure S56: Significant genes heatmap for THCA
57. Figure S57: Significant genes heatmap for UCEC
58. Figure S58: Common up-down regulated genes in all 21 cancer types
59. Figure S59: Common marker genes in all 21 cancer types
60. Figure S60: Common significant genes in all 21 cancer types
61. Figure S61: Enriched pathways in DLBC
62. Figure S62: Enriched pathways in SKCM
63. Figure S63: Enriched pathways in GBM
64. Figure S64: Enriched pathways in LGG
65. Figure S65: Enriched pathways in SKCM
66. Figure S66: Enriched pathways in BRCA
67. Figure S67: Enriched pathways in OV
68. Figure S68: Enriched pathways in PACA
69. Figure S69: Enriched pathways in PRAD
70. Table S1: OG list (Known) (top 500)
71. Table S2: TS list (Known) (top 500)
72. Table S3: LK,LY,ME,GL - OG, TS (top 500)
73. Table S4: SC - OG, TS (top 500)
74. Table S5: CA - OG, TS (top 500)
75. Table S6: LK,LY,ME,GL -DR/PA (top 500)
76. Table S7: SC -DR/PA (top 500)
77. Table S8: CA –DR/PA (top 500)
78. Table S9: List of fusions and parental proteins (top 500)
79. Table S10: List of community attachment scores (top 500)
80. Table S11: Communities as predicted using scores (top 500)
81. Table S12: OG and TS predicted by NBC, MutsigCV and 20/20+
82. Table S13: Prediction rate of unique OG and TS by NBC, MuSiC, OncodriveCLUST, OncodriveFM, 20/20+
83. Table S14: Overall comparison of the methods

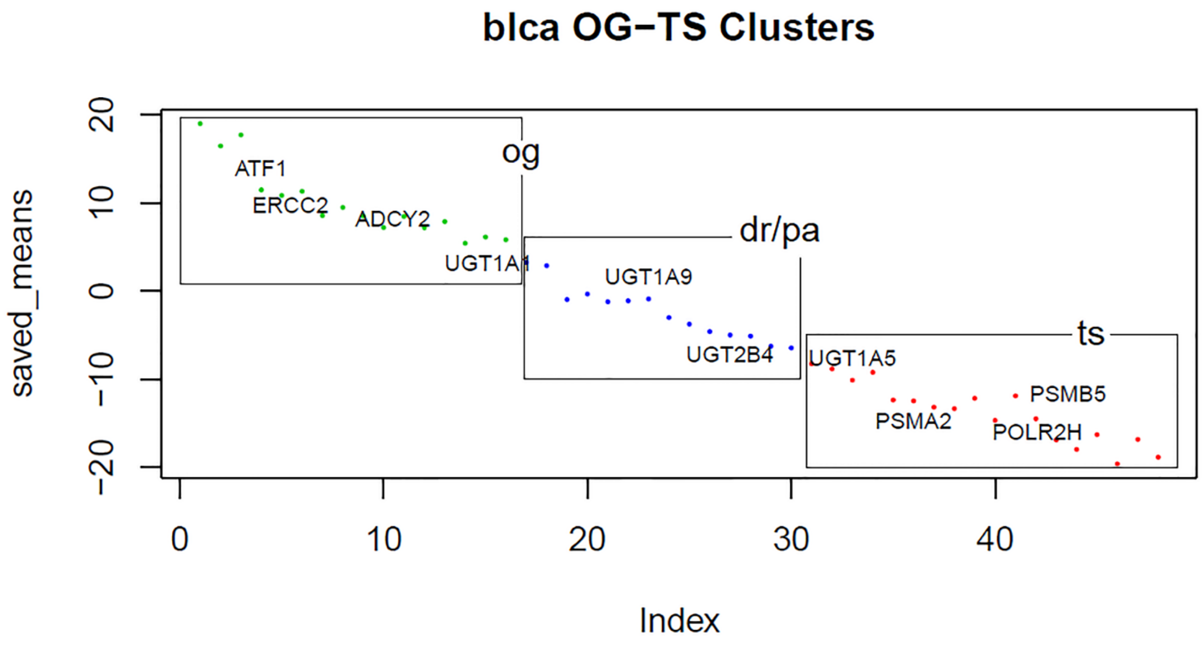

Figure S1: Cluster for BLCA

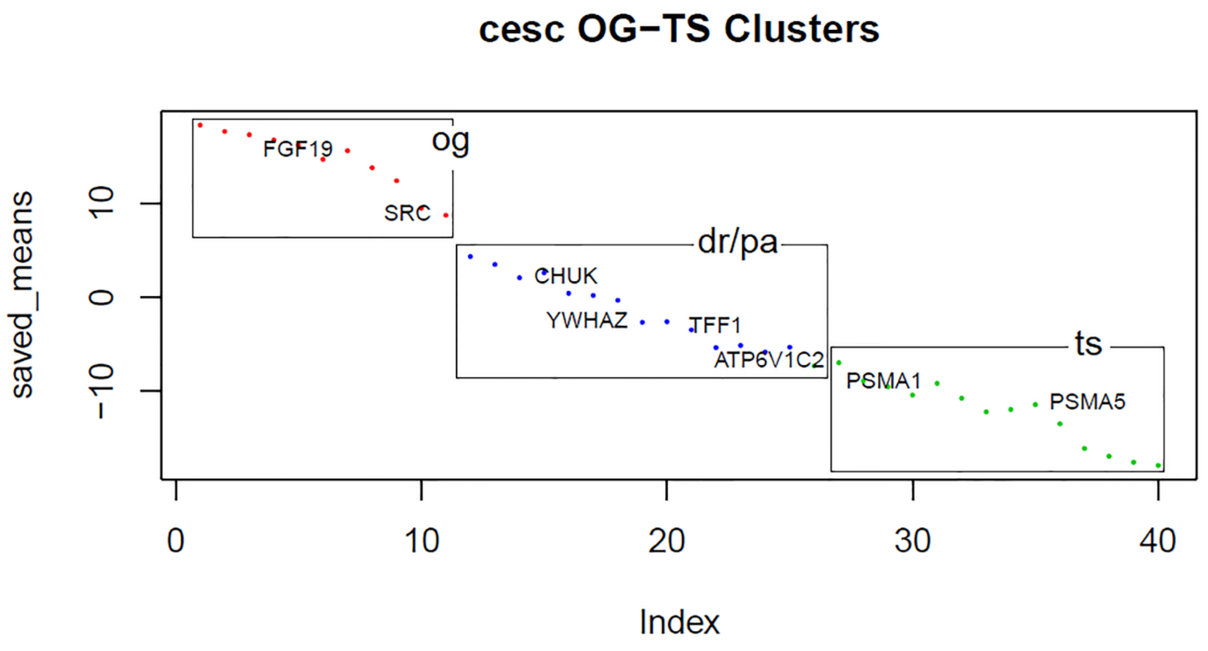

Figure S2: Cluster for CESC

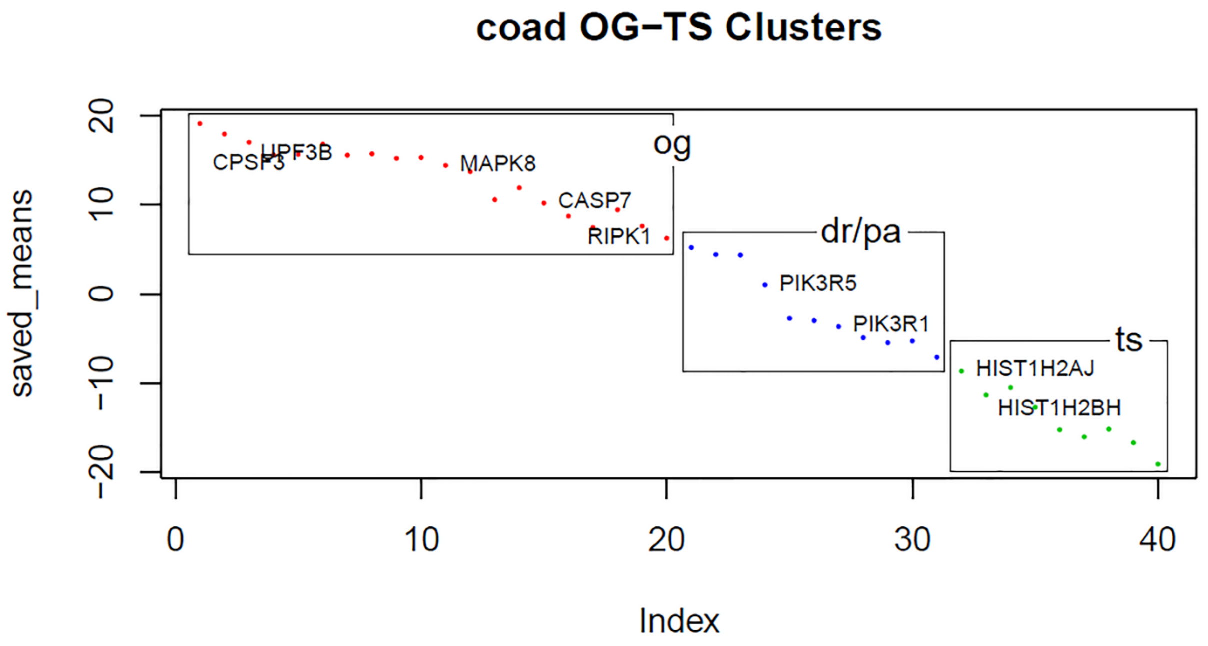

Figure S3: Cluster for COAD

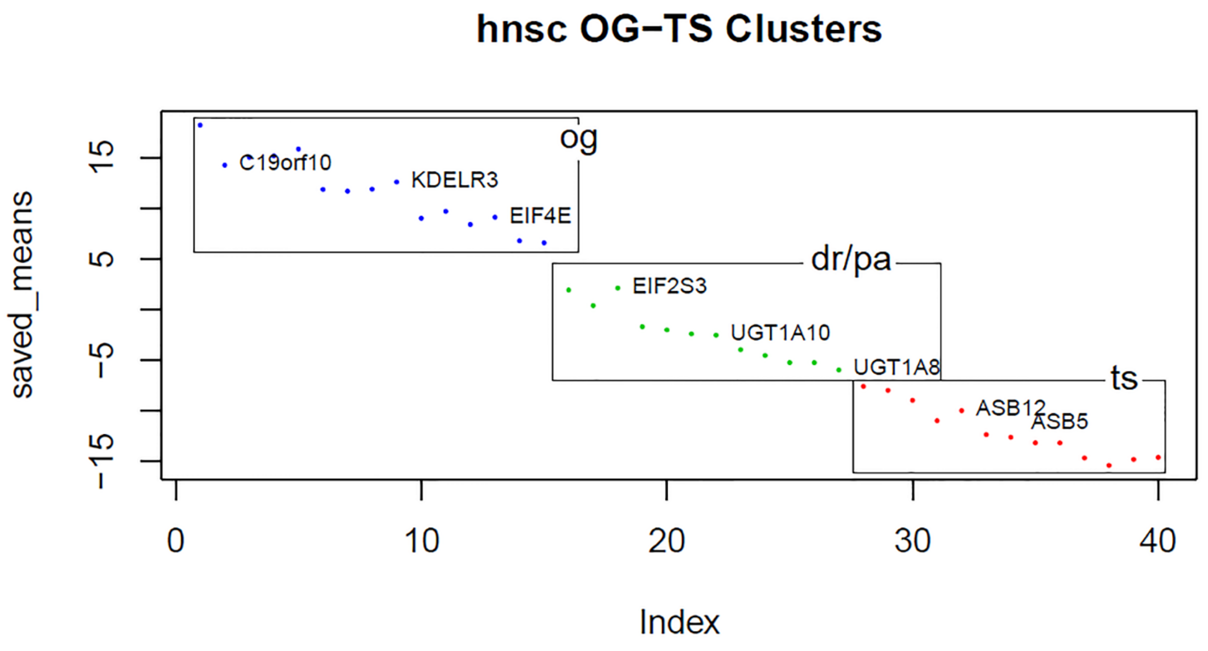

Figure S4: Cluster for HNSC

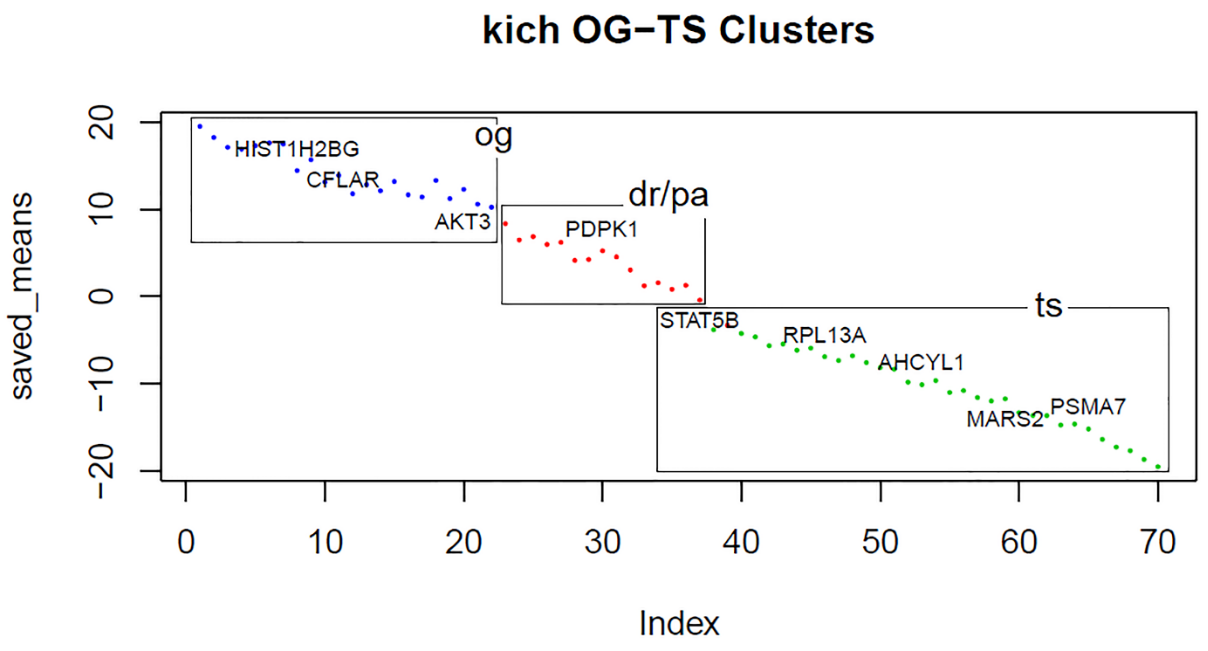

Figure S5: Cluster for KICH

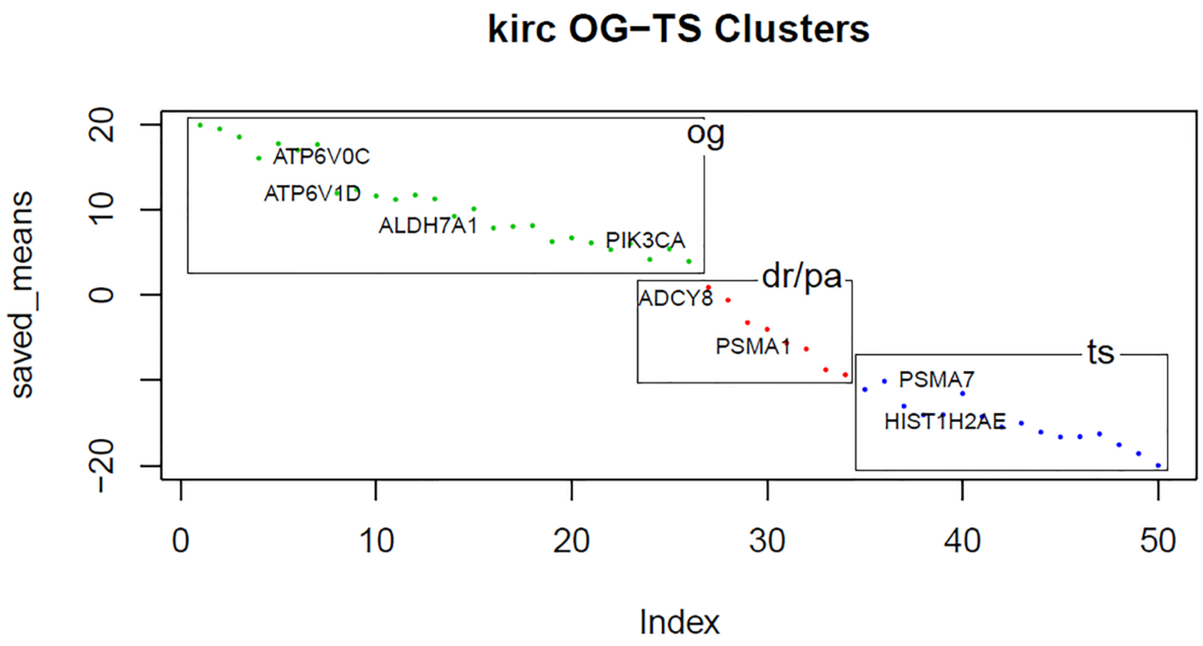

Figure S6: Cluster for KIRC

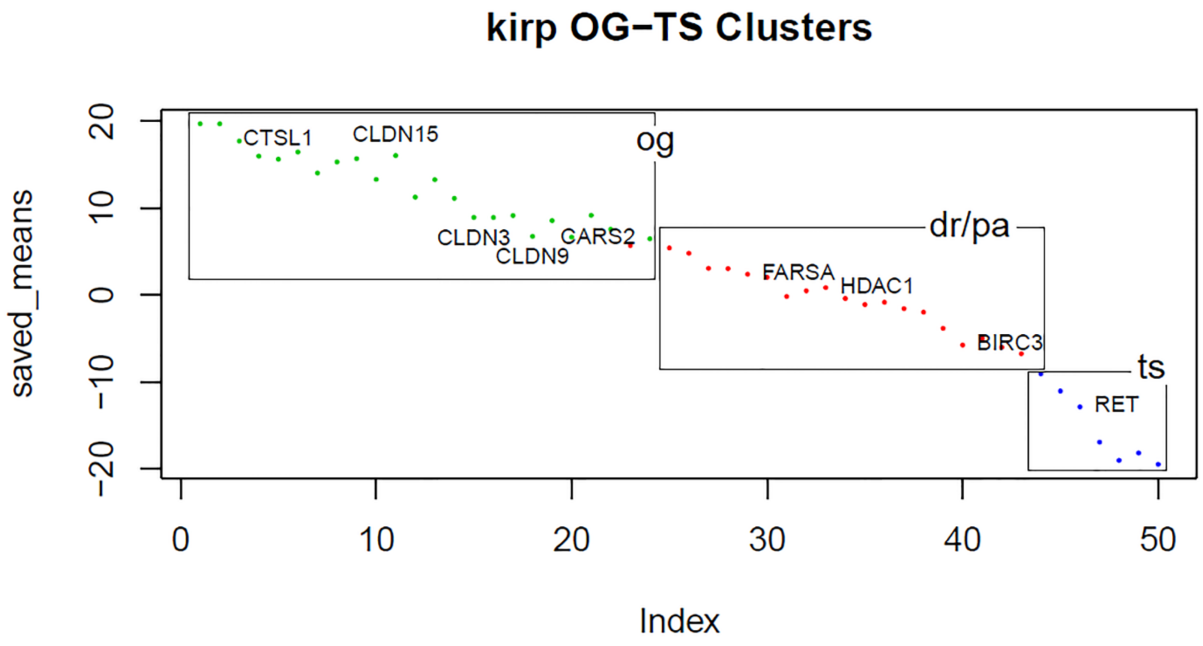

Figure S7: Cluster for KIRC

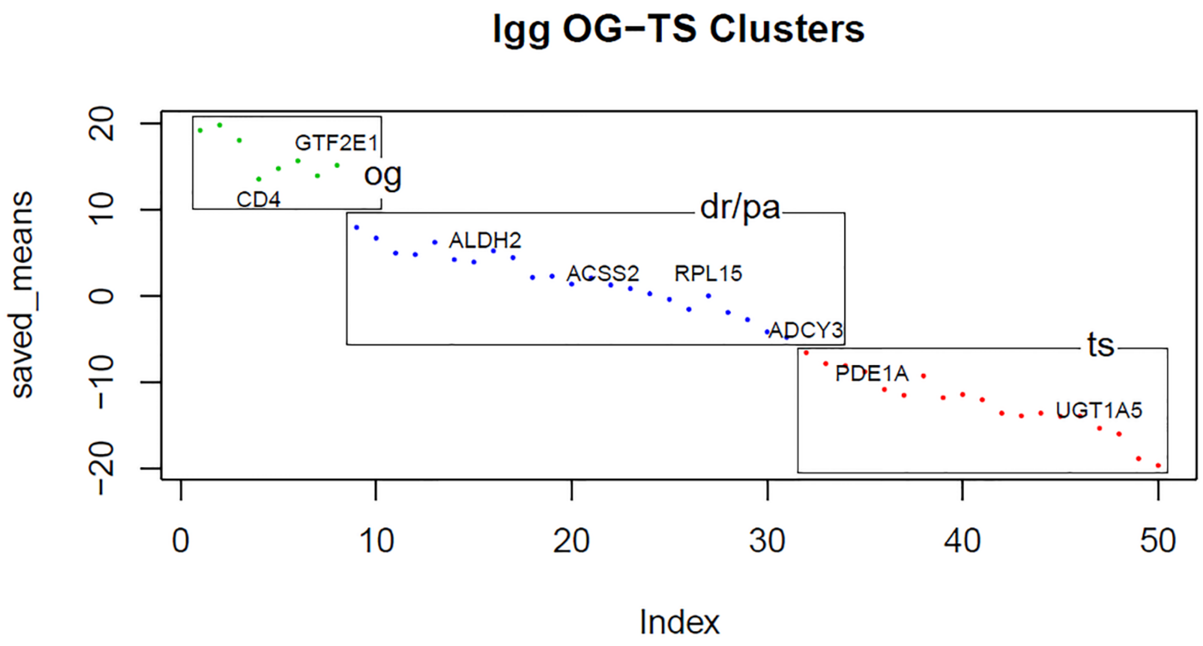

Figure S8: Cluster for LGG

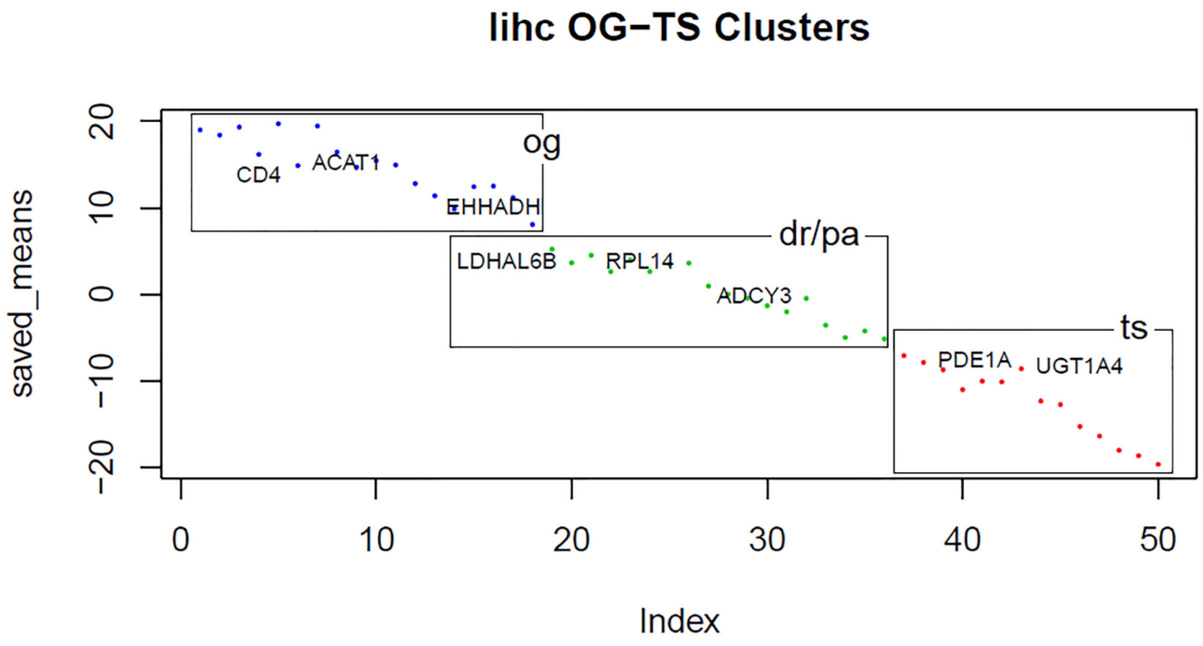

Figure S9: Cluster for LIHC

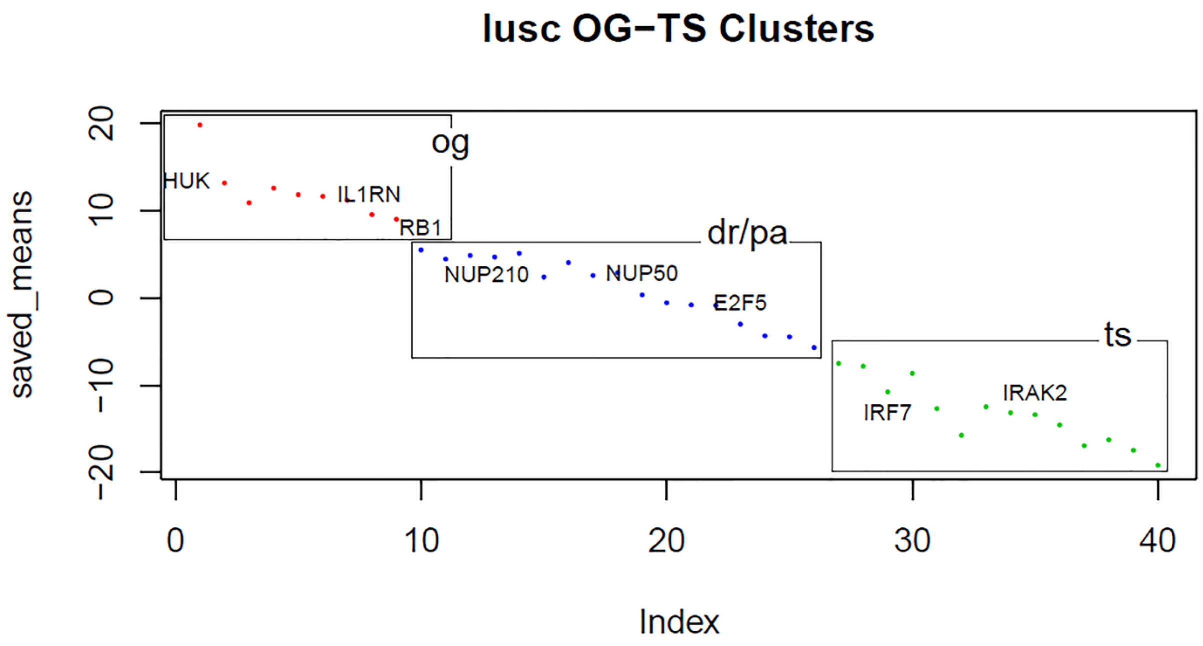

Figure S10: Cluster for LUSC

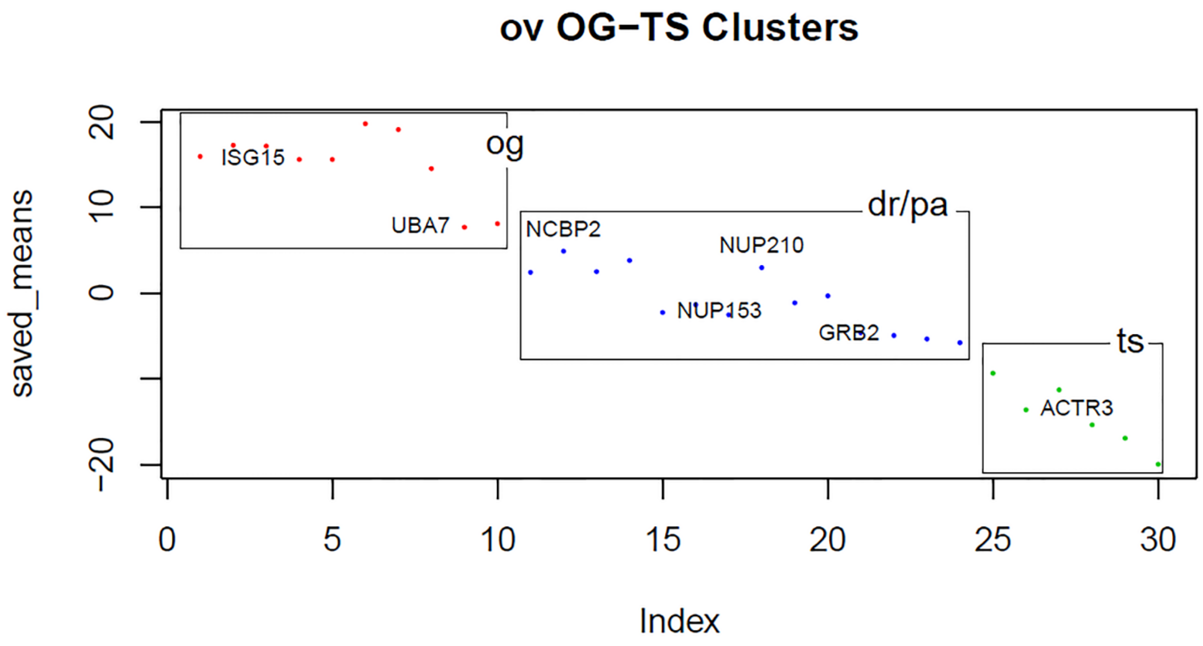

Figure S11: Cluster for OV

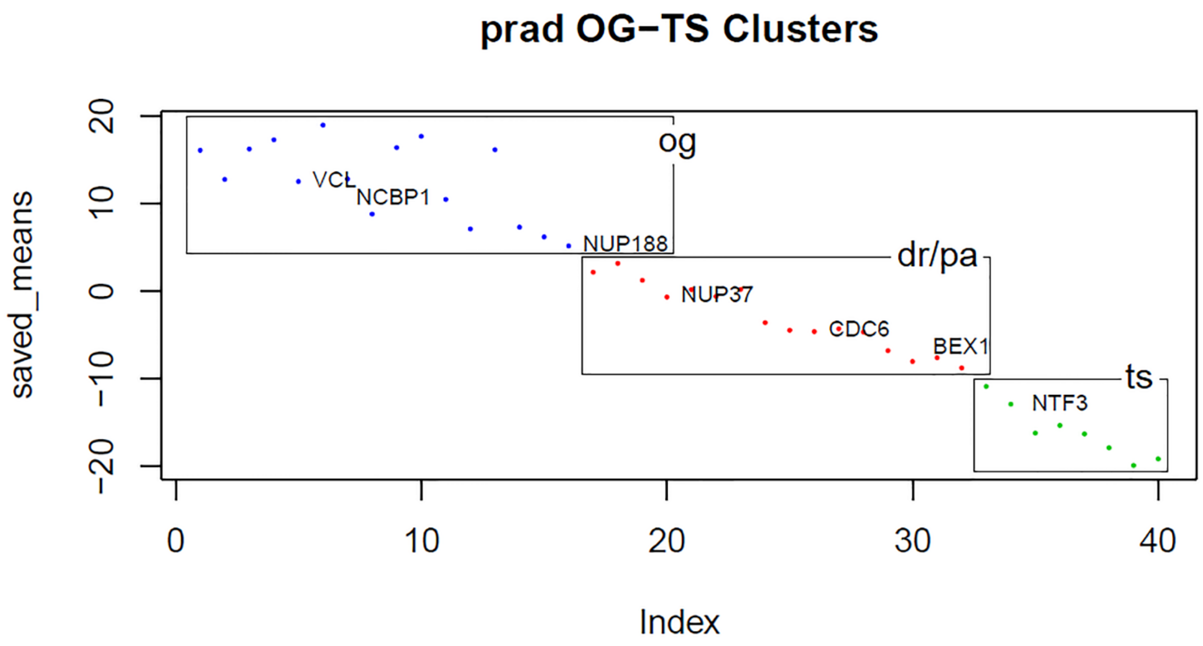

Figure S12: Cluster for PRAD

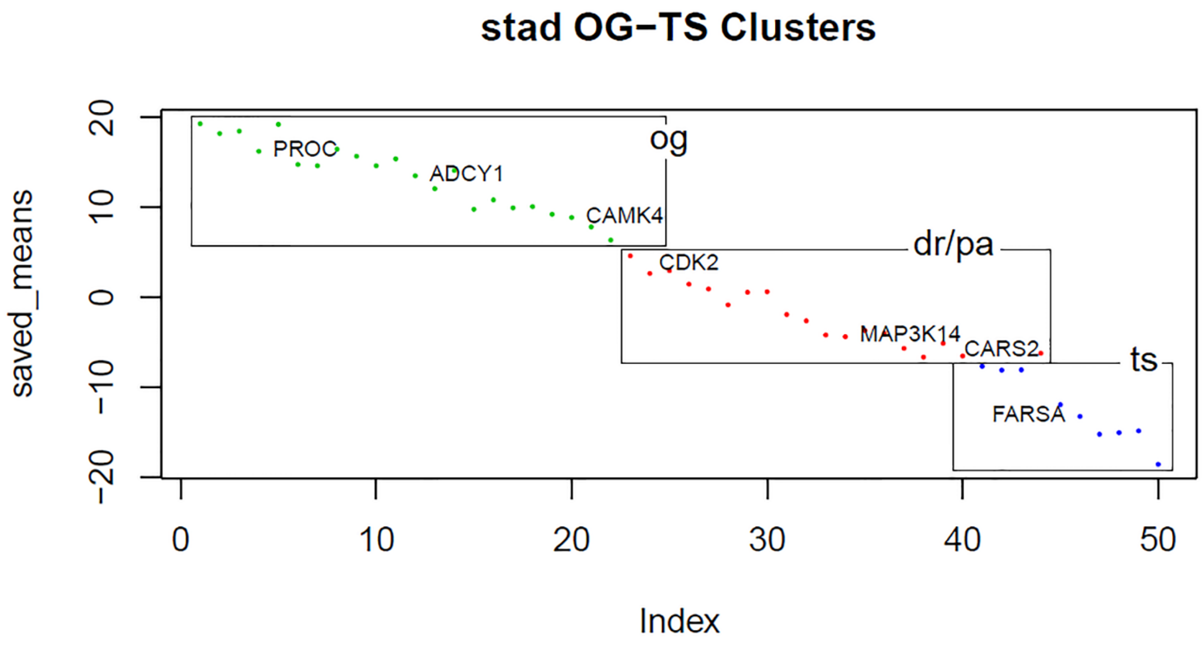

Figure S13: Cluster for STAD

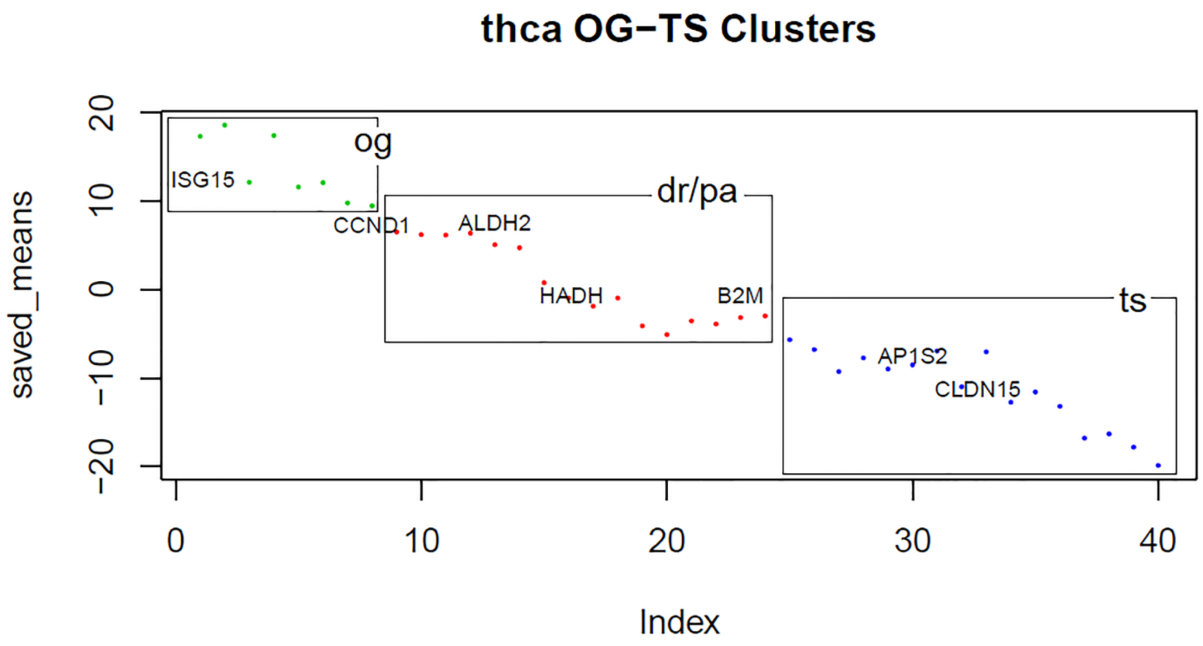

Figure S14: Cluster for THCA

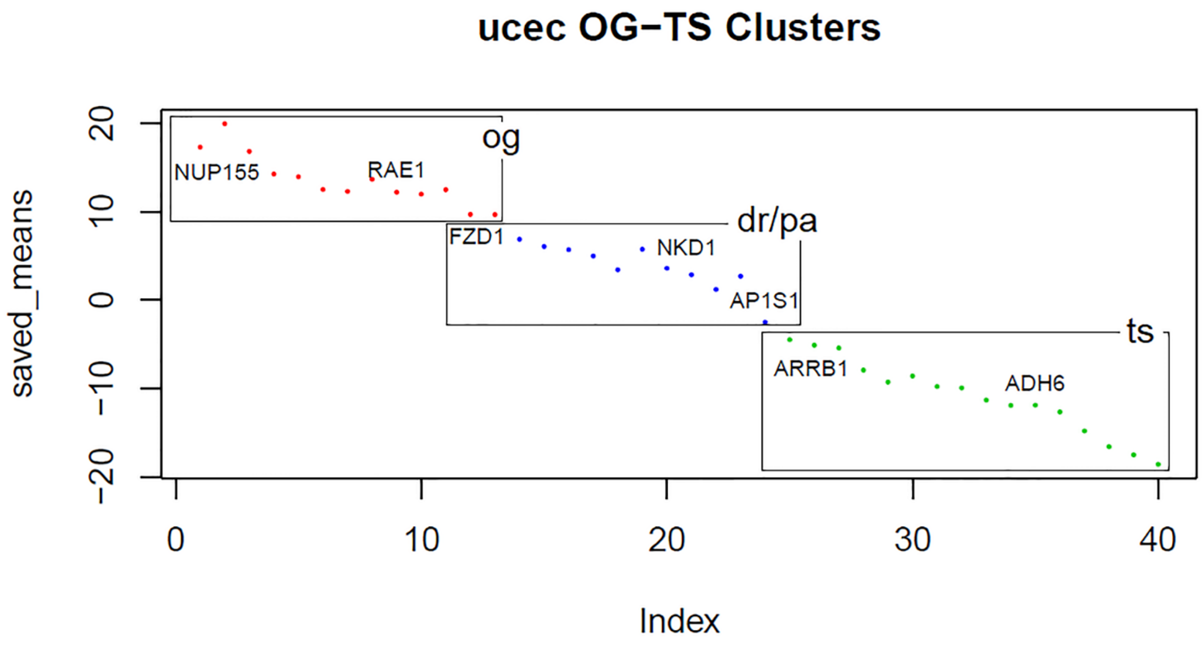

Figure S15: Cluster for UCEC

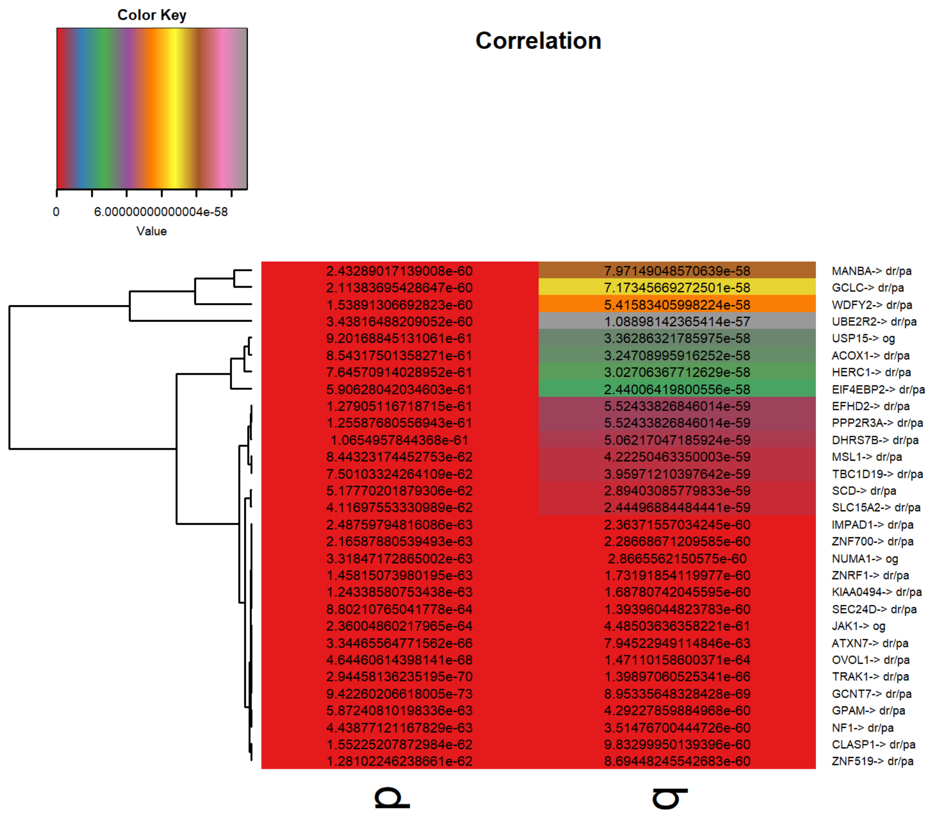

Figure S16: Marker genes heatmap for BLCA

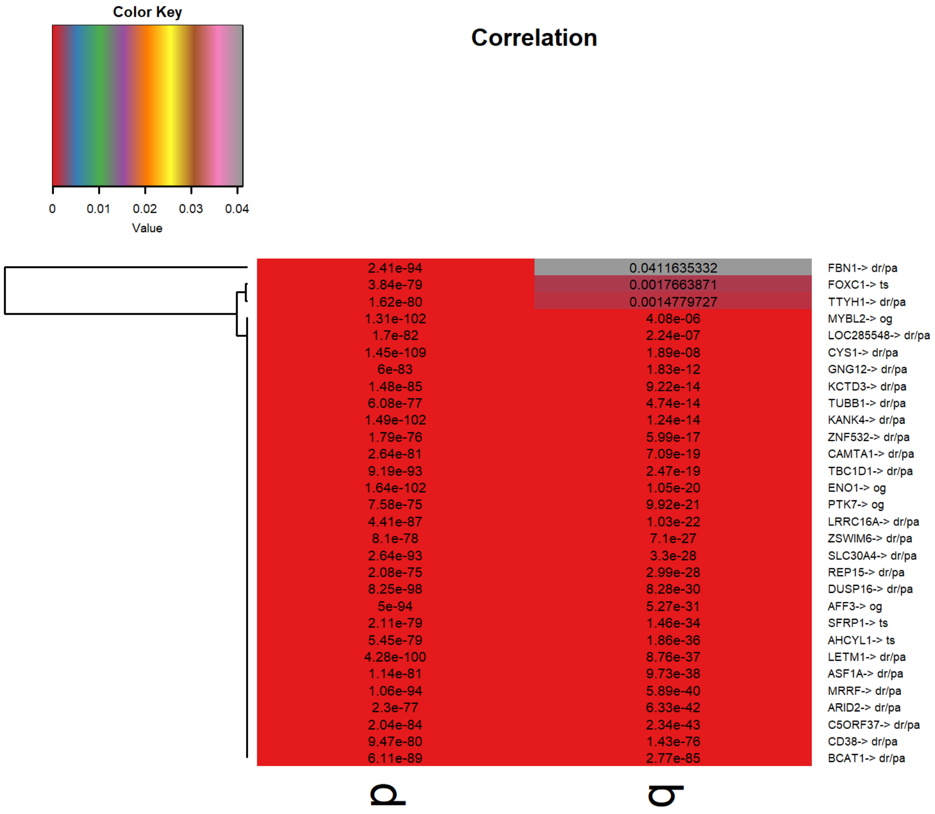

Figure S17: Marker genes heatmap for BRCA

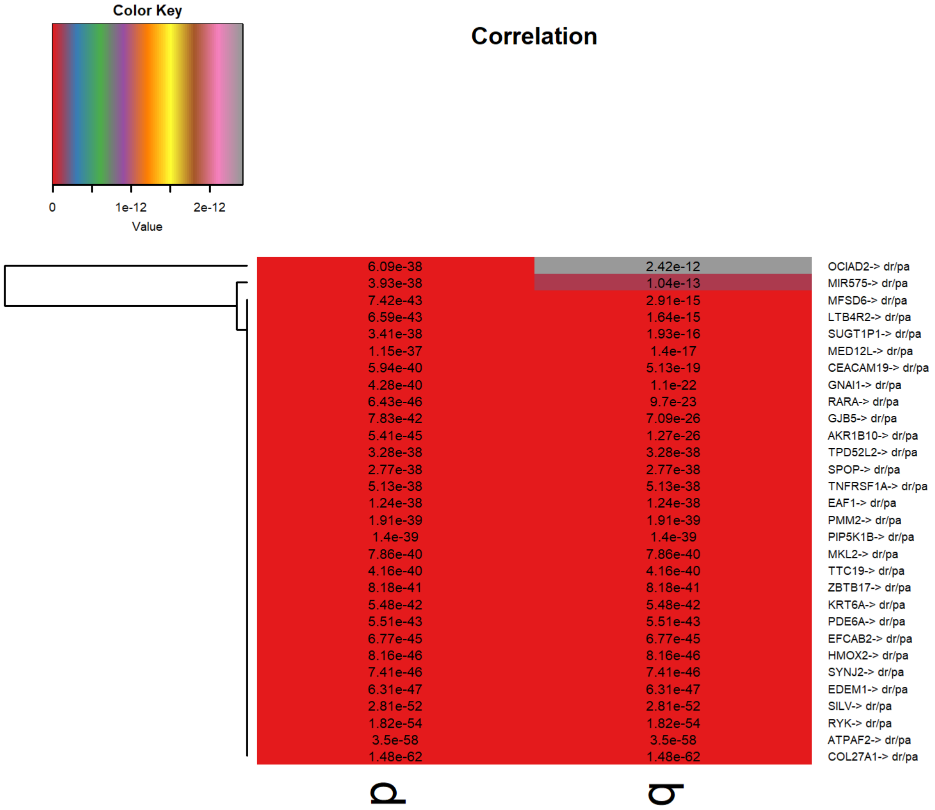

Figure S18: Marker genes heatmap for CESC

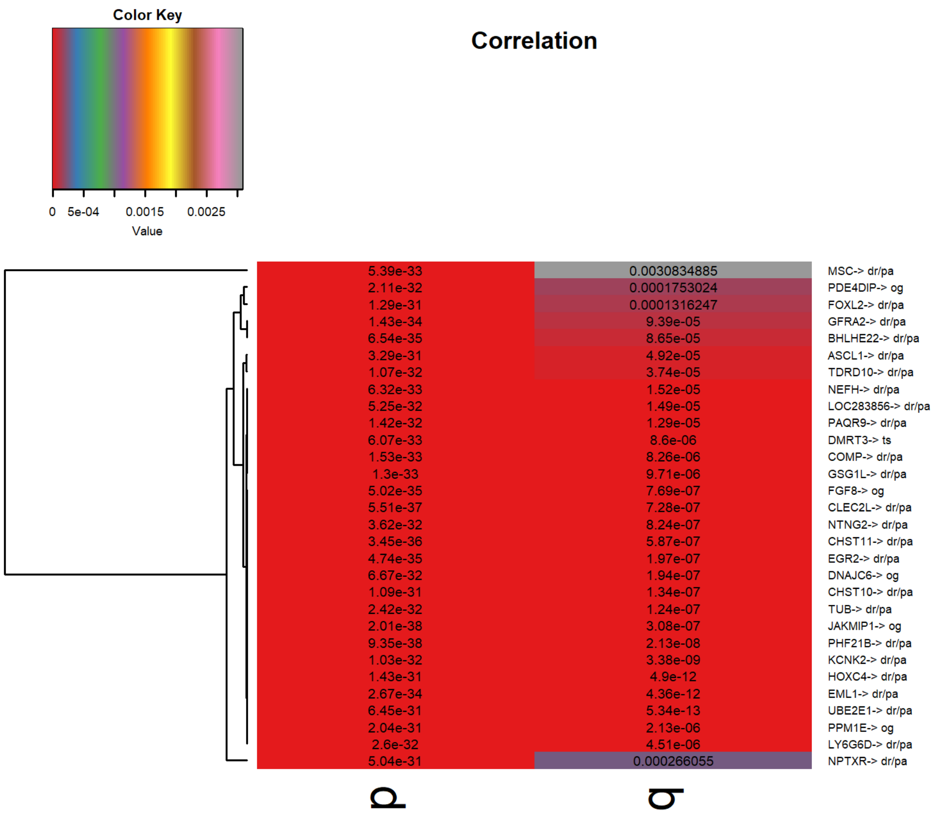

Figure S19: Marker genes heatmap for COAD

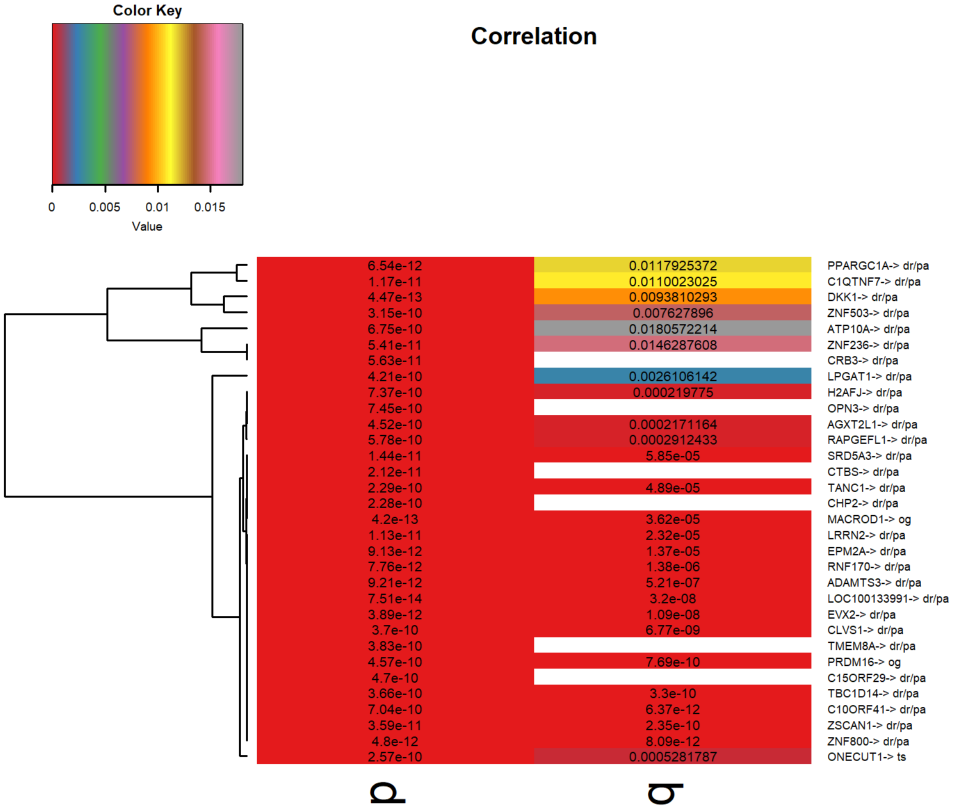

Figure S20: Marker genes heatmap for DLBC

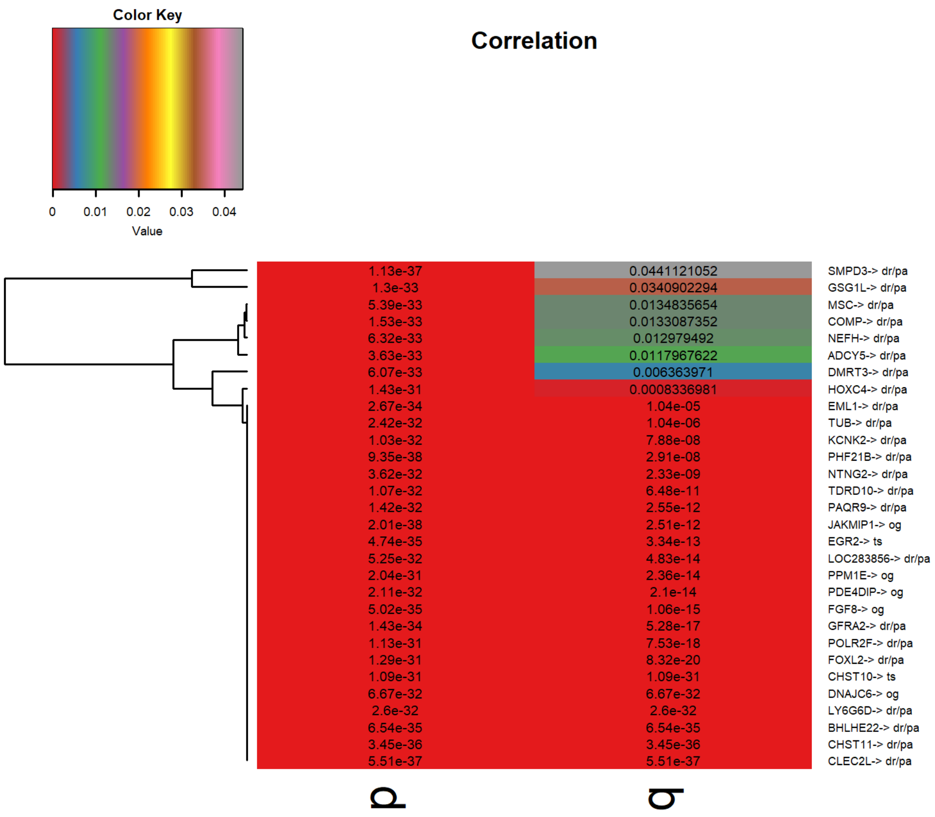

Figure S21: Marker genes heatmap for GBM

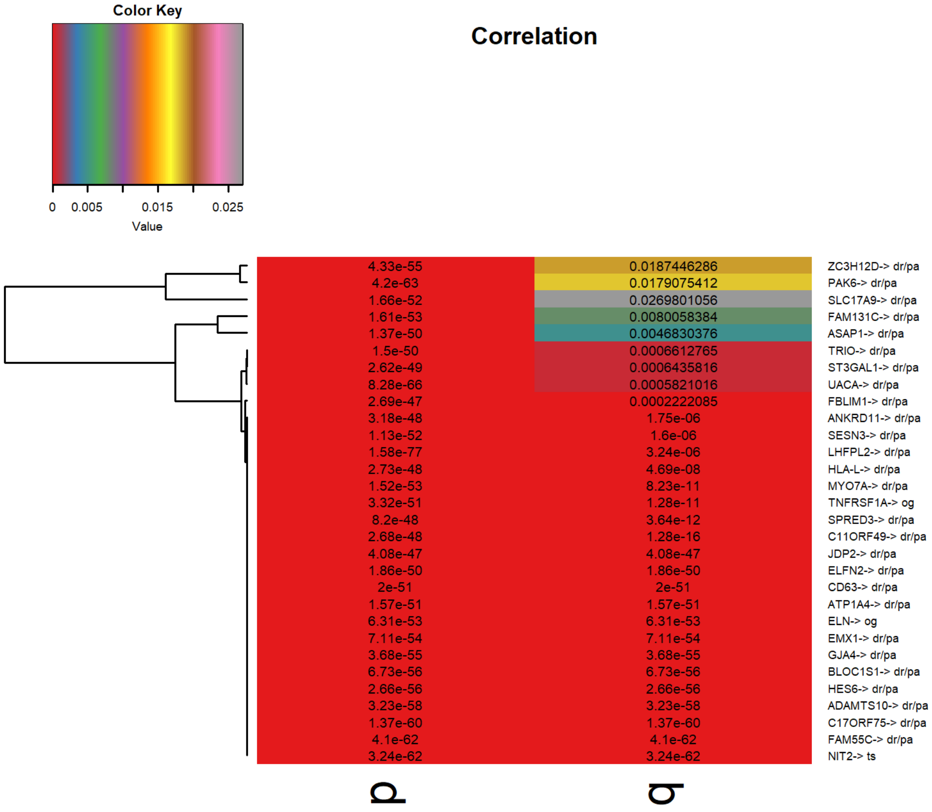

Figure S22: Marker genes heatmap for HNSC

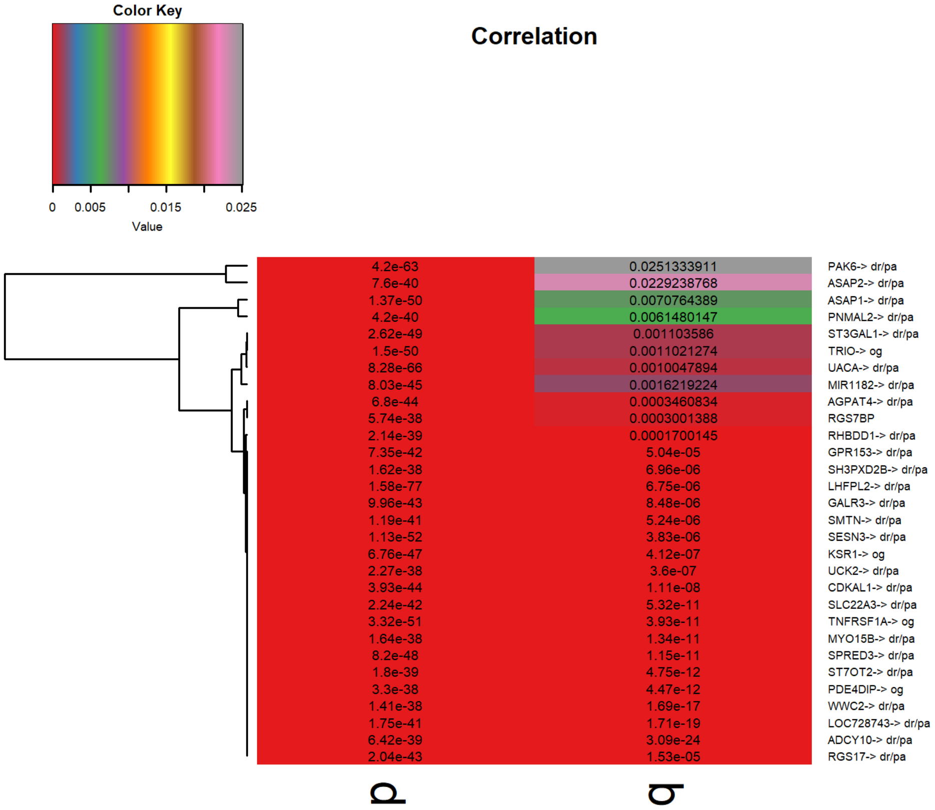

Figure S23: Marker genes heatmap for KICH

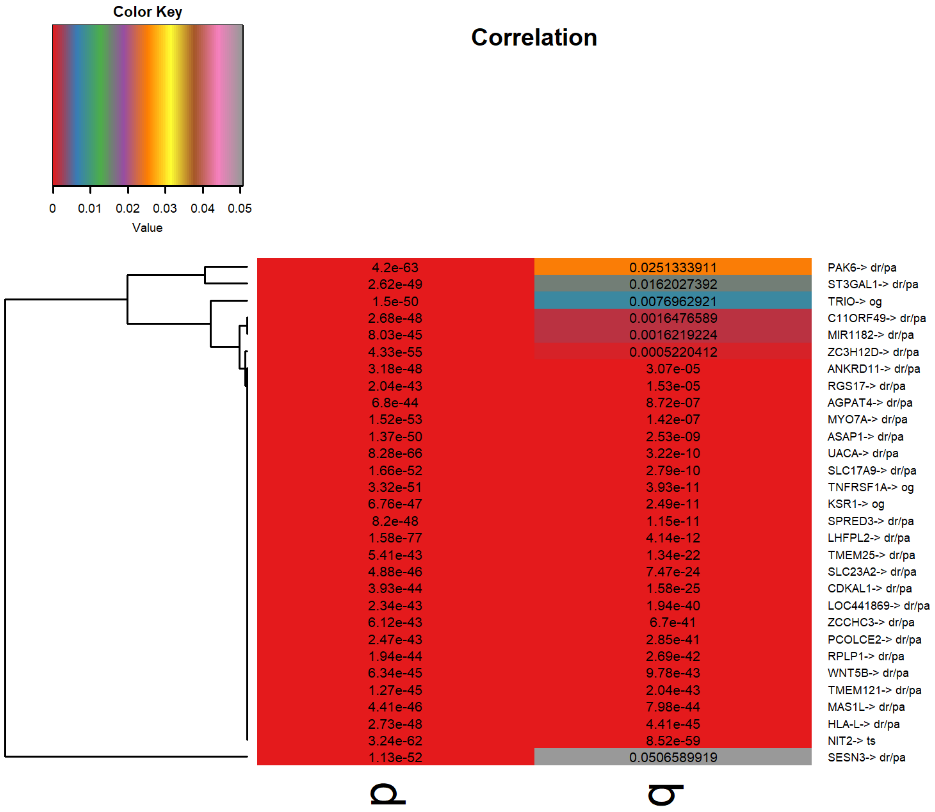

Figure S24: Marker genes heatmap for KIRC

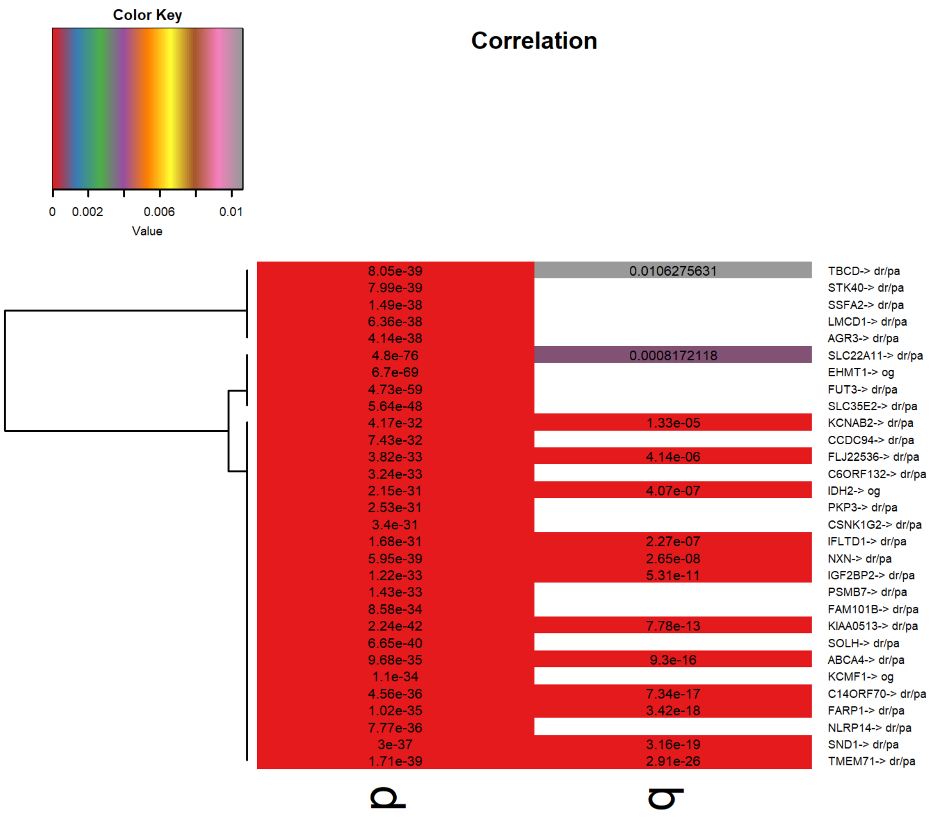

Figure S25: Marker genes heatmap for KIRP

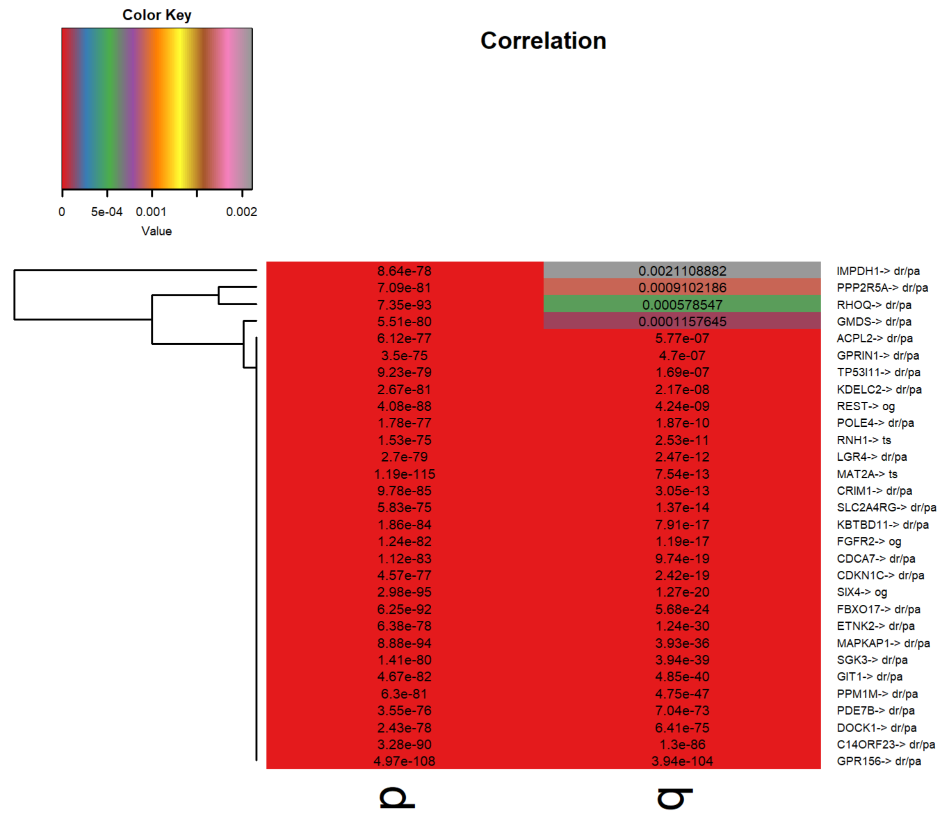

Figure S26: Marker genes heatmap for LGG

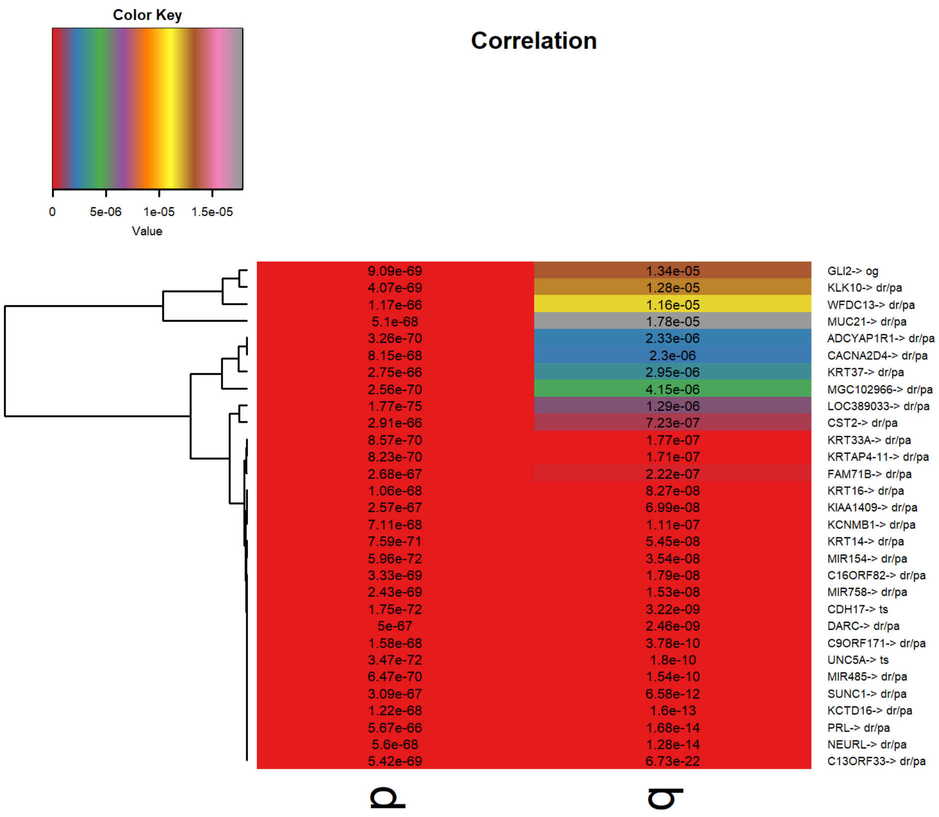

Figure S27: Marker genes heatmap for LIHC

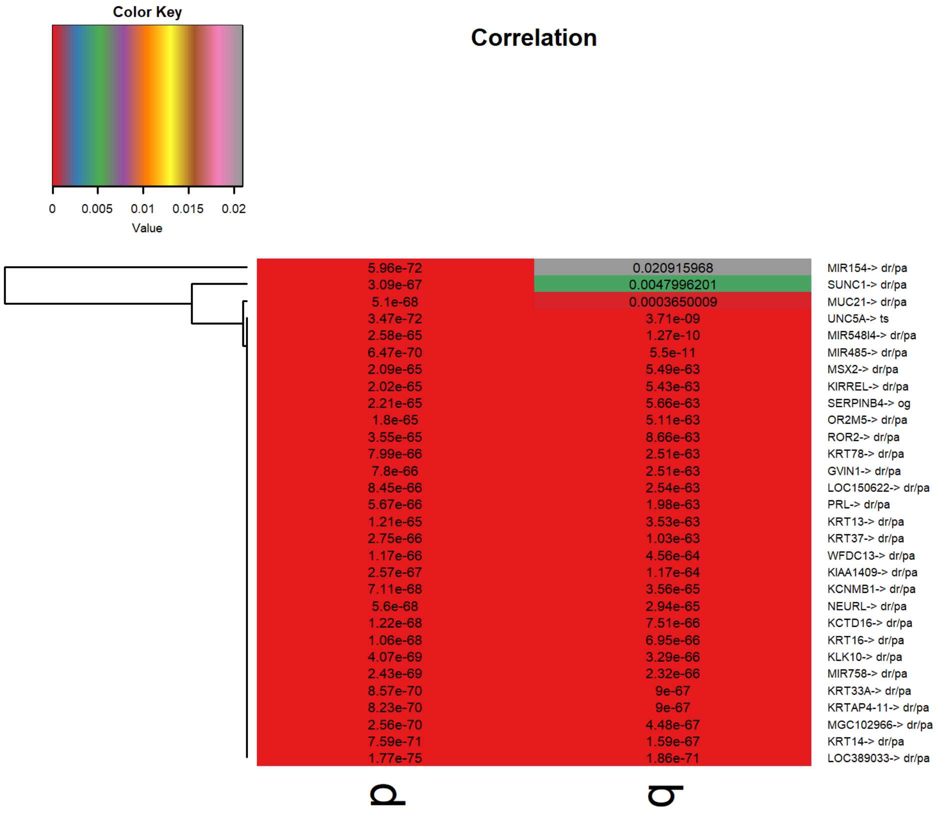

Figure S28: Marker genes heatmap for LUSC

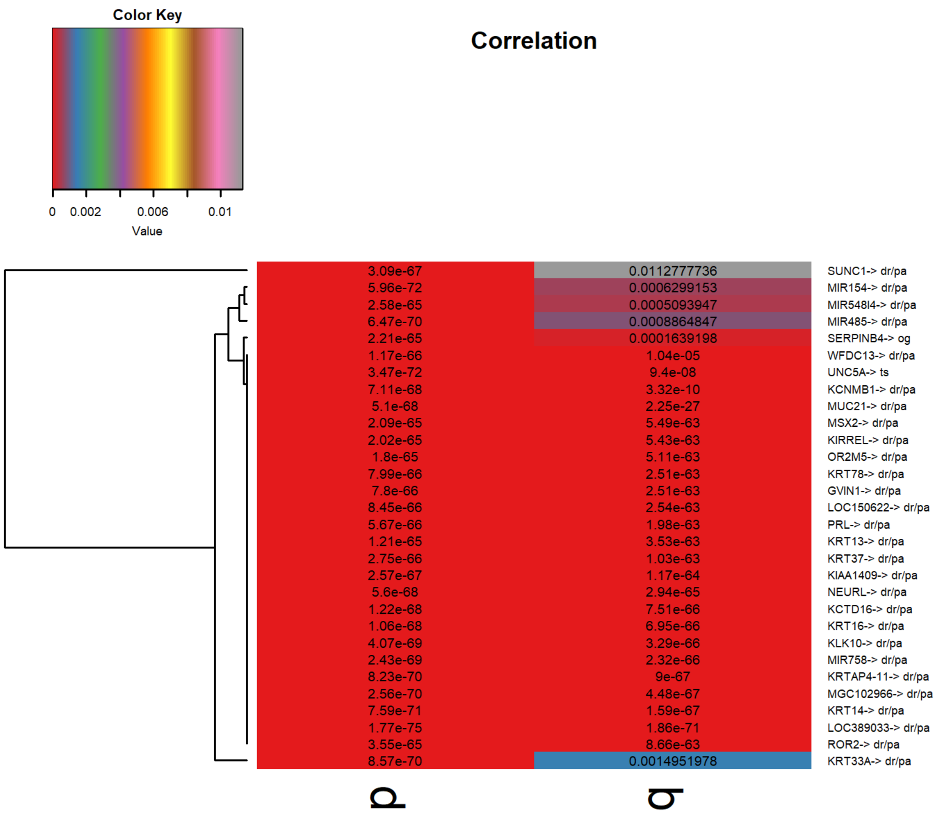

Figure S29: Marker genes heatmap for OV

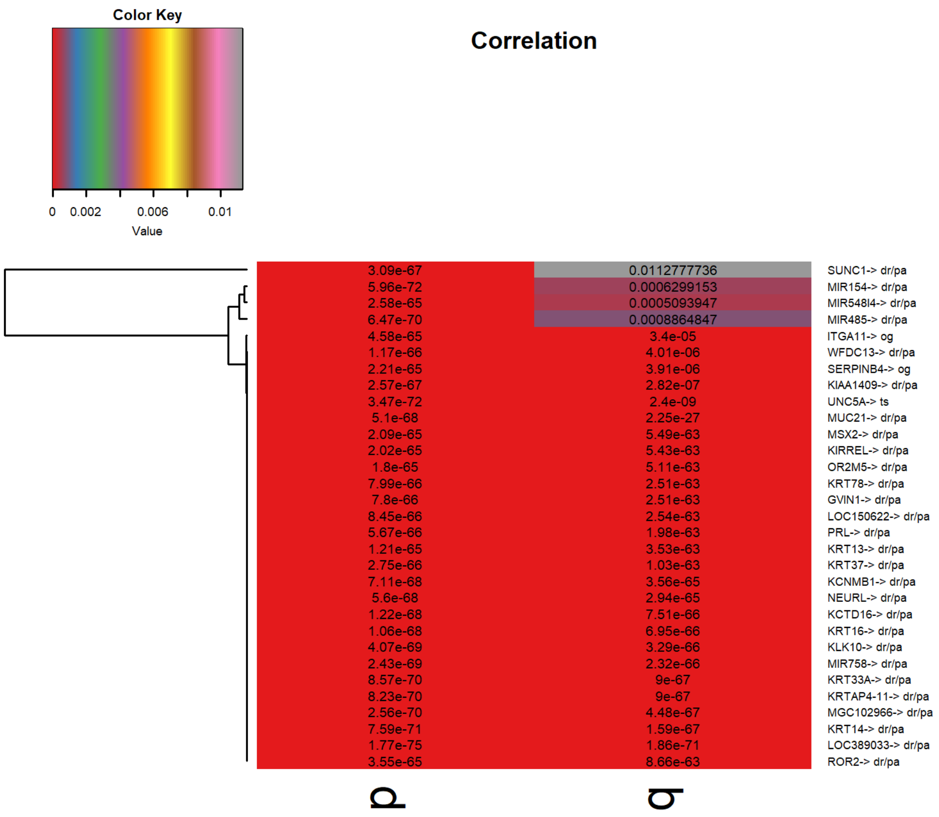

Figure S30: Marker genes heatmap for PACA

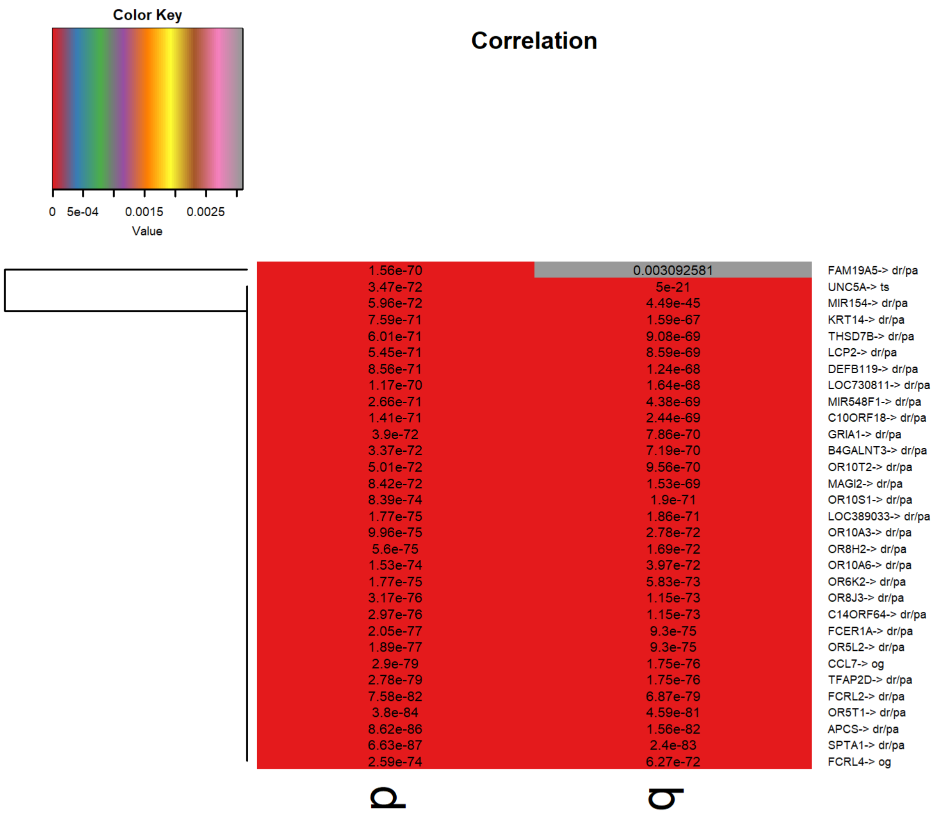

Figure S31: Marker genes heatmap for PRAD

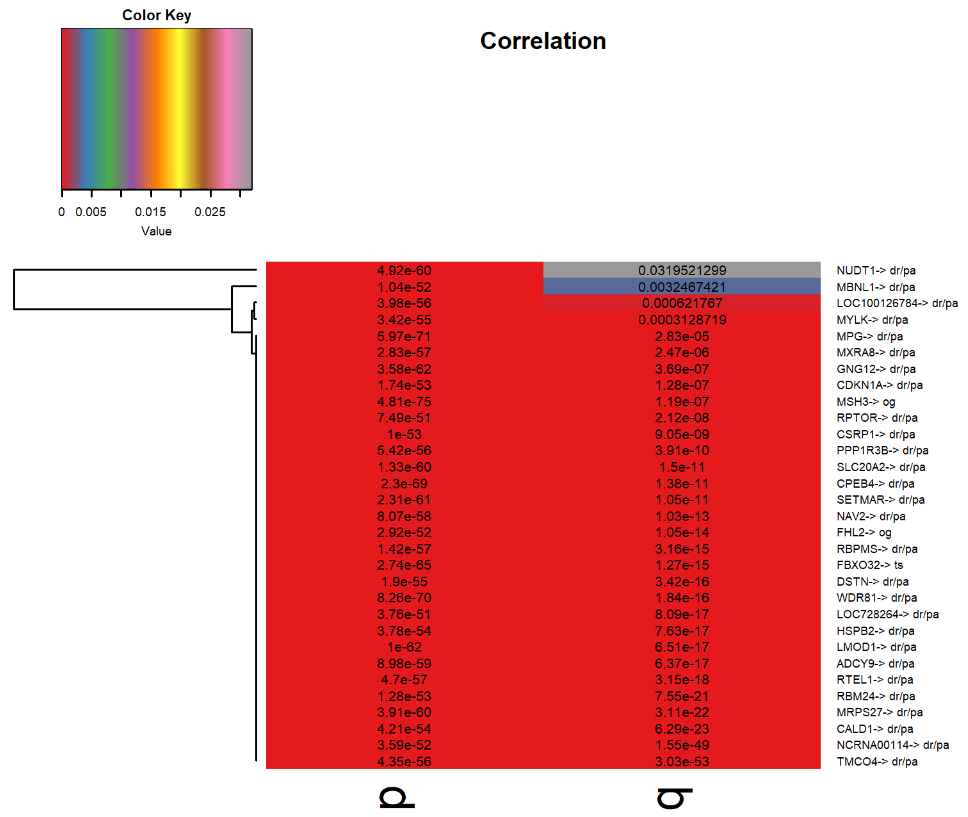

Figure S32: Marker genes heatmap for SARC

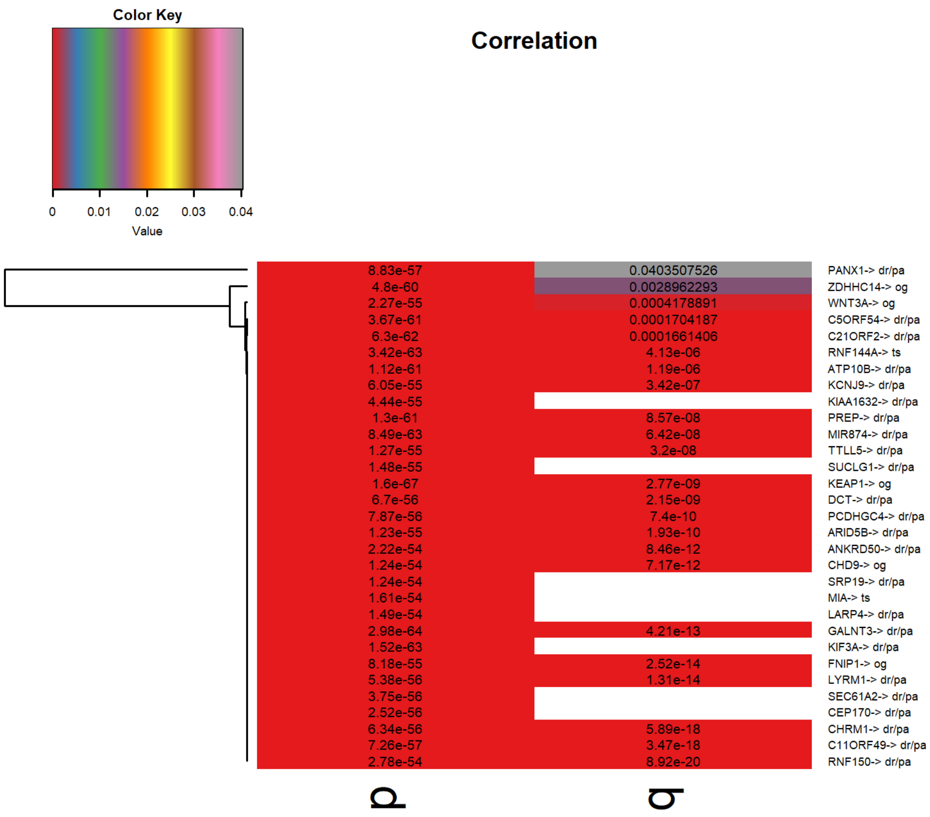

Figure S33: Marker genes heatmap for SKCM

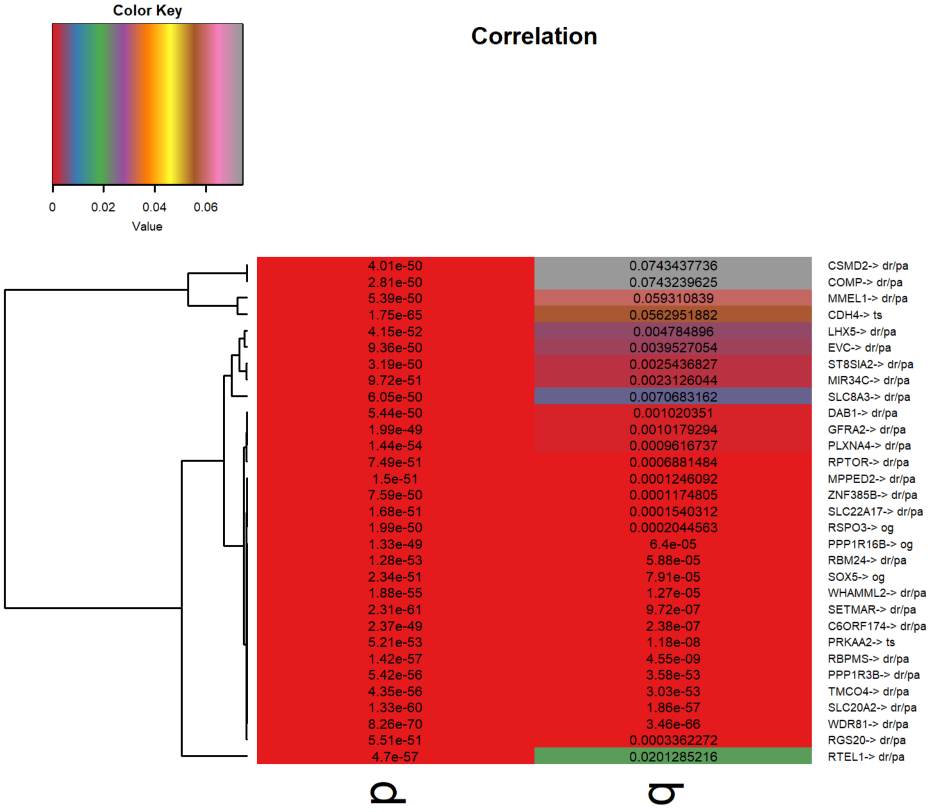

Figure S34: Marker genes heatmap for STAD

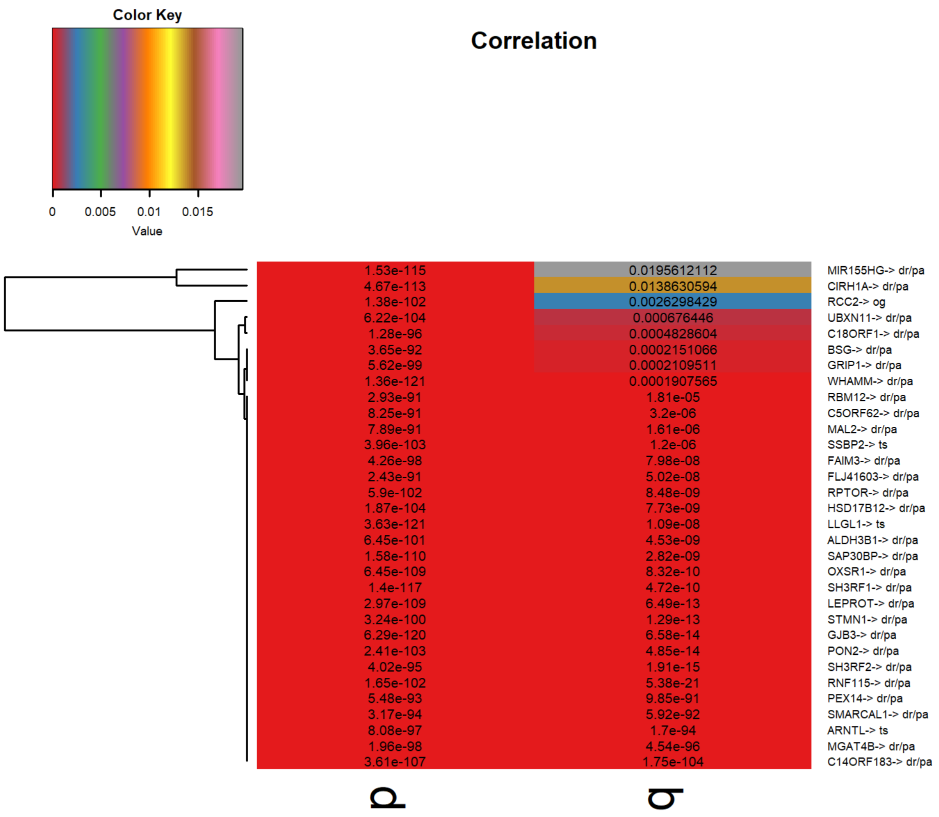

Figure S35: Marker genes heatmap for THCA

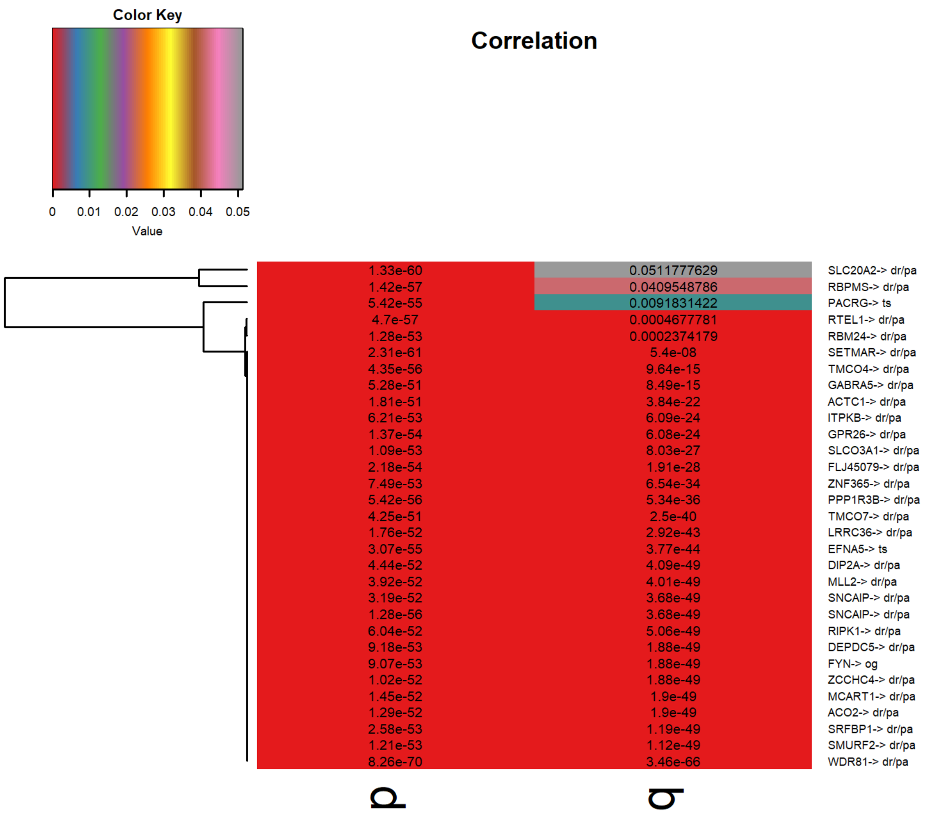

Figure S36: Marker genes heatmap for UCEC

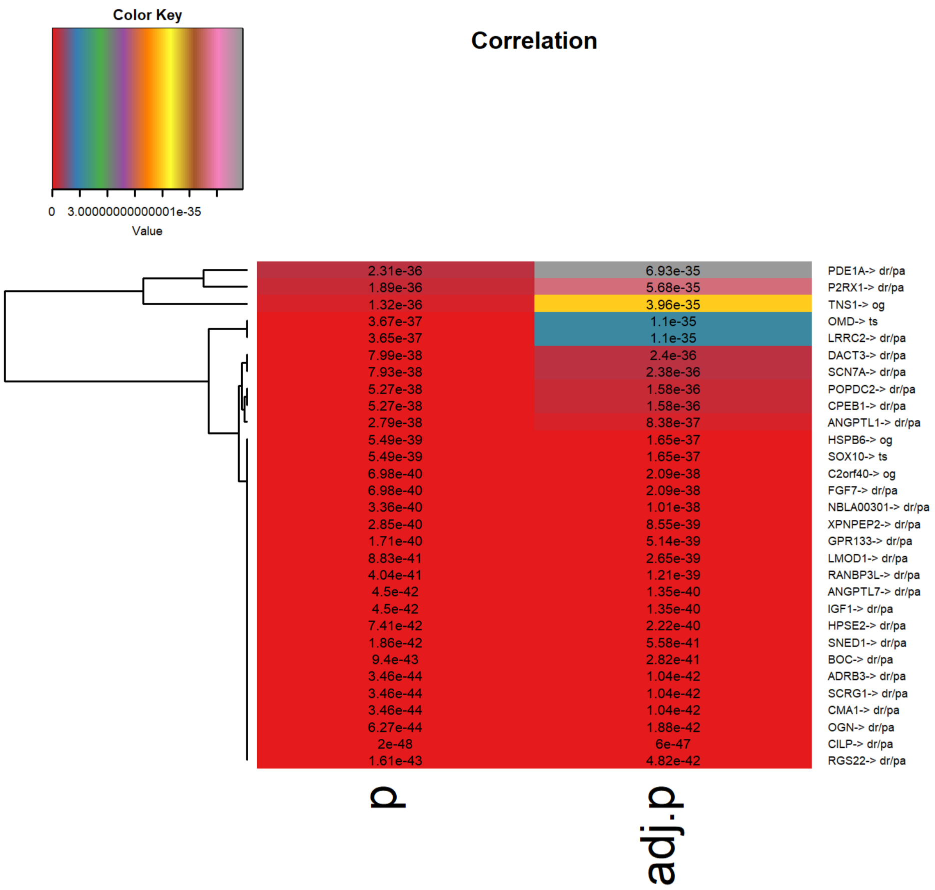

Figure S37: Significant genes heatmap for BLCA

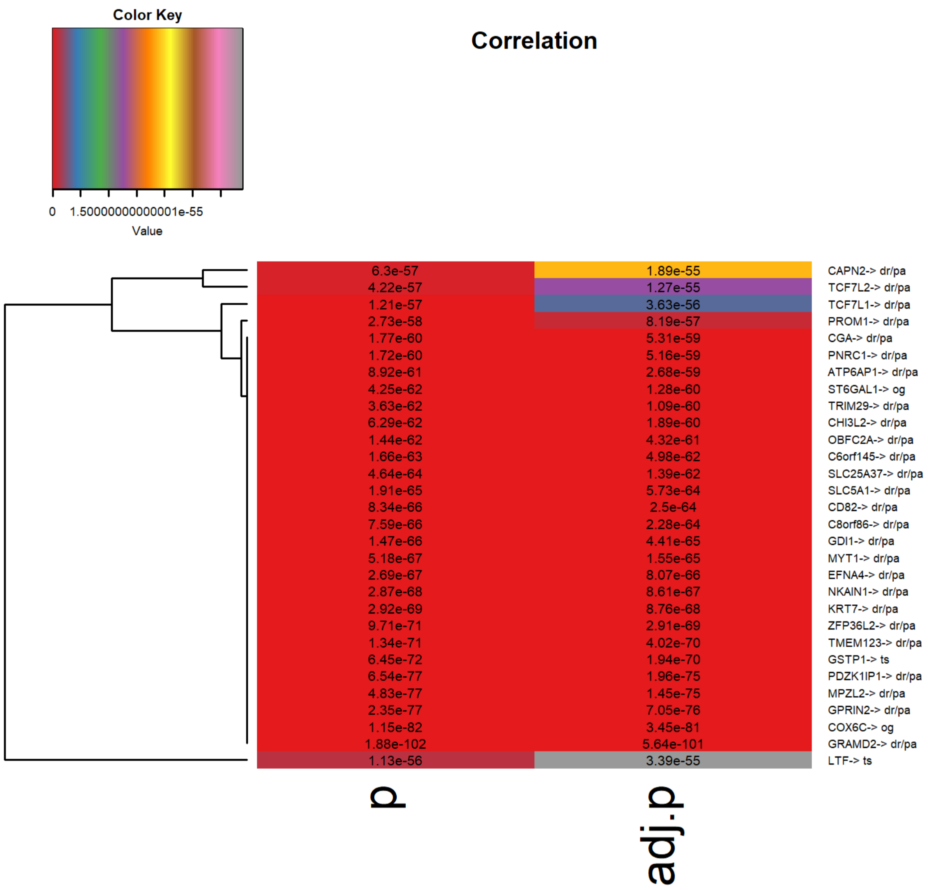

Figure S38: Significant genes heatmap for BRCA

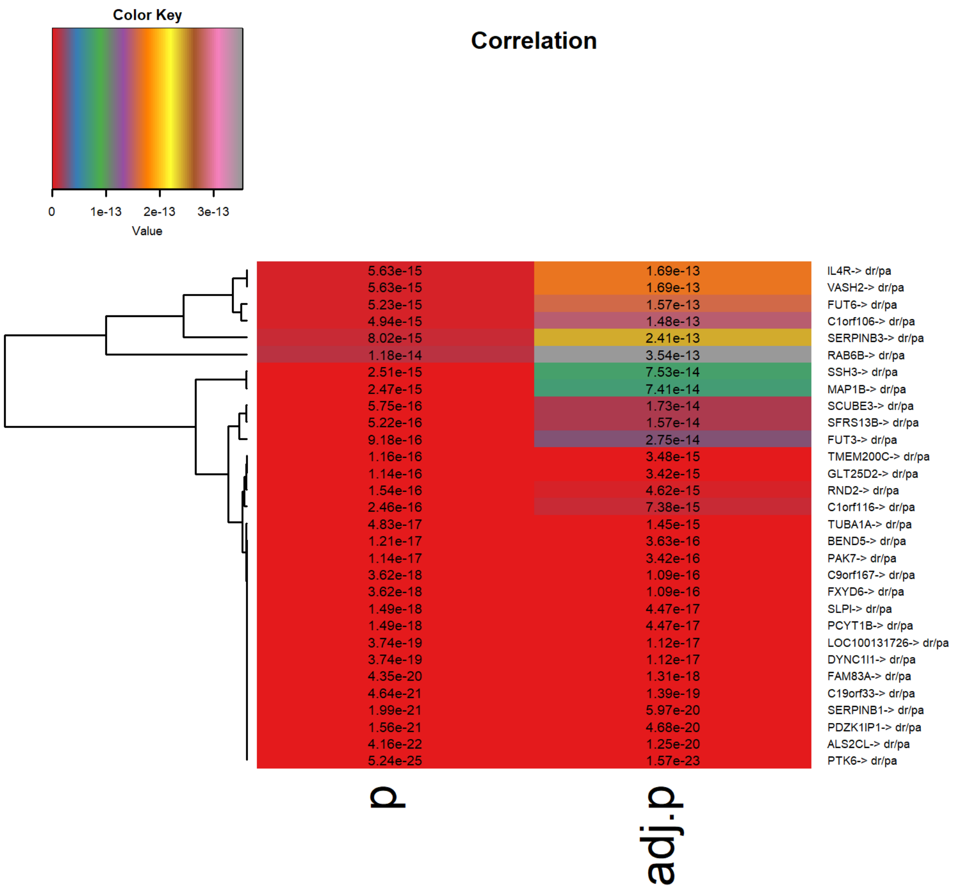

Figure S39: Significant genes heatmap for CESC

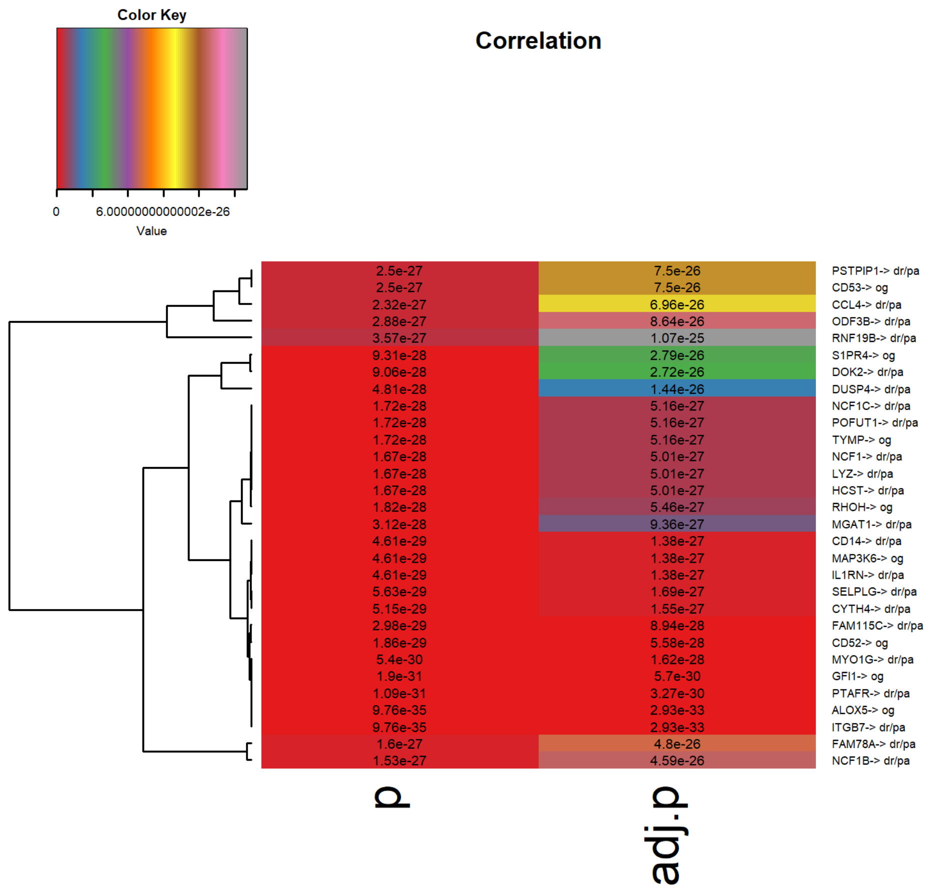

Figure S40: Significant genes heatmap for COAD

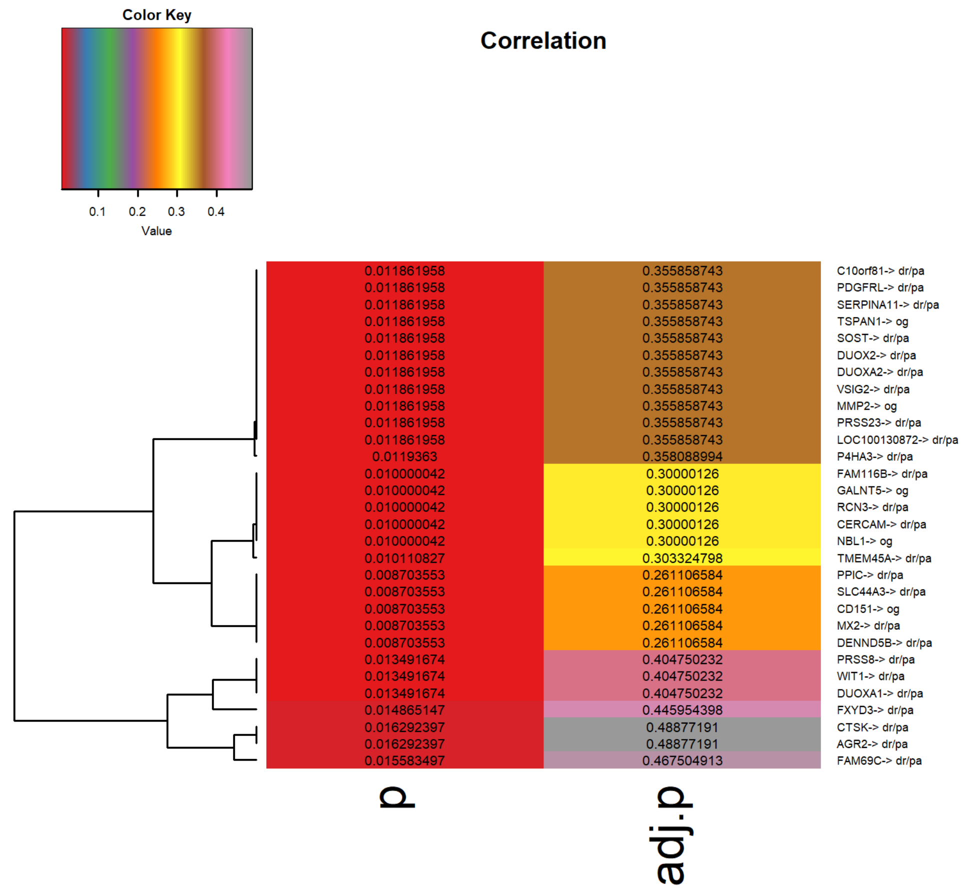

Figure S41: Significant genes heatmap for DLBC

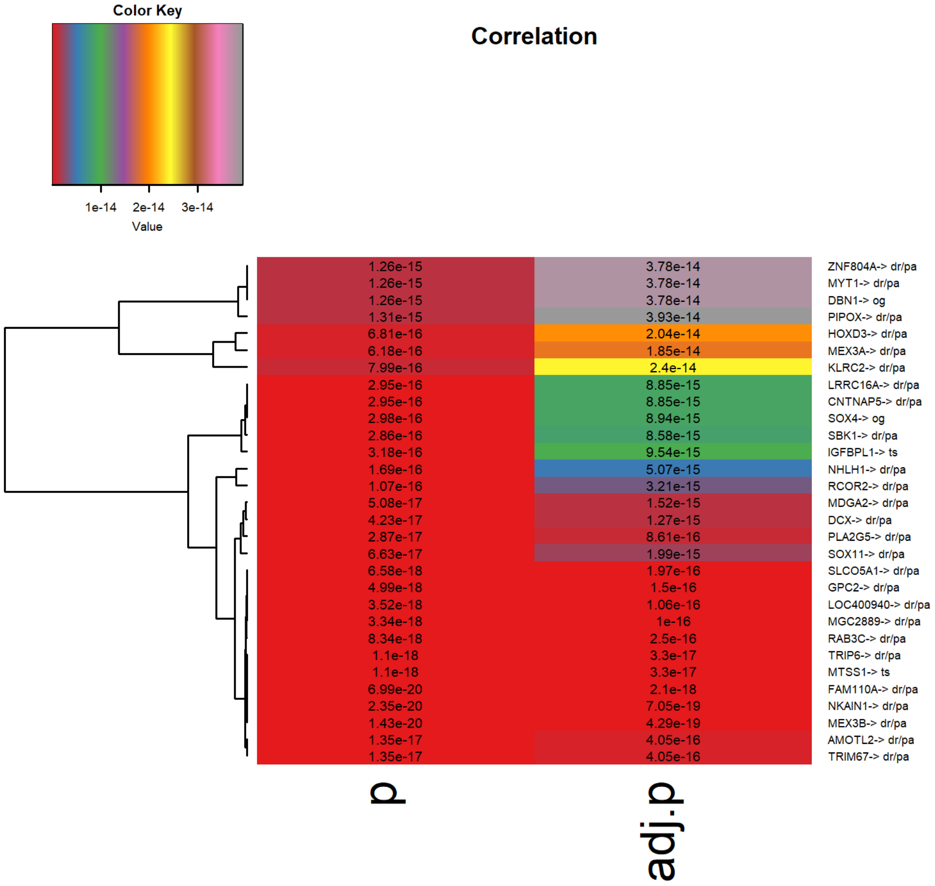

Figure S42: Significant genes heatmap for GBM

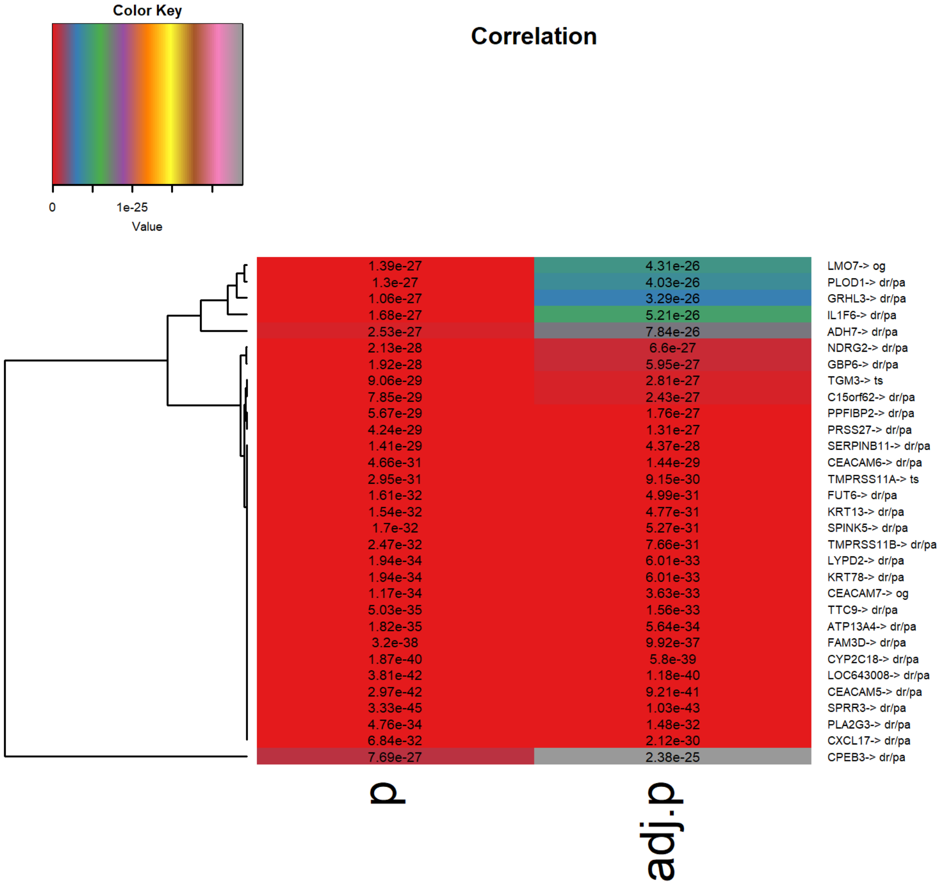

Figure S43: Significant genes heatmap for HNSC

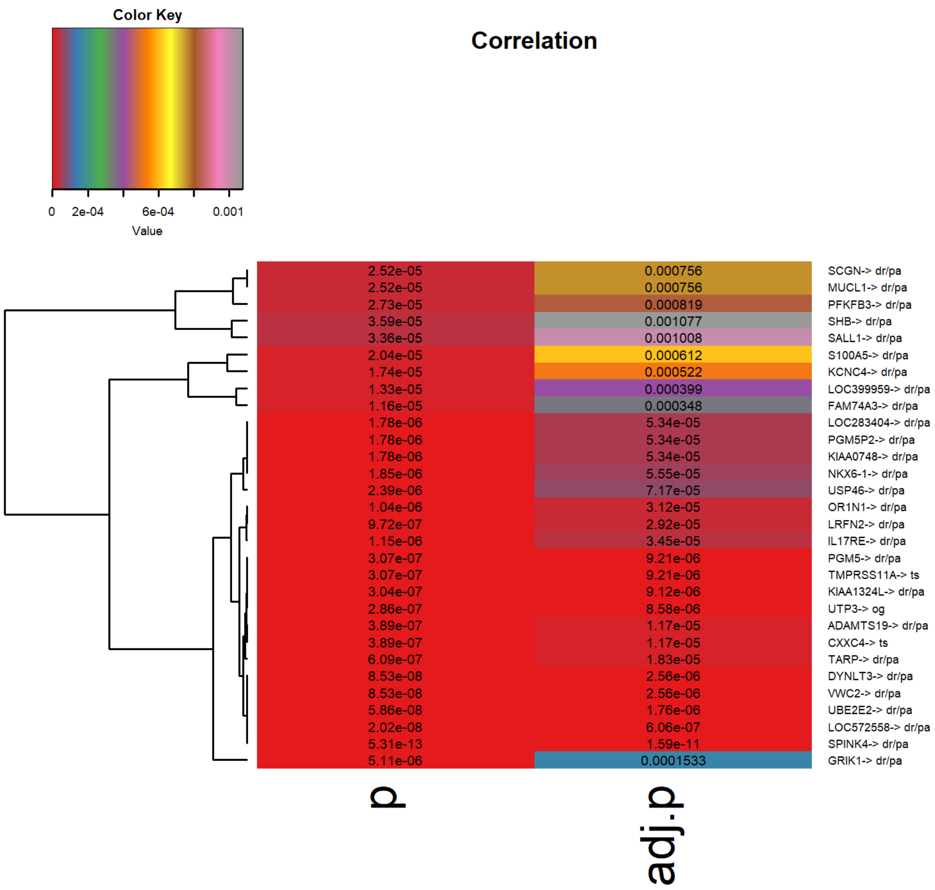

Figure S44: Significant genes heatmap for KICH

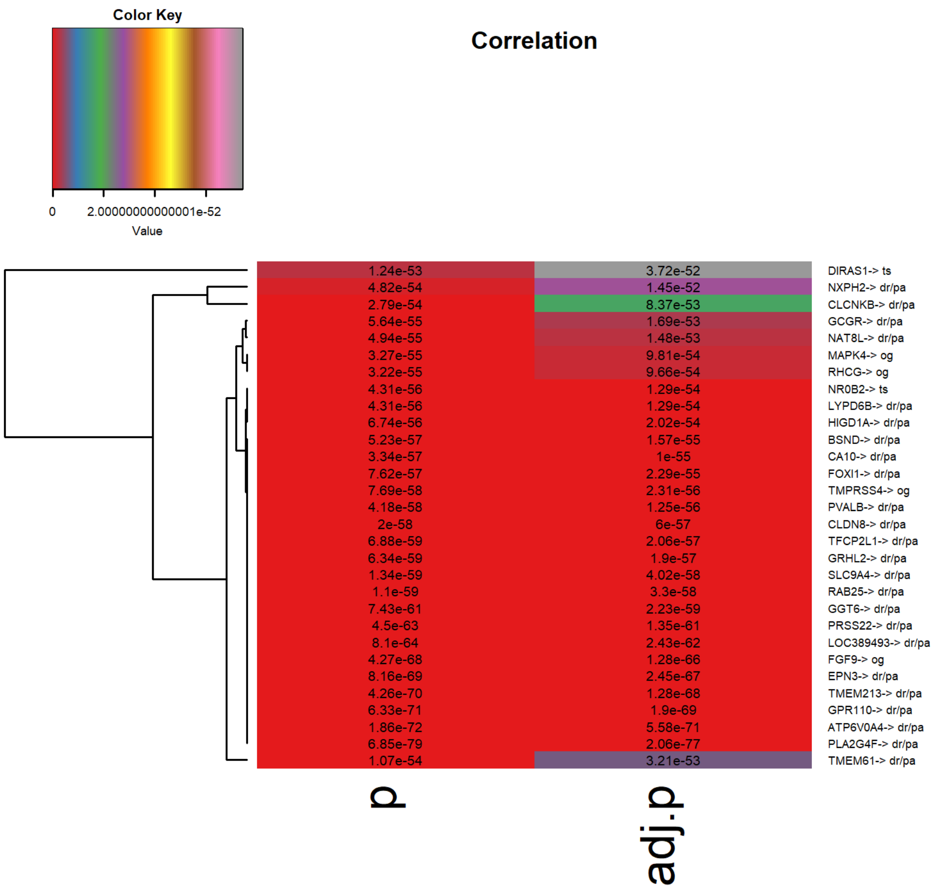

Figure S45: Significant genes heatmap for KIRC

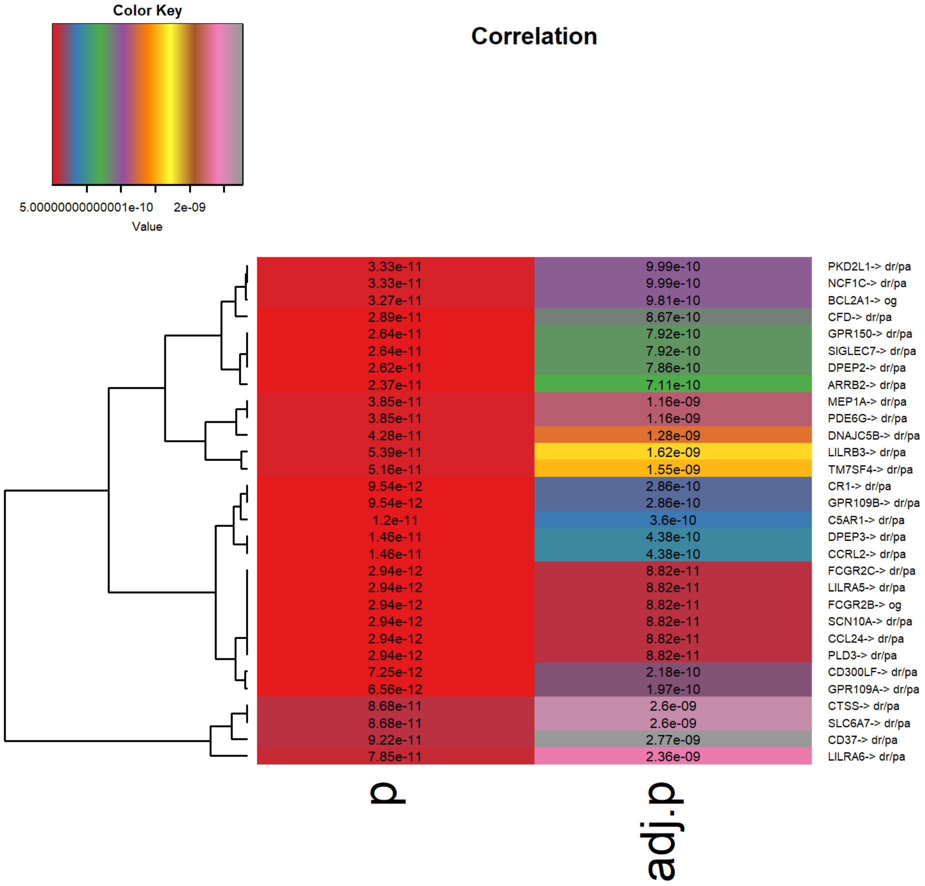

Figure S46: Significant genes heatmap for KIRP

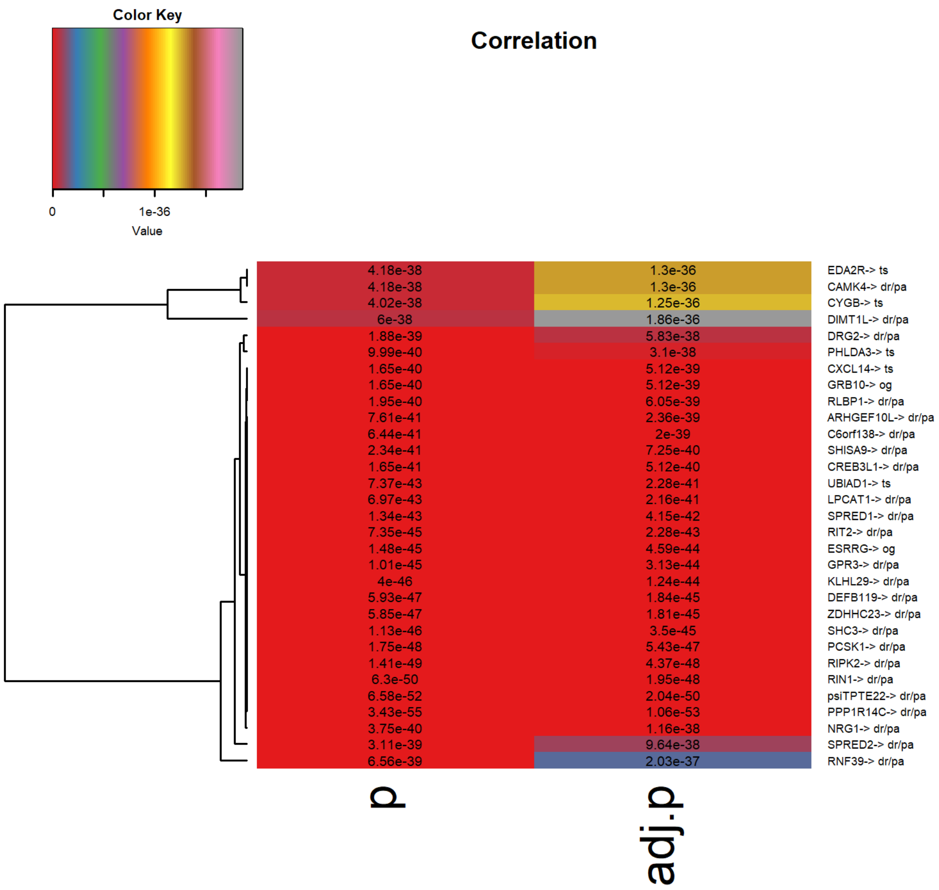

Figure S47: Significant genes heatmap for LGG

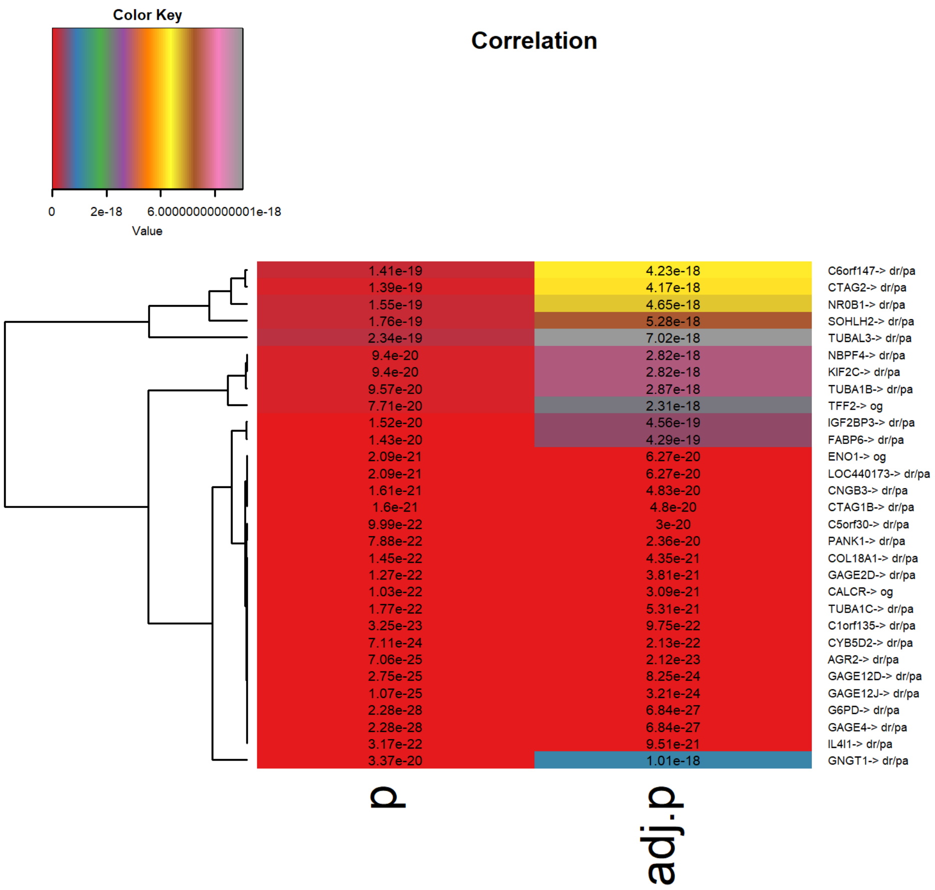

Figure S48: Significant genes heatmap for LIHC

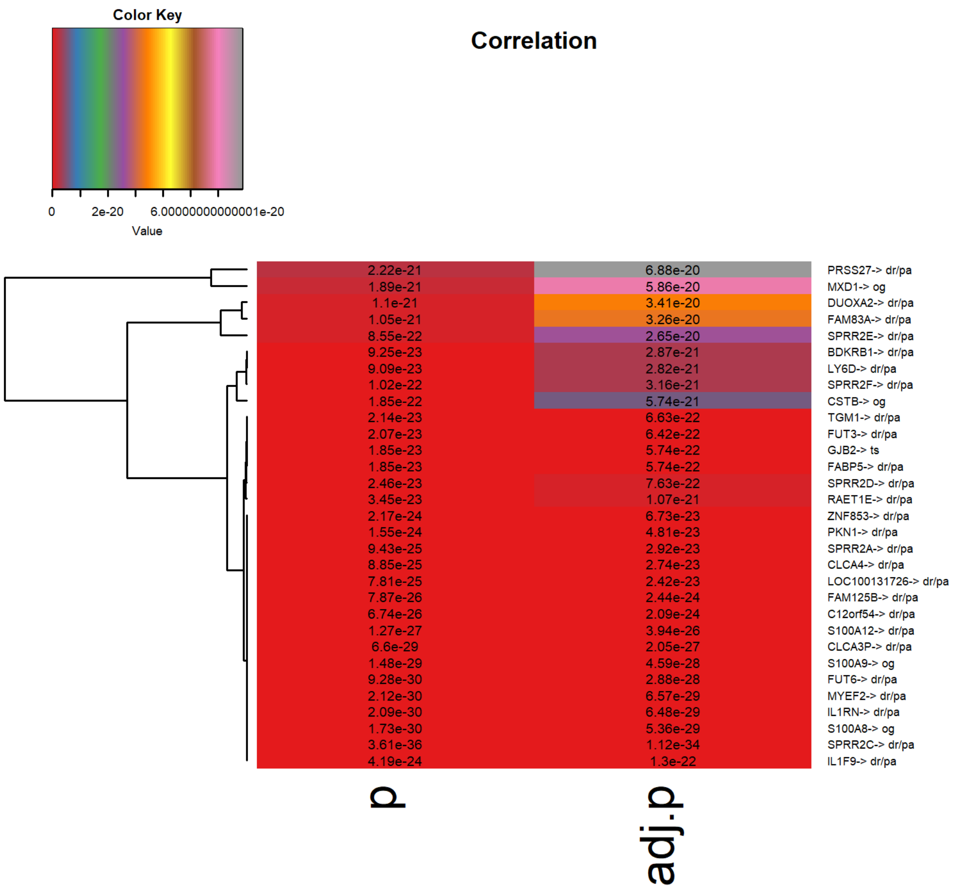

Figure S49: Significant genes heatmap for LUSC

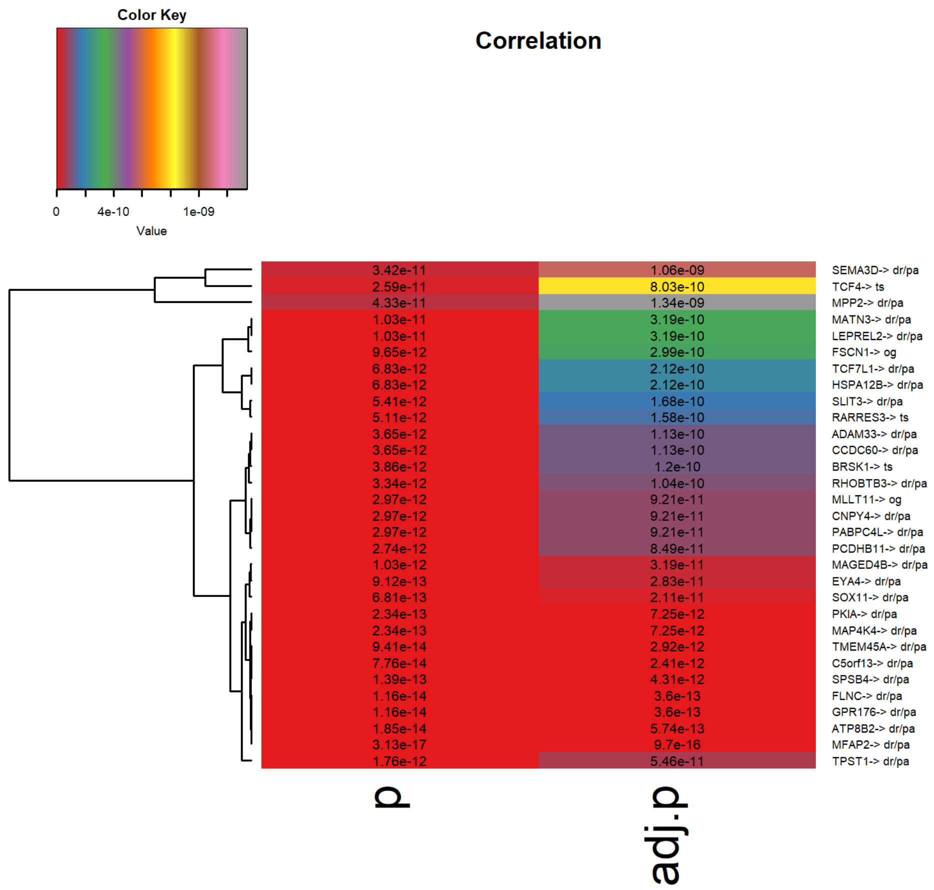

Figure S50: Significant genes heatmap for OV

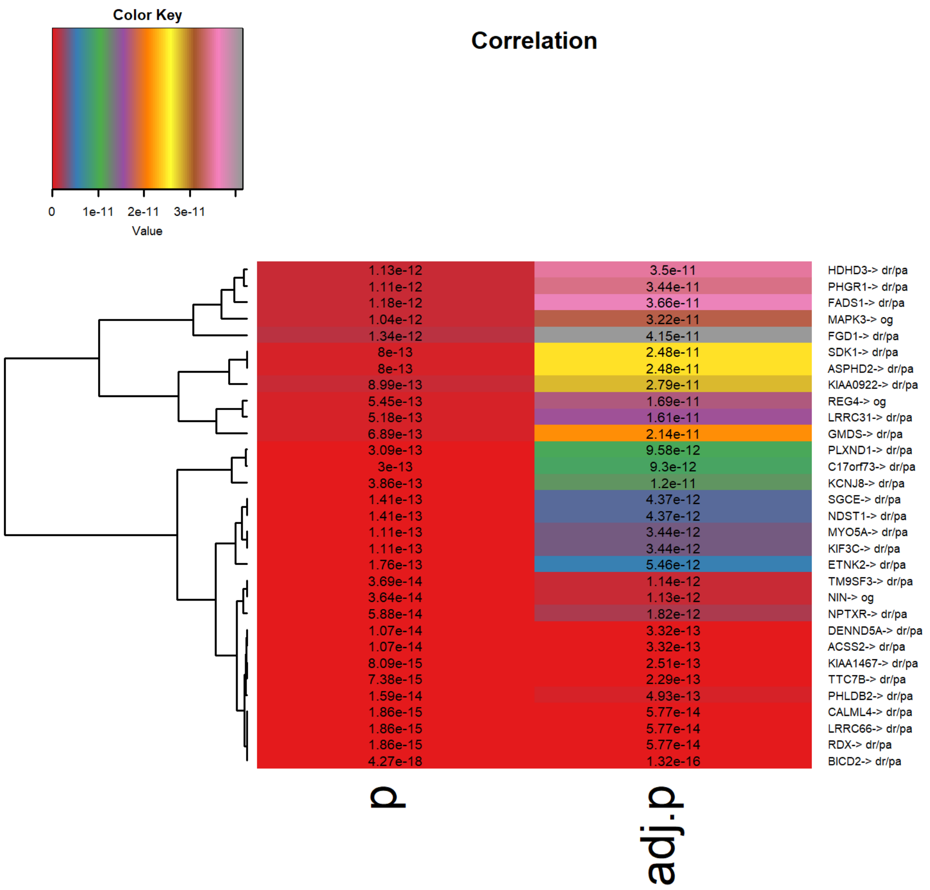

Figure S51: Significant genes heatmap for PACA

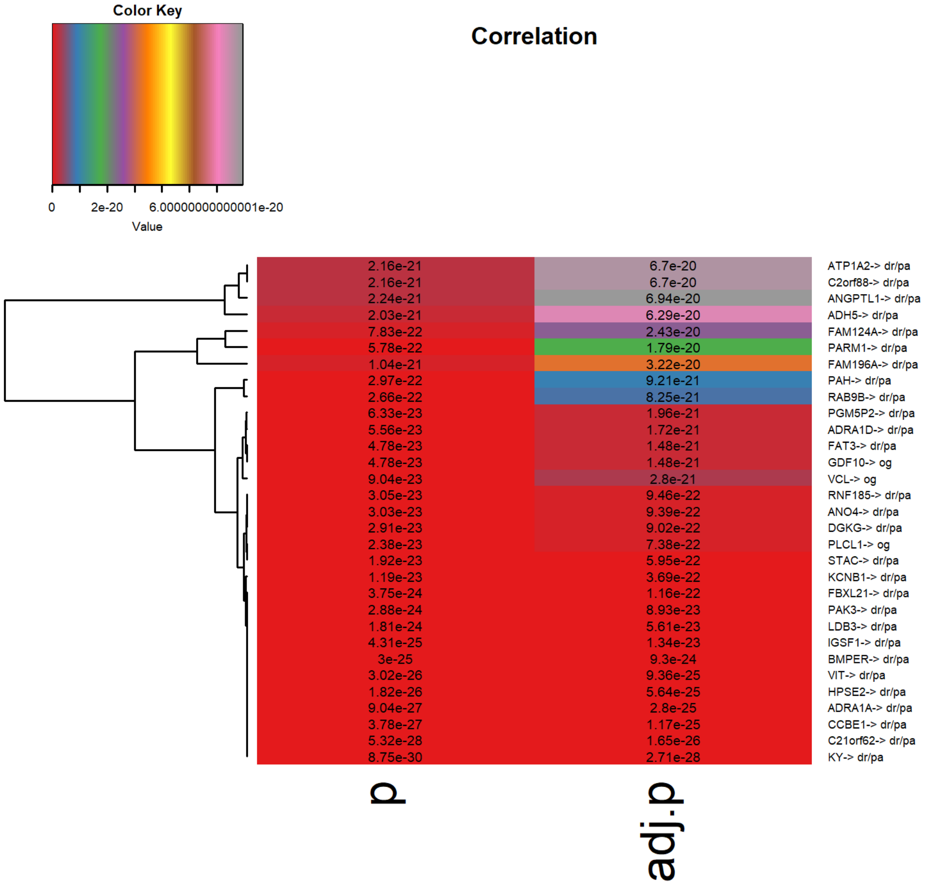

Figure S52: Significant genes heatmap for PRAD

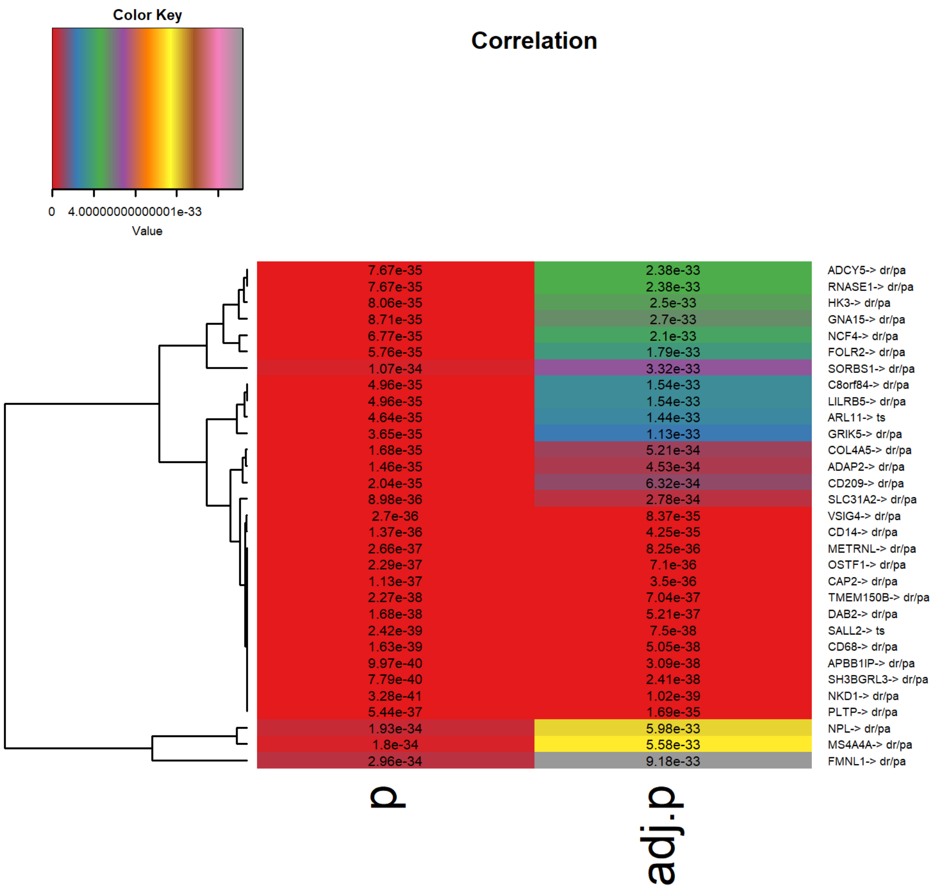

Figure S53: Significant genes heatmap for SARC

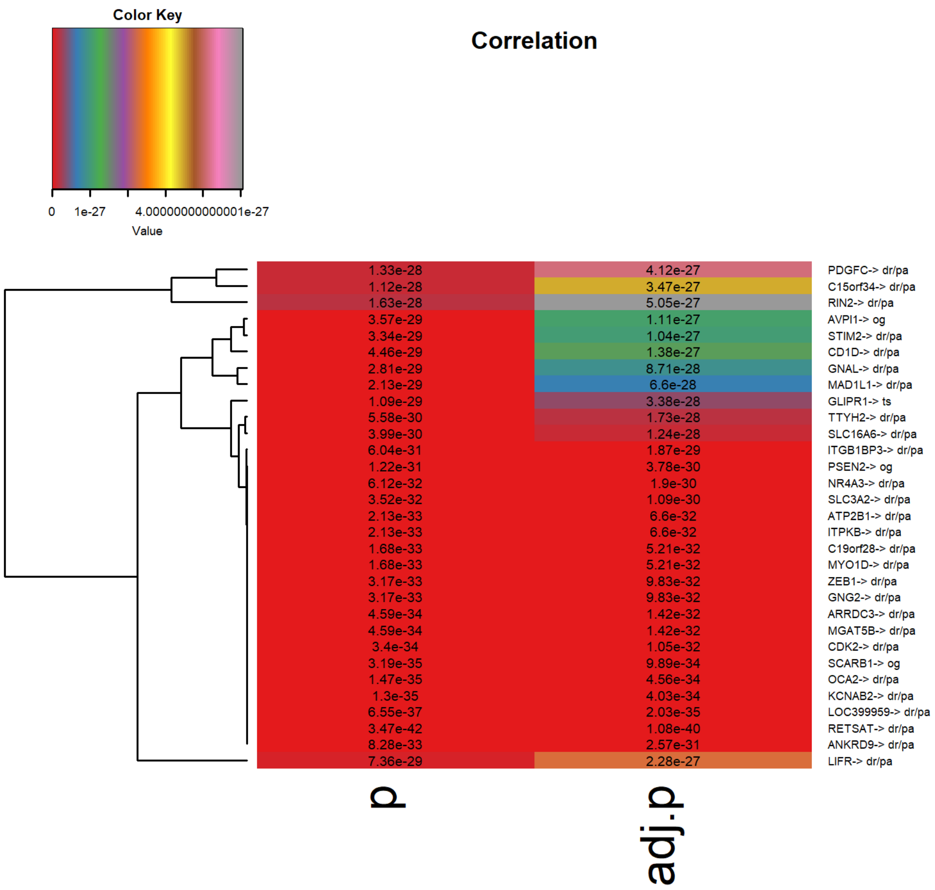

Figure S54: Significant genes heatmap for SKCM

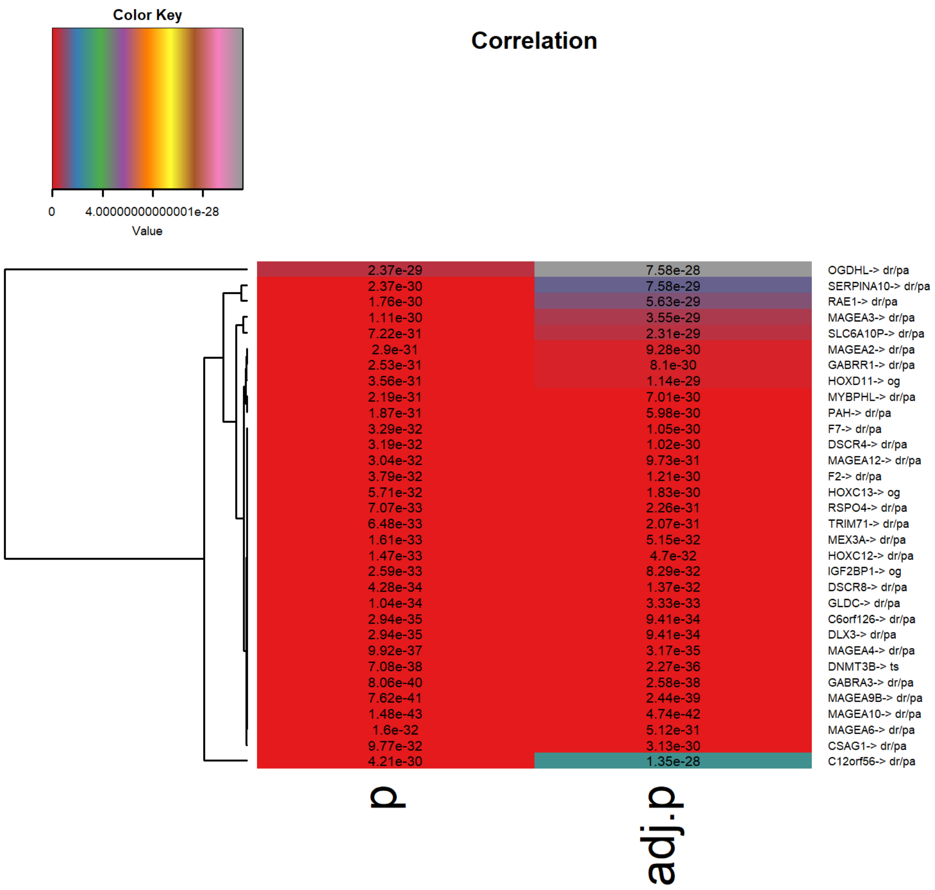

Figure S55: Significant genes heatmap for STAD

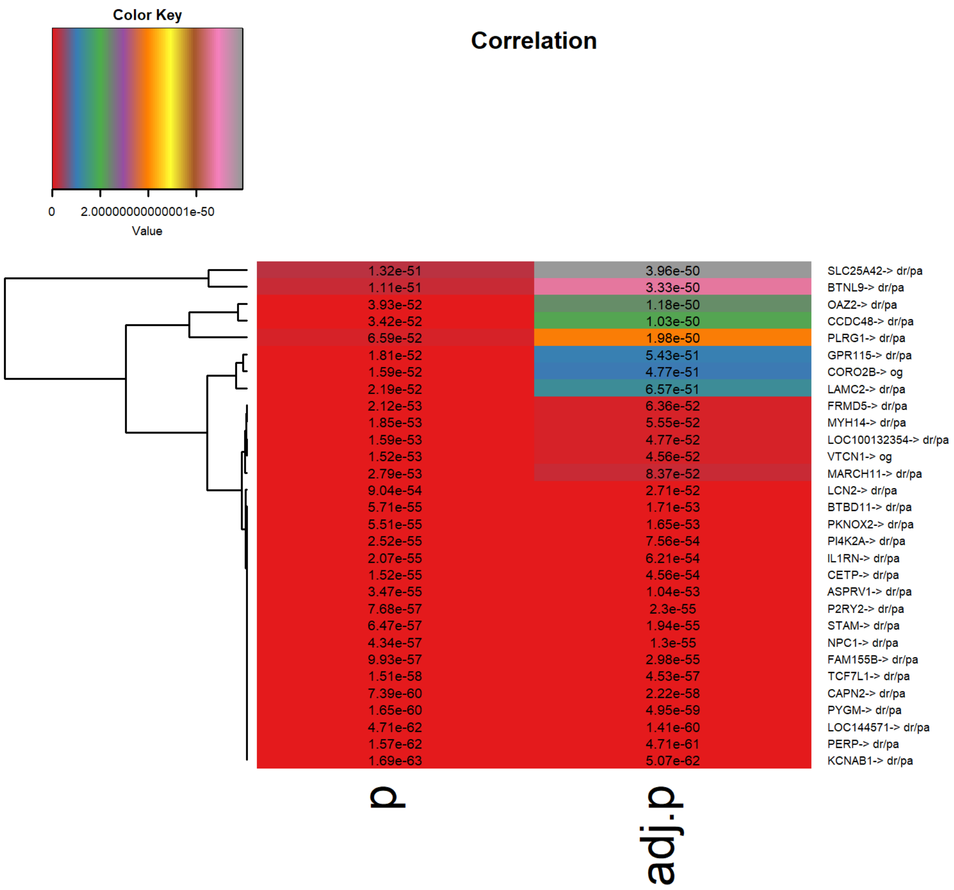

Figure S56: Significant genes heatmap for THCA

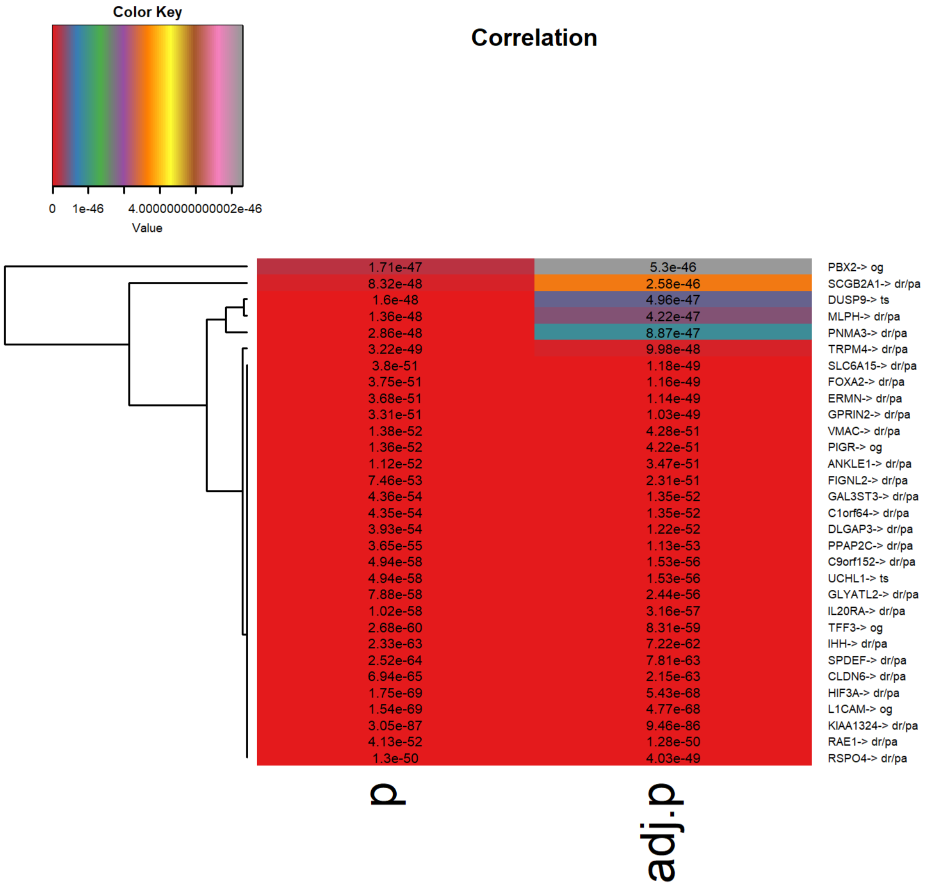

Figure S57: Significant genes heatmap for UCEC

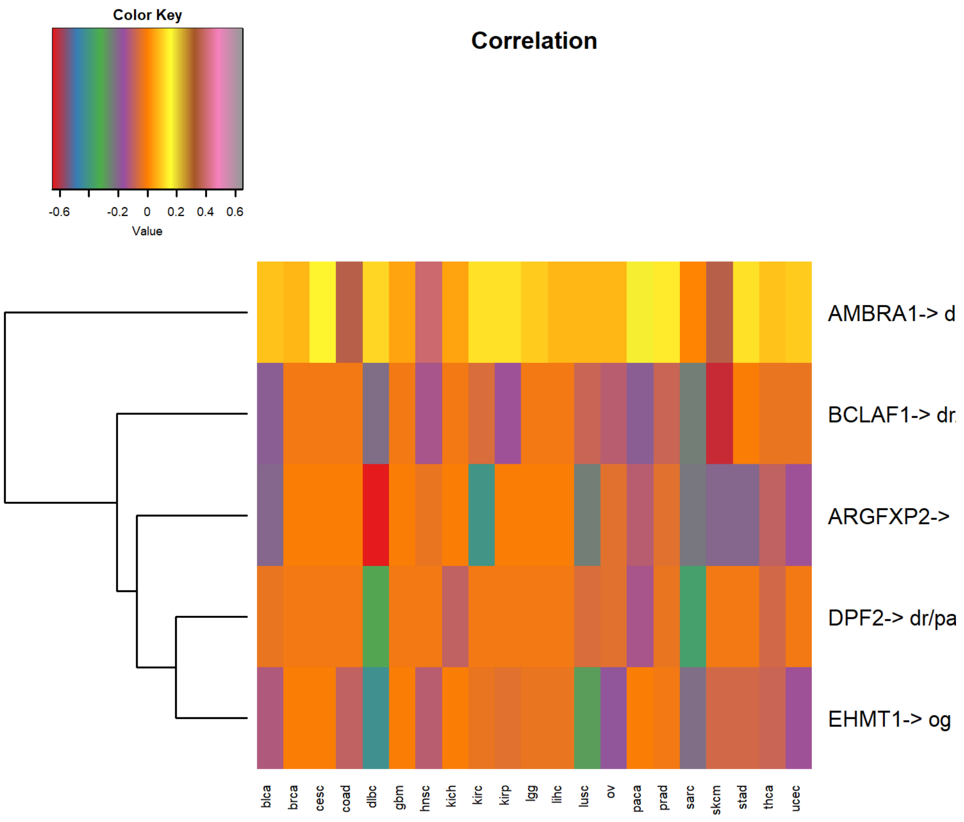

Figure S58: Common up-down regulated genes in all 21 cancer types

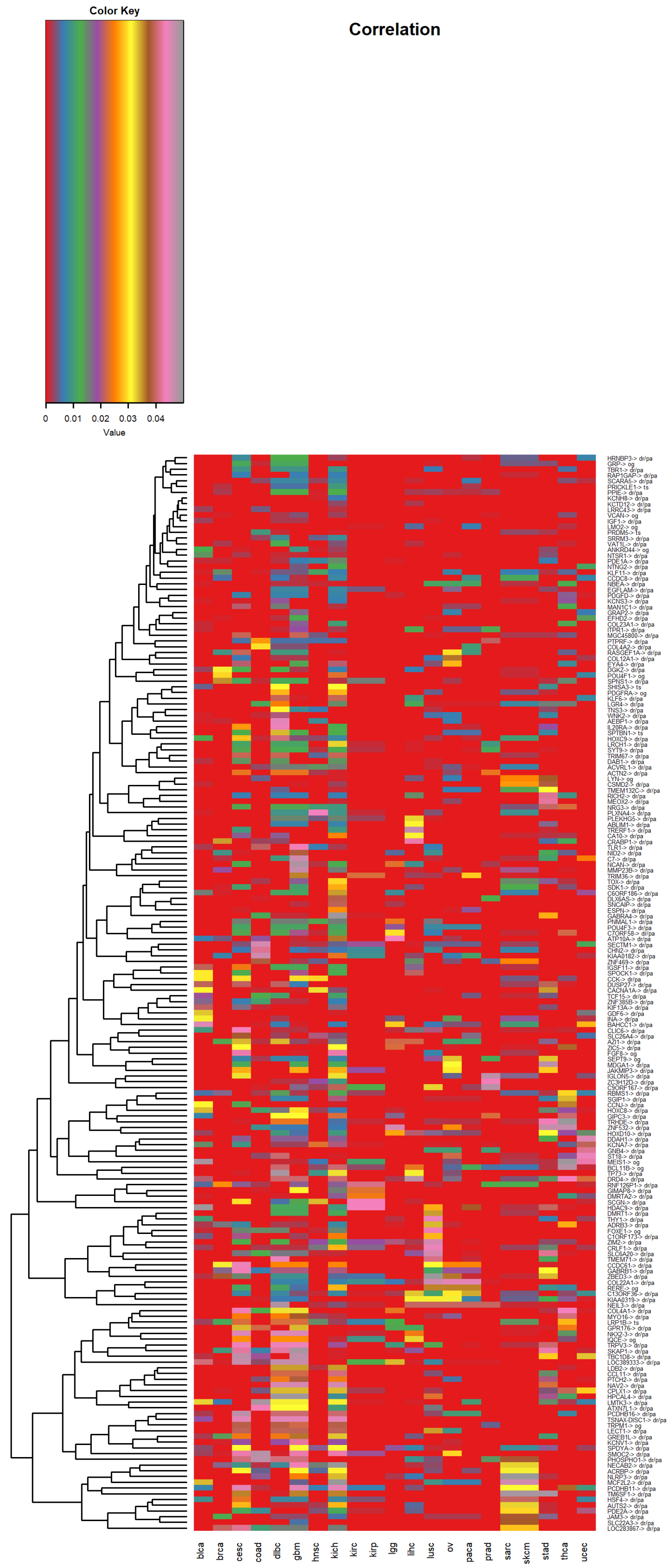

Figure S59: Common marker genes in all 21 cancer types

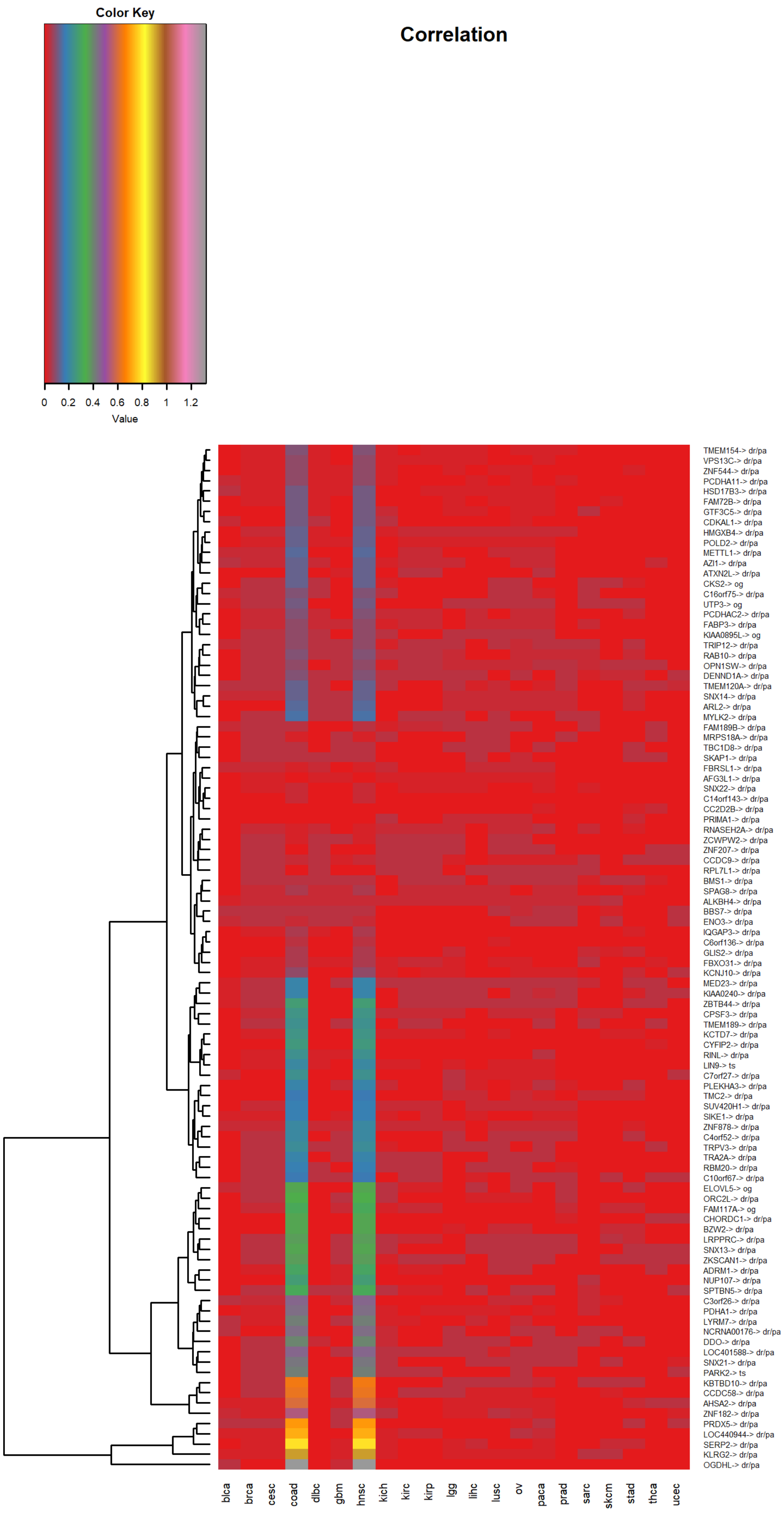

Figure S60: Common significant genes in all 21 cancer types

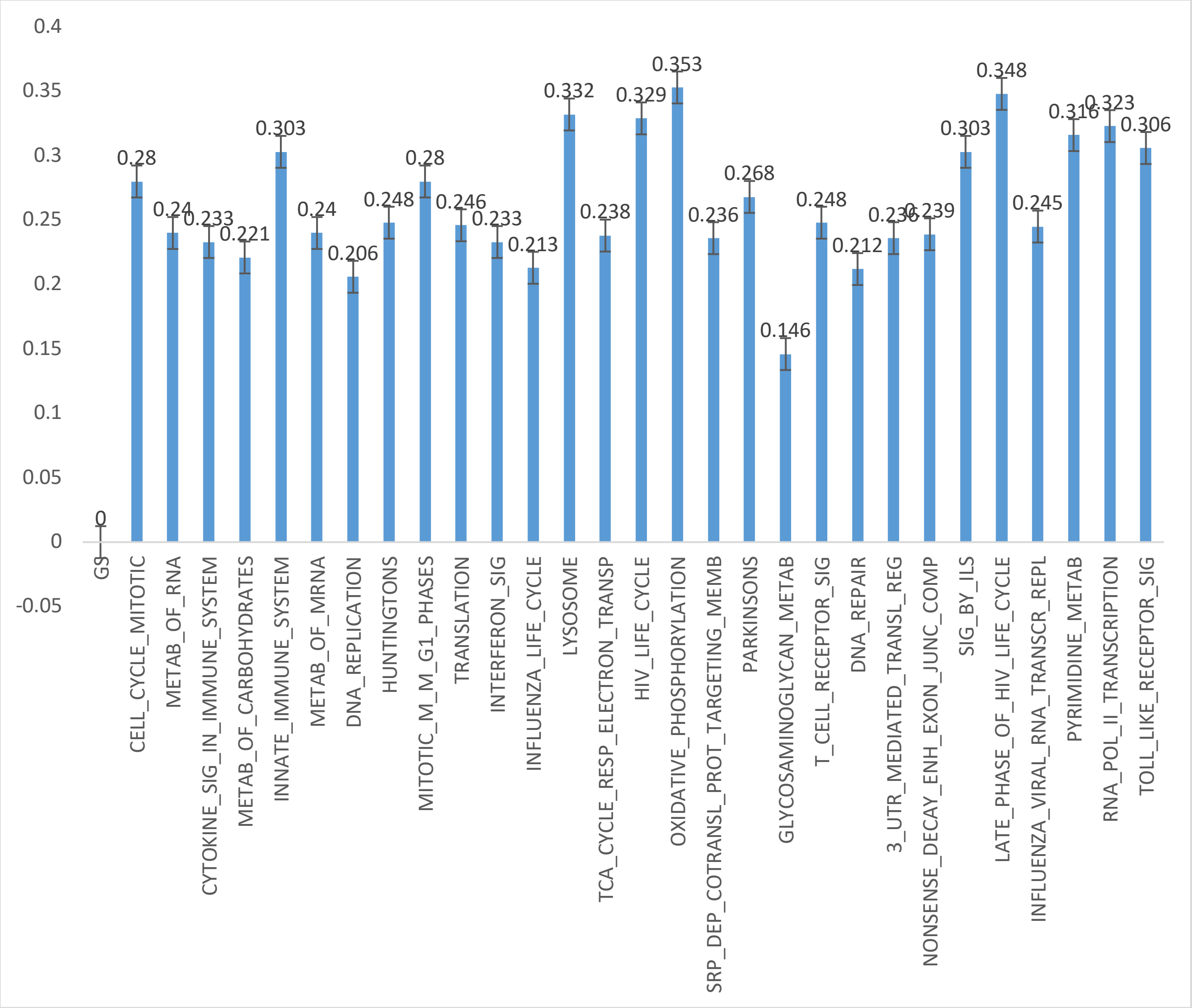

Figure S61: Enriched pathways in DLBC

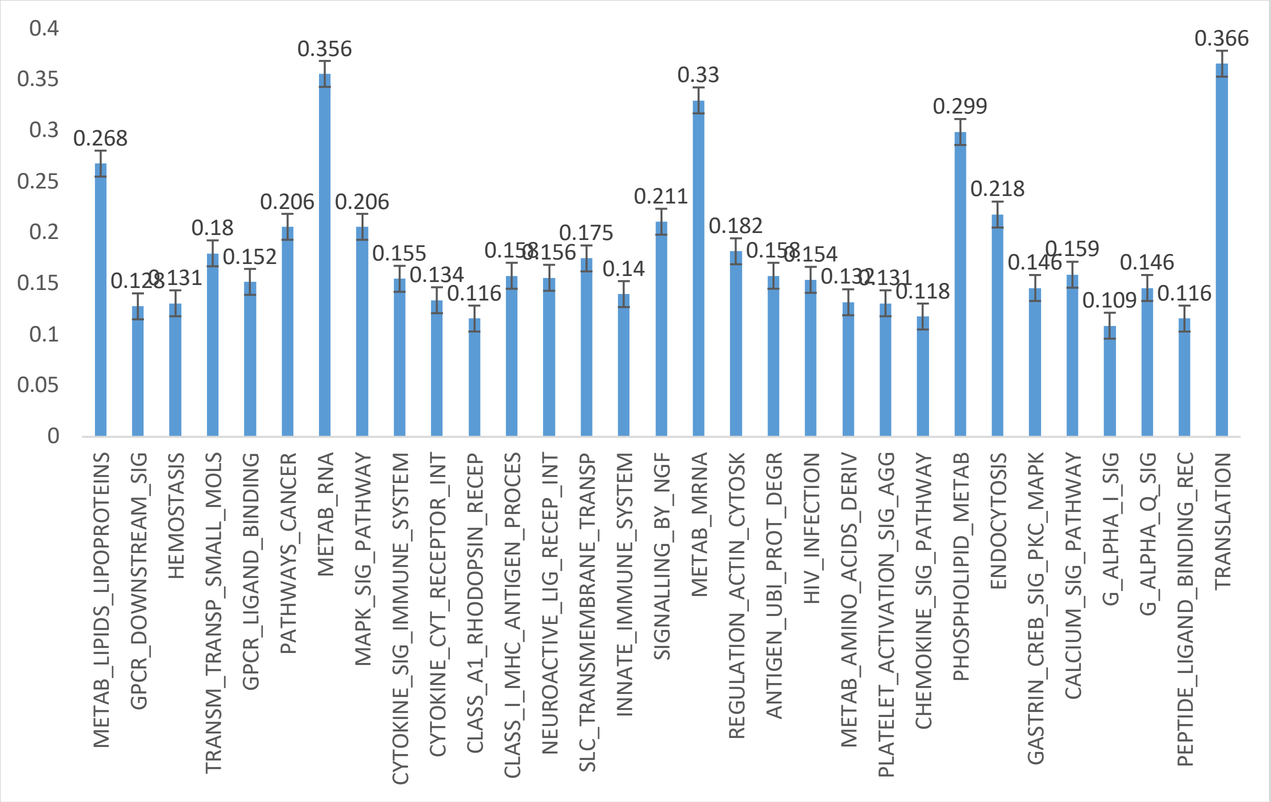

Figure S62: Enriched pathways in SKCM

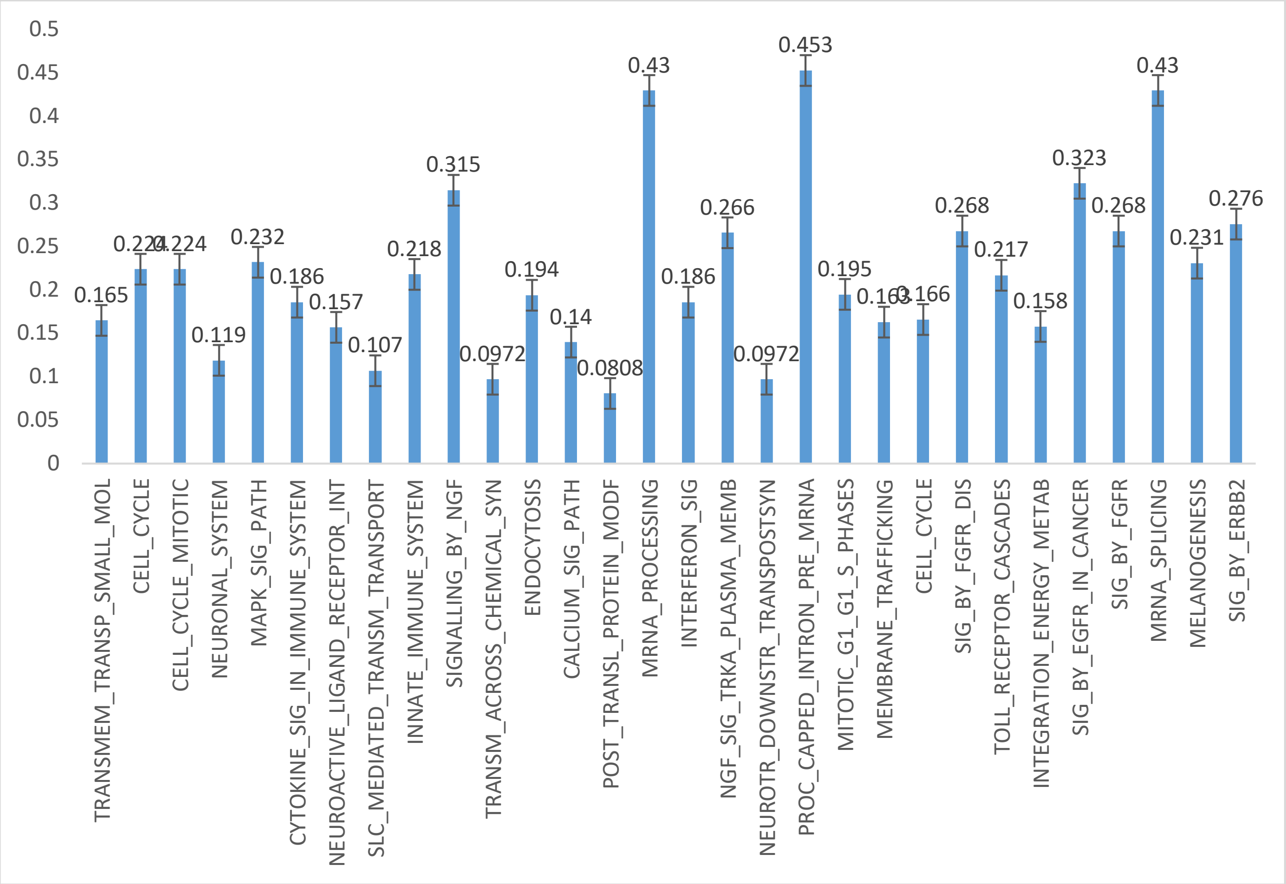

Figure S63: Enriched pathways in GBM

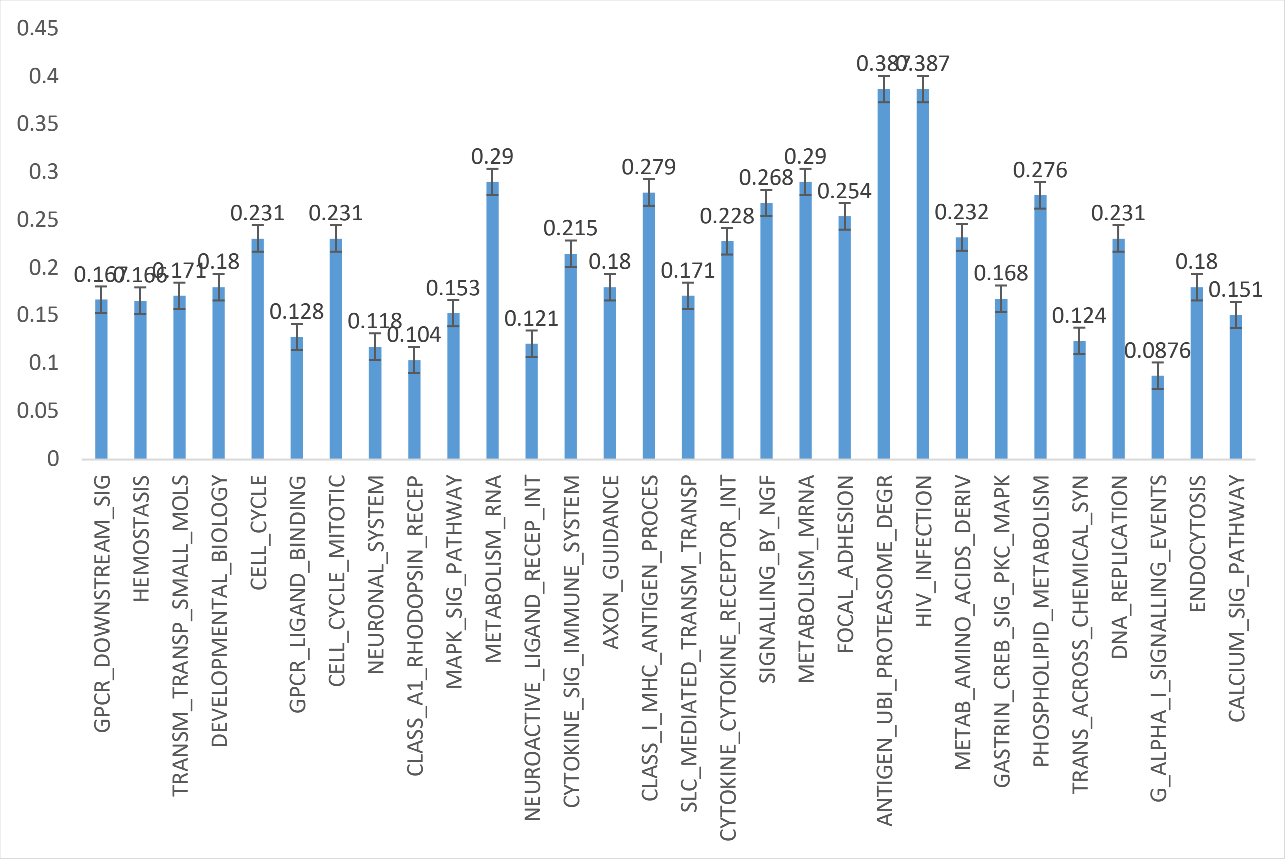

Figure S64: Enriched pathways in LGG

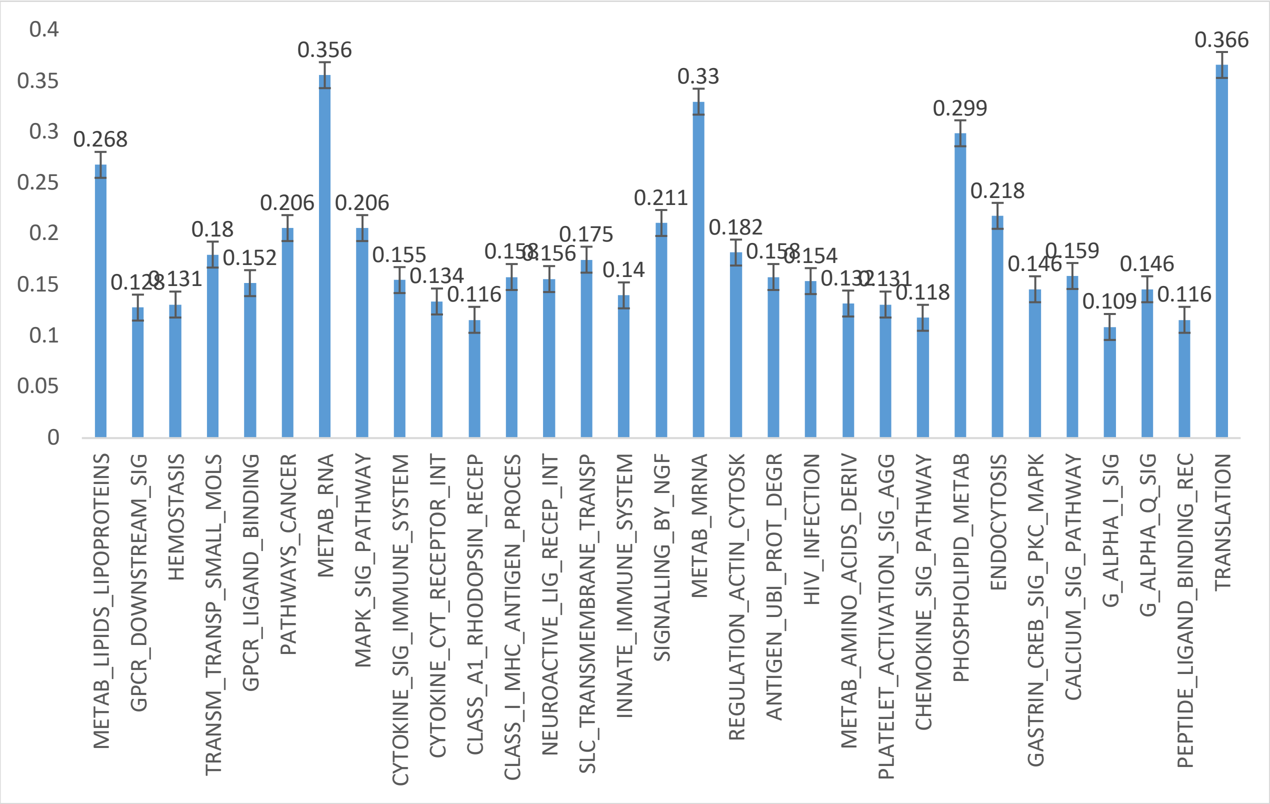

Figure S65: Enriched pathways in SKCM

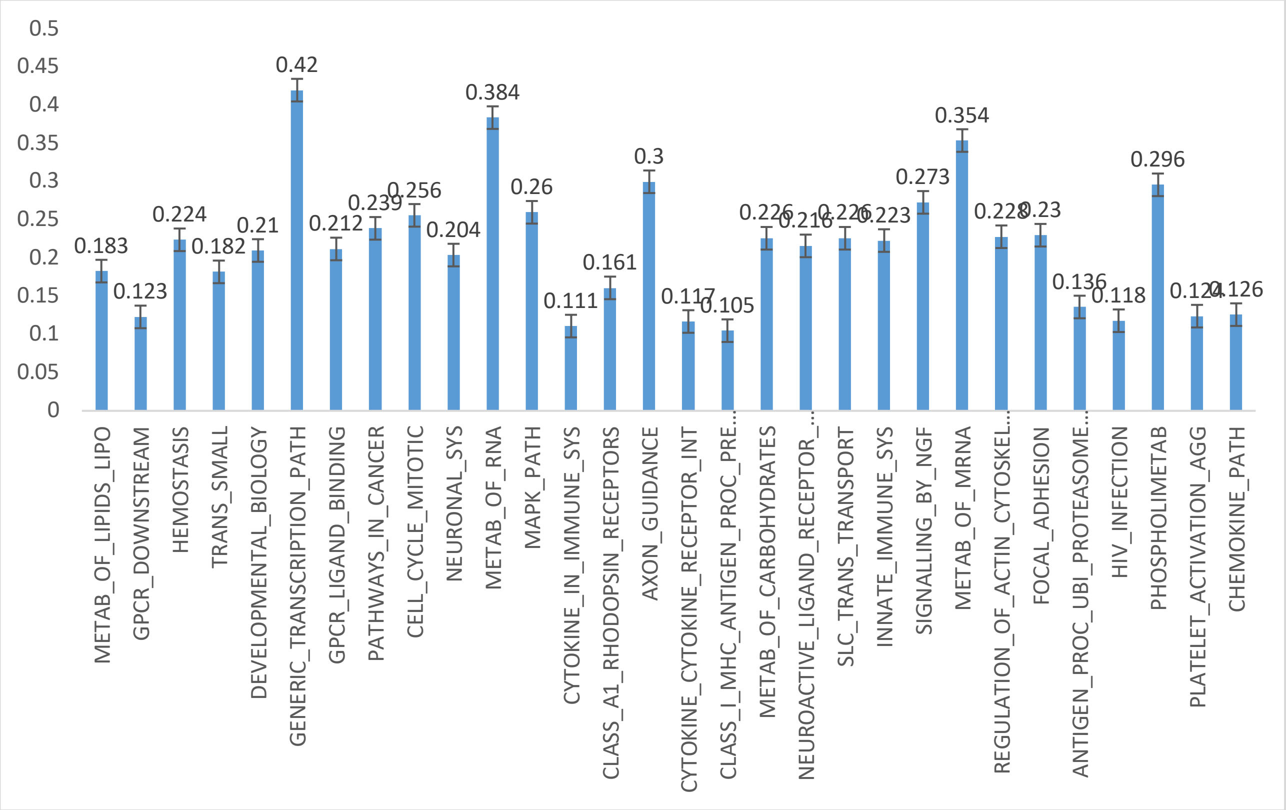

Figure S66: Enriched pathways in BRCA

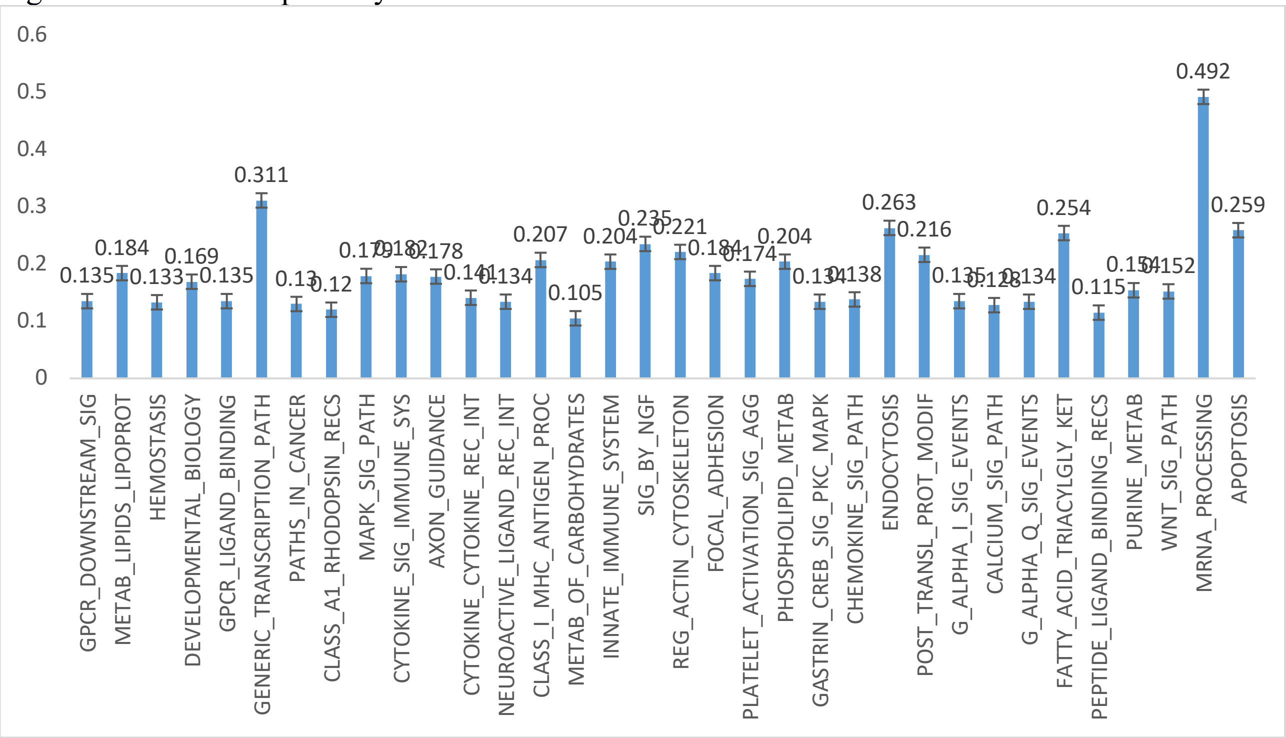

Figure S67: Enriched pathways in OV

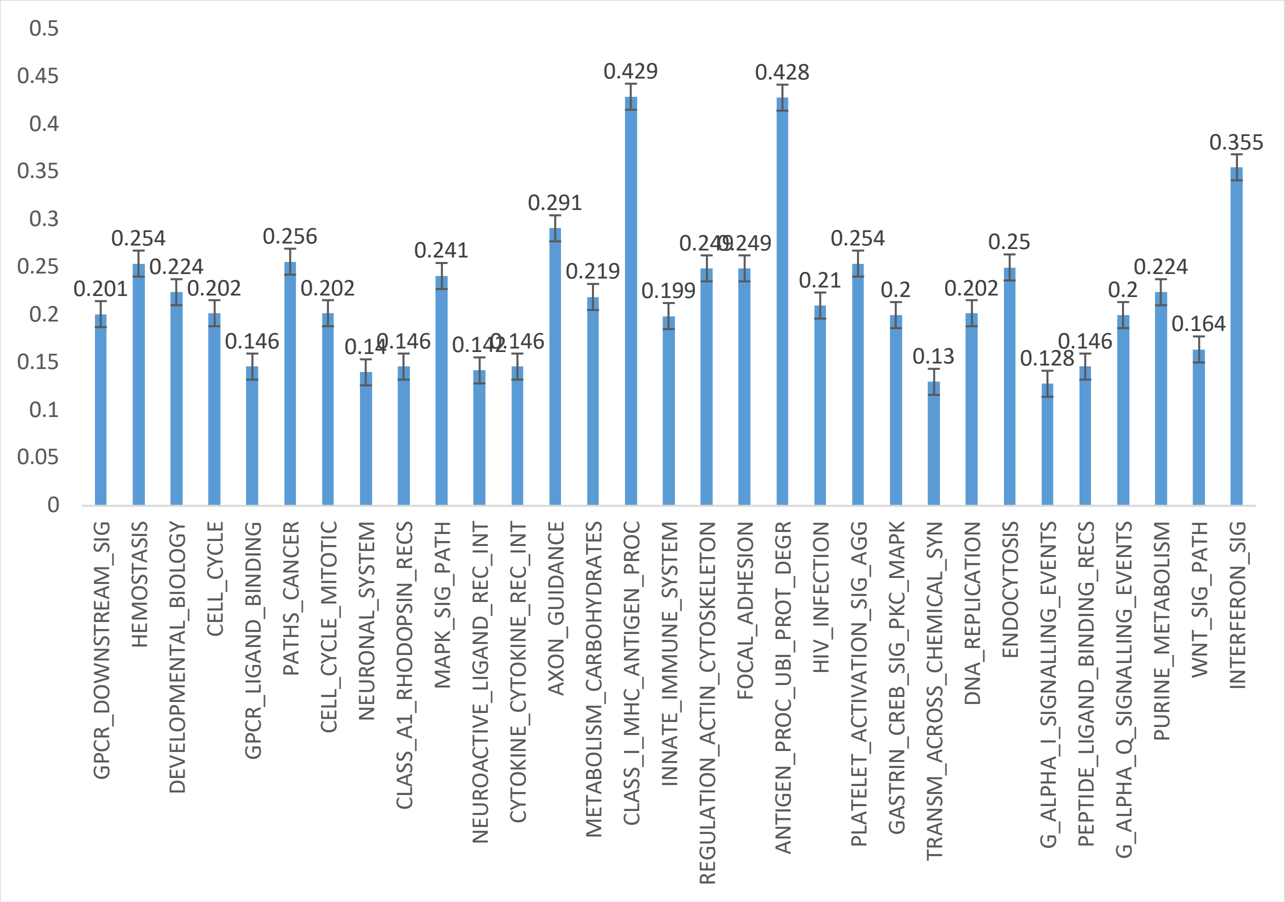

Figure S68: Enriched pathways in PACA

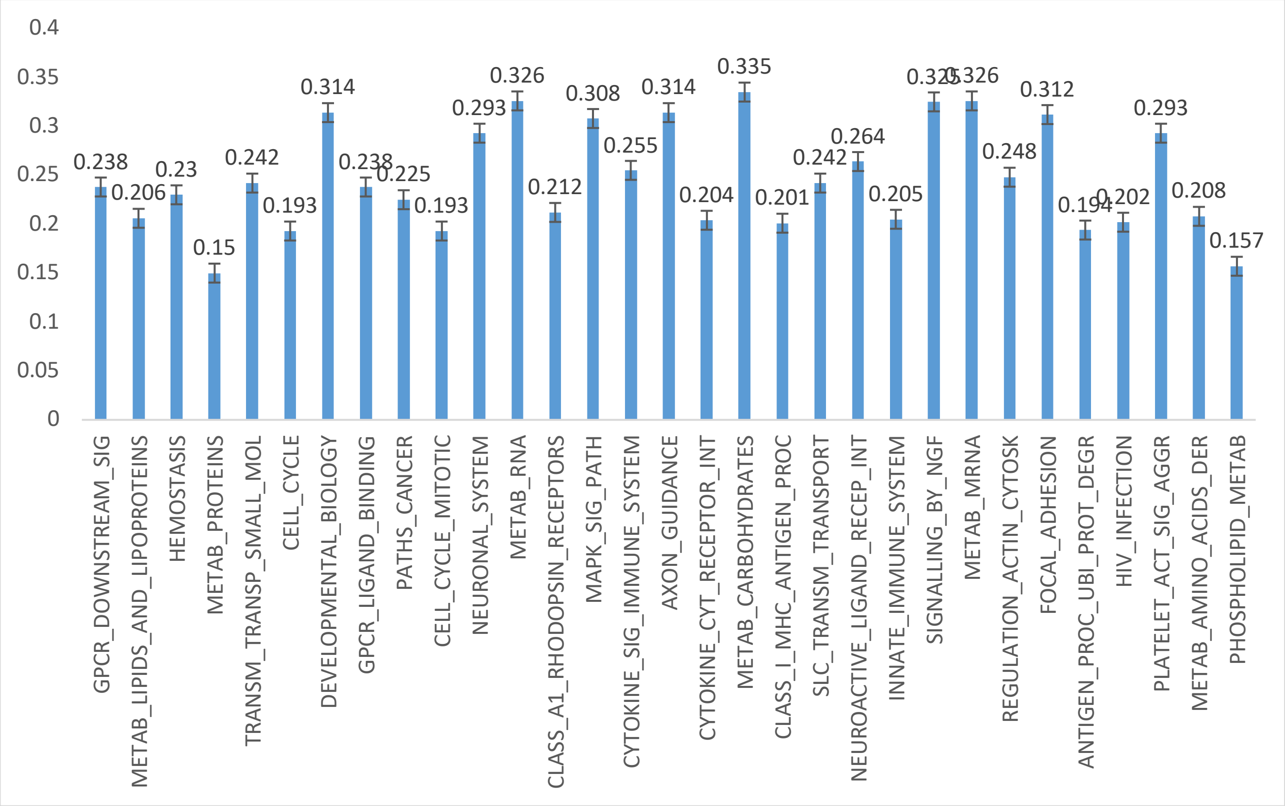

Figure S69: Enriched pathways in PRAD

Table S1: OG list (Known)

| symbol | info |
|---|---|
| NAT1 | N-acetyltransferase 1 |
| NAT2 | N-acetyltransferase 2 |
| ABCA1 | ATP binding cassette subfamily A member 1 |
| ABCA3 | ATP binding cassette subfamily A member 3 |
| ABL1 | ABL proto-oncogene 1, non-receptor tyrosine kinase |
| ABL2 | ABL proto-oncogene 2, non-receptor tyrosine kinase |
| ABO | ABO blood group (transferase A, alpha 1-3-N-acetylgalactosaminyltransferase; transferase B, alpha 1-3-galactosyltransferase) |
| ABR | active BCR-related |
| ACADM | acyl-CoA dehydrogenase, C-4 to C-12 straight chain |
| ACTB | actin beta |
| ACVR1B | activin A receptor type 1B |
| ACVR2A | activin A receptor type 2A |
| ADAM10 | ADAM metallopeptidase domain 10 |
| ADARB2 | adenosine deaminase, RNA specific B2 (inactive) |
| ADCY1 | adenylate cyclase 1 |
| ADD3 | adducin 3 |
| ADORA2A | adenosine A2a receptor |
| AGER | advanced glycosylation end-product specific receptor |
| JAG1 | jagged 1 |
| APLNR | apelin receptor |
| AKT1 | AKT serine/threonine kinase 1 |
| AKT2 | AKT serine/threonine kinase 2 |
| ALAD | aminolevulinate dehydratase |
| ALDH2 | aldehyde dehydrogenase 2 family (mitochondrial) |
| ALDOB | aldolase, fructose-bisphosphate B |
| ALK | anaplastic lymphoma receptor tyrosine kinase |
| ALOX12 | arachidonate 12-lipoxygenase, 12S type |
| ALOX5 | arachidonate 5-lipoxygenase |
| AMFR | autocrine motility factor receptor |
| AMPH | amphiphysin |
| ANG | angiogenin |
| ANXA5 | annexin A5 |
| APEH | acylaminoacyl-peptide hydrolase |
| BIRC2 | baculoviral IAP repeat containing 2 |
| BIRC3 | baculoviral IAP repeat containing 3 |
| AR | androgen receptor |
| ARAF | A-Raf proto-oncogene, serine/threonine kinase |
| AREG | amphiregulin |
| ARF6 | ADP ribosylation factor 6 |
| RHOC | ras homolog family member C |
| RHOG | ras homolog family member G |
| ARHGAP4 | Rho GTPase activating protein 4 |
| RHOH | ras homolog family member H |
| ARNT | aryl hydrocarbon receptor nuclear translocator |

| | |
|---|---|
| ASL | argininosuccinate lyase |
| ASNS | asparagine synthetase (glutamine-hydrolyzing) |
| ATF1 | activating transcription factor 1 |
| ATF4 | activating transcription factor 4 |
| ATIC | 5-aminoimidazole-4-carboxamide ribonucleotide formyltransferase/IMP cyclohydrolase |
| RERE | arginine-glutamic acid dipeptide repeats |
| ATRX | ATRX, chromatin remodeler |
| AUH | AU RNA binding methylglutaconyl-CoA hydratase |
| AXL | AXL receptor tyrosine kinase |
| B2M | beta-2-microglobulin |
| BAD | BCL2 associated agonist of cell death |
| BAG1 | BCL2 associated athanogene 1 |
| BAK1 | BCL2 antagonist/killer 1 |
| CCND1 | cyclin D1 |
| BCL2 | BCL2, apoptosis regulator |
| BCL2A1 | BCL2 related protein A1 |
| BCL2L1 | BCL2 like 1 |
| BCL2L2 | BCL2 like 2 |
| BCL3 | B-cell CLL/lymphoma 3 |
| BCL5 | B-cell CLL/lymphoma 5 |
| BCL6 | B-cell CLL/lymphoma 6 |
| NBEAP1 | neurobeachin pseudogene 1 |
| BCL9 | B-cell CLL/lymphoma 9 |
| TNFRSF17 | TNF receptor superfamily member 17 |
| BCRP4 | breakpoint cluster region pseudogene 4 |
| BGN | biglycan |
| BID | BH3 interacting domain death agonist |
| BLK | BLK proto-oncogene, Src family tyrosine kinase |
| BMI1 | BMI1 proto-oncogene, polycomb ring finger |
| BNIP3 | BCL2 interacting protein 3 |
| BOK | BOK, BCL2 family apoptosis regulator |
| DST | dystonin |
| BRAF | B-Raf proto-oncogene, serine/threonine kinase |
| ZFP36L1 | ZFP36 ring finger protein like 1 |
| BST2 | bone marrow stromal cell antigen 2 |
| BTG1 | BTG anti-proliferation factor 1 |
| BUB1 | BUB1 mitotic checkpoint serine/threonine kinase |
| BUB1B | BUB1 mitotic checkpoint serine/threonine kinase B |
| C1QA | complement C1q A chain |
| C3AR1 | complement C3a receptor 1 |
| PTTG1IP | pituitary tumor-transforming 1 interacting protein |
| DDR1 | discoidin domain receptor tyrosine kinase 1 |
| CAD | carbamoyl-phosphate synthetase 2, aspartate transcarbamylase, and dihydroorotase |
| CALCA | calcitonin related polypeptide alpha |
| CALCR | calcitonin receptor |
| CALR | calreticulin |
| CAMK2B | calcium/calmodulin dependent protein kinase II beta |
| CAMK2D | calcium/calmodulin dependent protein kinase II delta |
| CAST | calpastatin |
| CARS | cysteinyl-tRNA synthetase |
| CASP1 | caspase 1 |
| CASP9 | caspase 9 |
| RUNX1T1 | RUNX1 translocation partner 1 |
| CBLB | Cbl proto-oncogene B |

| | |
|---|---|
| CCNA2 | cyclin A2 |
| CCNB1 | cyclin B1 |
| CCND2 | cyclin D2 |
| CCND3 | cyclin D3 |
| CCNE1 | cyclin E1 |
| CCNF | cyclin F |
| CCNG1 | cyclin G1 |
| CCNH | cyclin H |
| CCNT1 | cyclin T1 |
| CD6 | CD6 molecule |
| CD9 | CD9 molecule |
| TNFRSF8 | TNF receptor superfamily member 8 |
| SCARB1 | scavenger receptor class B member 1 |
| CD53 | CD53 molecule |
| CD68 | CD68 molecule |
| CD69 | CD69 molecule |
| CD74 | CD74 molecule |
| CD79A | CD79a molecule |
| CD79B | CD79b molecule |
| ADGRE5 | adhesion G protein-coupled receptor E5 |
| CD151 | CD151 molecule (Raph blood group) |
| CDA | cytidine deaminase |
| CDK1 | cyclin dependent kinase 1 |
| CDK11B | cyclin dependent kinase 11B |
| LRBA | LPS responsive beige-like anchor protein |
| CDC6 | cell division cycle 6 |
| CDC20 | cell division cycle 20 |
| CDC25A | cell division cycle 25A |
| CDC25B | cell division cycle 25B |
| CDC25C | cell division cycle 25C |
| CDC27 | cell division cycle 27 |
| CDC34 | cell division cycle 34 |
| CDC42 | cell division cycle 42 |
| CDH10 | cadherin 10 |
| CDK4 | cyclin dependent kinase 4 |
| CDK5 | cyclin dependent kinase 5 |
| CDK7 | cyclin dependent kinase 7 |
| CDKN2D | cyclin dependent kinase inhibitor 2D |
| CDKN3 | cyclin dependent kinase inhibitor 3 |
| CD52 | CD52 molecule |
| CEBPB | CCAAT/enhancer binding protein beta |
| CENPF | centromere protein F |
| CEACAM7 | carcinoembryonic antigen related cell adhesion molecule 7 |
| CHD2 | chromodomain helicase DNA binding protein 2 |
| CHD3 | chromodomain helicase DNA binding protein 3 |
| CHD4 | chromodomain helicase DNA binding protein 4 |
| CHGA | chromogranin A |
| CHKA | choline kinase alpha |
| CHN1 | chimerin 1 |
| CHRNB4 | cholinergic receptor nicotinic beta 4 subunit |
| CIRBP | cold inducible RNA binding protein |
| ERCC8 | ERCC excision repair 8, CSA ubiquitin ligase complex subunit |
| CKS1B | CDC28 protein kinase regulatory subunit 1B |
| CKS2 | CDC28 protein kinase regulatory subunit 2 |

| | |
|---|---|
| CLTC | clathrin heavy chain |
| CCR1 | C-C motif chemokine receptor 1 |
| CCR7 | C-C motif chemokine receptor 7 |
| CMM | cutaneous malignant melanoma/dysplastic nevus |
| CNR2 | cannabinoid receptor 2 |
| COL1A1 | collagen type I alpha 1 chain |
| COL1A2 | collagen type I alpha 2 chain |
| COL3A1 | collagen type III alpha 1 chain |
| COL4A3 | collagen type IV alpha 3 chain |
| COL7A1 | collagen type VII alpha 1 chain |
| COL11A1 | collagen type XI alpha 1 chain |
| COL16A1 | collagen type XVI alpha 1 chain |
| COL19A1 | collagen type XIX alpha 1 chain |
| COX6C | cytochrome c oxidase subunit 6C |
| CLDN4 | claudin 4 |
| CR1L | complement C3b/C4b receptor 1 like |
| CR2 | complement C3d receptor 2 |
| CREB1 | cAMP responsive element binding protein 1 |
| ATF2 | activating transcription factor 2 |
| CRIP2 | cysteine rich protein 2 |
| CRK | CRK proto-oncogene, adaptor protein |
| CRKL | CRK like proto-oncogene, adaptor protein |
| HAPLN1 | hyaluronan and proteoglycan link protein 1 |
| CRYAA | crystallin alpha A |
| CRYAB | crystallin alpha B |
| MAPK14 | mitogen-activated protein kinase 14 |
| CSE1L | chromosome segregation 1 like |
| CSF1 | colony stimulating factor 1 |
| CSF1R | colony stimulating factor 1 receptor |
| CSF3R | colony stimulating factor 3 receptor |
| CSNK2A1 | casein kinase 2 alpha 1 |
| VCAN | versican |
| CSPG4 | chondroitin sulfate proteoglycan 4 |
| CSTA | cystatin A |
| CSTB | cystatin B |
| NKX2-5 | NK2 homeobox 5 |
| CTNNA1 | catenin alpha 1 |
| CTNNB1 | catenin beta 1 |
| CTNND2 | catenin delta 2 |
| CTSB | cathepsin B |
| CTSH | cathepsin H |
| CTSL | cathepsin L |
| CYP1A1 | cytochrome P450 family 1 subfamily A member 1 |
| CYP2A6 | cytochrome P450 family 2 subfamily A member 6 |
| CYP2D6 | cytochrome P450 family 2 subfamily D member 6 |
| CYP7A1 | cytochrome P450 family 7 subfamily A member 1 |
| CYP19A1 | cytochrome P450 family 19 subfamily A member 1 |
| CD55 | CD55 molecule (Cromer blood group) |
| DAXX | death domain associated protein |
| DBN1 | drebrin 1 |
| DDIT3 | DNA damage inducible transcript 3 |
| DDX1 | DEAD/H-box helicase 1 |
| DDX5 | DEAD-box helicase 5 |
| DDX6 | DEAD-box helicase 6 |

| | |
|---|---|
| DHX9 | DEAH-box helicase 9 |
| DDX10 | DEAD-box helicase 10 |
| NQO1 | NAD(P)H quinone dehydrogenase 1 |
| DIO2 | deiodinase, iodothyronine type II |
| DKC1 | dyskerin pseudouridine synthase 1 |
| DLST | dihydrolipoamide S-succinyltransferase |
| DLX4 | distal-less homeobox 4 |
| DNAH9 | dynein axonemal heavy chain 9 |
| DNASE1L3 | deoxyribonuclease 1 like 3 |
| DOCK2 | dedicator of cytokinesis 2 |
| DPT | dermatopontin |
| DSG2 | desmoglein 2 |
| DUSP2 | dual specificity phosphatase 2 |
| DYRK1A | dual specificity tyrosine phosphorylation regulated kinase 1A |
| E2F6 | E2F transcription factor 6 |
| EBF1 | early B-cell factor 1 |
| EBVS1 | Epstein Barr virus integration site 1 |
| TYMP | thymidine phosphorylase |
| ECM1 | extracellular matrix protein 1 |
| LPAR1 | lysophosphatidic acid receptor 1 |
| EEF1A2 | eukaryotic translation elongation factor 1 alpha 2 |
| EGF | epidermal growth factor |
| EGFR | epidermal growth factor receptor |
| EGR3 | early growth response 3 |
| EIF1AX | eukaryotic translation initiation factor 1A, X-linked |
| EIF4A2 | eukaryotic translation initiation factor 4A2 |
| EIF4E | eukaryotic translation initiation factor 4E |
| EIF4EBP1 | eukaryotic translation initiation factor 4E binding protein 1 |
| EIF4G1 | eukaryotic translation initiation factor 4 gamma 1 |
| CELA1 | chymotrypsin like elastase family member 1 |
| SERPINB1 | serpin family B member 1 |
| ELAVL1 | ELAV like RNA binding protein 1 |
| ELF3 | E74 like ETS transcription factor 3 |
| ELF4 | E74 like ETS transcription factor 4 |
| ELK1 | ELK1, ETS transcription factor |
| ELK2AP | ELK2A, member of ETS oncogene family, pseudogene |
| ELK3 | ELK3, ETS transcription factor |
| ELK4 | ELK4, ETS transcription factor |
| ELN | elastin |
| CTTN | cortactin |
| ENO1 | enolase 1 |
| SLC29A1 | solute carrier family 29 member 1 (Augustine blood group) |
| EP300 | E1A binding protein p300 |
| EPHA7 | EPH receptor A7 |
| EPO | erythropoietin |
| EPOR | erythropoietin receptor |
| EPS8 | epidermal growth factor receptor pathway substrate 8 |
| EPS15 | epidermal growth factor receptor pathway substrate 15 |
| NR2F6 | nuclear receptor subfamily 2 group F member 6 |
| ERBB2 | erb-b2 receptor tyrosine kinase 2 |
| ERBB3 | erb-b2 receptor tyrosine kinase 3 |
| ERCC1 | ERCC excision repair 1, endonuclease non-catalytic subunit |
| ERCC2 | ERCC excision repair 2, TFIIH core complex helicase subunit |
| ERCC3 | ERCC excision repair 3, TFIIH core complex helicase subunit |

| | |
|---|---|
| ERCC4 | ERCC excision repair 4, endonuclease catalytic subunit |
| ERCC5 | ERCC excision repair 5, endonuclease |
| ERCC6 | ERCC excision repair 6, chromatin remodeling factor |
| ERG | ERG, ETS transcription factor |
| ESRRA | estrogen related receptor alpha |
| ESRRG | estrogen related receptor gamma |
| ETS1 | ETS proto-oncogene 1, transcription factor |
| ETV1 | ETS variant 1 |
| ETV3 | ETS variant 3 |
| ETV4 | ETS variant 4 |
| ETV5 | ETS variant 5 |
| MECOM | MDS1 and EVI1 complex locus |
| EWSR1 | EWS RNA binding protein 1 |
| FANCA | Fanconi anemia complementation group A |
| FANCC | Fanconi anemia complementation group C |
| FANCD2 | Fanconi anemia complementation group D2 |
| FANCE | Fanconi anemia complementation group E |
| ACSL3 | acyl-CoA synthetase long-chain family member 3 |
| FANCF | Fanconi anemia complementation group F |
| FAT2 | FAT atypical cadherin 2 |
| FBN2 | fibrillin 2 |
| FCGR2B | Fc fragment of IgG receptor IIb |
| FEN1 | flap structure-specific endonuclease 1 |
| FES | FES proto-oncogene, tyrosine kinase |
| FGF1 | fibroblast growth factor 1 |
| FGF2 | fibroblast growth factor 2 |
| FGF3 | fibroblast growth factor 3 |
| FGF4 | fibroblast growth factor 4 |
| FGF5 | fibroblast growth factor 5 |
| FGF6 | fibroblast growth factor 6 |
| FGF7 | fibroblast growth factor 7 |
| FGF8 | fibroblast growth factor 8 |
| FGF9 | fibroblast growth factor 9 |
| FGF10 | fibroblast growth factor 10 |
| FGF11 | fibroblast growth factor 11 |
| FGF12 | fibroblast growth factor 12 |
| FGF13 | fibroblast growth factor 13 |
| FGF14 | fibroblast growth factor 14 |
| FGFR1 | fibroblast growth factor receptor 1 |
| FGFR3 | fibroblast growth factor receptor 3 |
| FGFR2 | fibroblast growth factor receptor 2 |
| FGFR4 | fibroblast growth factor receptor 4 |
| FGR | FGR proto-oncogene, Src family tyrosine kinase |
| FHL2 | four and a half LIM domains 2 |
| FOXG1 | forkhead box G1 |
| FOXL1 | forkhead box L1 |
| FOXC2 | forkhead box C2 |
| FOXE1 | forkhead box E1 |
| FOXM1 | forkhead box M1 |
| FLI1 | Fli-1 proto-oncogene, ETS transcription factor |
| FLII | FLII, actin remodeling protein |
| FLNB | filamin B |
| FLT1 | fms related tyrosine kinase 1 |
| FMOD | fibromodulin |

| | |
|---|---|
| FN1 | fibronectin 1 |
| FOS | Fos proto-oncogene, AP-1 transcription factor subunit |
| FOSB | FosB proto-oncogene, AP-1 transcription factor subunit |
| FOSL2 | FOS like 2, AP-1 transcription factor subunit |
| FRA16A | fragile site, folic acid type, rare, fra(16)(p13.11) |
| MTOR | mechanistic target of rapamycin |
| FUT8 | fucosyltransferase 8 |
| KDSR | 3-ketodihydrosphingosine reductase |
| FYN | FYN proto-oncogene, Src family tyrosine kinase |
| XRCC6 | X-ray repair cross complementing 6 |
| GAB1 | GRB2 associated binding protein 1 |
| GABPB1 | GA binding protein transcription factor beta subunit 1 |
| GAK | cyclin G associated kinase |
| GALNS | galactosamine (N-acetyl)-6-sulfatase |
| GAPDH | glyceraldehyde-3-phosphate dehydrogenase |
| GAS6 | growth arrest specific 6 |
| GATA1 | GATA binding protein 1 |
| GATA2 | GATA binding protein 2 |
| GATA3 | GATA binding protein 3 |
| GATA6 | GATA binding protein 6 |
| GDF10 | growth differentiation factor 10 |
| GFI1 | growth factor independent 1 transcriptional repressor |
| GGTA1P | glycoprotein, alpha-galactosyltransferase 1 pseudogene |
| GHR | growth hormone receptor |
| GHRH | growth hormone releasing hormone |
| GIP | gastric inhibitory polypeptide |
| GLB1 | galactosidase beta 1 |
| GLI2 | GLI family zinc finger 2 |
| GLI3 | GLI family zinc finger 3 |
| GLI4 | GLI family zinc finger 4 |
| GMFB | glia maturation factor beta |
| GML | glycosylphosphatidylinositol anchored molecule like |
| GNA11 | G protein subunit alpha 11 |
| GNA12 | G protein subunit alpha 12 |
| GNAI1 | G protein subunit alpha i1 |
| GNAI2 | G protein subunit alpha i2 |
| GNAQ | G protein subunit alpha q |
| GNAS | GNAS complex locus |
| GOT2 | glutamic-oxaloacetic transaminase 2 |
| CXCR3 | C-X-C motif chemokine receptor 3 |
| GPER1 | G protein-coupled estrogen receptor 1 |
| GPS2 | G protein pathway suppressor 2 |
| GPX1 | glutathione peroxidase 1 |
| GRB2 | growth factor receptor bound protein 2 |
| GRB10 | growth factor receptor bound protein 10 |
| RAPGEF1 | Rap guanine nucleotide exchange factor 1 |
| GRN | granulin |
| GRIN2D | glutamate ionotropic receptor NMDA type subunit 2D |
| ARHGAP35 | Rho GTPase activating protein 35 |
| CXCL1 | C-X-C motif chemokine ligand 1 |
| CXCL2 | C-X-C motif chemokine ligand 2 |
| CXCL3 | C-X-C motif chemokine ligand 3 |
| GRP | gastrin releasing peptide |
| GRPR | gastrin releasing peptide receptor |

| | |
|---|---|
| GSPT1 | G1 to S phase transition 1 |
| GSTM1 | glutathione S-transferase mu 1 |
| MSH6 | mutS homolog 6 |
| GTF2A1 | general transcription factor IIA subunit 1 |
| GTF2H1 | general transcription factor IIH subunit 1 |
| GUCY1A2 | guanylate cyclase 1 soluble subunit alpha 2 |
| HIST1H1C | histone cluster 1 H1 family member c |
| HIST1H1B | histone cluster 1 H1 family member b |
| H3F3A | H3 histone family member 3A |
| HAS2 | hyaluronan synthase 2 |
| HCK | HCK proto-oncogene, Src family tyrosine kinase |
| HDLBP | high density lipoprotein binding protein |
| CFH | complement factor H |
| HFE | hemochromatosis |
| HGF | hepatocyte growth factor |
| HHEX | hematopoietically expressed homeobox |
| HIP1 | huntingtin interacting protein 1 |
| HLA-A | major histocompatibility complex, class I, A |
| HLA-B | major histocompatibility complex, class I, B |
| MNX1 | motor neuron and pancreas homeobox 1 |
| HLF | HLF, PAR bZIP transcription factor |
| HMGB1 | high mobility group box 1 |
| HMGCR | 3-hydroxy-3-methylglutaryl-CoA reductase |
| HMGA1 | high mobility group AT-hook 1 |
| HNRNPA2B1 | heterogeneous nuclear ribonucleoprotein A2/B1 |
| HNRNPD | heterogeneous nuclear ribonucleoprotein D |
| HNRNPH1 | heterogeneous nuclear ribonucleoprotein H1 (H) |
| HNRNPK | heterogeneous nuclear ribonucleoprotein K |
| TLX1 | T-cell leukemia homeobox 1 |
| HOXA3 | homeobox A3 |
| HOXA7 | homeobox A7 |
| HOXA9 | homeobox A9 |
| HOXA11 | homeobox A11 |
| HOXA13 | homeobox A13 |
| HOXB8 | homeobox B8 |
| HOXC11 | homeobox C11 |
| HOXC13 | homeobox C13 |
| HOXD11 | homeobox D11 |
| HOXD13 | homeobox D13 |
| HPR | haptoglobin-related protein |
| HPV18I2 | human papillomavirus (type 18) integration site 2 |
| HRAS | HRas proto-oncogene, GTPase |
| ERAS | ES cell expressed Ras |
| HES1 | hes family bHLH transcription factor 1 |
| HSPA5 | heat shock protein family A (Hsp70) member 5 |
| HSPA8 | heat shock protein family A (Hsp70) member 8 |
| HSPA9 | heat shock protein family A (Hsp70) member 9 |
| HSPB1 | heat shock protein family B (small) member 1 |
| HSP90AA1 | heat shock protein 90 alpha family class A member 1 |
| HSP90AA2P | heat shock protein 90 alpha family class A member 2, pseudogene |
| HSP90AB1 | heat shock protein 90 alpha family class B member 1 |
| HSPG2 | heparan sulfate proteoglycan 2 |
| HUS1 | HUS1 checkpoint clamp component |
| HVBS7 | hepatitis B virus integration site 7 |

| | |
|---|---|
| TNC | tenascin C |
| HYAL1 | hyaluronoglucosaminidase 1 |
| ICAM1 | intercellular adhesion molecule 1 |
| ID2 | inhibitor of DNA binding 2, HLH protein |
| ID3 | inhibitor of DNA binding 3, HLH protein |
| IDH2 | isocitrate dehydrogenase (NADP(+)) 2, mitochondrial |
| IFI27 | interferon alpha inducible protein 27 |
| SP110 | SP110 nuclear body protein |
| IFNAR1 | interferon alpha and beta receptor subunit 1 |
| IFNGR2 | interferon gamma receptor 2 (interferon gamma transducer 1) |
| IGF1R | insulin like growth factor 1 receptor |
| IGF2 | insulin like growth factor 2 |
| IGFBP1 | insulin like growth factor binding protein 1 |
| IGFBP2 | insulin like growth factor binding protein 2 |
| IGFBP6 | insulin like growth factor binding protein 6 |
| CYR61 | cysteine rich angiogenic inducer 61 |
| IGH | immunoglobulin heavy locus |
| IGHG1 | immunoglobulin heavy constant gamma 1 (G1m marker) |
| IGHG2 | immunoglobulin heavy constant gamma 2 (G2m marker) |
| IGHG3 | immunoglobulin heavy constant gamma 3 (G3m marker) |
| IGHG4 | immunoglobulin heavy constant gamma 4 (G4m marker) |
| IGHGP | immunoglobulin heavy constant gamma P (non-functional) |
| IGKC | immunoglobulin kappa constant |
| IGL | immunoglobulin lambda locus |
| IL1B | interleukin 1 beta |
| IL2 | interleukin 2 |
| IL2RA | interleukin 2 receptor subunit alpha |
| IL2RB | interleukin 2 receptor subunit beta |
| IL2RG | interleukin 2 receptor subunit gamma |
| IL3 | interleukin 3 |
| IL3RA | interleukin 3 receptor subunit alpha |
| IL3RA | interleukin 3 receptor subunit alpha |
| IL4R | interleukin 4 receptor |
| IL5 | interleukin 5 |
| IL6 | interleukin 6 |
| IL6ST | interleukin 6 signal transducer |
| IL7 | interleukin 7 |
| IL9 | interleukin 9 |
| IL16 | interleukin 16 |
| ILF3 | interleukin enhancer binding factor 3 |
| IDO1 | indoleamine 2,3-dioxygenase 1 |
| INHA | inhibin alpha subunit |
| INPPL1 | inositol polyphosphate phosphatase like 1 |
| EIF3E | eukaryotic translation initiation factor 3 subunit E |
| ITGA6 | integrin subunit alpha 6 |
| IRAK2 | interleukin 1 receptor associated kinase 2 |
| ITGA9 | integrin subunit alpha 9 |
| ITGAX | integrin subunit alpha X |
| ITGB4 | integrin subunit beta 4 |
| ITK | IL2 inducible T-cell kinase |
| JAG2 | jagged 2 |
| JAK1 | Janus kinase 1 |
| JAK2 | Janus kinase 2 |
| JAK3 | Janus kinase 3 |

| | |
|---|---|
| JUN | Jun proto-oncogene, AP-1 transcription factor subunit |
| JUNB | JunB proto-oncogene, AP-1 transcription factor subunit |
| JUND | JunD proto-oncogene, AP-1 transcription factor subunit |
| KCNH1 | potassium voltage-gated channel subfamily H member 1 |
| KDR | kinase insert domain receptor |
| KIFC3 | kinesin family member C3 |
| KIT | KIT proto-oncogene receptor tyrosine kinase |
| KLK2 | kallikrein related peptidase 2 |
| KLRB1 | killer cell lectin like receptor B1 |
| KPNA5 | karyopherin subunit alpha 5 |
| KRASP1 | KRAS proto-oncogene, GTPase pseudogene 1 |
| KRAS | KRAS proto-oncogene, GTPase |
| KTN1 | kinectin 1 |
| L1CAM | L1 cell adhesion molecule |
| AFF3 | AF4/FMR2 family member 3 |
| LASP1 | LIM and SH3 protein 1 |
| LCK | LCK proto-oncogene, Src family tyrosine kinase |
| LCP1 | lymphocyte cytosolic protein 1 |
| LEPR | leptin receptor |
| LFNG | LFNG O-fucosylpeptide 3-beta-N-acetylglucosaminyltransferase |
| LGALS3 | galectin 3 |
| LGALS3BP | galectin 3 binding protein |
| LHCGR | luteinizing hormone/choriogonadotropin receptor |
| LIMK1 | LIM domain kinase 1 |
| LMO1 | LIM domain only 1 |
| LMO2 | LIM domain only 2 |
| LMO7 | LIM domain 7 |
| LNPEP | leucyl and cystinyl aminopeptidase |
| LOXL2 | lysyl oxidase like 2 |
| LPP | LIM domain containing preferred translocation partner in lipoma |
| LRP2 | LDL receptor related protein 2 |
| LRP5 | LDL receptor related protein 5 |
| LTA | lymphotoxin alpha |
| LTB | lymphotoxin beta |
| LTBP3 | latent transforming growth factor beta binding protein 3 |
| LTBR | lymphotoxin beta receptor |
| LUM | lumican |
| LY6E | lymphocyte antigen 6 complex, locus E |
| LYL1 | LYL1, basic helix-loop-helix family member |
| LYN | LYN proto-oncogene, Src family tyrosine kinase |
| TACSTD2 | tumor-associated calcium signal transducer 2 |
| TM4SF1 | transmembrane 4 L six family member 1 |
| EPCAM | epithelial cell adhesion molecule |
| NBR1 | NBR1, autophagy cargo receptor |
| MXD1 | MAX dimerization protein 1 |
| MAD2L1 | MAD2 mitotic arrest deficient-like 1 (yeast) |
| SMAD3 | SMAD family member 3 |
| MAF | MAF bZIP transcription factor |
| MAFG | MAF bZIP transcription factor G |
| MAP2 | microtubule associated protein 2 |
| MAP4 | microtubule associated protein 4 |
| MARS | methionyl-tRNA synthetase |
| MAS1 | MAS1 proto-oncogene, G protein-coupled receptor |
| MATK | megakaryocyte-associated tyrosine kinase |

| | |
|---|---|
| MC2R | melanocortin 2 receptor |
| MCF2 | MCF.2 cell line derived transforming sequence |
| MCL1 | BCL2 family apoptosis regulator |
| MCM2 | minichromosome maintenance complex component 2 |
| MCM5 | minichromosome maintenance complex component 5 |
| CD46 | CD46 molecule |
| MDK | midkine (neurite growth-promoting factor 2) |
| MDM2 | MDM2 proto-oncogene |
| MDM4 | MDM4, p53 regulator |
| MEF2A | myocyte enhancer factor 2A |
| BORCS8-MEF2B | BORCS8-MEF2B readthrough |
| MEF2C | myocyte enhancer factor 2C |
| MEF2D | myocyte enhancer factor 2D |
| MEIS1 | Meis homeobox 1 |
| MAP3K1 | mitogen-activated protein kinase kinase kinase 1 |
| MAP3K3 | mitogen-activated protein kinase kinase kinase 3 |
| MAP3K5 | mitogen-activated protein kinase kinase kinase 5 |
| RAB8A | RAB8A, member RAS oncogene family |
| MET | MET proto-oncogene, receptor tyrosine kinase |
| KITLG | KIT ligand |
| CIITA | class II major histocompatibility complex transactivator |
| MID1 | midline 1 |
| MIF | macrophage migration inhibitory factor (glycosylation-inhibiting factor) |
| MITF | melanogenesis associated transcription factor |
| MLF1 | myeloid leukemia factor 1 |
| KMT2A | lysine methyltransferase 2A |
| MLLT1 | MLLT1, super elongation complex subunit |
| AFF1 | AF4/FMR2 family member 1 |
| MLLT3 | MLLT3, super elongation complex subunit |
| AFDN | afadin, adherens junction formation factor |
| MLLT6 | MLLT6, PHD finger domain containing |
| NR3C2 | nuclear receptor subfamily 3 group C member 2 |
| TRPM1 | transient receptor potential cation channel subfamily M member 1 |
| MMP2 | matrix metallopeptidase 2 |
| MMP9 | matrix metallopeptidase 9 |
| MMP11 | matrix metallopeptidase 11 |
| MMP14 | matrix metallopeptidase 14 |
| MN1 | MN1 proto-oncogene, transcriptional regulator |
| MNAT1 | MNAT1, CDK activating kinase assembly factor |
| MOS | v-mos Moloney murine sarcoma viral oncogene homolog |
| CD200 | CD200 molecule |
| MPL | MPL proto-oncogene, thrombopoietin receptor |
| MPP1 | membrane palmitoylated protein 1 |
| MRC1 | mannose receptor, C type 1 |
| MRE11A | MRE11 homolog A, double strand break repair nuclease |
| ABCC1 | ATP binding cassette subfamily C member 1 |
| MSH3 | mutS homolog 3 |
| MSN | moesin |
| MTCP1 | mature T-cell proliferation 1 |
| MTHFR | methylenetetrahydrofolate reductase |
| MUC1 | mucin 1, cell surface associated |
| MUC2 | mucin 2, oligomeric mucus/gel-forming |
| MUC4 | mucin 4, cell surface associated |
| MUC5AC | mucin 5AC, oligomeric mucus/gel-forming |

| | |
|---|---|
| MUC6 | mucin 6, oligomeric mucus/gel-forming |
| TRIM37 | tripartite motif containing 37 |
| MUTYH | mutY DNA glycosylase |
| MYB | MYB proto-oncogene, transcription factor |
| MYBL1 | MYB proto-oncogene like 1 |
| MYBL2 | MYB proto-oncogene like 2 |
| MYC | v-myc avian myelocytomatosis viral oncogene homolog |
| MYCL | v-myc avian myelocytomatosis viral oncogene lung carcinoma derived homolog |
| MYCLP1 | MYCL pseudogene 1 |
| MYCLK1 | v-myc avian myelocytomatosis viral oncogene homolog-like 1 |
| MYCN | v-myc avian myelocytomatosis viral oncogene neuroblastoma derived homolog |
| MYD88 | myeloid differentiation primary response 88 |
| MYH1 | myosin heavy chain 1 |
| MYH11 | myosin heavy chain 11 |
| MYOD1 | myogenic differentiation 1 |
| NAB2 | NGFI-A binding protein 2 |
| NACA | nascent polypeptide-associated complex alpha subunit |
| NBL1 | neuroblastoma 1, DAN family BMP antagonist |
| NCK1 | NCK adaptor protein 1 |
| NEB | nebulin |
| SEPT2 | septin 2 |
| NFATC1 | nuclear factor of activated T-cells 1 |
| NFATC3 | nuclear factor of activated T-cells 3 |
| NFE2L2 | nuclear factor, erythroid 2 like 2 |
| NFIB | nuclear factor I B |
| NFIC | nuclear factor I C |
| NFIX | nuclear factor I X |
| NFKB2 | nuclear factor kappa B subunit 2 |
| NFKBIA | NFKB inhibitor alpha |
| NFKBIE | NFKB inhibitor epsilon |
| NKX2-2 | NK2 homeobox 2 |
| NME3 | NME/NM23 nucleoside diphosphate kinase 3 |
| NMT1 | N-myristoyltransferase 1 |
| NNMT | nicotinamide N-methyltransferase |
| NONO | non-POU domain containing, octamer-binding |
| NPY | neuropeptide Y |
| NOTCH4 | notch 4 |
| NPY1R | neuropeptide Y receptor Y1 |
| NRAS | neuroblastoma RAS viral oncogene homolog |
| YBX1 | Y-box binding protein 1 |
| NTRK1 | neurotrophic receptor tyrosine kinase 1 |
| NTRK2 | neurotrophic receptor tyrosine kinase 2 |
| NUMA1 | nuclear mitotic apparatus protein 1 |
| NR4A2 | nuclear receptor subfamily 4 group A member 2 |
| OMD | osteomodulin |
| TNFRSF11B | TNF receptor superfamily member 11b |
| SLC22A18 | solute carrier family 22 member 18 |
| OTX2 | orthodenticle homeobox 2 |
| P2RX7 | purinergic receptor P2X 7 |
| PAFAH1B2 | platelet activating factor acetylhydrolase 1b catalytic subunit 2 |
| PRDX1 | peroxiredoxin 1 |
| PAK1 | p21 (RAC1) activated kinase 1 |
| PAM | peptidylglycine alpha-amidating monooxygenase |
| REG3A | regenerating family member 3 alpha |

| | |
|---|---|
| PAX2 | paired box 2 |
| PAX3 | paired box 3 |
| PAX7 | paired box 7 |
| PAX9 | paired box 9 |
| PBX1 | PBX homeobox 1 |
| PBX2 | PBX homeobox 2 |
| PCBP1 | poly(rC) binding protein 1 |
| PCM1 | pericentriolar material 1 |
| PCNA | proliferating cell nuclear antigen |
| CDK16 | cyclin dependent kinase 16 |
| CDK18 | cyclin dependent kinase 18 |
| PDE4A | phosphodiesterase 4A |
| PDE4D | phosphodiesterase 4D |
| PDGFA | platelet derived growth factor subunit A |
| PDGFB | platelet derived growth factor subunit B |
| PDGFRA | platelet derived growth factor receptor alpha |
| PDGFRB | platelet derived growth factor receptor beta |
| ENPP2 | ectonucleotide pyrophosphatase/phosphodiesterase 2 |
| PECAM1 | platelet and endothelial cell adhesion molecule 1 |
| SERPINF1 | serpin family F member 1 |
| PER1 | period circadian clock 1 |
| CFP | complement factor properdin |
| ATP8B1 | ATPase phospholipid transporting 8B1 |
| PFKFB3 | 6-phosphofructo-2-kinase/fructose-2,6-biphosphatase 3 |
| PGD | phosphogluconate dehydrogenase |
| PGF | placental growth factor |
| ABCB1 | ATP binding cassette subfamily B member 1 |
| ABCB4 | ATP binding cassette subfamily B member 4 |
| SERPINA1 | serpin family A member 1 |
| PIGR | polymeric immunoglobulin receptor |
| PIK3CA | phosphatidylinositol-4,5-bisphosphate 3-kinase catalytic subunit alpha |
| PIK3CB | phosphatidylinositol-4,5-bisphosphate 3-kinase catalytic subunit beta |
| PIM1 | Pim-1 proto-oncogene, serine/threonine kinase |
| PIK3CD | phosphatidylinositol-4,5-bisphosphate 3-kinase catalytic subunit delta |
| PIK3CG | phosphatidylinositol-4,5-bisphosphate 3-kinase catalytic subunit gamma |
| PIK3R1 | phosphoinositide-3-kinase regulatory subunit 1 |
| PIK3R2 | phosphoinositide-3-kinase regulatory subunit 2 |
| PITPNA | phosphatidylinositol transfer protein alpha |
| PITX1 | paired like homeodomain 1 |
| PKHD1 | polycystic kidney and hepatic disease 1 (autosomal recessive) |
| PKM | pyruvate kinase, muscle |
| PLA2G4A | phospholipase A2 group IVA |
| PLAG1 | PLAG1 zinc finger |
| PLAGL2 | PLAG1 like zinc finger 2 |
| PLAUR | plasminogen activator, urokinase receptor |
| PLCB2 | phospholipase C beta 2 |
| PLCL1 | phospholipase C like 1 |
| PLCG1 | phospholipase C gamma 1 |
| PLEC | plectin |
| PLXNB1 | plexin B1 |
| PMS1 | PMS1 homolog 1, mismatch repair system component |
| PMS2 | PMS1 homolog 2, mismatch repair system component |
| PRRX1 | paired related homeobox 1 |
| UBL3 | ubiquitin like 3 |

| | |
|---|---|
| SEPT5 | septin 5 |
| POLH | DNA polymerase eta |
| POU2AF1 | POU class 2 associating factor 1 |
| POU2F2 | POU class 2 homeobox 2 |
| POU4F1 | POU class 4 homeobox 1 |
| POU5F1 | POU class 5 homeobox 1 |
| PPARD | peroxisome proliferator activated receptor delta |
| PPP1CC | protein phosphatase 1 catalytic subunit gamma |
| PPP1R1A | protein phosphatase 1 regulatory inhibitor subunit 1A |
| PPP2R1A | protein phosphatase 2 scaffold subunit Aalpha |
| PRCC | papillary renal cell carcinoma (translocation-associated) |
| PRF1 | perforin 1 |
| PRKAB1 | protein kinase AMP-activated non-catalytic subunit beta 1 |
| PRKD1 | protein kinase D1 |
| MAPK1 | mitogen-activated protein kinase 1 |
| MAPK3 | mitogen-activated protein kinase 3 |
| MAPK4 | mitogen-activated protein kinase 4 |
| MAPK6 | mitogen-activated protein kinase 6 |
| MAPK7 | mitogen-activated protein kinase 7 |
| MAPK8 | mitogen-activated protein kinase 8 |
| MAPK13 | mitogen-activated protein kinase 13 |
| MAP2K1 | mitogen-activated protein kinase kinase 1 |
| MAP2K5 | mitogen-activated protein kinase kinase 5 |
| EIF2AK2 | eukaryotic translation initiation factor 2 alpha kinase 2 |
| PRLR | prolactin receptor |
| PROS1 | protein S (alpha) |
| PRPS1 | phosphoribosyl pyrophosphate synthetase 1 |
| KLK7 | kallikrein related peptidase 7 |
| PSAP | prosaposin |
| PSEN1 | presenilin 1 |
| PSEN2 | presenilin 2 |
| PSMA5 | proteasome subunit alpha 5 |
| PYY | peptide YY |
| PSMD2 | proteasome 26S subunit, non-ATPase 2 |
| PSPH | phosphoserine phosphatase |
| PTBP1 | polypyrimidine tract binding protein 1 |
| PTGIS | prostaglandin I2 synthase |
| PTH | parathyroid hormone |
| PTGS2 | prostaglandin-endoperoxide synthase 2 |
| PTHLH | parathyroid hormone like hormone |
| PTK7 | protein tyrosine kinase 7 (inactive) |
| PTMA | prothymosin, alpha |
| PTN | pleiotrophin |
| PTPN7 | protein tyrosine phosphatase, non-receptor type 7 |
| PTPN14 | protein tyrosine phosphatase, non-receptor type 14 |
| PTPRE | protein tyrosine phosphatase, receptor type E |
| PTPRG | protein tyrosine phosphatase, receptor type G |
| PTPRH | protein tyrosine phosphatase, receptor type H |
| PVT1 | Pvt1 oncogene (non-protein coding) |
| PXN | paxillin |
| RAB1A | RAB1A, member RAS oncogene family |
| RAB2A | RAB2A, member RAS oncogene family |
| RAB3A | RAB3A, member RAS oncogene family |
| RAB3B | RAB3B, member RAS oncogene family |

| | |
|---|---|
| RAB4A | RAB4A, member RAS oncogene family |
| RAB5A | RAB5A, member RAS oncogene family |
| RAB5B | RAB5B, member RAS oncogene family |
| RAB6A | RAB6A, member RAS oncogene family |
| MAP4K2 | mitogen-activated protein kinase kinase kinase kinase 2 |
| RAB13 | RAB13, member RAS oncogene family |
| RAB27A | RAB27A, member RAS oncogene family |
| RAB27B | RAB27B, member RAS oncogene family |
| RABGGTB | Rab geranylgeranyltransferase beta subunit |
| RAB5C | RAB5C, member RAS oncogene family |
| RAC1 | ras-related C3 botulinum toxin substrate 1 (rho family, small GTP binding protein Rac1) |
| RAC2 | ras-related C3 botulinum toxin substrate 2 (rho family, small GTP binding protein Rac2) |
| RAC3 | ras-related C3 botulinum toxin substrate 3 (rho family, small GTP binding protein Rac3) |
| RAD9A | RAD9 checkpoint clamp component A |
| RAD21 | RAD21 cohesin complex component |
| RAD23A | RAD23 homolog A, nucleotide excision repair protein |
| RAD51B | RAD51 paralog B |
| RAD51D | RAD51 paralog D |
| RAD52 | RAD52 homolog, DNA repair protein |
| RAF1 | Raf-1 proto-oncogene, serine/threonine kinase |
| RALA | RAS like proto-oncogene A |
| RALB | RAS like proto-oncogene B |
| RALGDS | ral guanine nucleotide dissociation stimulator |
| RAN | RAN, member RAS oncogene family |
| RANBP2 | RAN binding protein 2 |
| RAP1AP | RAP1A, member of RAS oncogene family pseudogene |
| RAP1B | RAP1B, member of RAS oncogene family |
| RAP1GDS1 | Rap1 GTPase-GDP dissociation stimulator 1 |
| RAP2A | RAP2A, member of RAS oncogene family |
| RAP2B | RAP2B, member of RAS oncogene family |
| RARA | retinoic acid receptor alpha |
| RARG | retinoic acid receptor gamma |
| RARRES1 | retinoic acid receptor responder 1 |
| RASA1 | RAS p21 protein activator 1 |
| RASGRF1 | Ras protein specific guanine nucleotide releasing factor 1 |
| RASGRF2 | Ras protein specific guanine nucleotide releasing factor 2 |
| ARID4A | AT-rich interaction domain 4A |
| RBBP4 | RB binding protein 4, chromatin remodeling factor |
| RBBP5 | RB binding protein 5, histone lysine methyltransferase complex subunit |
| RBBP6 | RB binding protein 6, ubiquitin ligase |
| RECQL | RecQ like helicase |
| REL | REL proto-oncogene, NF-kB subunit |
| RELA | RELA proto-oncogene, NF-kB subunit |
| RELB | RELB proto-oncogene, NF-kB subunit |
| REST | RE1 silencing transcription factor |
| RET | ret proto-oncogene |
| TRIM27 | tripartite motif containing 27 |
| RFX2 | regulatory factor X2 |
| RFX5 | regulatory factor X5 |
| RGS1 | regulator of G-protein signaling 1 |
| RGS2 | regulator of G-protein signaling 2 |
| RGS3 | regulator of G-protein signaling 3 |

| | |
|---|---|
| RHEB | Ras homolog enriched in brain |
| RLF | rearranged L-myc fusion |
| RLN2 | relaxin 2 |
| RMRP | RNA component of mitochondrial RNA processing endoribonuclease |
| BRD2 | bromodomain containing 2 |
| RORC | RAR related orphan receptor C |
| ROS1 | ROS proto-oncogene 1, receptor tyrosine kinase |
| RPGR | retinitis pigmentosa GTPase regulator |
| RPA2 | replication protein A2 |
| RPL22 | ribosomal protein L22 |
| RPL26 | ribosomal protein L26 |
| RPN1 | ribophorin I |
| RPS6KA1 | ribosomal protein S6 kinase A1 |
| RRAS | related RAS viral (r-ras) oncogene homolog |
| RREB1 | ras responsive element binding protein 1 |
| RRM1 | ribonucleotide reductase catalytic subunit M1 |
| CLIP1 | CAP-Gly domain containing linker protein 1 |
| RXRA | retinoid X receptor alpha |
| RYK | receptor-like tyrosine kinase |
| S100A7 | S100 calcium binding protein A7 |
| S100A8 | S100 calcium binding protein A8 |
| S100A9 | S100 calcium binding protein A9 |
| S100A10 | S100 calcium binding protein A10 |
| S100A13 | S100 calcium binding protein A13 |
| S100B | S100 calcium binding protein B |
| S100P | S100 calcium binding protein P |
| SAG | S-antigen visual arrestin |
| SAI1 | suppression of anchorage independence 1 |
| MAPK12 | mitogen-activated protein kinase 12 |
| SATB1 | SATB homeobox 1 |
| SERPINB3 | serpin family B member 3 |
| SERPINB4 | serpin family B member 4 |
| SCNN1B | sodium channel epithelial 1 beta subunit |
| CCL7 | C-C motif chemokine ligand 7 |
| SDC1 | syndecan 1 |
| SDC4 | syndecan 4 |
| SDHC | succinate dehydrogenase complex subunit C |
| SEA | S13 erythroblastosis (avian) oncogene homolog |
| SEL1L | SEL1L ERAD E3 ligase adaptor subunit |
| SELE | selectin E |
| SET | SET nuclear proto-oncogene |
| SFPQ | splicing factor proline and glutamine rich |
| SRSF2 | serine and arginine rich splicing factor 2 |
| SRSF3 | serine and arginine rich splicing factor 3 |
| SRSF6 | serine and arginine rich splicing factor 6 |
| SGK1 | serum/glucocorticoid regulated kinase 1 |
| SH3BP2 | SH3 domain binding protein 2 |
| SH3GL1 | SH3 domain containing GRB2 like 1, endophilin A2 |
| SH3GL2 | SH3 domain containing GRB2 like 2, endophilin A1 |
| SHBG | sex hormone binding globulin |
| SHC1 | SHC adaptor protein 1 |
| FBXW4 | F-box and WD repeat domain containing 4 |
| SHH | sonic hedgehog |
| SIAH2 | siah E3 ubiquitin protein ligase 2 |

| | |
|---|---|
| ST6GAL1 | ST6 beta-galactoside alpha-2,6-sialyltransferase 1 |
| ST3GAL1 | ST3 beta-galactoside alpha-2,3-sialyltransferase 1 |
| PMEL | premelanosome protein |
| STIL | SCL/TAL1 interrupting locus |
| SIPA1 | signal-induced proliferation-associated 1 |
| SIX1 | SIX homeobox 1 |
| SKI | SKI proto-oncogene |
| SKP1 | S-phase kinase associated protein 1 |
| SLC4A1 | solute carrier family 4 member 1 (Diego blood group) |
| SLC5A5 | solute carrier family 5 member 5 |
| SLC6A3 | solute carrier family 6 member 3 |
| SLC9A2 | solute carrier family 9 member A2 |
| SLC16A1 | solute carrier family 16 member 1 |
| SLC19A1 | solute carrier family 19 member 1 |
| SNAI2 | snail family transcriptional repressor 2 |
| SMO | smoothened, frizzled class receptor |
| SNAI1 | snail family transcriptional repressor 1 |
| SNAPC3 | small nuclear RNA activating complex polypeptide 3 |
| SNCG | synuclein gamma |
| FSCN1 | fascin actin-bundling protein 1 |
| SORL1 | sortilin related receptor 1 |
| SOS1 | SOS Ras/Rac guanine nucleotide exchange factor 1 |
| SOX2 | SRY-box 2 |
| SOX3 | SRY-box 3 |
| SOX4 | SRY-box 4 |
| SOX5 | SRY-box 5 |
| SOX9 | SRY-box 9 |
| SOX10 | SRY-box 10 |
| SPAM1 | sperm adhesion molecule 1 |
| SPG7 | SPG7, paraplegin matrix AAA peptidase subunit |
| SPINT1 | serine peptidase inhibitor, Kunitz type 1 |
| SPP1 | secreted phosphoprotein 1 |
| SPTAN1 | spectrin alpha, non-erythrocytic 1 |
| SRC | SRC proto-oncogene, non-receptor tyrosine kinase |
| SRI | sorcin |
| SRPK2 | SRSF protein kinase 2 |
| SSTR2 | somatostatin receptor 2 |
| SSX1 | SSX family member 1 |
| SSX2 | SSX family member 2 |
| SSX4 | SSX family member 4 |
| SS18 | SS18, nBAF chromatin remodeling complex subunit |
| ST3 | suppression of tumorigenicity 3 |
| ST8 | suppression of tumorigenicity 8 (ovarian) |
| STAT2 | signal transducer and activator of transcription 2 |
| STAT5B | signal transducer and activator of transcription 5B |
| STAT6 | signal transducer and activator of transcription 6 |
| STK4 | serine/threonine kinase 4 |
| AURKA | aurora kinase A |
| STRN | striatin |
| STX4 | syntaxin 4 |
| TAC1 | tachykinin precursor 1 |
| TACC1 | transforming acidic coiled-coil containing protein 1 |
| TAF1 | TATA-box binding protein associated factor 1 |
| MAP3K7 | mitogen-activated protein kinase kinase kinase 7 |

| | |
|---|---|
| TAL1 | TAL bHLH transcription factor 1, erythroid differentiation factor |
| TAL2 | TAL bHLH transcription factor 2 |
| TAP1 | transporter 1, ATP binding cassette subfamily B member |
| TAP2 | transporter 2, ATP binding cassette subfamily B member |
| TBX2 | T-box 2 |
| TCEA1 | transcription elongation factor A1 |
| TCEB1 | transcription elongation factor B subunit 1 |
| TBX3 | T-box 3 |
| HNF1A | HNF1 homeobox A |
| TCF7 | transcription factor 7 (T-cell specific, HMG-box) |
| TCF12 | transcription factor 12 |
| TRA | T-cell receptor alpha locus |
| TRB | T cell receptor beta locus |
| TRD | T cell receptor delta locus |
| TRG | T cell receptor gamma locus |
| TCTA | T-cell leukemia translocation altered |
| TDGF1P3 | teratocarcinoma-derived growth factor 1 pseudogene 3 |
| TECTA | tectorin alpha |
| TEK | TEK receptor tyrosine kinase |
| TERT | telomerase reverse transcriptase |
| TFAP4 | transcription factor AP-4 |
| NR2F1 | nuclear receptor subfamily 2 group F member 1 |
| TFDP1 | transcription factor Dp-1 |
| TFE3 | transcription factor binding to IGHM enhancer 3 |
| TFF1 | trefoil factor 1 |
| TFF2 | trefoil factor 2 |
| TFF3 | trefoil factor 3 |
| TFRC | transferrin receptor |
| TG | thyroglobulin |
| TGFA | transforming growth factor alpha |
| TGFB2 | transforming growth factor beta 2 |
| TGFB3 | transforming growth factor beta 3 |
| TGFBR1 | transforming growth factor beta receptor 1 |
| TGIF1 | TGFB induced factor homeobox 1 |
| THBS2 | thrombospondin 2 |
| THBS3 | thrombospondin 3 |
| THPO | thrombopoietin |
| THRSP | thyroid hormone responsive |
| TIAM1 | T-cell lymphoma invasion and metastasis 1 |
| TIE1 | tyrosine kinase with immunoglobulin like and EGF like domains 1 |
| TIMP1 | TIMP metallopeptidase inhibitor 1 |
| TIMP2 | TIMP metallopeptidase inhibitor 2 |
| NKX2-1 | NK2 homeobox 1 |
| TK1 | thymidine kinase 1 |
| ICAM5 | intercellular adhesion molecule 5 |
| TSPAN8 | tetraspanin 8 |
| TMPRSS2 | transmembrane protease, serine 2 |
| TNF | tumor necrosis factor |
| TNFAIP1 | TNF alpha induced protein 1 |
| TNFRSF1A | TNF receptor superfamily member 1A |
| TNFRSF1B | TNF receptor superfamily member 1B |
| TNR | tenascin R |
| TOP1 | topoisomerase (DNA) I |
| TOP2A | topoisomerase (DNA) II alpha |

| | |
|---|---|
| TOP2B | topoisomerase (DNA) II beta |
| TOP3A | topoisomerase (DNA) III alpha |
| TPD52 | tumor protein D52 |
| TPM3 | tropomyosin 3 |
| TPM4 | tropomyosin 4 |
| TPR | translocated promoter region, nuclear basket protein |
| TRAF1 | TNF receptor associated factor 1 |
| TRAF3 | TNF receptor associated factor 3 |
| TRHR | thyrotropin releasing hormone receptor |
| TRIO | trio Rho guanine nucleotide exchange factor |
| TRPS1 | transcriptional repressor GATA binding 1 |
| TSHR | thyroid stimulating hormone receptor |
| TWIST1 | twist family bHLH transcription factor 1 |
| TXN | thioredoxin |
| TYRO3 | TYRO3 protein tyrosine kinase |
| TYS | sclerotylosis |
| U2AF1 | U2 small nuclear RNA auxiliary factor 1 |
| UBA7 | ubiquitin like modifier activating enzyme 7 |
| UBE2D2 | ubiquitin conjugating enzyme E2 D2 |
| UBE3A | ubiquitin protein ligase E3A |
| USP4 | ubiquitin specific peptidase 4 |
| NR1H2 | nuclear receptor subfamily 1 group H member 2 |
| UQCRC2 | ubiquinol-cytochrome c reductase core protein II |
| USF1 | upstream transcription factor 1 |
| VAV1 | vav guanine nucleotide exchange factor 1 |
| VAV2 | vav guanine nucleotide exchange factor 2 |
| VCL | vinculin |
| VGF | VGF nerve growth factor inducible |
| VIS1 | viral integration site 1 |
| TRPV1 | transient receptor potential cation channel subfamily V member 1 |
| VRK1 | vaccinia related kinase 1 |
| WAS | Wiskott-Aldrich syndrome |
| CORO2A | coronin 2A |
| WHSC1 | Wolf-Hirschhorn syndrome candidate 1 |
| WNT1 | Wnt family member 1 |
| WNT2 | Wnt family member 2 |
| WNT3 | Wnt family member 3 |
| WNT6 | Wnt family member 6 |
| WNT10B | Wnt family member 10B |
| WRN | Werner syndrome RecQ like helicase |
| XDH | xanthine dehydrogenase |
| XPA | XPA, DNA damage recognition and repair factor |
| XPC | XPC complex subunit, DNA damage recognition and repair factor |
| XRCC3 | X-ray repair cross complementing 3 |
| YES1 | YES proto-oncogene 1, Src family tyrosine kinase |
| YWHAE | tyrosine 3-monooxygenase/tryptophan 5-monooxygenase activation protein epsilon |
| CNBP | CCHC-type zinc finger nucleic acid binding protein |
| ZNF35 | zinc finger protein 35 |
| ZNF91 | zinc finger protein 91 |
| MKRN3 | makorin ring finger protein 3 |
| ZNF132 | zinc finger protein 132 |
| ZNF146 | zinc finger protein 146 |
| ZMYM2 | zinc finger MYM-type containing 2 |
| ZNF217 | zinc finger protein 217 |

| | |
|---|---|
| LAPTM5 | lysosomal protein transmembrane 5 |
| CSDE1 | cold shock domain containing E1 |
| PCAP | predisposing for prostate cancer |
| TUBA1A | tubulin alpha 1a |
| PAX8 | paired box 8 |
| IFRD2 | interferon related developmental regulator 2 |
| MANF | mesencephalic astrocyte derived neurotrophic factor |
| USP7 | ubiquitin specific peptidase 7 |
| DEK | DEK proto-oncogene |
| TFEB | transcription factor EB |
| RNF217-AS1 | RNF217 antisense RNA 1 (head to head) |
| MAFK | MAF bZIP transcription factor K |
| KAT6A | lysine acetyltransferase 6A |
| PSCA | prostate stem cell antigen |
| BRD3 | bromodomain containing 3 |
| NUP214 | nucleoporin 214 |
| MLLT10 | myeloid/lymphoid or mixed-lineage leukemia; translocated to, 10 |
| CUBN | cubilin |
| CCDC6 | coiled-coil domain containing 6 |
| ADAM12 | ADAM metallopeptidase domain 12 |
| FOSL1 | FOS like 1, AP-1 transcription factor subunit |
| BRCATA | Breast cancer, 11;22 translocation associated |
| SSPN | sarcospan |
| KMT2D | lysine methyltransferase 2D |
| HMGA2 | high mobility group AT-hook 2 |
| GPR68 | G protein-coupled receptor 68 |
| TCL1A | T-cell leukemia/lymphoma 1A |
| TAF15 | TATA-box binding protein associated factor 15 |
| ELL | elongation factor for RNA polymerase II |
| ZNF239 | zinc finger protein 239 |
| NCOA3 | nuclear receptor coactivator 3 |
| NRIP1 | nuclear receptor interacting protein 1 |
| LZTR1 | leucine zipper like transcription regulator 1 |
| CLTCL1 | clathrin heavy chain like 1 |
| KDM5C | lysine demethylase 5C |
| SMC1A | structural maintenance of chromosomes 1A |
| NAA10 | N(alpha)-acetyltransferase 10, NatA catalytic subunit |
| HIST1H4I | histone cluster 1 H4 family member i |
| PICALM | phosphatidylinositol binding clathrin assembly protein |
| GFI1B | growth factor independent 1B transcriptional repressor |
| HIST1H2BG | histone cluster 1 H2B family member g |
| HIST1H3B | histone cluster 1 H3 family member b |
| PIP5K1A | phosphatidylinositol-4-phosphate 5-kinase type 1 alpha |
| BCAR3 | breast cancer anti-estrogen resistance 3 |
| BFSP2 | beaded filament structural protein 2 |
| CUL4B | cullin 4B |
| SORBS2 | sorbin and SH3 domain containing 2 |
| PPM1D | protein phosphatase, Mg2+/Mn2+ dependent 1D |
| PPFIBP1 | PPFIA binding protein 1 |
| RANBP3 | RAN binding protein 3 |
| SLC43A1 | solute carrier family 43 member 1 |
| ENC1 | ectodermal-neural cortex 1 |
| GAS7 | growth arrest specific 7 |
| DGKE | diacylglycerol kinase epsilon |

| | |
|---|---|
| CBX4 | chromobox 4 |
| DENR | density regulated re-initiation and release factor |
| THOC5 | THO complex 5 |
| PDXK | pyridoxal (pyridoxine, vitamin B6) kinase |
| RUVBL1 | RuvB like AAA ATPase 1 |
| PLPP3 | phospholipid phosphatase 3 |
| AKR1C3 | aldo-keto reductase family 1 member C3 |
| KCNK5 | potassium two pore domain channel subfamily K member 5 |
| NCOA1 | nuclear receptor coactivator 1 |
| IRS2 | insulin receptor substrate 2 |
| EIF3A | eukaryotic translation initiation factor 3 subunit A |
| EIF3C | eukaryotic translation initiation factor 3 subunit C |
| EIF3I | eukaryotic translation initiation factor 3 subunit I |
| VAMP8 | vesicle associated membrane protein 8 |
| HYAL2 | hyaluronoglucosaminidase 2 |
| S1PR4 | sphingosine-1-phosphate receptor 4 |
| RNGTT | RNA guanylyltransferase and 5'-phosphatase |
| TNFSF10 | tumor necrosis factor superfamily member 10 |
| ADAM23 | ADAM metallopeptidase domain 23 |
| ADAM9 | ADAM metallopeptidase domain 9 |
| TNFRSF14 | TNF receptor superfamily member 14 |
| RAB11A | RAB11A, member RAS oncogene family |
| TNFRSF6B | TNF receptor superfamily member 6b |
| NAPA | NSF attachment protein alpha |
| TNFRSF11A | TNF receptor superfamily member 11a |
| TNFRSF10C | TNF receptor superfamily member 10c |
| GMPS | guanine monophosphate synthase |
| SOCS2 | suppressor of cytokine signaling 2 |
| GGH | gamma-glutamyl hydrolase |
| CFLAR | CASP8 and FADD like apoptosis regulator |
| WISP2 | WNT1 inducible signaling pathway protein 2 |
| KSR1 | kinase suppressor of ras 1 |
| CDK5R1 | cyclin dependent kinase 5 regulatory subunit 1 |
| LDB1 | LIM domain binding 1 |
| SQSTM1 | sequestosome 1 |
| FUBP1 | far upstream element binding protein 1 |
| MTMR3 | myotubularin related protein 3 |
| PRPF4B | pre-mRNA processing factor 4B |
| PHOX2B | paired like homeobox 2b |
| MBD2 | methyl-CpG binding domain protein 2 |
| RAB29 | RAB29, member RAS oncogene family |
| BTRC | beta-transducin repeat containing E3 ubiquitin protein ligase |
| TNFSF18 | tumor necrosis factor superfamily member 18 |
| NOL3 | nucleolar protein 3 |
| KALRN | kalirin, RhoGEF kinase |
| MAP3K6 | mitogen-activated protein kinase kinase kinase 6 |
| USP6 | ubiquitin specific peptidase 6 |
| MTA1 | metastasis associated 1 |
| SLC16A3 | solute carrier family 16 member 3 |
| SMC3 | structural maintenance of chromosomes 3 |
| AIFM1 | apoptosis inducing factor, mitochondria associated 1 |
| RABEP1 | rabaptin, RAB GTPase binding effector protein 1 |
| RRP9 | ribosomal RNA processing 9, small subunit (SSU) processome component, homolog (yeast) |
| DYRK1B | dual specificity tyrosine phosphorylation regulated kinase 1B |

| | |
|---|---|
| PCSK7 | proprotein convertase subtilisin/kexin type 7 |
| EBAG9 | estrogen receptor binding site associated, antigen, 9 |
| TMSB10 | thymosin beta 10 |
| ARHGEF2 | Rho/Rac guanine nucleotide exchange factor 2 |
| REPS2 | RALBP1 associated Eps domain containing 2 |
| LRRFIP1 | LRR binding FLII interacting protein 1 |
| LGI1 | leucine rich glioma inactivated 1 |
| DLGAP1 | DLG associated protein 1 |
| RAB11B | RAB11B, member RAS oncogene family |
| PTTG1 | pituitary tumor-transforming 1 |
| PNMA1 | paraneoplastic Ma antigen 1 |
| GCNT3 | glucosaminyl (N-acetyl) transferase 3, mucin type |
| DHRS3 | dehydrogenase/reductase 3 |
| TSPOAP1 | TSPO associated protein 1 |
| MFHAS1 | malignant fibrous histiocytoma amplified sequence 1 |
| MAPKAPK2 | mitogen-activated protein kinase-activated protein kinase 2 |
| TRIP11 | thyroid hormone receptor interactor 11 |
| B4GALT5 | beta-1,4-galactosyltransferase 5 |
| RAB33A | RAB33A, member RAS oncogene family |
| RAB28 | RAB28, member RAS oncogene family |
| RAB9BP1 | RAB9B, member RAS oncogene family pseudogene 1 |
| RAB9A | RAB9A, member RAS oncogene family |
| NRXN2 | neurexin 2 |
| OTOF | otoferlin |
| RECQL5 | RecQ like helicase 5 |
| RECQL4 | RecQ like helicase 4 |
| TJP2 | tight junction protein 2 |
| CYP7B1 | cytochrome P450 family 7 subfamily B member 1 |
| MED17 | mediator complex subunit 17 |
| ARHGEF6 | Rac/Cdc42 guanine nucleotide exchange factor 6 |
| PICK1 | protein interacting with PRKCA 1 |
| MAPK8IP1 | mitogen-activated protein kinase 8 interacting protein 1 |
| ADAMTS1 | ADAM metallopeptidase with thrombospondin type 1 motif 1 |
| GDF15 | growth differentiation factor 15 |
| BAG3 | BCL2 associated athanogene 3 |
| RAB3D | RAB3D, member RAS oncogene family |
| BCAR1 | BCAR1, Cas family scaffolding protein |
| BRE | brain and reproductive organ-expressed (TNFRSF1A modulator) |
| RBM39 | RNA binding motif protein 39 |
| WTAP | Wilms tumor 1 associated protein |
| IER2 | immediate early response 2 |
| RAB36 | RAB36, member RAS oncogene family |
| NCOR1 | nuclear receptor corepressor 1 |
| NCOR2 | nuclear receptor corepressor 2 |
| RNF7 | ring finger protein 7 |
| TRAF4 | TNF receptor associated factor 4 |
| KLK4 | kallikrein related peptidase 4 |
| TCL1B | T-cell leukemia/lymphoma 1B |
| PLCH2 | phospholipase C eta 2 |
| PDE4DIP | phosphodiesterase 4D interacting protein |
| SDC3 | syndecan 3 |
| NUP93 | nucleoporin 93 |
| HERPUD1 | homocysteine inducible ER protein with ubiquitin like domain 1 |
| ARHGAP32 | Rho GTPase activating protein 32 |

| | |
|---|---|
| ACAP1 | ArfGAP with coiled-coil, ankyrin repeat and PH domains 1 |
| STARD8 | StAR related lipid transfer domain containing 8 |
| KMT2B | lysine methyltransferase 2B |
| HDAC4 | histone deacetylase 4 |
| BCLAF1 | BCL2 associated transcription factor 1 |
| ATG13 | autophagy related 13 |
| CTIF | cap binding complex dependent translation initiation factor |
| KEAP1 | kelch like ECH associated protein 1 |
| ARMCX2 | armadillo repeat containing, X-linked 2 |
| DNAJC6 | DnaJ heat shock protein family (Hsp40) member C6 |
| ZEB2 | zinc finger E-box binding homeobox 2 |
| ELMO1 | engulfment and cell motility 1 |
| GAB2 | GRB2 associated binding protein 2 |
| C2CD5 | C2 calcium dependent domain containing 5 |
| RHOBTB1 | Rho related BTB domain containing 1 |
| RABGAP1L | RAB GTPase activating protein 1 like |
| NCAPD2 | non-SMC condensin I complex subunit D2 |
| KIF14 | kinesin family member 14 |
| P2RY14 | purinergic receptor P2Y14 |
| MAFB | MAF bZIP transcription factor B |
| ARHGAP25 | Rho GTPase activating protein 25 |
| GOLGA5 | golgin A5 |
| USP15 | ubiquitin specific peptidase 15 |
| USP3 | ubiquitin specific peptidase 3 |
| MVP | major vault protein |
| TNFSF15 | tumor necrosis factor superfamily member 15 |
| THRAP3 | thyroid hormone receptor associated protein 3 |
| MED12 | mediator complex subunit 12 |
| MED13 | mediator complex subunit 13 |
| RBX1 | ring-box 1 |
| DOPEY2 | dopey family member 2 |
| ELK2BP | ELK2B, member of ETS oncogene family, pseudogene |
| AKT3 | AKT serine/threonine kinase 3 |
| ABI1 | abl interactor 1 |
| ZBTB33 | zinc finger and BTB domain containing 33 |
| HDAC6 | histone deacetylase 6 |
| PDCD6 | programmed cell death 6 |
| FRAT1 | frequently rearranged in advanced T-cell lymphomas 1 |
| HMGXB4 | HMG-box containing 4 |
| TOM1 | target of myb1 membrane trafficking protein |
| HUWE1 | HECT, UBA and WWE domain containing 1, E3 ubiquitin protein ligase |
| PTPRU | protein tyrosine phosphatase, receptor type U |
| TSPAN1 | tetraspanin 1 |
| RASGRP1 | RAS guanyl releasing protein 1 |
| DNAL4 | dynein axonemal light chain 4 |
| TRAP1 | TNF receptor associated protein 1 |
| AKAP9 | A-kinase anchoring protein 9 |
| WASF2 | WAS protein family member 2 |
| LHFP | lipoma HMGIC fusion partner |
| PATJ | PATJ, crumbs cell polarity complex component |
| OLIG2 | oligodendrocyte lineage transcription factor 2 |
| PRG4 | proteoglycan 4 |
| GPA33 | glycoprotein A33 |
| RASGRP2 | RAS guanyl releasing protein 2 |

| | |
|---|---|
| GPHN | gephyrin |
| RAMP1 | receptor activity modifying protein 1 |
| FSTL3 | follistatin like 3 |
| STAG1 | stromal antigen 1 |
| NET1 | neuroepithelial cell transforming 1 |
| RGS19 | regulator of G-protein signaling 19 |
| EIF1B | eukaryotic translation initiation factor 1B |
| APC2 | APC2, WNT signaling pathway regulator |
| TCIRG1 | T-cell immune regulator 1, ATPase H+ transporting V0 subunit a3 |
| TFG | TRK-fused gene |
| PKDREJ | polycystin (PKD) family receptor for egg jelly |
| CORO2B | coronin 2B |
| NOD1 | nucleotide binding oligomerization domain containing 1 |
| PIAS3 | protein inhibitor of activated STAT 3 |
| NSA2 | NSA2, ribosome biogenesis homolog |
| IFI30 | IFI30, lysosomal thiol reductase |
| VAV3 | vav guanine nucleotide exchange factor 3 |
| GPNMB | glycoprotein nmb |
| TACC3 | transforming acidic coiled-coil containing protein 3 |
| MERTK | MER proto-oncogene, tyrosine kinase |
| FST | follistatin |
| CARM1 | coactivator associated arginine methyltransferase 1 |
| NCOA2 | nuclear receptor coactivator 2 |
| SEMA4D | semaphorin 4D |
| SEMA4B | semaphorin 4B |
| BATF | basic leucine zipper ATF-like transcription factor |
| ANP32B | acidic nuclear phosphoprotein 32 family member B |
| SLC34A2 | solute carrier family 34 member 2 |
| PRPF8 | pre-mRNA processing factor 8 |
| SH2B2 | SH2B adaptor protein 2 |
| HEXIM1 | hexamethylene bisacetamide inducible 1 |
| IGF2BP1 | insulin like growth factor 2 mRNA binding protein 1 |
| CAMKK2 | calcium/calmodulin dependent protein kinase kinase 2 |
| TNFSF13B | tumor necrosis factor superfamily member 13b |
| USP39 | ubiquitin specific peptidase 39 |
| MGEA5 | meningioma expressed antigen 5 (hyaluronidase) |
| NFAT5 | nuclear factor of activated T-cells 5 |
| STAG2 | stromal antigen 2 |
| CHL1 | cell adhesion molecule L1 like |
| TOB2 | transducer of ERBB2, 2 |
| SEPT9 | septin 9 |
| CCR9 | C-C motif chemokine receptor 9 |
| SDCCAG8 | serologically defined colon cancer antigen 8 |
| HSPH1 | heat shock protein family H (Hsp110) member 1 |
| CPLX2 | complexin 2 |
| NEU3 | neuraminidase 3 |
| PPP1R13L | protein phosphatase 1 regulatory subunit 13 like |
| HPSE | heparanase |
| RUVBL2 | RuvB like AAA ATPase 2 |
| ARID5A | AT-rich interaction domain 5A |
| ME3 | malic enzyme 3 |
| RAB10 | RAB10, member RAS oncogene family |
| MALT1 | MALT1 paracaspase |
| APOBEC2 | apolipoprotein B mRNA editing enzyme catalytic subunit 2 |

| | |
|---|---|
| AP3M2 | adaptor related protein complex 3 mu 2 subunit |
| STARD3 | StAR related lipid transfer domain containing 3 |
| SERINC3 | serine incorporator 3 |
| MLLT11 | myeloid/lymphoid or mixed-lineage leukemia; translocated to, 11 |
| RAB40B | RAB40B, member RAS oncogene family |
| CLP1 | cleavage and polyadenylation factor I subunit 1 |
| RAB32 | RAB32, member RAS oncogene family |
| METAP2 | methionyl aminopeptidase 2 |
| KLK11 | kallikrein related peptidase 11 |
| RAB35 | RAB35, member RAS oncogene family |
| RAB31 | RAB31, member RAS oncogene family |
| PIM2 | Pim-2 proto-oncogene, serine/threonine kinase |
| ABHD2 | abhydrolase domain containing 2 |
| WWP1 | WW domain containing E3 ubiquitin protein ligase 1 |
| CNTRL | centriolin |
| UBE2C | ubiquitin conjugating enzyme E2 C |
| ADAM29 | ADAM metallopeptidase domain 29 |
| PTPN21 | protein tyrosine phosphatase, non-receptor type 21 |
| FGFR1OP | FGFR1 oncogene partner |
| CDC42EP1 | CDC42 effector protein 1 |
| PKIG | protein kinase (cAMP-dependent, catalytic) inhibitor gamma |
| GLMN | glomulin, FKBP associated protein |
| CORO1A | coronin 1A |
| PTP4A3 | protein tyrosine phosphatase type IVA, member 3 |
| RABL2B | RAB, member of RAS oncogene family-like 2B |
| RABL2A | RAB, member of RAS oncogene family-like 2A |
| NUDT6 | nudix hydrolase 6 |
| PSIP1 | PC4 and SFRS1 interacting protein 1 |
| ABCB8 | ATP binding cassette subfamily B member 8 |
| AKAP13 | A-kinase anchoring protein 13 |
| DUSP10 | dual specificity phosphatase 10 |
| GALNT5 | polypeptide N-acetylgalactosaminyltransferase 5 |
| RNF139 | ring finger protein 139 |
| PADI2 | peptidyl arginine deiminase 2 |
| PACSIN2 | protein kinase C and casein kinase substrate in neurons 2 |
| CASC3 | cancer susceptibility candidate 3 |
| RRAS2 | related RAS viral (r-ras) oncogene homolog 2 |
| ITGA11 | integrin subunit alpha 11 |
| MRAS | muscle RAS oncogene homolog |
| ATF5 | activating transcription factor 5 |
| RASA3 | RAS p21 protein activator 3 |
| PHLDA1 | pleckstrin homology like domain family A member 1 |
| PPM1E | protein phosphatase, Mg2+/Mn2+ dependent 1E |
| CARD8 | caspase recruitment domain family member 8 |
| KLRK1 | killer cell lectin like receptor K1 |
| MAPRE1 | microtubule associated protein RP/EB family member 1 |
| RAB18 | RAB18, member RAS oncogene family |
| DIP2C | disco interacting protein 2 homolog C |
| RAB21 | RAB21, member RAS oncogene family |
| STK38L | serine/threonine kinase 38 like |
| SPEN | spen family transcriptional repressor |
| RBM34 | RNA binding motif protein 34 |
| FNBP1 | formin binding protein 1 |
| NCOA6 | nuclear receptor coactivator 6 |

| | |
|---|---|
| UHRF1BP1L | UHRF1 binding protein 1 like |
| SWAP70 | SWAP switching B-cell complex 70kDa subunit |
| KDM4C | lysine demethylase 4C |
| ERC1 | ELKS/RAB6-interacting/CAST family member 1 |
| PEG10 | paternally expressed 10 |
| ZNF423 | zinc finger protein 423 |
| ARHGAP26 | Rho GTPase activating protein 26 |
| TAB2 | TGF-beta activated kinase 1/MAP3K7 binding protein 2 |
| N4BP3 | NEDD4 binding protein 3 |
| MAST2 | microtubule associated serine/threonine kinase 2 |
| SEPT6 | septin 6 |
| RGL1 | ral guanine nucleotide dissociation stimulator like 1 |
| FBXL7 | F-box and leucine rich repeat protein 7 |
| GSE1 | Gse1 coiled-coil protein |
| SYNE2 | spectrin repeat containing nuclear envelope protein 2 |
| DNAJC9 | DnaJ heat shock protein family (Hsp40) member C9 |
| PLCB1 | phospholipase C beta 1 |
| ASTN2 | astrotactin 2 |
| BOP1 | block of proliferation 1 |
| ANKRD12 | ankyrin repeat domain 12 |
| ADGRL2 | adhesion G protein-coupled receptor L2 |
| IQCE | IQ motif containing E |
| SMG6 | SMG6, nonsense mediated mRNA decay factor |
| ACSL6 | acyl-CoA synthetase long-chain family member 6 |
| SYNE1 | spectrin repeat containing nuclear envelope protein 1 |
| USP24 | ubiquitin specific peptidase 24 |
| FNBP4 | formin binding protein 4 |
| CRTC1 | CREB regulated transcription coactivator 1 |
| SIK3 | SIK family kinase 3 |
| COTL1 | coactosin like F-actin binding protein 1 |
| GPR161 | G protein-coupled receptor 161 |
| SF3B1 | splicing factor 3b subunit 1 |
| ABCB10 | ATP binding cassette subfamily B member 10 |
| HEY1 | hes related family bHLH transcription factor with YRPW motif 1 |
| BRD4 | bromodomain containing 4 |
| TNFRSF13B | TNF receptor superfamily member 13B |
| MACF1 | microtubule-actin crosslinking factor 1 |
| ZFYVE26 | zinc finger FYVE-type containing 26 |
| KCTD2 | potassium channel tetramerization domain containing 2 |
| KAT6B | lysine acetyltransferase 6B |
| ZNF281 | zinc finger protein 281 |
| PIK3R5 | phosphoinositide-3-kinase regulatory subunit 5 |
| CDK20 | cyclin dependent kinase 20 |
| PATZ1 | POZ/BTB and AT hook containing zinc finger 1 |
| CORO1C | coronin 1C |
| ZMYND8 | zinc finger MYND-type containing 8 |
| CBLC | Cbl proto-oncogene C |
| LDOC1 | leucine zipper down-regulated in cancer 1 |
| SSBP3 | single stranded DNA binding protein 3 |
| RAB38 | RAB38, member RAS oncogene family |
| NPAP1 | nuclear pore associated protein 1 |
| ZNF318 | zinc finger protein 318 |
| SLC39A6 | solute carrier family 39 member 6 |
| KLK5 | kallikrein related peptidase 5 |

| | |
|---|---|
| NIPBL | NIPBL, cohesin loading factor |
| RAB26 | RAB26, member RAS oncogene family |
| ABTB2 | ankyrin repeat and BTB domain containing 2 |
| ZNF521 | zinc finger protein 521 |
| CLIC4 | chloride intracellular channel 4 |
| SIN3A | SIN3 transcription regulator family member A |
| EGFL6 | EGF like domain multiple 6 |
| OSBPL3 | oxysterol binding protein like 3 |
| SS18L1 | SS18L1, nBAF chromatin remodeling complex subunit |
| SETBP1 | SET binding protein 1 |
| PPP1R16B | protein phosphatase 1 regulatory subunit 16B |
| ERC2 | ELKS/RAB6-interacting/CAST family member 2 |
| GGA1 | golgi associated, gamma adaptin ear containing, ARF binding protein 1 |
| EDRF1 | erythroid differentiation regulatory factor 1 |
| FGFR1OP2 | FGFR1 oncogene partner 2 |
| TTLL3 | tubulin tyrosine ligase like 3 |
| FBXO5 | F-box protein 5 |
| PPP1R14B | protein phosphatase 1 regulatory inhibitor subunit 14B |
| PLEK2 | pleckstrin 2 |
| CNNM4 | cyclin and CBS domain divalent metal cation transport mediator 4 |
| CHIC2 | cysteine rich hydrophobic domain 2 |
| AATF | apoptosis antagonizing transcription factor |
| MYEOV | myeloma overexpressed |
| STEAP1 | STEAP family member 1 |
| PABPC1 | poly(A) binding protein cytoplasmic 1 |
| TCL6 | T-cell leukemia/lymphoma 6 (non-protein coding) |
| TPK1 | thiamin pyrophosphokinase 1 |
| LYPD3 | LY6/PLAUR domain containing 3 |
| AFF4 | AF4/FMR2 family member 4 |
| SNX5 | sorting nexin 5 |
| CNTN6 | contactin 6 |
| RAB30 | RAB30, member RAS oncogene family |
| EML4 | echinoderm microtubule associated protein like 4 |
| CECR5 | cat eye syndrome chromosome region, candidate 5 |
| CDH20 | cadherin 20 |
| MACROD1 | MACRO domain containing 1 |
| CFAP20 | cilia and flagella associated protein 20 |
| CD274 | CD274 molecule |
| PARVB | parvin beta |
| TFPT | TCF3 fusion partner |
| TBX21 | T-box 21 |
| TLX3 | T-cell leukemia homeobox 3 |
| SOX8 | SRY-box 8 |
| ERVW-1 | endogenous retrovirus group W member 1 |
| STOML2 | stomatin like 2 |
| IL21R | interleukin 21 receptor |
| IL22 | interleukin 22 |
| ARHGEF4 | Rho guanine nucleotide exchange factor 4 |
| IGK | immunoglobulin kappa locus |
| CDON | cell adhesion associated, oncogene regulated |
| PDE11A | phosphodiesterase 11A |
| TBX22 | T-box 22 |
| BHD | Beukes familial hip dysplasia |
| ADIPOR1 | adiponectin receptor 1 |

| | |
|---|---|
| APH1A | aph-1 homolog A, gamma-secretase subunit |
| SBDS | SBDS ribosome assembly guanine nucleotide exchange factor |
| CYB5R4 | cytochrome b5 reductase 4 |
| LEF1 | lymphoid enhancer binding factor 1 |
| NIN | ninein |
| RAB9B | RAB9B, member RAS oncogene family |
| PHF20 | PHD finger protein 20 |
| KLF3 | Kruppel like factor 3 |
| ZDHHC3 | zinc finger DHHC-type containing 3 |
| ARMCX1 | armadillo repeat containing, X-linked 1 |
| UCHL5 | ubiquitin C-terminal hydrolase L5 |
| PRRX2 | paired related homeobox 2 |
| RHCG | Rh family C glycoprotein |
| EVL | Enah/Vasp-like |
| NELFCD | negative elongation factor complex member C/D |
| TRIAP1 | TP53 regulated inhibitor of apoptosis 1 |
| NCKIPSD | NCK interacting protein with SH3 domain |
| RAB14 | RAB14, member RAS oncogene family |
| RAB6B | RAB6B, member RAS oncogene family |
| IL23A | interleukin 23 subunit alpha |
| ARMCX3 | armadillo repeat containing, X-linked 3 |
| TRIM33 | tripartite motif containing 33 |
| ERGIC3 | ERGIC and golgi 3 |
| ACSL5 | acyl-CoA synthetase long-chain family member 5 |
| RAB23 | RAB23, member RAS oncogene family |
| CMPK1 | cytidine/uridine monophosphate kinase 1 |
| GHRL | ghrelin and obestatin prepropeptide |
| RTEL1 | regulator of telomere elongation helicase 1 |
| CDK12 | cyclin dependent kinase 12 |
| RAB8B | RAB8B, member RAS oncogene family |
| INPP5K | inositol polyphosphate-5-phosphatase K |
| RSF1 | remodeling and spacing factor 1 |
| SIX4 | SIX homeobox 4 |
| BCL11A | B-cell CLL/lymphoma 11A |
| SPA17 | sperm autoantigenic protein 17 |
| FXYD5 | FXYD domain containing ion transport regulator 5 |
| RAB4B | RAB4B, member RAS oncogene family |
| RAB24 | RAB24, member RAS oncogene family |
| SLC38A2 | solute carrier family 38 member 2 |
| SEMA5B | semaphorin 5B |
| ANLN | anillin actin binding protein |
| NLE1 | notchless homolog 1 |
| RNF216 | ring finger protein 216 |
| RHOF | ras homolog family member F, filopodia associated |
| SPATA6 | spermatogenesis associated 6 |
| PAF1 | PAF1 homolog, Paf1/RNA polymerase II complex component |
| RAB39A | RAB39A, member RAS oncogene family |
| FEV | FEV, ETS transcription factor |
| AHI1 | Abelson helper integration site 1 |
| DYM | dymeclin |
| BCAS3 | BCAS3, microtubule associated cell migration factor |
| BSPRY | B-box and SPRY domain containing |
| TMEM104 | transmembrane protein 104 |
| BCOR | BCL6 corepressor |

| | |
|---|---|
| RNF43 | ring finger protein 43 |
| CYP2W1 | cytochrome P450 family 2 subfamily W member 1 |
| SDHAF2 | succinate dehydrogenase complex assembly factor 2 |
| PHIP | pleckstrin homology domain interacting protein |
| RALGPS2 | Ral GEF with PH domain and SH3 binding motif 2 |
| WRAP53 | WD repeat containing antisense to TP53 |
| FAIM | Fas apoptotic inhibitory molecule |
| RNF220 | ring finger protein 220 |
| SBNO1 | strawberry notch homolog 1 |
| DRAM1 | DNA damage regulated autophagy modulator 1 |
| ZNF331 | zinc finger protein 331 |
| TRPV6 | transient receptor potential cation channel subfamily V member 6 |
| DDX43 | DEAD-box helicase 43 |
| IL17RB | interleukin 17 receptor B |
| FERMT1 | fermitin family member 1 |
| ZDHHC7 | zinc finger DHHC-type containing 7 |
| RAB20 | RAB20, member RAS oncogene family |
| BCAS4 | breast carcinoma amplified sequence 4 |
| NLRP2 | NLR family pyrin domain containing 2 |
| RABL6 | RAB, member RAS oncogene family-like 6 |
| VAC14 | Vac14, PIKFYVE complex component |
| ATF7IP | activating transcription factor 7 interacting protein |
| ZFP64 | ZFP64 zinc finger protein |
| ENAH | enabled homolog (Drosophila) |
| PARVA | parvin alpha |
| NUP133 | nucleoporin 133 |
| SCN3B | sodium voltage-gated channel beta subunit 3 |
| ST6GALNAC1 | ST6 N-acetylgalactosaminide alpha-2,6-sialyltransferase 1 |
| KIZ | kizuna centrosomal protein |
| LMO3 | LIM domain only 3 |
| ERBIN | erbb2 interacting protein |
| RCC2 | regulator of chromosome condensation 2 |
| FAM212B | family with sequence similarity 212 member B |
| SULF2 | sulfatase 2 |
| PCDHB15 | protocadherin beta 15 |
| KCNQ5 | potassium voltage-gated channel subfamily Q member 5 |
| TMPRSS4 | transmembrane protease, serine 4 |
| MUC13 | mucin 13, cell surface associated |
| KCMF1 | potassium channel modulatory factor 1 |
| SPPL2B | signal peptide peptidase like 2B |
| PMEPA1 | prostate transmembrane protein, androgen induced 1 |
| EMSY | EMSY, BRCA2 interacting transcriptional repressor |
| ACKR3 | atypical chemokine receptor 3 |
| RSRP1 | arginine and serine rich protein 1 |
| UTP3 | UTP3, small subunit processome component homolog (S. cerevisiae) |
| KNL1 | kinetochore scaffold 1 |
| GOPC | golgi associated PDZ and coiled-coil motif containing |
| CD248 | CD248 molecule |
| RHBG | Rh family B glycoprotein (gene/pseudogene) |
| ZMIZ1 | zinc finger MIZ-type containing 1 |
| ADAMTSL3 | ADAMTS like 3 |
| KIAA1147 | KIAA1147 |
| KAT14 | lysine acetyltransferase 14 |
| S100A14 | S100 calcium binding protein A14 |

| | |
|---|---|
| RAB22A | RAB22A, member RAS oncogene family |
| BIRC6 | baculoviral IAP repeat containing 6 |
| ARID1B | AT-rich interaction domain 1B |
| MTA3 | metastasis associated 1 family member 3 |
| AARS2 | alanyl-tRNA synthetase 2, mitochondrial |
| ZNF608 | zinc finger protein 608 |
| ARHGAP20 | Rho GTPase activating protein 20 |
| UNC79 | unc-79 homolog (C. elegans) |
| PREX1 | phosphatidylinositol-3,4,5-trisphosphate dependent Rac exchange factor 1 |
| MKL1 | megakaryoblastic leukemia (translocation) 1 |
| EP400 | E1A binding protein p400 |
| KIAA1524 | KIAA1524 |
| KIAA1549 | KIAA1549 |
| RNF213 | ring finger protein 213 |
| CHD8 | chromodomain helicase DNA binding protein 8 |
| MAGEE1 | MAGE family member E1 |
| MIER1 | MIER1 transcriptional regulator |
| SFMBT2 | Scm-like with four mbt domains 2 |
| METTL14 | methyltransferase like 14 |
| EPG5 | ectopic P-granules autophagy protein 5 homolog |
| MARK4 | microtubule affinity regulating kinase 4 |
| RAB40C | RAB40C, member RAS oncogene family |
| CCNB1IP1 | cyclin B1 interacting protein 1 |
| RAP2C | RAP2C, member of RAS oncogene family |
| PLEKHB1 | pleckstrin homology domain containing B1 |
| NLRC4 | NLR family CARD domain containing 4 |
| SCAF1 | SR-related CTD associated factor 1 |
| EPS15L1 | epidermal growth factor receptor pathway substrate 15 like 1 |
| IL22RA1 | interleukin 22 receptor subunit alpha 1 |
| PLEKHA2 | pleckstrin homology domain containing A2 |
| LGR6 | leucine rich repeat containing G protein-coupled receptor 6 |
| AVPI1 | arginine vasopressin induced 1 |
| EXOC4 | exocyst complex component 4 |
| BACH2 | BTB domain and CNC homolog 2 |
| ELOVL5 | ELOVL fatty acid elongase 5 |
| SAV1 | salvador family WW domain containing protein 1 |
| ELAC2 | elaC ribonuclease Z 2 |
| PAPPA2 | pappalysin 2 |
| SMAP1 | small ArfGAP 1 |
| SLC22A23 | solute carrier family 22 member 23 |
| THADA | THADA, armadillo repeat containing |
| TNN | tenascin N |
| PRDM16 | PR/SET domain 16 |
| PBLD | phenazine biosynthesis like protein domain containing |
| CRLF2 | cytokine receptor-like factor 2 |
| CRLF2 | cytokine receptor-like factor 2 |
| RAB17 | RAB17, member RAS oncogene family |
| SOX17 | SRY-box 17 |
| NSD1 | nuclear receptor binding SET domain protein 1 |
| ARHGAP9 | Rho GTPase activating protein 9 |
| TMPRSS3 | transmembrane protease, serine 3 |
| SMYD3 | SET and MYND domain containing 3 |
| CREB3L2 | cAMP responsive element binding protein 3 like 2 |
| RMND5B | required for meiotic nuclear division 5 homolog B |

| | |
|---|---|
| RBM15 | RNA binding motif protein 15 |
| CRTC3 | CREB regulated transcription coactivator 3 |
| LMF1 | lipase maturation factor 1 |
| PLEKHG2 | pleckstrin homology and RhoGEF domain containing G2 |
| RANBP17 | RAN binding protein 17 |
| BCL11B | B-cell CLL/lymphoma 11B |
| SLC30A5 | solute carrier family 30 member 5 |
| RAPH1 | Ras association (RalGDS/AF-6) and pleckstrin homology domains 1 |
| MARCKSL1 | MARCKS like 1 |
| MPPE1 | metallophosphoesterase 1 |
| RASL11B | RAS like family 11 member B |
| ASPSCR1 | ASPSCR1, UBX domain containing tether for SLC2A4 |
| WDR77 | WD repeat domain 77 |
| CHCHD7 | coiled-coil-helix-coiled-coil-helix domain containing 7 |
| RHBDF2 | rhomboid 5 homolog 2 |
| NEIL1 | nei like DNA glycosylase 1 |
| HSPBAP1 | HSPB1 associated protein 1 |
| VEPH1 | ventricular zone expressed PH domain containing 1 |
| VTCN1 | V-set domain containing T cell activation inhibitor 1 |
| ZDHHC14 | zinc finger DHHC-type containing 14 |
| TBL1XR1 | transducin beta like 1 X-linked receptor 1 |
| EHMT1 | euchromatic histone lysine methyltransferase 1 |
| FAM57A | family with sequence similarity 57 member A |
| BAALC | brain and acute leukemia, cytoplasmic |
| CBLL1 | Cbl proto-oncogene like 1 |
| ZNF442 | zinc finger protein 442 |
| PGGHG | protein-glucosylgalactosylhydroxylysine glucosidase |
| FBXO11 | F-box protein 11 |
| CHD9 | chromodomain helicase DNA binding protein 9 |
| ORAI2 | ORAI calcium release-activated calcium modulator 2 |
| TET1 | tet methylcytosine dioxygenase 1 |
| WDR82 | WD repeat domain 82 |
| PDCD1LG2 | programmed cell death 1 ligand 2 |
| SLC44A4 | solute carrier family 44 member 4 |
| COL18A1 | collagen type XVIII alpha 1 chain |
| SLC38A1 | solute carrier family 38 member 1 |
| FAM49A | family with sequence similarity 49 member A |
| FAM117A | family with sequence similarity 117 member A |
| NECTIN4 | nectin cell adhesion molecule 4 |
| FIP1L1 | factor interacting with PAPOLA and CPSF1 |
| CDT1 | chromatin licensing and DNA replication factor 1 |
| TRIM7 | tripartite motif containing 7 |
| RAB1B | RAB1B, member RAS oncogene family |
| FCRL5 | Fc receptor like 5 |
| FCRL4 | Fc receptor like 4 |
| RAB33B | RAB33B, member RAS oncogene family |
| BCL2L12 | BCL2 like 12 |
| SH3BGRL2 | SH3 domain binding glutamate rich protein like 2 |
| INHBE | inhibin beta E subunit |
| FRMD8 | FERM domain containing 8 |
| RAB34 | RAB34, member RAS oncogene family |
| MIXL1 | Mix paired-like homeobox |
| BRIP1 | BRCA1 interacting protein C-terminal helicase 1 |
| REG4 | regenerating family member 4 |

| | |
|---|---|
| OBSCN | obscurin, cytoskeletal calmodulin and titin-interacting RhoGEF |
| FSCB | fibrous sheath CABYR binding protein |
| RAB6C | RAB6C, member RAS oncogene family |
| ARID5B | AT-rich interaction domain 5B |
| KIAA1109 | KIAA1109 |
| LOXL4 | lysyl oxidase like 4 |
| TRAF7 | TNF receptor associated factor 7 |
| ZDHHC18 | zinc finger DHHC-type containing 18 |
| SARNP | SAP domain containing ribonucleoprotein |
| HOOK3 | hook microtubule tethering protein 3 |
| CARD11 | caspase recruitment domain family member 11 |
| MAML2 | mastermind like transcriptional coactivator 2 |
| DOT1L | DOT1 like histone lysine methyltransferase |
| ZC3H8 | zinc finger CCCH-type containing 8 |
| USP38 | ubiquitin specific peptidase 38 |
| USP32 | ubiquitin specific peptidase 32 |
| KDM2B | lysine demethylase 2B |
| ACCS | 1-aminocyclopropane-1-carboxylate synthase homolog (inactive) |
| LOXL3 | lysyl oxidase like 3 |
| PLEKHA8 | pleckstrin homology domain containing A8 |
| MINA | MYC induced nuclear antigen |
| RSPO3 | R-spondin 3 |
| AIFM2 | apoptosis inducing factor, mitochondria associated 2 |
| DIRC2 | disrupted in renal carcinoma 2 |
| RAB2B | RAB2B, member RAS oncogene family |
| AJUBA | ajuba LIM protein |
| UBL7 | ubiquitin like 7 |
| SYDE1 | synapse defective Rho GTPase homolog 1 |
| SLC45A3 | solute carrier family 45 member 3 |
| RSPH1 | radial spoke head 1 homolog |
| WNT3A | Wnt family member 3A |
| NAV3 | neuron navigator 3 |
| NAV1 | neuron navigator 1 |
| COX19 | COX19, cytochrome c oxidase assembly factor |
| FMNL3 | formin like 3 |
| PCED1B | PC-esterase domain containing 1B |
| ANKRD44 | ankyrin repeat domain 44 |
| BOC | BOC cell adhesion associated, oncogene regulated |
| ZNF300 | zinc finger protein 300 |
| SNX29 | sorting nexin 29 |
| SPECC1 | sperm antigen with calponin homology and coiled-coil domains 1 |
| NEURL3 | neuralized E3 ubiquitin protein ligase 3 |
| IGSF8 | immunoglobulin superfamily member 8 |
| TRMT10A | tRNA methyltransferase 10A |
| CADPS2 | calcium dependent secretion activator 2 |
| C12orf9 | chromosome 12 open reading frame 9 |
| MUC16 | mucin 16, cell surface associated |
| FNIP1 | folliculin interacting protein 1 |
| CMTM7 | CKLF like MARVEL transmembrane domain containing 7 |
| TP53RK | TP53 regulating kinase |
| AHNAK2 | AHNAK nucleoprotein 2 |
| SCAMP4 | secretory carrier membrane protein 4 |
| DTX2 | deltex E3 ubiquitin ligase 2 |
| LACTB | lactamase beta |

| | |
|---|---|
| NLRP3 | NLR family pyrin domain containing 3 |
| CSMD3 | CUB and Sushi multiple domains 3 |
| RNF157 | ring finger protein 157 |
| SLAMF6 | SLAM family member 6 |
| MARCH3 | membrane associated ring-CH-type finger 3 |
| RAB42 | RAB42, member RAS oncogene family |
| RAB3C | RAB3C, member RAS oncogene family |
| CTHRC1 | collagen triple helix repeat containing 1 |
| RMI2 | RecQ mediated genome instability 2 |
| DIRC1 | disrupted in renal carcinoma 1 |
| RAB39B | RAB39B, member RAS oncogene family |
| MAS1L | MAS1 proto-oncogene like, G protein-coupled receptor |
| ARAP2 | ArfGAP with RhoGAP domain, ankyrin repeat and PH domain 2 |
| AGAP2 | ArfGAP with GTPase domain, ankyrin repeat and PH domain 2 |
| DACH2 | dachshund family transcription factor 2 |
| DNAJC24 | DnaJ heat shock protein family (Hsp40) member C24 |
| LRRK2 | leucine rich repeat kinase 2 |
| SPPL3 | signal peptide peptidase like 3 |
| MSI2 | musashi RNA binding protein 2 |
| CANT1 | calcium activated nucleotidase 1 |
| NLRP8 | NLR family pyrin domain containing 8 |
| TNFAIP8L1 | TNF alpha induced protein 8 like 1 |
| LRRC25 | leucine rich repeat containing 25 |
| M1AP | meiosis 1 associated protein |
| SCLT1 | sodium channel and clathrin linker 1 |
| RAB5CP2 | RAB5C, member RAS oncogene family pseudogene 2 |
| CD109 | CD109 molecule |
| LRGUK | leucine rich repeats and guanylate kinase domain containing |
| CKS1BP7 | CDC28 protein kinase regulatory subunit 1B pseudogene 7 |
| MUC17 | mucin 17, cell surface associated |
| MIB2 | mindbomb E3 ubiquitin protein ligase 2 |
| RAB40A | RAB40A, member RAS oncogene family |
| VTI1A | vesicle transport through interaction with t-SNAREs 1A |
| TMEM86A | transmembrane protein 86A |
| PTGR2 | prostaglandin reductase 2 |
| ZNF569 | zinc finger protein 569 |
| BTBD19 | BTB domain containing 19 |
| C1orf64 | chromosome 1 open reading frame 64 |
| TTL | tubulin tyrosine ligase |
| PUS10 | pseudouridylate synthase 10 |
| METTL21A | methyltransferase like 21A |
| XXYLT1 | xyloside xylosyltransferase 1 |
| CNTN4 | contactin 4 |
| JAKMIP1 | janus kinase and microtubule interacting protein 1 |
| NKAIN2 | Na+/K+ transporting ATPase interacting 2 |
| AMOT | angiomotin |
| ERICH1 | glutamate rich 1 |
| TMEM74 | transmembrane protein 74 |
| MAMDC4 | MAM domain containing 4 |
| TTC39B | tetratricopeptide repeat domain 39B |
| PPTC7 | PTC7 protein phosphatase homolog |
| SYNE3 | spectrin repeat containing nuclear envelope family member 3 |
| CITED4 | Cbp/p300 interacting transactivator with Glu/Asp rich carboxy-terminal domain 4 |
| SDE2 | SDE2 telomere maintenance homolog |

| | |
|---|---|
| XIRP1 | xin actin binding repeat containing 1 |
| RASSF6 | Ras association domain family member 6 |
| GIMAP7 | GTPase, IMAP family member 7 |
| IDO2 | indoleamine 2,3-dioxygenase 2 |
| SSBP4 | single stranded DNA binding protein 4 |
| ADAMTS15 | ADAM metallopeptidase with thrombospondin type 1 motif 15 |
| ZNF384 | zinc finger protein 384 |
| PTCRA | pre T-cell antigen receptor alpha |
| B3GNT6 | UDP-GlcNAc:betaGal beta-1,3-N-acetylglucosaminyltransferase 6 |
| DTX3 | deltex E3 ubiquitin ligase 3 |
| ALG14 | ALG14, UDP-N-acetylglucosaminyltransferase subunit |
| CRTC2 | CREB regulated transcription coactivator 2 |
| RAB12 | RAB12, member RAS oncogene family |
| SUSD3 | sushi domain containing 3 |
| ORAOV1 | oral cancer overexpressed 1 |
| HNRNPA3 | heterogeneous nuclear ribonucleoprotein A3 |
| AK9 | adenylate kinase 9 |
| RANP1 | RAN, member RAS oncogene family pseudogene 1 |
| JAZF1 | JAZF zinc finger 1 |
| BRAT1 | BRCA1 associated ATM activator 1 |
| BRWD3 | bromodomain and WD repeat domain containing 3 |
| NUTM1 | NUT midline carcinoma family member 1 |
| TAB3 | TGF-beta activated kinase 1/MAP3K7 binding protein 3 |
| ALS2CL | ALS2 C-terminal like |
| ASPM | abnormal spindle microtubule assembly |
| MDS2 | myelodysplastic syndrome 2 translocation associated |
| STEAP2 | STEAP2 metalloreductase |
| RAB40AL | RAB40A, member RAS oncogene family-like |
| HMGN2P46 | high mobility group nucleosomal binding domain 2 pseudogene 46 |
| GSDMA | gasdermin A |
| SLC26A11 | solute carrier family 26 member 11 |
| ACTL9 | actin like 9 |
| ZNF844 | zinc finger protein 844 |
| HKR1 | HKR1, GLI-Kruppel zinc finger family member |
| RSPO1 | R-spondin 1 |
| FRG1BP | FSHD region gene 1 family member B, pseudogene |
| SLC9A9 | solute carrier family 9 member A9 |
| RABL3 | RAB, member of RAS oncogene family like 3 |
| FRYL | FRY like transcription coactivator |
| CAGE1 | cancer antigen 1 |
| CRB2 | crumbs 2, cell polarity complex component |
| P2RY8 | purinergic receptor P2Y8 |
| P2RY8 | purinergic receptor P2Y8 |
| KLHL10 | kelch like family member 10 |
| KRT73 | keratin 73 |
| RAB37 | RAB37, member RAS oncogene family |
| RAB7B | RAB7B, member RAS oncogene family |
| C1QTNF9 | C1q and tumor necrosis factor related protein 9 |
| RNASE10 | ribonuclease A family member 10 (inactive) |
| RAB43 | RAB43, member RAS oncogene family |
| ENPP7 | ectonucleotide pyrophosphatase/phosphodiesterase 7 |
| COL28A1 | collagen type XXVIII alpha 1 chain |
| ABCB5 | ATP binding cassette subfamily B member 5 |
| RSPO2 | R-spondin 2 |

| | |
|---|---|
| ZSCAN22 | zinc finger and SCAN domain containing 22 |
| RAB41 | RAB41, member RAS oncogene family |
| GEN1 | GEN1, Holliday junction 5' flap endonuclease |
| KRT26 | keratin 26 |
| IRF2BP2 | interferon regulatory factor 2 binding protein 2 |
| PEAR1 | platelet endothelial aggregation receptor 1 |
| RAB15 | RAB15, member RAS oncogene family |
| SLC27A1 | solute carrier family 27 member 1 |
| MALAT1 | metastasis associated lung adenocarcinoma transcript 1 (non-protein coding) |
| RPSAP58 | ribosomal protein SA pseudogene 58 |
| MAFA | MAF bZIP transcription factor A |
| SHC4 | SHC adaptor protein 4 |
| BCAR4 | breast cancer anti-estrogen resistance 4 (non-protein coding) |
| BCRP2 | breakpoint cluster region pseudogene 2 |
| RAB44 | RAB44, member RAS oncogene family |
| RAB19 | RAB19, member RAS oncogene family |
| LOC402641 | v-ral simian leukemia viral oncogene homolog A (ras related) pseudogene |
| CUEDC1 | CUE domain containing 1 |
| MIR10B | microRNA 10b |
| MIR21 | microRNA 21 |
| MIR221 | microRNA 221 |
| IDNK | IDNK, gluconokinase |
| PIM3 | Pim-3 proto-oncogene, serine/threonine kinase |
| LOC440059 | cell adhesion associated, oncogene regulated pseudogene |
| H3F3C | H3 histone family member 3C |
| RAB1C | RAB1C, member RAS oncogene family pseudogene |
| LOC441768 | RAB31, member RAS oncogene family pseudogene |
| LOC642550 | v-ral simian leukemia viral oncogene homolog A (ras related) pseudogene |
| RAP1BL | RAP1B, member of RAS oncogene family pseudogene |
| LOC643916 | RAB1A, member RAS oncogene family pseudogene |
| ARAFP2 | ARAF pseudogene 2 |
| BCRP3 | breakpoint cluster region pseudogene 3 |
| RAB42P1 | RAB42, member RAS oncogene family, pseudogene 1 |
| ELK1P1 | ELK1, member of ETS oncogene family pseudogene 1 |
| KIAA0895L | KIAA0895 like |
| TLCD2 | TLC domain containing 2 |
| POTEF | POTE ankyrin domain family member F |
| SKP1P2 | S-phase kinase associated protein 1 pseudogene 2 |
| CDK11A | cyclin dependent kinase 11A |
| NUTM2B | NUT family member 2B |
| DIRC3 | disrupted in renal carcinoma 3 |
| ZNF814 | zinc finger protein 814 |
| BMI1P1 | BMI1 proto-oncogene, polycomb ring finger pseudogene 1 |
| LOC100128122 | Cbl proto-oncogene like 1 pseudogene |
| RAP1BP2 | RAP1B, member of RAS oncogene family pseudogene 2 |
| RANP6 | RAN, member RAS oncogene family pseudogene 6 |
| RAB28P5 | RAB28, member RAS oncogene family pseudogene 5 |
| LOC100128800 | cell adhesion associated, oncogene regulated pseudogene |
| RAB28P2 | RAB28, member RAS oncogene family pseudogene 2 |
| RANP5 | RAN, member RAS oncogene family pseudogene 5 |
| LOC100129672 | cell adhesion associated, oncogene regulated pseudogene |
| RAB11AP1 | RAB11A, member RAS oncogene family pseudogene 1 |
| LOC100130203 | cell adhesion associated, oncogene regulated pseudogene |
| RANP4 | RAN, member RAS oncogene family pseudogene 4 |

| | |
|---|---|
| LOC100130841 | MDM2 oncogene, E3 ubiquitin protein ligase pseudogene |
| LOC100131294 | RAB13, member RAS oncogene family pseudogene |
| LOC100131617 | v-ets avian erythroblastosis virus E26 oncogene homolog 2 pseudogene |
| NBPF10 | neuroblastoma breakpoint family member 10 |
| RAB28P3 | RAB28, member RAS oncogene family pseudogene 3 |
| RAB28P4 | RAB28, member RAS oncogene family pseudogene 4 |
| RAP1BP3 | RAP1B, member of RAS oncogene family pseudogene 3 |
| RAB28P1 | RAB28, member RAS oncogene family pseudogene 1 |
| LOC100133211 | related RAS viral (r-ras) oncogene homolog 2 pseudogene |
| MEF2B | myocyte enhancer factor 2B |
| CMC4 | C-X9-C motif containing 4 |
| RAB9AP1 | RAB9A, member RAS oncogene family pseudogene 1 |
| RAB9AP5 | RAB9A, member RAS oncogene family pseudogene 5 |
| RAB9AP2 | RAB9A, member RAS oncogene family pseudogene 2 |
| RAB9AP3 | RAB9A, member RAS oncogene family pseudogene 3 |
| RAB9AP4 | RAB9A, member RAS oncogene family pseudogene 4 |
| DUX4 | double homeobox 4 |
| LOC100335030 | FGFR1 oncogene partner 2 pseudogene |
| LOC100384885 | related RAS viral (r-ras) oncogene homolog 2 pseudogene |
| LOC100418497 | MAS1 proto-oncogene like, G protein-coupled receptor pseudogene |
| LOC100418498 | MAS1 proto-oncogene like, G protein-coupled receptor pseudogene |
| LOC100418499 | MAS1 proto-oncogene like, G protein-coupled receptor pseudogene |
| LOC100418500 | MAS1 proto-oncogene like, G protein-coupled receptor pseudogene |
| LOC100418501 | MAS1 proto-oncogene like, G protein-coupled receptor pseudogene |
| LOC100418502 | MAS1 proto-oncogene like, G protein-coupled receptor pseudogene |
| LOC100418503 | MAS1 proto-oncogene like, G protein-coupled receptor pseudogene |
| LOC100418504 | MAS1 proto-oncogene like, G protein-coupled receptor pseudogene |
| LOC100418505 | MAS1 proto-oncogene like, G protein-coupled receptor pseudogene |
| LOC100418506 | MAS1 proto-oncogene like, G protein-coupled receptor pseudogene |
| LOC100418507 | MAS1 proto-oncogene like, G protein-coupled receptor pseudogene |
| LOC100418508 | MAS1 proto-oncogene like, G protein-coupled receptor pseudogene |
| LOC100418509 | MAS1 proto-oncogene like, G protein-coupled receptor pseudogene |
| LOC100418510 | MAS1 proto-oncogene like, G protein-coupled receptor pseudogene |
| LOC100418578 | RAN, member RAS oncogene family pseudogene |
| LOC100418579 | RAN, member RAS oncogene family pseudogene |
| LOC100418580 | RAN, member RAS oncogene family pseudogene |
| LOC100418581 | RAN, member RAS oncogene family pseudogene |
| LOC100418582 | RAN, member RAS oncogene family pseudogene |
| LOC100419932 | FGFR1 oncogene partner 2 pseudogene |
| RANP8 | RAN, member RAS oncogene family pseudogene 8 |
| LOC100420571 | RAB1A, member RAS oncogene family pseudogene |
| LOC100420673 | RAB5A, member RAS oncogene family pseudogene |
| RAB5CP1 | RAB5C, member RAS oncogene family pseudogene 1 |
| LOC100421737 | ubiquitin specific peptidase 4 (proto-oncogene) pseudogene |
| LOC100422456 | v-rel avian reticuloendotheliosis viral oncogene homolog A pseudogene |
| RAP1BP1 | RAP1B, member of RAS oncogene family pseudogene 1 |
| SEPT2P1 | septin 2 pseudogene 1 |
| FGFR1OP2P1 | FGFR1 oncogene partner 2 pseudogene 1 |
| RANP2 | RAN, member RAS oncogene family pseudogene 2 |
| RANP7 | RAN, member RAS oncogene family pseudogene 7 |
| RANP9 | RAN, member RAS oncogene family pseudogene 9 |
| RANP3 | RAN, member RAS oncogene family pseudogene 3 |
| LOC101060112 | RAP1A, member of RAS oncogene family pseudogene |
| BLACAT1 | bladder cancer associated transcript 1 (non-protein coding) |

| | |
|---|---|
| RAB11AP2 | RAB11A, member RAS oncogene family pseudogene 2 |

Table S2: TS list (Known)

| symbol | info |
|---|---|
| ACHE | acetylcholinesterase (Cartwright blood group) |
| ACY1 | aminoacylase 1 |
| ADARB1 | adenosine deaminase, RNA specific B1 |
| ADPRH | ADP-ribosylarginine hydrolase |
| PARP1 | poly(ADP-ribose) polymerase 1 |
| AGTR1 | angiotensin II receptor type 1 |
| AHR | aryl hydrocarbon receptor |
| AIF1 | allograft inflammatory factor 1 |
| AKR1B1 | aldo-keto reductase family 1 member B |
| ALOX15 | arachidonate 15-lipoxygenase |
| ALOX15B | arachidonate 15-lipoxygenase, type B |
| ALPL | alkaline phosphatase, liver/bone/kidney |
| AMH | anti-Mullerian hormone |
| BIN1 | bridging integrator 1 |
| ANXA1 | annexin A1 |
| ANXA7 | annexin A7 |
| APAF1 | apoptotic peptidase activating factor 1 |
| APC | APC, WNT signaling pathway regulator |
| FAS | Fas cell surface death receptor |
| ARF1 | ADP ribosylation factor 1 |
| ARG1 | arginase 1 |
| RHOA | ras homolog family member A |
| RHOB | ras homolog family member B |
| RND3 | Rho family GTPase 3 |
| PHOX2A | paired like homeobox 2a |
| ARNTL | aryl hydrocarbon receptor nuclear translocator like |
| ASCL1 | achaete-scute family bHLH transcription factor 1 |
| ASS1 | argininosuccinate synthase 1 |
| ZFHX3 | zinc finger homeobox 3 |
| ATF3 | activating transcription factor 3 |
| ATM | ATM serine/threonine kinase |
| ATR | ATR serine/threonine kinase |
| AZGP1 | alpha-2-glycoprotein 1, zinc-binding |
| BARD1 | BRCA1 associated RING domain 1 |
| BAX | BCL2 associated X, apoptosis regulator |
| BCL7A | BCL tumor suppressor 7A |
| BCR | BCR, RhoGEF and GTPase activating protein |
| CEACAM1 | carcinoembryonic antigen related cell adhesion molecule 1 |
| BIK | BCL2 interacting killer |
| PRDM1 | PR/SET domain 1 |
| BLM | Bloom syndrome RecQ like helicase |
| BMP2 | bone morphogenetic protein 2 |
| BMP4 | bone morphogenetic protein 4 |
| BMPR1A | bone morphogenetic protein receptor type 1A |
| BMPR2 | bone morphogenetic protein receptor type 2 |
| BNIP3L | BCL2 interacting protein 3 like |
| FOXL2 | forkhead box L2 |
| BRCA1 | BRCA1, DNA repair associated |
| BRCA2 | BRCA2, DNA repair associated |

| | |
|---|---|
| ZFP36L2 | ZFP36 ring finger protein like 2 |
| KLF5 | Kruppel like factor 5 |
| BTK | Bruton tyrosine kinase |
| CAPG | capping actin protein, gelsolin like |
| CASP2 | caspase 2 |
| CASP5 | caspase 5 |
| CASP8 | caspase 8 |
| CAT | catalase |
| CAV1 | caveolin 1 |
| RUNX2 | runt related transcription factor 2 |
| RUNX1 | runt related transcription factor 1 |
| CBFA2T3 | CBFA2/RUNX1 translocation partner 3 |
| RUNX3 | runt related transcription factor 3 |
| CBFB | core-binding factor beta subunit |
| CBL | Cbl proto-oncogene |
| KRIT1 | KRIT1, ankyrin repeat containing |
| CCNC | cyclin C |
| CD4 | CD4 molecule |
| CD44 | CD44 molecule (Indian blood group) |
| CDH1 | cadherin 1 |
| CDH4 | cadherin 4 |
| CDH5 | cadherin 5 |
| CDH11 | cadherin 11 |
| CDH13 | cadherin 13 |
| CDH17 | cadherin 17 |
| CDK2 | cyclin dependent kinase 2 |
| CDK6 | cyclin dependent kinase 6 |
| CDKN1A | cyclin dependent kinase inhibitor 1A |
| CDKN1B | cyclin dependent kinase inhibitor 1B |
| CDKN1C | cyclin dependent kinase inhibitor 1C |
| CDKN2A | cyclin dependent kinase inhibitor 2A |
| CDKN2B | cyclin dependent kinase inhibitor 2B |
| CDKN2C | cyclin dependent kinase inhibitor 2C |
| CDO1 | cysteine dioxygenase type 1 |
| CDX2 | caudal type homeobox 2 |
| CEBPA | CCAAT/enhancer binding protein alpha |
| CEBPD | CCAAT/enhancer binding protein delta |
| CFTR | cystic fibrosis transmembrane conductance regulator |
| CHD1 | chromodomain helicase DNA binding protein 1 |
| CHEK1 | checkpoint kinase 1 |
| CHUK | conserved helix-loop-helix ubiquitous kinase |
| CLU | clusterin |
| CNN1 | calponin 1 |
| KLF6 | Kruppel like factor 6 |
| MAP3K8 | mitogen-activated protein kinase kinase kinase 8 |
| CREBBP | CREB binding protein |
| CREM | cAMP responsive element modulator |
| CSF2 | colony stimulating factor 2 |
| CSNK1A1 | casein kinase 1 alpha 1 |
| CST5 | cystatin D |
| CST6 | cystatin E/M |
| CTGF | connective tissue growth factor |
| CTNNA2 | catenin alpha 2 |
| CTNND1 | catenin delta 1 |

| | |
|---|---|
| CUX1 | cut like homeobox 1 |
| CYB5A | cytochrome b5 type A |
| CYLD | CYLD lysine 63 deubiquitinase |
| DAB2 | DAB2, clathrin adaptor protein |
| DACH1 | dachshund family transcription factor 1 |
| DAPK1 | death associated protein kinase 1 |
| DAPK3 | death associated protein kinase 3 |
| BRINP1 | BMP/retinoic acid inducible neural specific 1 |
| DCC | DCC netrin 1 receptor |
| DCN | decorin |
| DDB2 | damage specific DNA binding protein 2 |
| GADD45A | growth arrest and DNA damage inducible alpha |
| DDX3X | DEAD-box helicase 3, X-linked |
| DEFB1 | defensin beta 1 |
| DFFA | DNA fragmentation factor subunit alpha |
| DFNA5 | DFNA5, deafness associated tumor suppressor |
| DLG1 | discs large MAGUK scaffold protein 1 |
| DMBT1 | deleted in malignant brain tumors 1 |
| DMD | dystrophin |
| DNMT1 | DNA methyltransferase 1 |
| DNMT3A | DNA methyltransferase 3 alpha |
| DNMT3B | DNA methyltransferase 3 beta |
| DOK1 | docking protein 1 |
| DPH1 | diphthamide biosynthesis 1 |
| DPP4 | dipeptidyl peptidase 4 |
| DSC3 | desmocollin 3 |
| DSP | desmoplakin |
| DUSP1 | dual specificity phosphatase 1 |
| DUSP5 | dual specificity phosphatase 5 |
| DUSP6 | dual specificity phosphatase 6 |
| DUSP9 | dual specificity phosphatase 9 |
| E2F1 | E2F transcription factor 1 |
| E2F2 | E2F transcription factor 2 |
| E2F3 | E2F transcription factor 3 |
| E2F4 | E2F transcription factor 4 |
| ECT2 | epithelial cell transforming 2 |
| EDNRB | endothelin receptor type B |
| EEF1A1 | eukaryotic translation elongation factor 1 alpha 1 |
| EFNA5 | ephrin A5 |
| EGR1 | early growth response 1 |
| EGR2 | early growth response 2 |
| EPHA2 | EPH receptor A2 |
| EMP1 | epithelial membrane protein 1 |
| EMP2 | epithelial membrane protein 2 |
| EPAS1 | endothelial PAS domain protein 1 |
| EPB41 | erythrocyte membrane protein band 4.1 |
| EPHA1 | EPH receptor A1 |
| EPHA3 | EPH receptor A3 |
| EPHB2 | EPH receptor B2 |
| EPHB3 | EPH receptor B3 |
| EPHB4 | EPH receptor B4 |
| EPHB6 | EPH receptor B6 |
| ERBB4 | erb-b2 receptor tyrosine kinase 4 |
| EYA4 | EYA transcriptional coactivator and phosphatase 4 |

| | |
|---|---|
| ERF | ETS2 repressor factor |
| ESR1 | estrogen receptor 1 |
| ESR2 | estrogen receptor 2 |
| ESRRB | estrogen related receptor beta |
| ETS2 | ETS proto-oncogene 2, transcription factor |
| ETV6 | ETS variant 6 |
| EXT1 | exostosin glycosyltransferase 1 |
| EXT2 | exostosin glycosyltransferase 2 |
| EXTL1 | exostosin like glycosyltransferase 1 |
| EXTL2 | exostosin like glycosyltransferase 2 |
| EXTL3 | exostosin like glycosyltransferase 3 |
| EZH1 | enhancer of zeste 1 polycomb repressive complex 2 subunit |
| EZH2 | enhancer of zeste 2 polycomb repressive complex 2 subunit |
| FABP3 | fatty acid binding protein 3 |
| FANCG | Fanconi anemia complementation group G |
| FBLN1 | fibulin 1 |
| FAT1 | FAT atypical cadherin 1 |
| FBP1 | fructose-bisphosphatase 1 |
| GPC5 | glypican 5 |
| FH | fumarate hydratase |
| FHIT | fragile histidine triad |
| FHL1 | four and a half LIM domains 1 |
| FOXC1 | forkhead box C1 |
| FOXO1 | forkhead box O1 |
| FOXO3 | forkhead box O3 |
| FLNA | filamin A |
| FLT3 | fms related tyrosine kinase 3 |
| FXN | frataxin |
| FRK | fyn related Src family tyrosine kinase |
| FUS | FUS RNA binding protein |
| GALR1 | galanin receptor 1 |
| GAS1 | growth arrest specific 1 |
| GATA4 | GATA binding protein 4 |
| GBP1 | guanylate binding protein 1 |
| GJA1 | gap junction protein alpha 1 |
| GJB2 | gap junction protein beta 2 |
| GPC3 | glypican 3 |
| GLI1 | GLI family zinc finger 1 |
| GNAT1 | G protein subunit alpha transducin 1 |
| SFN | stratifin |
| GPX3 | glutathione peroxidase 3 |
| GRIN2A | glutamate ionotropic receptor NMDA type subunit 2A |
| GSK3B | glycogen synthase kinase 3 beta |
| GSN | gelsolin |
| GSTP1 | glutathione S-transferase pi 1 |
| GSTT1 | glutathione S-transferase theta 1 |
| BRF1 | BRF1, RNA polymerase III transcription initiation factor 90 kDa subunit |
| GUCY2C | guanylate cyclase 2C |
| H2AFX | H2A histone family member X |
| HDAC1 | histone deacetylase 1 |
| HDAC2 | histone deacetylase 2 |
| HIC1 | hypermethylated in cancer 1 |
| HIF1A | hypoxia inducible factor 1 alpha subunit |
| HINT1 | histidine triad nucleotide binding protein 1 |

| | |
|---|---|
| HIVEP1 | human immunodeficiency virus type I enhancer binding protein 1 |
| ZBTB48 | zinc finger and BTB domain containing 48 |
| NR4A1 | nuclear receptor subfamily 4 group A member 1 |
| FOXA1 | forkhead box A1 |
| FOXA2 | forkhead box A2 |
| HNF4A | hepatocyte nuclear factor 4 alpha |
| ONECUT1 | one cut homeobox 1 |
| HPGD | hydroxyprostaglandin dehydrogenase 15-(NAD) |
| HRG | histidine rich glycoprotein |
| HSPD1 | heat shock protein family D (Hsp60) member 1 |
| DNAJB1 | DnaJ heat shock protein family (Hsp40) member B1 |
| IRF8 | interferon regulatory factor 8 |
| ID4 | inhibitor of DNA binding 4, HLH protein |
| IDH1 | isocitrate dehydrogenase (NADP(+)) 1, cytosolic |
| IFI16 | interferon gamma inducible protein 16 |
| IGF1 | insulin like growth factor 1 |
| IGF2R | insulin like growth factor 2 receptor |
| IGFALS | insulin like growth factor binding protein acid labile subunit |
| IGFBP3 | insulin like growth factor binding protein 3 |
| IGFBP4 | insulin like growth factor binding protein 4 |
| IGFBP5 | insulin like growth factor binding protein 5 |
| IGFBP7 | insulin like growth factor binding protein 7 |
| CXCR2 | C-X-C motif chemokine receptor 2 |
| IL17A | interleukin 17A |
| ILK | integrin linked kinase |
| ING1 | inhibitor of growth family member 1 |
| ING2 | inhibitor of growth family member 2 |
| CXCL10 | C-X-C motif chemokine ligand 10 |
| PDX1 | pancreatic and duodenal homeobox 1 |
| IRF1 | interferon regulatory factor 1 |
| IRF3 | interferon regulatory factor 3 |
| IRF4 | interferon regulatory factor 4 |
| IRF5 | interferon regulatory factor 5 |
| ITGA5 | integrin subunit alpha 5 |
| ITGA7 | integrin subunit alpha 7 |
| ITGAV | integrin subunit alpha V |
| ITGB1 | integrin subunit beta 1 |
| ITGB3 | integrin subunit beta 3 |
| JUP | junction plakoglobin |
| CD82 | CD82 molecule |
| KISS1 | KiSS-1 metastasis-suppressor |
| KRT19 | keratin 19 |
| LGALS7 | galectin 7 |
| LIFR | leukemia inhibitory factor receptor alpha |
| LLGL1 | LLGL1, scribble cell polarity complex component |
| VWA5A | von Willebrand factor A domain containing 5A |
| LOX | lysyl oxidase |
| LRMP | lymphoid restricted membrane protein |
| LSAMP | limbic system-associated membrane protein |
| LTF | lactotransferrin |
| MARCKS | myristoylated alanine rich protein kinase C substrate |
| SMAD2 | SMAD family member 2 |
| SMAD4 | SMAD family member 4 |
| MAL | mal, T-cell differentiation protein |

| | |
|---|---|
| MAT2A | methionine adenosyltransferase 2A |
| MAX | MYC associated factor X |
| MCC | mutated in colorectal cancers |
| MAP3K4 | mitogen-activated protein kinase kinase kinase 4 |
| MEN1 | menin 1 |
| MIA2 | melanoma inhibitory activity 2 |
| MLH1 | mutL homolog 1 |
| FOXO4 | forkhead box O4 |
| MME | membrane metalloendopeptidase |
| MNT | MAX network transcriptional repressor |
| MSH2 | mutS homolog 2 |
| MSMB | microseminoprotein beta |
| MST1 | macrophage stimulating 1 |
| MST1R | macrophage stimulating 1 receptor |
| MT1F | metallothionein 1F |
| MT1G | metallothionein 1G |
| MT1M | metallothionein 1M |
| MT2A | metallothionein 2A |
| MTAP | methylthioadenosine phosphorylase |
| MXI1 | MAX interactor 1, dimerization protein |
| GADD45B | growth arrest and DNA damage inducible beta |
| MYH9 | myosin heavy chain 9 |
| MYO1A | myosin IA |
| NBN | nibrin |
| NDN | necdin, MAGE family member |
| NEDD4 | neural precursor cell expressed, developmentally down-regulated 4, E3 ubiquitin protein ligase |
| NF1 | neurofibromin 1 |
| NF2 | neurofibromin 2 |
| NFATC2 | nuclear factor of activated T-cells 2 |
| NFKB1 | nuclear factor kappa B subunit 1 |
| NGFR | nerve growth factor receptor |
| NINJ1 | ninjurin 1 |
| NKX3-1 | NK3 homeobox 1 |
| NNAT | neuronatin |
| NME1 | NME/NM23 nucleoside diphosphate kinase 1 |
| CNOT3 | CCR4-NOT transcription complex subunit 3 |
| NOTCH1 | notch 1 |
| NOTCH2 | notch 2 |
| NOTCH3 | notch 3 |
| NOV | nephroblastoma overexpressed |
| NPAS2 | neuronal PAS domain protein 2 |
| NPM1 | nucleophosmin |
| NRCAM | neuronal cell adhesion molecule |
| NRF1 | nuclear respiratory factor 1 |
| NTRK3 | neurotrophic receptor tyrosine kinase 3 |
| ROR2 | receptor tyrosine kinase like orphan receptor 2 |
| DDR2 | discoidin domain receptor tyrosine kinase 2 |
| NUP98 | nucleoporin 98 |
| OPCML | opioid binding protein/cell adhesion molecule like |
| PEBP1 | phosphatidylethanolamine binding protein 1 |
| PAEP | progestagen associated endometrial protein |
| PAFAH1B1 | platelet activating factor acetylhydrolase 1b regulatory subunit 1 |
| PARK2 | parkin RBR E3 ubiquitin protein ligase |
| PAWR | pro-apoptotic WT1 regulator |

| | |
|---|---|
| PAX4 | paired box 4 |
| PAX5 | paired box 5 |
| PAX6 | paired box 6 |
| PCDHGC3 | protocadherin gamma subfamily C, 3 |
| PCDH8 | protocadherin 8 |
| PCDH9 | protocadherin 9 |
| PDGFRL | platelet derived growth factor receptor like |
| PEG3 | paternally expressed 3 |
| PF4 | platelet factor 4 |
| PFN1 | profilin 1 |
| PGR | progesterone receptor |
| PHB | prohibitin |
| SERPINB5 | serpin family B member 5 |
| SERPINI2 | serpin family I member 2 |
| PIN1 | peptidylprolyl cis/trans isomerase, NIMA-interacting 1 |
| PKD1 | polycystin 1, transient receptor potential channel interacting |
| PKNOX1 | PBX/knotted 1 homeobox 1 |
| PLA2G2A | phospholipase A2 group IIA |
| PLAGL1 | PLAG1 like zinc finger 1 |
| PLCB3 | phospholipase C beta 3 |
| PLCD1 | phospholipase C delta 1 |
| PLD1 | phospholipase D1 |
| PLK1 | polo like kinase 1 |
| PML | promyelocytic leukemia |
| PNN | pinin, desmosome associated protein |
| SEPT4 | septin 4 |
| PPARA | peroxisome proliferator activated receptor alpha |
| PPARG | peroxisome proliferator activated receptor gamma |
| PPM1A | protein phosphatase, Mg2+/Mn2+ dependent 1A |
| PPP1CA | protein phosphatase 1 catalytic subunit alpha |
| PPP2CA | protein phosphatase 2 catalytic subunit alpha |
| PPP2CB | protein phosphatase 2 catalytic subunit beta |
| PPP2R1B | protein phosphatase 2 scaffold subunit Abeta |
| PPP2R2C | protein phosphatase 2 regulatory subunit Bgamma |
| PTPA | protein phosphatase 2 phosphatase activator |
| PPP2R5C | protein phosphatase 2 regulatory subunit B'gamma |
| PPP3CC | protein phosphatase 3 catalytic subunit gamma |
| PRKAA1 | protein kinase AMP-activated catalytic subunit alpha 1 |
| PRKAA2 | protein kinase AMP-activated catalytic subunit alpha 2 |
| PRKAR1A | protein kinase cAMP-dependent type I regulatory subunit alpha |
| PRKCB | protein kinase C beta |
| PRKCD | protein kinase C delta |
| PRKCE | protein kinase C epsilon |
| MAPK9 | mitogen-activated protein kinase 9 |
| MAPK10 | mitogen-activated protein kinase 10 |
| PRODH | proline dehydrogenase 1 |
| PROX1 | prospero homeobox 1 |
| KLK6 | kallikrein related peptidase 6 |
| HTRA1 | HtrA serine peptidase 1 |
| KLK10 | kallikrein related peptidase 10 |
| PTCH1 | patched 1 |
| PTEN | phosphatase and tensin homolog |
| PTGDR | prostaglandin D2 receptor |
| PTPN1 | protein tyrosine phosphatase, non-receptor type 1 |

| | |
|---|---|
| PTPN2 | protein tyrosine phosphatase, non-receptor type 2 |
| PTPN6 | protein tyrosine phosphatase, non-receptor type 6 |
| PTPN11 | protein tyrosine phosphatase, non-receptor type 11 |
| PTPN12 | protein tyrosine phosphatase, non-receptor type 12 |
| PTPN13 | protein tyrosine phosphatase, non-receptor type 13 |
| PTPRC | protein tyrosine phosphatase, receptor type C |
| PTPRD | protein tyrosine phosphatase, receptor type D |
| PTPRJ | protein tyrosine phosphatase, receptor type J |
| PTPRK | protein tyrosine phosphatase, receptor type K |
| RAD23B | RAD23 homolog B, nucleotide excision repair protein |
| RAD51C | RAD51 paralog C |
| RAP1A | RAP1A, member of RAS oncogene family |
| RAP1GAP | RAP1 GTPase activating protein |
| RARB | retinoic acid receptor beta |
| RARRES3 | retinoic acid receptor responder 3 |
| RB1 | RB transcriptional corepressor 1 |
| KDM5A | lysine demethylase 5A |
| RBBP7 | RB binding protein 7, chromatin remodeling factor |
| RBBP8 | RB binding protein 8, endonuclease |
| RBL1 | RB transcriptional corepressor like 1 |
| RBL2 | RB transcriptional corepressor like 2 |
| RBM4 | RNA binding motif protein 4 |
| RBP1 | retinol binding protein 1 |
| RNASEL | ribonuclease L |
| RNH1 | ribonuclease/angiogenin inhibitor 1 |
| ROBO1 | roundabout guidance receptor 1 |
| RPA1 | replication protein A1 |
| RPL5 | ribosomal protein L5 |
| RPL10 | ribosomal protein L10 |
| RPL11 | ribosomal protein L11 |
| RPS6KA2 | ribosomal protein S6 kinase A2 |
| S100A2 | S100 calcium binding protein A2 |
| S100A11 | S100 calcium binding protein A11 |
| SAA1 | serum amyloid A1 |
| SAFB | scaffold attachment factor B |
| SALL2 | spalt like transcription factor 2 |
| CXCL12 | C-X-C motif chemokine ligand 12 |
| SDHA | succinate dehydrogenase complex flavoprotein subunit A |
| SDHB | succinate dehydrogenase complex iron sulfur subunit B |
| SDHD | succinate dehydrogenase complex subunit D |
| SEMA3F | semaphorin 3F |
| MAP2K4 | mitogen-activated protein kinase kinase 4 |
| SFRP1 | secreted frizzled related protein 1 |
| SFRP2 | secreted frizzled related protein 2 |
| SFRP4 | secreted frizzled related protein 4 |
| SFRP5 | secreted frizzled related protein 5 |
| SIAH1 | siah E3 ubiquitin protein ligase 1 |
| SKIL | SKI like proto-oncogene |
| SKP2 | S-phase kinase associated protein 2 |
| SMARCA2 | SWI/SNF related, matrix associated, actin dependent regulator of chromatin, subfamily a, member 2 |
| HLTF | helicase like transcription factor |
| SMARCA4 | SWI/SNF related, matrix associated, actin dependent regulator of chromatin, subfamily a, member 4 |
| SMARCB1 | SWI/SNF related, matrix associated, actin dependent regulator of chromatin, |

| | |
|---|---|
| | subfamily b, member 1 |
| SMARCC1 | SWI/SNF related, matrix associated, actin dependent regulator of chromatin subfamily c member 1 |
| SOD2 | superoxide dismutase 2, mitochondrial |
| SOX1 | SRY-box 1 |
| SOX11 | SRY-box 11 |
| SOX15 | SRY-box 15 |
| SP100 | SP100 nuclear antigen |
| SPARC | secreted protein acidic and cysteine rich |
| SPI1 | Spi-1 proto-oncogene |
| SPTBN1 | spectrin beta, non-erythrocytic 1 |
| ST2 | suppression of tumorigenicity 2 |
| ST5 | suppression of tumorigenicity 5 |
| ST13 | suppression of tumorigenicity 13 (colon carcinoma) (Hsp70 interacting protein) |
| STAT1 | signal transducer and activator of transcription 1 |
| STAT3 | signal transducer and activator of transcription 3 |
| STAT5A | signal transducer and activator of transcription 5A |
| STK10 | serine/threonine kinase 10 |
| STK11 | serine/threonine kinase 11 |
| SYK | spleen associated tyrosine kinase |
| TAGLN | transgelin |
| TAT | tyrosine aminotransferase |
| TBX5 | T-box 5 |
| TCEB3 | transcription elongation factor B subunit 3 |
| TCF4 | transcription factor 4 |
| TCF3 | transcription factor 3 |
| TCF7L2 | transcription factor 7 like 2 |
| TDGF1 | teratocarcinoma-derived growth factor 1 |
| TFAP2A | transcription factor AP-2 alpha |
| TGFB1 | transforming growth factor beta 1 |
| LEFTY2 | left-right determination factor 2 |
| TGFBI | transforming growth factor beta induced |
| TGFBR2 | transforming growth factor beta receptor 2 |
| TGFBR3 | transforming growth factor beta receptor 3 |
| TGM3 | transglutaminase 3 |
| THBD | thrombomodulin |
| THBS1 | thrombospondin 1 |
| THRA | thyroid hormone receptor, alpha |
| THRB | thyroid hormone receptor beta |
| THY1 | Thy-1 cell surface antigen |
| KLF10 | Kruppel like factor 10 |
| TIMP3 | TIMP metallopeptidase inhibitor 3 |
| TNFAIP3 | TNF alpha induced protein 3 |
| TP53 | tumor protein p53 |
| TP53BP1 | tumor protein p53 binding protein 1 |
| TP53BP2 | tumor protein p53 binding protein 2 |
| TP73 | tumor protein p73 |
| NR2C2 | nuclear receptor subfamily 2 group C member 2 |
| HSP90B1 | heat shock protein 90 beta family member 1 |
| TSC1 | tuberous sclerosis 1 |
| TSC2 | tuberous sclerosis 2 |
| TSG101 | tumor susceptibility 101 |
| TSSC1 | tumor suppressing subtransferable candidate 1 |
| PHLDA2 | pleckstrin homology like domain family A member 2 |
| TTC4 | tetratricopeptide repeat domain 4 |

| | |
|---|---|
| TTF1 | transcription termination factor 1 |
| HIRA | histone cell cycle regulator |
| UCHL1 | ubiquitin C-terminal hydrolase L1 |
| KDM6A | lysine demethylase 6A |
| UVRAG | UV radiation resistance associated |
| VDR | vitamin D (1,25- dihydroxyvitamin D3) receptor |
| VEGFA | vascular endothelial growth factor A |
| VHL | von Hippel-Lindau tumor suppressor |
| VIL1 | villin 1 |
| VIM | vimentin |
| VSNL1 | visinin like 1 |
| LAT2 | linker for activation of T-cells family member 2 |
| WNT5A | Wnt family member 5A |
| WNT7A | Wnt family member 7A |
| WNT11 | Wnt family member 11 |
| WT1 | Wilms tumor 1 |
| XIST | X inactive specific transcript (non-protein coding) |
| XRCC5 | X-ray repair cross complementing 5 |
| ZFP36 | ZFP36 ring finger protein |
| ZIC1 | Zic family member 1 |
| PCGF2 | polycomb group ring finger 2 |
| ZBTB16 | zinc finger and BTB domain containing 16 |
| ZNF185 | zinc finger protein 185 (LIM domain) |
| ZYX | zyxin |
| PRDM2 | PR/SET domain 2 |
| BTG2 | BTG anti-proliferation factor 2 |
| SEMA3B | semaphorin 3B |
| RAB7A | RAB7A, member RAS oncogene family |
| PLA2G7 | phospholipase A2 group VII |
| AIMP2 | aminoacyl tRNA synthetase complex interacting multifunctional protein 2 |
| TFPI2 | tissue factor pathway inhibitor 2 |
| ST7 | suppression of tumorigenicity 7 |
| TUSC3 | tumor suppressor candidate 3 |
| NR4A3 | nuclear receptor subfamily 4 group A member 3 |
| NCOA4 | nuclear receptor coactivator 4 |
| CUL5 | cullin 5 |
| CDK2AP1 | cyclin dependent kinase 2 associated protein 1 |
| IFT88 | intraflagellar transport 88 |
| ANP32A | acidic nuclear phosphoprotein 32 family member A |
| MIA | melanoma inhibitory activity |
| MLRL | Myeloid leukemia-related gene (myeloid tumor suppressor) |
| ARID1A | AT-rich interaction domain 1A |
| AXIN1 | axin 1 |
| AXIN2 | axin 2 |
| BAP1 | BRCA1 associated protein 1 |
| MAD1L1 | MAD1 mitotic arrest deficient like 1 |
| SPARCL1 | SPARC like 1 |
| SPOP | speckle type BTB/POZ protein |
| SRPX | sushi repeat containing protein, X-linked |
| NR0B2 | nuclear receptor subfamily 0 group B member 2 |
| RECK | reversion inducing cysteine rich protein with kazal motifs |
| RASAL1 | RAS protein activator like 1 |
| CUL2 | cullin 2 |
| CUL1 | cullin 1 |

| | |
|---|---|
| ST11 | suppression of tumorigenicity 11 (pancreas) |
| PIAS1 | protein inhibitor of activated STAT 1 |
| MADD | MAP kinase activating death domain |
| PDLIM4 | PDZ and LIM domain 4 |
| TMEFF1 | transmembrane protein with EGF like and two follistatin like domains 1 |
| TP63 | tumor protein p63 |
| UNC5C | unc-5 netrin receptor C |
| RNASET2 | ribonuclease T2 |
| PTCH2 | patched 2 |
| NUMB | NUMB, endocytic adaptor protein |
| SOCS1 | suppressor of cytokine signaling 1 |
| EIF3F | eukaryotic translation initiation factor 3 subunit F |
| BECN1 | beclin 1 |
| PEA15 | phosphoprotein enriched in astrocytes 15 |
| TNK1 | tyrosine kinase non receptor 1 |
| EED | embryonic ectoderm development |
| TNFSF12 | tumor necrosis factor superfamily member 12 |
| TNFSF9 | tumor necrosis factor superfamily member 9 |
| FADD | Fas associated via death domain |
| TNFRSF18 | TNF receptor superfamily member 18 |
| DLK1 | delta like non-canonical Notch ligand 1 |
| TNFRSF10B | TNF receptor superfamily member 10b |
| TNFRSF10A | TNF receptor superfamily member 10a |
| TRIM24 | tripartite motif containing 24 |
| INPP4B | inositol polyphosphate-4-phosphatase type II B |
| WISP3 | WNT1 inducible signaling pathway protein 3 |
| HDAC3 | histone deacetylase 3 |
| DLEU2 | deleted in lymphocytic leukemia 2 (non-protein coding) |
| TSC22D1 | TSC22 domain family member 1 |
| ALDH1A2 | aldehyde dehydrogenase 1 family member A2 |
| NR1I2 | nuclear receptor subfamily 1 group I member 2 |
| PER2 | period circadian clock 2 |
| IER3 | immediate early response 3 |
| BCL10 | B-cell CLL/lymphoma 10 |
| MBD4 | methyl-CpG binding domain 4, DNA glycosylase |
| SELENBP1 | selenium binding protein 1 |
| LIMD1 | LIM domains containing 1 |
| SOCS3 | suppressor of cytokine signaling 3 |
| RNF8 | ring finger protein 8 |
| DOK2 | docking protein 2 |
| AIP | aryl hydrocarbon receptor interacting protein |
| GPRC5A | G protein-coupled receptor class C group 5 member A |
| CLDN1 | claudin 1 |
| DIRAS3 | DIRAS family GTPase 3 |
| DNAJA3 | DnaJ heat shock protein family (Hsp40) member A3 |
| LATS1 | large tumor suppressor kinase 1 |
| PDCD5 | programmed cell death 5 |
| NEURL1 | neuralized E3 ubiquitin protein ligase 1 |
| AIMP1 | aminoacyl tRNA synthetase complex interacting multifunctional protein 1 |
| BCL7C | BCL tumor suppressor 7C |
| BCL7B | BCL tumor suppressor 7B |
| KLF4 | Kruppel like factor 4 |
| COPS2 | COP9 signalosome subunit 2 |
| SLIT2 | slit guidance ligand 2 |

| | |
|---|---|
| KL | klotho |
| SLC9A3R1 | SLC9A3 regulator 1 |
| ARHGAP29 | Rho GTPase activating protein 29 |
| ABCG2 | ATP binding cassette subfamily G member 2 (Junior blood group) |
| AIM2 | absent in melanoma 2 |
| HOMER2 | homer scaffolding protein 2 |
| RASAL2 | RAS protein activator like 2 |
| CHST10 | carbohydrate sulfotransferase 10 |
| LITAF | lipopolysaccharide induced TNF factor |
| EEF1E1 | eukaryotic translation elongation factor 1 epsilon 1 |
| EI24 | EI24, autophagy associated transmembrane protein |
| CXCL14 | C-X-C motif chemokine ligand 14 |
| AKAP12 | A-kinase anchoring protein 12 |
| ISG15 | ISG15 ubiquitin-like modifier |
| MDC1 | mediator of DNA damage checkpoint 1 |
| SAFB2 | scaffold attachment factor B2 |
| LZTS3 | leucine zipper tumor suppressor family member 3 |
| RASSF2 | Ras association domain family member 2 |
| RNF144A | ring finger protein 144A |
| MTSS1 | MTSS1, I-BAR domain containing |
| RB1CC1 | RB1 inducible coiled-coil 1 |
| NUAK1 | NUAK family kinase 1 |
| SRGAP3 | SLIT-ROBO Rho GTPase activating protein 3 |
| DCLRE1A | DNA cross-link repair 1A |
| DLEC1 | deleted in lung and esophageal cancer 1 |
| DMTF1 | cyclin D binding myb like transcription factor 1 |
| TANK | TRAF family member associated NFKB activator |
| BCL2L11 | BCL2 like 11 |
| SH2B3 | SH2B adaptor protein 3 |
| RANBP9 | RAN binding protein 9 |
| TSPAN32 | tetraspanin 32 |
| TSSC4 | tumor suppressing subtransferable candidate 4 |
| ABI2 | abl interactor 2 |
| PLXNC1 | plexin C1 |
| RBM6 | RNA binding motif protein 6 |
| RBM5 | RNA binding motif protein 5 |
| TRIM13 | tripartite motif containing 13 |
| TOPORS | TOP1 binding arginine/serine rich protein |
| CTDSPL | CTD small phosphatase like |
| SPRY2 | sprouty RTK signaling antagonist 2 |
| STUB1 | STIP1 homology and U-box containing protein 1 |
| UBE4B | ubiquitination factor E4B |
| DLEU1 | deleted in lymphocytic leukemia 1 |
| IKZF1 | IKAROS family zinc finger 1 |
| MRVI1 | murine retrovirus integration site 1 homolog |
| CITED2 | Cbp/p300 interacting transactivator with Glu/Asp rich carboxy-terminal domain 2 |
| DLC1 | DLC1 Rho GTPase activating protein |
| NDRG1 | N-myc downstream regulated 1 |
| RACK1 | receptor for activated C kinase 1 |
| BASP1 | brain abundant membrane attached signal protein 1 |
| YAP1 | Yes associated protein 1 |
| PGRMC2 | progesterone receptor membrane component 2 |
| RBM14 | RNA binding motif protein 14 |
| ZBTB18 | zinc finger and BTB domain containing 18 |

| | |
|---|---|
| HOXB13 | homeobox B13 |
| MYBBP1A | MYB binding protein 1a |
| KAT5 | lysine acetyltransferase 5 |
| ARL6IP5 | ADP ribosylation factor like GTPase 6 interacting protein 5 |
| HTATIP2 | HIV-1 Tat interactive protein 2 |
| OLFM4 | olfactomedin 4 |
| TRIM3 | tripartite motif containing 3 |
| TXNIP | thioredoxin interacting protein |
| RASL10A | RAS like family 10 member A |
| LEFTY1 | left-right determination factor 1 |
| NPRL2 | NPR2-like, GATOR1 complex subunit |
| SPINT2 | serine peptidase inhibitor, Kunitz type 2 |
| CTCF | CCCTC-binding factor |
| AHCYL1 | adenosylhomocysteinase like 1 |
| PLK2 | polo like kinase 2 |
| ZMYND11 | zinc finger MYND-type containing 11 |
| IQGAP2 | IQ motif containing GTPase activating protein 2 |
| FRS3 | fibroblast growth factor receptor substrate 3 |
| BLCAP | bladder cancer associated protein |
| GADD45G | growth arrest and DNA damage inducible gamma |
| BTG3 | BTG anti-proliferation factor 3 |
| IL24 | interleukin 24 |
| GLIPR1 | GLI pathogenesis related 1 |
| CYB561D2 | cytochrome b561 family member D2 |
| TRIM31 | tripartite motif containing 31 |
| DNAJB4 | DnaJ heat shock protein family (Hsp40) member B4 |
| DIDO1 | death inducer-obliterator 1 |
| ADAMTS8 | ADAM metallopeptidase with thrombospondin type 1 motif 8 |
| PRDM5 | PR/SET domain 5 |
| PRDM4 | PR/SET domain 4 |
| PTPRT | protein tyrosine phosphatase, receptor type T |
| PLA2G16 | phospholipase A2 group XVI |
| LZTS1 | leucine zipper tumor suppressor 1 |
| MAP4K1 | mitogen-activated protein kinase kinase kinase kinase 1 |
| RASSF1 | Ras association domain family member 1 |
| PTENP1 | phosphatase and tensin homolog pseudogene 1 |
| WIF1 | WNT inhibitory factor 1 |
| CHEK2 | checkpoint kinase 2 |
| TREX2 | three prime repair exonuclease 2 |
| RASSF8 | Ras association domain family member 8 |
| POU6F2 | POU class 6 homeobox 2 |
| PARK7 | Parkinsonism associated deglycase |
| TUSC2 | tumor suppressor candidate 2 |
| GABARAP | GABA type A receptor-associated protein |
| IKZF3 | IKAROS family zinc finger 3 |
| IKZF2 | IKAROS family zinc finger 2 |
| ZHX2 | zinc fingers and homeoboxes 2 |
| PLA2R1 | phospholipase A2 receptor 1 |
| SIRT2 | sirtuin 2 |
| DKK1 | dickkopf WNT signaling pathway inhibitor 1 |
| TRIM32 | tripartite motif containing 32 |
| USP33 | ubiquitin specific peptidase 33 |
| PHLPP2 | PH domain and leucine rich repeat protein phosphatase 2 |
| ZNF292 | zinc finger protein 292 |

| | |
|---|---|
| PDS5B | PDS5 cohesin associated factor B |
| TRIM35 | tripartite motif containing 35 |
| KIF1B | kinesin family member 1B |
| EPB41L3 | erythrocyte membrane protein band 4.1 like 3 |
| CIC | capicua transcriptional repressor |
| KANK1 | KN motif and ankyrin repeat domains 1 |
| GANAB | glucosidase II alpha subunit |
| RHOBTB2 | Rho related BTB domain containing 2 |
| PHLPP1 | PH domain and leucine rich repeat protein phosphatase 1 |
| CAMTA1 | calmodulin binding transcription activator 1 |
| MTUS2 | microtubule associated tumor suppressor candidate 2 |
| ATMIN | ATM interactor |
| NEDD4L | neural precursor cell expressed, developmentally down-regulated 4-like, E3 ubiquitin protein ligase |
| SASH1 | SAM and SH3 domain containing 1 |
| SYNM | synemin |
| SMCHD1 | structural maintenance of chromosomes flexible hinge domain containing 1 |
| ARHGEF12 | Rho guanine nucleotide exchange factor 12 |
| UFL1 | UFM1 specific ligase 1 |
| DICER1 | dicer 1, ribonuclease III |
| SIRT4 | sirtuin 4 |
| SIRT3 | sirtuin 3 |
| SIRT1 | sirtuin 1 |
| CBX5 | chromobox 5 |
| SUZ12 | SUZ12 polycomb repressive complex 2 subunit |
| SCRIB | scribbled planar cell polarity protein |
| SEC14L2 | SEC14 like lipid binding 2 |
| GTPBP4 | GTP binding protein 4 |
| CCNDBP1 | cyclin D1 binding protein 1 |
| DDX58 | DEXD/H-box helicase 58 |
| DAPK2 | death associated protein kinase 2 |
| PHLDA3 | pleckstrin homology like domain family A member 3 |
| SSBP2 | single stranded DNA binding protein 2 |
| TMEFF2 | transmembrane protein with EGF like and two follistatin like domains 2 |
| CADM1 | cell adhesion molecule 1 |
| CIZ1 | CDKN1A interacting zinc finger protein 1 |
| POU2F3 | POU class 2 homeobox 3 |
| BRMS1 | breast cancer metastasis suppressor 1 |
| RCHY1 | ring finger and CHY zinc finger domain containing 1 |
| PTPN23 | protein tyrosine phosphatase, non-receptor type 23 |
| LRIG1 | leucine rich repeats and immunoglobulin like domains 1 |
| CHD5 | chromodomain helicase DNA binding protein 5 |
| CNTNAP2 | contactin associated protein-like 2 |
| TES | testin LIM domain protein |
| NKX2-8 | NK2 homeobox 8 |
| FBXO25 | F-box protein 25 |
| EHF | ETS homologous factor |
| LHX6 | LIM homeobox 6 |
| NUPR1 | nuclear protein 1, transcriptional regulator |
| INTS6 | integrator complex subunit 6 |
| LATS2 | large tumor suppressor kinase 2 |
| GREM1 | gremlin 1, DAN family BMP antagonist |
| TBL2 | transducin beta like 2 |
| SNORD50A | small nucleolar RNA, C/D box 50A |
| HBP1 | HMG-box transcription factor 1 |

| | |
|---|---|
| FOXD3 | forkhead box D3 |
| MLH3 | mutL homolog 3 |
| TSPAN13 | tetraspanin 13 |
| FOXP1 | forkhead box P1 |
| BBC3 | BCL2 binding component 3 |
| DKK3 | dickkopf WNT signaling pathway inhibitor 3 |
| HSPB7 | heat shock protein family B (small) member 7 |
| CPNE7 | copine 7 |
| GLS2 | glutaminase 2 |
| SLC39A1 | solute carrier family 39 member 1 |
| GNMT | glycine N-methyltransferase |
| PDCD4 | programmed cell death 4 (neoplastic transformation inhibitor) |
| PCDH17 | protocadherin 17 |
| BMP10 | bone morphogenetic protein 10 |
| RBMS3 | RNA binding motif single stranded interacting protein 3 |
| RBMX | RNA binding motif protein, X-linked |
| RPS6KA6 | ribosomal protein S6 kinase A6 |
| HTRA2 | HtrA serine peptidase 2 |
| HIPK2 | homeodomain interacting protein kinase 2 |
| SETD2 | SET domain containing 2 |
| PYCARD | PYD and CARD domain containing |
| BRD7 | bromodomain containing 7 |
| CTNNA3 | catenin alpha 3 |
| BLNK | B-cell linker |
| UBIAD1 | UbiA prenyltransferase domain containing 1 |
| OSGIN1 | oxidative stress induced growth inhibitor 1 |
| NRBP1 | nuclear receptor binding protein 1 |
| GLTSCR2 | glioma tumor suppressor candidate region gene 2 |
| GLTSCR1 | glioma tumor suppressor candidate region gene 1 |
| EHD3 | EH domain containing 3 |
| CRNN | cornulin |
| G0S2 | G0/G1 switch 2 |
| DEC1 | deleted in esophageal cancer 1 |
| FOXP3 | forkhead box P3 |
| TSG11 | Tumor suppressor gene on chromosome 11 |
| SH3GLB1 | SH3 domain containing GRB2 like endophilin B1 |
| ANGPTL4 | angiopoietin like 4 |
| ING4 | inhibitor of growth family member 4 |
| PLEKHO1 | pleckstrin homology domain containing O1 |
| PLCE1 | phospholipase C epsilon 1 |
| ZDHHC2 | zinc finger DHHC-type containing 2 |
| MZB1 | marginal zone B and B1 cell specific protein |
| PAIP2 | poly(A) binding protein interacting protein 2 |
| TNFRSF12A | TNF receptor superfamily member 12A |
| DACT1 | dishevelled binding antagonist of beta catenin 1 |
| FZR1 | fizzy/cell division cycle 20 related 1 |
| ZMYND10 | zinc finger MYND-type containing 10 |
| NOL7 | nucleolar protein 7 |
| DCDC2 | doublecortin domain containing 2 |
| LIMA1 | LIM domain and actin binding 1 |
| CXXC5 | CXXC finger protein 5 |
| SIRT6 | sirtuin 6 |
| SUFU | SUFU negative regulator of hedgehog signaling |
| HECA | hdc homolog, cell cycle regulator |

| | |
|---|---|
| CYB5R2 | cytochrome b5 reductase 2 |
| UIMC1 | ubiquitin interaction motif containing 1 |
| WWOX | WW domain containing oxidoreductase |
| KDM3B | lysine demethylase 3B |
| LRP1B | LDL receptor related protein 1B |
| ERRFI1 | ERBB receptor feedback inhibitor 1 |
| ING3 | inhibitor of growth family member 3 |
| EGLN1 | egl-9 family hypoxia inducible factor 1 |
| XAF1 | XIAP associated factor 1 |
| IL17RD | interleukin 17 receptor D |
| BTG4 | BTG anti-proliferation factor 4 |
| RNF111 | ring finger protein 111 |
| TET2 | tet methylcytosine dioxygenase 2 |
| TRIT1 | tRNA isopentenyltransferase 1 |
| ESRP1 | epithelial splicing regulatory protein 1 |
| WHSC1L1 | Wolf-Hirschhorn syndrome candidate 1-like 1 |
| BANP | BTG3 associated nuclear protein |
| HRASLS2 | HRAS like suppressor 2 |
| PINX1 | PIN2/TERF1 interacting, telomerase inhibitor 1 |
| PIWIL2 | piwi like RNA-mediated gene silencing 2 |
| SHQ1 | SHQ1, H/ACA ribonucleoprotein assembly factor |
| PBRM1 | polybromo 1 |
| TRIM62 | tripartite motif containing 62 |
| MOB1A | MOB kinase activator 1A |
| CASC1 | cancer susceptibility candidate 1 |
| VPS53 | VPS53, GARP complex subunit |
| FBXW7 | F-box and WD repeat domain containing 7 |
| MEG3 | maternally expressed 3 (non-protein coding) |
| CAMK2N1 | calcium/calmodulin dependent protein kinase II inhibitor 1 |
| RBM38 | RNA binding motif protein 38 |
| VEZT | vezatin, adherens junctions transmembrane protein |
| PRR5 | proline rich 5 |
| SLC39A4 | solute carrier family 39 member 4 |
| TMEM127 | transmembrane protein 127 |
| WDR11 | WD repeat domain 11 |
| DNAJC11 | DnaJ heat shock protein family (Hsp40) member C11 |
| CHFR | checkpoint with forkhead and ring finger domains |
| CNDP2 | CNDP dipeptidase 2 (metallopeptidase M20 family) |
| CCAR1 | cell division cycle and apoptosis regulator 1 |
| CACNA2D3 | calcium voltage-gated channel auxiliary subunit alpha2delta 3 |
| KDM3A | lysine demethylase 3A |
| EAF2 | ELL associated factor 2 |
| THSD1 | thrombospondin type 1 domain containing 1 |
| AJAP1 | adherens junctions associated protein 1 |
| GKN1 | gastrokine 1 |
| RPRM | reprimo, TP53 dependent G2 arrest mediator candidate |
| DIABLO | diablo IAP-binding mitochondrial protein |
| PANX2 | pannexin 2 |
| TCEAL7 | transcription elongation factor A like 7 |
| LXN | latexin |
| DUSP22 | dual specificity phosphatase 22 |
| NIT2 | nitrilase family member 2 |
| PRDM11 | PR/SET domain 11 |
| CTNNBIP1 | catenin beta interacting protein 1 |

| | |
|---|---|
| ADAMTS9 | ADAM metallopeptidase with thrombospondin type 1 motif 9 |
| PDSS2 | prenyl (decaprenyl) diphosphate synthase, subunit 2 |
| RAB25 | RAB25, member RAS oncogene family |
| RTN4 | reticulon 4 |
| SALL4 | spalt like transcription factor 4 |
| SCYL1 | SCY1 like pseudokinase 1 |
| NDRG2 | NDRG family member 2 |
| AHRR | aryl-hydrocarbon receptor repressor |
| MTUS1 | microtubule associated tumor suppressor 1 |
| XPO5 | exportin 5 |
| HACE1 | HECT domain and ankyrin repeat containing E3 ubiquitin protein ligase 1 |
| PCDH10 | protocadherin 10 |
| SYT13 | synaptotagmin 13 |
| WDR48 | WD repeat domain 48 |
| ZBTB4 | zinc finger and BTB domain containing 4 |
| NCOA5 | nuclear receptor coactivator 5 |
| SCUBE2 | signal peptide, CUB domain and EGF like domain containing 2 |
| CCAR2 | cell cycle and apoptosis regulator 2 |
| CADM3 | cell adhesion molecule 3 |
| WFDC1 | WAP four-disulfide core domain 1 |
| KMT2C | lysine methyltransferase 2C |
| HIVEP3 | human immunodeficiency virus type I enhancer binding protein 3 |
| EDA2R | ectodysplasin A2 receptor |
| RINT1 | RAD50 interactor 1 |
| GAS5 | growth arrest specific 5 (non-protein coding) |
| BCORL1 | BCL6 corepressor-like 1 |
| LRRC4 | leucine rich repeat containing 4 |
| RFWD2 | ring finger and WD repeat domain 2 |
| CSMD1 | CUB and Sushi multiple domains 1 |
| NDST4 | N-deacetylase and N-sulfotransferase 4 |
| CSRNP1 | cysteine and serine rich nuclear protein 1 |
| ANAPC1 | anaphase promoting complex subunit 1 |
| CCDC136 | coiled-coil domain containing 136 |
| CDCP1 | CUB domain containing protein 1 |
| NDRG4 | NDRG family member 4 |
| WNK2 | WNK lysine deficient protein kinase 2 |
| PHACTR4 | phosphatase and actin regulator 4 |
| DUSP26 | dual specificity phosphatase 26 (putative) |
| AHNAK | AHNAK nucleoprotein |
| IRX1 | iroquois homeobox 1 |
| BHLHE41 | basic helix-loop-helix family member e41 |
| CDC73 | cell division cycle 73 |
| TNFAIP8L2 | TNF alpha induced protein 8 like 2 |
| FAT4 | FAT atypical cadherin 4 |
| MCPH1 | microcephalin 1 |
| PALB2 | partner and localizer of BRCA2 |
| ZNF668 | zinc finger protein 668 |
| FBXO31 | F-box protein 31 |
| ARMC5 | armadillo repeat containing 5 |
| KDM8 | lysine demethylase 8 |
| GGNBP2 | gametogenetin binding protein 2 |
| DOK3 | docking protein 3 |
| DENND2D | DENN domain containing 2D |
| PHC3 | polyhomeotic homolog 3 |

| | |
|---|---|
| FAM188A | family with sequence similarity 188 member A |
| NRSN2 | neurensin 2 |
| MUS81 | MUS81 structure-specific endonuclease subunit |
| FER1L4 | fer-1 like family member 4, pseudogene |
| CXXC4 | CXXC finger protein 4 |
| SPRY4 | sprouty RTK signaling antagonist 4 |
| RASSF5 | Ras association domain family member 5 |
| SOX7 | SRY-box 7 |
| YPEL3 | yippee like 3 |
| MARVELD1 | MARVEL domain containing 1 |
| ARMC10 | armadillo repeat containing 10 |
| RASSF4 | Ras association domain family member 4 |
| FAM172A | family with sequence similarity 172 member A |
| PPP1R1B | protein phosphatase 1 regulatory inhibitor subunit 1B |
| TCHP | trichoplein keratin filament binding |
| ING5 | inhibitor of growth family member 5 |
| PHF6 | PHD finger protein 6 |
| BRMS1L | breast cancer metastasis-suppressor 1-like |
| C2orf40 | chromosome 2 open reading frame 40 |
| LZTS2 | leucine zipper tumor suppressor 2 |
| BRSK1 | BR serine/threonine kinase 1 |
| SLX4 | SLX4 structure-specific endonuclease subunit |
| HOPX | HOP homeobox |
| AFAP1L2 | actin filament associated protein 1 like 2 |
| SPINK7 | serine peptidase inhibitor, Kazal type 7 (putative) |
| MYO18B | myosin XVIIIB |
| BEX2 | brain expressed X-linked 2 |
| RTN4IP1 | reticulon 4 interacting protein 1 |
| ZC3H10 | zinc finger CCCH-type containing 10 |
| MFSD2A | major facilitator superfamily domain containing 2A |
| TBRG1 | transforming growth factor beta regulator 1 |
| ZNF382 | zinc finger protein 382 |
| RITA1 | RBPJ interacting and tubulin associated 1 |
| EAF1 | ELL associated factor 1 |
| TSLP | thymic stromal lymphopoietin |
| TRIM15 | tripartite motif containing 15 |
| LHX4 | LIM homeobox 4 |
| UNC5A | unc-5 netrin receptor A |
| BMF | Bcl2 modifying factor |
| GADD45GIP1 | GADD45G interacting protein 1 |
| STARD13 | StAR related lipid transfer domain containing 13 |
| CREB3L1 | cAMP responsive element binding protein 3 like 1 |
| L3MBTL4 | l(3)mbt-like 4 (Drosophila) |
| RASL10B | RAS like family 10 member B |
| CABLES1 | Cdk5 and Abl enzyme substrate 1 |
| SCGB3A1 | secretoglobin family 3A member 1 |
| STRADA | STE20-related kinase adaptor alpha |
| GORAB | golgin, RAB6 interacting |
| MOB1B | MOB kinase activator 1B |
| TPTE2 | transmembrane phosphoinositide 3-phosphatase and tensin homolog 2 |
| HTRA3 | HtrA serine peptidase 3 |
| TP53INP1 | tumor protein p53 inducible nuclear protein 1 |
| EGLN3 | egl-9 family hypoxia inducible factor 3 |
| PRKCDBP | protein kinase C delta binding protein |

| | |
|---|---|
| CYGB | cytoglobin |
| SMYD4 | SET and MYND domain containing 4 |
| FBXO32 | F-box protein 32 |
| UHRF2 | ubiquitin like with PHD and ring finger domains 2 |
| ARL11 | ADP ribosylation factor like GTPase 11 |
| BATF2 | basic leucine zipper ATF-like transcription factor 2 |
| LRRC3B | leucine rich repeat containing 3B |
| CMTM5 | CKLF like MARVEL transmembrane domain containing 5 |
| MIA2 | melanoma inhibitory activity 2 |
| TWIST2 | twist family bHLH transcription factor 2 |
| LRIG3 | leucine rich repeats and immunoglobulin like domains 3 |
| JDP2 | Jun dimerization protein 2 |
| DCUN1D3 | defective in cullin neddylation 1 domain containing 3 |
| CMTM3 | CKLF like MARVEL transmembrane domain containing 3 |
| OVCA2 | ovarian tumor suppressor candidate 2 |
| PLK5 | polo like kinase 5 |
| OSCP1 | organic solute carrier partner 1 |
| ACVR1C | activin A receptor type 1C |
| UBE2QL1 | ubiquitin conjugating enzyme E2 Q family like 1 |
| PACRG | PARK2 coregulated |
| CLDN23 | claudin 23 |
| UNC5D | unc-5 netrin receptor D |
| AMER1 | APC membrane recruitment protein 1 |
| GATA5 | GATA binding protein 5 |
| CTCFL | CCCTC-binding factor like |
| ST13P5 | suppression of tumorigenicity 13 (colon carcinoma) (Hsp70 interacting protein) pseudogene 5 |
| PRICKLE1 | prickle planar cell polarity protein 1 |
| ST13P3 | suppression of tumorigenicity 13 (colon carcinoma) (Hsp70 interacting protein) pseudogene 3 |
| ST13P4 | suppression of tumorigenicity 13 (colon carcinoma) (Hsp70 interacting protein) pseudogene 4 |
| TOM1L2 | target of myb1 like 2 membrane trafficking protein |
| DIRAS1 | DIRAS family GTPase 1 |
| PYHIN1 | pyrin and HIN domain family member 1 |
| SIK1 | salt inducible kinase 1 |
| PPM1L | protein phosphatase, Mg2+/Mn2+ dependent 1L |
| SHISA3 | shisa family member 3 |
| DAB2IP | DAB2 interacting protein |
| ST13P7 | suppression of tumorigenicity 13 (colon carcinoma) (Hsp70 interacting protein) pseudogene 7 |
| ST13P6 | suppression of tumorigenicity 13 (colon carcinoma) (Hsp70 interacting protein) pseudogene 6 |
| DEUP1 | deuterosome assembly protein 1 |
| LMNTD1 | lamin tail domain containing 1 |
| SLC5A8 | solute carrier family 5 member 8 |
| TMPRSS6 | transmembrane protease, serine 6 |
| ZNF366 | zinc finger protein 366 |
| ADAMTS18 | ADAM metallopeptidase with thrombospondin type 1 motif 18 |
| ASXL1 | additional sex combs like 1, transcriptional regulator |
| SYNPO2 | synaptopodin 2 |
| ARID2 | AT-rich interaction domain 2 |
| CADM4 | cell adhesion molecule 4 |
| GKN2 | gastrokine 2 |
| FLCN | folliculin |
| ZBTB7C | zinc finger and BTB domain containing 7C |

| | |
|---|---|
| SAMD9L | sterile alpha motif domain containing 9 like |
| USP12 | ubiquitin specific peptidase 12 |
| UNC5B | unc-5 netrin receptor B |
| HEPACAM | hepatic and glial cell adhesion molecule |
| FBXL13 | F-box and leucine rich repeat protein 13 |
| NAPEPLD | N-acyl phosphatidylethanolamine phospholipase D |
| CADM2 | cell adhesion molecule 2 |
| EBF3 | early B-cell factor 3 |
| MCM9 | minichromosome maintenance 9 homologous recombination repair factor |
| CASC2 | cancer susceptibility candidate 2 (non-protein coding) |
| BCL6B | B-cell CLL/lymphoma 6B |
| SHPRH | SNF2 histone linker PHD RING helicase |
| SGMS1 | sphingomyelin synthase 1 |
| ITS | Insulinoma tumor suppressor gene locus |
| H19 | H19, imprinted maternally expressed transcript (non-protein coding) |
| RASSF3 | Ras association domain family member 3 |
| KCNRG | potassium channel regulator |
| ZFP82 | ZFP82 zinc finger protein |
| TUSC7 | tumor suppressor candidate 7 (non-protein coding) |
| RNF180 | ring finger protein 180 |
| TUSC1 | tumor suppressor candidate 1 |
| TUSC5 | tumor suppressor candidate 5 |
| LIN9 | lin-9 DREAM MuvB core complex component |
| MT1DP | metallothionein 1D, pseudogene |
| HCAR2 | hydroxycarboxylic acid receptor 2 |
| ST13P8 | suppression of tumorigenicity 13 (colon carcinoma) (Hsp70 interacting protein) pseudogene 8 |
| TMPRSS11A | transmembrane protease, serine 11A |
| ST13P2 | suppression of tumorigenicity 13 (colon carcinoma) (Hsp70 interacting protein) pseudogene 2 |
| IGFBPL1 | insulin like growth factor binding protein like 1 |
| PWAR4 | Prader Willi/Angelman region RNA 4 |
| TUSC2P1 | tumor suppressor candidate 2 pseudogene 1 |
| DND1 | DND microRNA-mediated repression inhibitor 1 |
| KIF7 | kinesin family member 7 |
| CENPS | centromere protein S |
| LINC-PINT | long intergenic non-protein coding RNA, p53 induced transcript |
| RASL11A | RAS like family 11 member A |
| ST13P9 | suppression of tumorigenicity 13 (colon carcinoma) (Hsp70 interacting protein) pseudogene 9 |
| VHLL | von Hippel-Lindau tumor suppressor like |
| ST13P10 | suppression of tumorigenicity 13 (colon carcinoma) (Hsp70 interacting protein) pseudogene 10 |
| TUSC8 | tumor suppressor candidate 8 (non-protein coding) |
| ST20 | suppressor of tumorigenicity 20 |
| LOC401317 | uncharacterized LOC401317 |
| HACD4 | 3-hydroxyacyl-CoA dehydratase 4 |
| ST13P13 | suppression of tumorigenicity 13 (colon carcinoma) (Hsp70 interacting protein) pseudogene 13 |
| MIRLET7A1 | microRNA let-7a-1 |
| MIRLET7A2 | microRNA let-7a-2 |
| MIRLET7A3 | microRNA let-7a-3 |
| MIRLET7B | microRNA let-7b |
| MIRLET7C | microRNA let-7c |
| MIRLET7D | microRNA let-7d |
| MIRLET7E | microRNA let-7e |

| | |
|---|---|
| MIRLET7F1 | microRNA let-7f-1 |
| MIRLET7F2 | microRNA let-7f-2 |
| MIRLET7G | microRNA let-7g |
| MIRLET7I | microRNA let-7i |
| MIR100 | microRNA 100 |
| MIR101-1 | microRNA 101-1 |
| MIR101-2 | microRNA 101-2 |
| MIR106A | microRNA 106a |
| MIR107 | microRNA 107 |
| MIR10A | microRNA 10a |
| MIR1-1 | microRNA 1-1 |
| MIR1-2 | microRNA 1-2 |
| MIR122 | microRNA 122 |
| MIR124-1 | microRNA 124-1 |
| MIR124-2 | microRNA 124-2 |
| MIR124-3 | microRNA 124-3 |
| MIR125A | microRNA 125a |
| MIR125B1 | microRNA 125b-1 |
| MIR125B2 | microRNA 125b-2 |
| MIR126 | microRNA 126 |
| MIR127 | microRNA 127 |
| MIR129-1 | microRNA 129-1 |
| MIR129-2 | microRNA 129-2 |
| MIR130A | microRNA 130a |
| MIR132 | microRNA 132 |
| MIR133A1 | microRNA 133a-1 |
| MIR133A2 | microRNA 133a-2 |
| MIR134 | microRNA 134 |
| MIR135A1 | microRNA 135a-1 |
| MIR135A2 | microRNA 135a-2 |
| MIR136 | microRNA 136 |
| MIR137 | microRNA 137 |
| MIR138-1 | microRNA 138-1 |
| MIR138-2 | microRNA 138-2 |
| MIR140 | microRNA 140 |
| MIR141 | microRNA 141 |
| MIR142 | microRNA 142 |
| MIR143 | microRNA 143 |
| MIR145 | microRNA 145 |
| MIR146A | microRNA 146a |
| MIR147A | microRNA 147a |
| MIR148A | microRNA 148a |
| MIR149 | microRNA 149 |
| MIR150 | microRNA 150 |
| MIR152 | microRNA 152 |
| MIR155 | microRNA 155 |
| MIR15A | microRNA 15a |
| MIR16-1 | microRNA 16-1 |
| MIR16-2 | microRNA 16-2 |
| MIR17 | microRNA 17 |
| MIR18A | microRNA 18a |
| MIR181A2 | microRNA 181a-2 |
| MIR181B1 | microRNA 181b-1 |
| MIR181B2 | microRNA 181b-2 |

| | |
|---|---|
| MIR181C | microRNA 181c |
| MIR182 | microRNA 182 |
| MIR183 | microRNA 183 |
| MIR185 | microRNA 185 |
| MIR186 | microRNA 186 |
| MIR187 | microRNA 187 |
| MIR192 | microRNA 192 |
| MIR193A | microRNA 193a |
| MIR194-1 | microRNA 194-1 |
| MIR194-2 | microRNA 194-2 |
| MIR195 | microRNA 195 |
| MIR196A2 | microRNA 196a-2 |
| MIR198 | microRNA 198 |
| MIR199A1 | microRNA 199a-1 |
| MIR20A | microRNA 20a |
| MIR200A | microRNA 200a |
| MIR200B | microRNA 200b |
| MIR200C | microRNA 200c |
| MIR203A | microRNA 203a |
| MIR204 | microRNA 204 |
| MIR205 | microRNA 205 |
| MIR206 | microRNA 206 |
| MIR210 | microRNA 210 |
| MIR211 | microRNA 211 |
| MIR181A1 | microRNA 181a-1 |
| MIR214 | microRNA 214 |
| MIR215 | microRNA 215 |
| MIR217 | microRNA 217 |
| MIR218-1 | microRNA 218-1 |
| MIR218-2 | microRNA 218-2 |
| MIR219A1 | microRNA 219a-1 |
| MIR22 | microRNA 22 |
| MIR222 | microRNA 222 |
| MIR223 | microRNA 223 |
| MIR23A | microRNA 23a |
| MIR23B | microRNA 23b |
| MIR24-1 | microRNA 24-1 |
| MIR25 | microRNA 25 |
| MIR26A1 | microRNA 26a-1 |
| MIR26A2 | microRNA 26a-2 |
| MIR26B | microRNA 26b |
| MIR27A | microRNA 27a |
| MIR27B | microRNA 27b |
| MIR28 | microRNA 28 |
| MIR29A | microRNA 29a |
| MIR296 | microRNA 296 |
| MIR29B1 | microRNA 29b-1 |
| MIR29C | microRNA 29c |
| MIR30A | microRNA 30a |
| MIR30C1 | microRNA 30c-1 |
| MIR31 | microRNA 31 |
| MIR320A | microRNA 320a |
| MIR33A | microRNA 33a |
| MIR34A | microRNA 34a |

| | |
|---|---|
| MIR34B | microRNA 34b |
| MIR34C | microRNA 34c |
| MIR7-1 | microRNA 7-1 |
| MIR7-2 | microRNA 7-2 |
| MIR7-3 | microRNA 7-3 |
| MIR9-1 | microRNA 9-1 |
| MIR9-2 | microRNA 9-2 |
| MIR9-3 | microRNA 9-3 |
| MIR98 | microRNA 98 |
| MIR99A | microRNA 99a |
| BLID | BH3-like motif containing, cell death inducer |
| LOC440311 | glioma tumor suppressor candidate region gene 2 pseudogene |
| ZFAS1 | ZNFX1 antisense RNA 1 |
| ST13P17 | suppression of tumorigenicity 13 (colon carcinoma) (Hsp70 interacting protein) pseudogene 17 |
| MIR148B | microRNA 148b |
| MIR302B | microRNA 302b |
| MIR326 | microRNA 326 |
| MIR335 | microRNA 335 |
| MIR338 | microRNA 338 |
| MIR340 | microRNA 340 |
| MIR367 | microRNA 367 |
| MIR370 | microRNA 370 |
| MIR196B | microRNA 196b |
| JST | tumor suppressor gene JST |
| MIR375 | microRNA 375 |
| MIR378A | microRNA 378a |
| MIR383 | microRNA 383 |
| MIR422A | microRNA 422a |
| MIR424 | microRNA 424 |
| MIR449A | microRNA 449a |
| MIR18B | microRNA 18b |
| MIR329-1 | microRNA 329-1 |
| MIR451A | microRNA 451a |
| MIR409 | microRNA 409 |
| MIR410 | microRNA 410 |
| MIR490 | microRNA 490 |
| MIR511 | microRNA 511 |
| MIR493 | microRNA 493 |
| MIR494 | microRNA 494 |
| MIR495 | microRNA 495 |
| MIR193B | microRNA 193b |
| MIR497 | microRNA 497 |
| MIR520B | microRNA 520b |
| MIR520C | microRNA 520c |
| MIR517A | microRNA 517a |
| MIR519D | microRNA 519d |
| MIR502 | microRNA 502 |
| MIR503 | microRNA 503 |
| MIR504 | microRNA 504 |
| MIR505 | microRNA 505 |
| MIR508 | microRNA 508 |
| MIR483 | microRNA 483 |
| MIR486-1 | microRNA 486-1 |
| ST13P21 | suppression of tumorigenicity 13 (colon carcinoma) (Hsp70 interacting protein) |

| | |
|---|---|
| | pseudogene 21 |
| ST13P18 | suppression of tumorigenicity 13 (colon carcinoma) (Hsp70 interacting protein) pseudogene 18 |
| TSG1 | tumor suppressor TSG1 |
| RASSF10 | Ras association domain family member 10 |
| CCDC154 | coiled-coil domain containing 154 |
| TSSC2 | tumor suppressing subtransferable candidate 2 pseudogene |
| CHES1L1 | checkpoint suppressor 1-like 1 |
| MIR487B | microRNA 487b |
| MIR449B | microRNA 449b |
| MIR551A | microRNA 551a |
| MIR574 | microRNA 574 |
| MIR615 | microRNA 615 |
| MIR636 | microRNA 636 |
| ST13P11 | suppression of tumorigenicity 13 (colon carcinoma) (Hsp70 interacting protein) pseudogene 11 |
| ST13P20 | suppression of tumorigenicity 13 (colon carcinoma) (Hsp70 interacting protein) pseudogene 20 |
| ST13P1 | suppression of tumorigenicity 13 (colon carcinoma) (Hsp70 interacting protein) pseudogene 1 |
| TDRG1 | testis development related 1 (non-protein coding) |
| VTRNA2-1 | vault RNA 2-1 |
| MIR888 | microRNA 888 |
| MIR941-1 | microRNA 941-1 |
| MIR708 | microRNA 708 |
| MIR509-3 | microRNA 509-3 |
| MIR874 | microRNA 874 |
| ST13P12 | suppression of tumorigenicity 13 (colon carcinoma) (Hsp70 interacting protein) pseudogene 12 |
| ST13P19 | suppression of tumorigenicity 13 (colon carcinoma) (Hsp70 interacting protein) pseudogene 19 |
| FOXO6 | forkhead box O6 |
| MIR1247 | microRNA 1247 |
| MIR1297 | microRNA 1297 |
| MIR1291 | microRNA 1291 |
| MIR1226 | microRNA 1226 |
| ST13P14 | suppression of tumorigenicity 13 (colon carcinoma) (Hsp70 interacting protein) pseudogene 14 |
| ST13P15 | suppression of tumorigenicity 13 (colon carcinoma) (Hsp70 interacting protein) pseudogene 15 |
| ST13P16 | suppression of tumorigenicity 13 (colon carcinoma) (Hsp70 interacting protein) pseudogene 16 |
| ADAMTS9-AS2 | ADAMTS9 antisense RNA 2 |
| ST13P22 | suppression of tumorigenicity 13 (colon carcinoma) (Hsp70 interacting protein) pseudogene 22 |
| PTCSC3 | papillary thyroid carcinoma susceptibility candidate 3 (non-protein coding) |
| PANO1 | proapoptotic nucleolar protein 1 |
| TP53COR1 | tumor protein p53 pathway corepressor 1 (non-protein coding) |
| HOTS | H19 opposite tumor suppressor |

Table S3: LK, LY, ME, GL - OG, TS

| | Proteins | OG | TS |
|---|---|---|---|
| BCR-ABL1 | xxxxxxxxxx | | |
| RUNX1-RUNX1T1 | HDAC1 | | √ |
| | BRCA1 | | √ |

| | | | |
|---|---|---|---|
| | KMT2A | √ | |
| | SMARCC1 | | √ |
| | HDAC2 | | √ |
| | CREBBP | | √ |
| | CTBP1 | | √ |
| | EP300 | | √ |
| | SMARCA4 | √ | √ |
| | NCOR2 | √ | |
| | NCOR1 | √ | |
| | xxxxx | | |
| | VDR | | √ |
| | SMARCA4 | √ | √ |
| | xxxxxxxxxx | | |
| KMT2A-MLLT10 | SMARCC2 | | |
| | KMT2A | √ | |
| | HDAC2 | | √ |
| | CHD3 | √ | |
| | SMARCC1 | | √ |
| | POLR2A | | |
| | SMARCA2 | | √ |
| | CREBBP | | √ |
| | SIN3A | √ | |
| | xxxxx | | |
| | CTBP1 | | √ |
| | KMT2A | √ | |
| | CREBBP | | √ |
| | xxxxxxxxxx | | |
| IGH-BCL2 | TP53 | | √ |
| | CASP3 | | |
| | PARP1 | | √ |
| | CASP8 | | √ |
| | HIF1A | | √ |
| | BCL2 | √ | |
| | xxxxx | | |
| | BAG3 | √ | |
| | PARP1 | | √ |
| | BCL2 | √ | |
| | xxxxxxxxxx | | |
| KMT2A-AFF1 | SMARCC2 | | |
| | KMT2A | √ | |
| | HDAC2 | | √ |
| | CHD3 | √ | |
| | SMARCC1 | | √ |
| | POLR2A | | |
| | SMARCA2 | √ | |
| | CREBBP | | √ |
| | SIN3A | √ | |
| | xxxxx | | |
| | CTBP1 | | √ |
| | KMT2A | √ | |
| | CREBBP | | √ |
| | xxxxxxxxxx | | |
| PICALM-MLLT10 | FN1 | | √ |
| | EEF1A1 | | √ |

| | | | |
|---|---|---|---|
| | EGFR | √ | |
| | NTRK1 | √ | |
| | PLCG1 | √ | |
| | DNM2 | | |
| | ILVBL | | |
| | PICALM | √ | |
| | xxxxx | | |
| | HNRNPD | √ | |
| | FUS | √ | √ |
| | FN1 | | √ |
| | DDX1 | | √ |
| | xxxxx | | |
| | SEC24D | | |
| | PICALM | √ | |
| | NTRK1 | √ | |
| | SEC24C | | |
| | xxxxxxxxxx | | |
| PML-RARA | NCOA2 | √ | |
| | NR3C1 | | |
| | NR4A1 | | √ |
| | KAT2B | | |
| | RXRA | √ | |
| | PPARG | √ | √ |
| | TP53 | | √ |
| | MDM2 | √ | |
| | EP300 | | √ |
| | SMARCA4 | √ | √ |
| | RELA | √ | |
| | RARA | √ | |
| | NCOA3 | √ | |
| | NCOA1 | √ | |
| | STAT3 | √ | √ |
| | ARNT | √ | |
| | NPAS2 | | √ |
| | PARP1 | | √ |
| | NFKB1 | √ | √ |
| | TRIP4 | | |
| | CREBBP | | √ |
| | xxxxx | | |
| | NFKB1 | √ | √ |
| | EP300 | | √ |
| | PARP1 | | √ |
| | xxxxxxxxxx | | |
| KMT2A-MLLT3 | SMARCC2 | | |
| | KMT2A | √ | |
| | HDAC2 | | √ |
| | CHD3 | √ | |
| | SMARCC1 | | √ |
| | POLR2A | | |
| | SMARCA2 | √ | |
| | CREBBP | | √ |
| | SIN3A | √ | |
| | xxxxx | | |
| | CTBP1 | | √ |

| | | | |
|---|---|---|---|
| | KMT2A | √ | |
| | CREBBP | | √ |
| | xxxxxxxxxx | | |
| KMT2A-AFDN | SMARCC2 | | |
| | KMT2A | √ | |
| | HDAC2 | | √ |
| | CHD3 | √ | |
| | SMARCC1 | | √ |
| | POLR2A | | |
| | SMARCA2 | √ | |
| | CREBBP | | √ |
| | SIN3A | √ | |
| | xxxxx | | |
| | CTBP1 | | √ |
| | KMT2A | √ | |
| | CREBBP | | √ |
| | xxxxxxxxxx | | |
| CBFB-MYH11 | ACTA2 | | |
| | MYH11 | √ | |
| | MYO1E | | |
| | RPA1 | | √ |
| | RPA2 | √ | |
| | ACTB | √ | |
| | xxxxx | | |
| | RPA1 | | √ |
| | ELAVL1 | √ | |
| | RPA2 | √ | |
| | xxxxxxxxxx | | |
| IGH-MYC | | | |
| | xxxxx | | |
| | xxxxxxxxxx | | |
| NUP98-DDX10 | SIRT7 | | |
| | DDX10 | √ | |
| | APP | | |
| | DDX56 | | |
| | NTRK1 | √ | |
| | DDX54 | | |
| | PUM3 | | |
| | PWP1 | | |
| | CSNK2A1 | √ | |
| | xxxxx | | |
| | HDAC1 | | √ |
| | CTNNB1 | √ | |
| | MAPK8 | √ | |
| | EP300 | | √ |
| | CREBBP | | √ |
| | xxxxx | | |
| | PUM3 | | |
| | NXF1 | | |
| | EED | | √ |
| | KPNB1 | | |
| | HNRNPUL1 | | |
| | xxxxx | | |

| | | | |
|---|---|---|---|
| | HDAC1 | | √ |
| | CREBBP | | √ |
| | APC | √ | √ |
| | CSNK2A1 | √ | |
| | xxxxx | | |
| | CTNNB1 | √ | |
| | APC | √ | √ |
| | CREBBP | | √ |
| | xxxxx | | |
| | USP7 | √ | |
| | NTRK1 | √ | |
| | CDC37 | | |
| | xxxxxxxxxx | | |
| PCM1-JAK2 | ERBB2 | √ | |
| | ERBB3 | √ | |
| | VAV1 | √ | |
| | TEC | | |
| | EGFR | √ | |
| | INSR | | |
| | PLCG1 | √ | |
| | JAK2 | √ | |
| | xxxxx | | |
| | STAT5A | √ | √ |
| | JAK2 | √ | |
| | INSR | | |
| | xxxxxxxxxx | | |
| KMT2A-SEPT9 | SMARCC2 | | |
| | KMT2A | √ | |
| | HDAC2 | | √ |
| | CHD3 | √ | |
| | SMARCC1 | | √ |
| | POLR2A | | |
| | SMARCA2 | √ | |
| | CREBBP | | √ |
| | SIN3A | √ | |
| | xxxxx | | |
| | CTBP1 | | √ |
| | KMT2A | √ | |
| | CREBBP | | √ |
| | xxxxxxxxxx | | |
| FUS-ERG | RPA1 | | √ |
| | SF3B2 | | |
| | PRKDC | | |
| | PRPF8 | √ | |
| | SF3A2 | | |
| | DHX15 | | |
| | RPA2 | √ | |
| | CUL3 | | |
| | xxxxx | | |
| | ABL1 | √ | |
| | PARP1 | | √ |
| | PRKDC | | |
| | xxxxxxxxxx | | |
| NUP98-HOXA9 | HDAC1 | | √ |

| | | | |
|---|---|---|---|
| | TP53 | | √ |
| | SMAD4 | √ | √ |
| | CTNNB1 | √ | |
| | CSNK2A1 | √ | |
| | MAPK8 | √ | |
| | EP300 | | √ |
| | CREBBP | | √ |
| | xxxxx | | |
| | CTNNB1 | √ | |
| | APC | √ | √ |
| | CREBBP | | √ |
| | xxxxxxxxxx | | |
| ETV6-ABL1 | ERBB2 | √ | |
| | CBLB | √ | |
| | UBASH3B | | |
| | SOS1 | √ | |
| | ERBB4 | √ | √ |
| | SRC | √ | |
| | VAV1 | √ | |
| | EGFR | √ | |
| | CBL | √ | √ |
| | PLCG1 | √ | |
| | PIK3R2 | √ | |
| | PIK3R1 | √ | |
| | ABL1 | √ | |
| | xxxxx | | |
| | ABL1 | √ | |
| | ABL2 | √ | |
| | JAK1 | √ | |
| | xxxxxxxxxx | | |
| SET-NUP214 | NXF1 | | |
| | FAF1 | | |
| | SUPT5H | | |
| | GART | | |
| | CUL2 | | √ |
| | CUL3 | | |
| | xxxxx | | |
| | NXF1 | | |
| | CUL2 | | √ |
| | RANBP2 | √ | |
| | xxxxxxxxxx | | |
| MNX1-ETV6 | HDAC3 | √ | √ |
| | ETV6 | √ | √ |
| | HDAC9 | | |
| | PIN1 | | √ |
| | NCOR1 | √ | |
| | SIN3A | √ | |
| | xxxxx | | |
| | L3MBTL1 | | |
| | ETV7 | | |
| | ETV6 | √ | √ |
| | xxxxxxxxxx | | |

| | | | |
|---|---|---|---|
| KMT2A-MLLT1 | SMARCC2 | | |
| | KMT2A | √ | |
| | HDAC2 | | √ |
| | CHD3 | √ | |
| | SMARCC1 | | √ |
| | POLR2A | | |
| | SMARCA2 | √ | |
| | CREBBP | | √ |
| | SIN3A | √ | |
| | xxxxx | | |
| | CTBP1 | | √ |
| | KMT2A | √ | |
| | CREBBP | | √ |
| | xxxxxxxxxx | | |
| KMT2A-MLLT6 | SMARCC2 | | |
| | KMT2A | √ | |
| | HDAC2 | | √ |
| | CHD3 | √ | |
| | SMARCC1 | | √ |
| | POLR2A | | |
| | SMARCA2 | √ | |
| | CREBBP | | √ |
| | SIN3A | √ | |
| | xxxxx | | |
| | CTBP1 | | √ |
| | KMT2A | √ | |
| | CREBBP | | √ |
| | xxxxxxxxxx | | |
| ETV6-ACSL6 | HDAC3 | √ | √ |
| | ETV6 | √ | √ |
| | HDAC9 | | |
| | PIN1 | | √ |
| | NCOR1 | √ | |
| | SIN3A | √ | |
| | xxxxx | | |
| | L3MBTL1 | | |
| | ETV7 | | |
| | ETV6 | √ | √ |
| | xxxxxxxxxx | | |
| ETV6-MECOM | HDAC1 | | √ |
| | UBE2I | | |
| | HDAC3 | √ | √ |
| | EHMT2 | | |
| | ELAVL1 | √ | |
| | MECOM | √ | |
| | SMAD1 | | |
| | SMAD2 | √ | √ |

| | | | |
|---|---|---|---|
| | SMAD3 | √ | |
| | KAT2B | | |
| | SUV39H1 | | |
| | NCOR1 | √ | |
| | CTBP1 | | √ |
| | CREBBP | | √ |
| | xxxxx | | |
| | MECOM | √ | |
| | SUV39H1 | | |
| | HDAC4 | √ | |
| | xxxxxxxxxx | | |
| KMT2A-EPS15 | SMARCC2 | | |
| | KMT2A | √ | |
| | HDAC2 | | √ |
| | CHD3 | √ | |
| | SMARCC1 | | √ |
| | POLR2A | | |
| | SMARCA2 | √ | |
| | CREBBP | | √ |
| | SIN3A | √ | |
| | xxxxx | | |
| | CTBP1 | | √ |
| | KMT2A | √ | |
| | CREBBP | | √ |
| | xxxxxxxxxx | | |
| KMT2A-GAS7 | SMARCC2 | | |
| | KMT2A | √ | |
| | HDAC2 | | √ |
| | CHD3 | √ | |
| | SMARCC1 | | √ |
| | POLR2A | | |
| | SMARCA2 | √ | |
| | CREBBP | | √ |
| | SIN3A | √ | |
| | xxxxx | | |
| | CTBP1 | | √ |
| | KMT2A | √ | |
| | CREBBP | | √ |
| | xxxxxxxxxx | | |
| KMT2A-ABL1 | SMARCC2 | | |
| | KMT2A | √ | |
| | SMARCC1 | | √ |
| | CHD3 | √ | |
| | POLR2A | | |
| | SMARCA2 | √ | |
| | CREBBP | | √ |
| | xxxxx | | |
| | ABL1 | √ | |
| | CBLB | √ | |
| | CBL | √ | √ |
| | xxxxxxxxxx | | |
| KMT2A-MLLT11 | SMARCC2 | | |
| | KMT2A | √ | |
| | HDAC2 | | √ |

| | | | |
|---|---|---|---|
| | CHD3 | √ | |
| | SMARCC1 | | √ |
| | POLR2A | | |
| | SMARCA2 | √ | |
| | CREBBP | | √ |
| | SIN3A | √ | |
| | xxxxx | | |
| | CTBP1 | | √ |
| | KMT2A | √ | |
| | CREBBP | | √ |
| | xxxxxxxxxx | | |
| MN1-ETV6 | HDAC3 | √ | √ |
| | ETV6 | √ | √ |
| | HDAC9 | | |
| | PIN1 | | √ |
| | EP300 | | √ |
| | NCOR1 | √ | |
| | SIN3A | √ | |
| | xxxxx | | |
| | HDAC3 | √ | √ |
| | EP300 | | √ |
| | KAT5 | √ | √ |
| | xxxxxxxxxx | | |
| KMT2A-MAML2 | SMARCC2 | | |
| | KMT2A | √ | |
| | SMARCC1 | | √ |
| | CHD3 | √ | |
| | SMARCA2 | √ | |
| | CREBBP | | √ |
| | xxxxx | | |
| | SMARCA2 | √ | |
| | EP300 | | √ |
| | CREBBP | | √ |
| | xxxxxxxxxx | | |
| KMT2A-FOXO4 | SMARCC2 | | |
| | KMT2A | √ | |
| | HDAC2 | | √ |
| | CHD3 | √ | |
| | SMARCC1 | | √ |
| | POLR2A | | |
| | SMARCA2 | √ | |
| | CREBBP | | √ |
| | SIN3A | √ | |
| | xxxxx | | |
| | CTBP1 | | √ |
| | KMT2A | √ | |
| | CREBBP | | √ |
| | xxxxxxxxxx | | |
| DEK-NUP214 | ESR1 | √ | √ |
| | KAT2B | | |
| | SMAD2 | √ | √ |
| | SMAD3 | √ | |
| | CDK2 | √ | √ |
| | DEK | √ | |

| | | | |
|---|---|---|---|
| | EP300 | | √ |
| | CREBBP | | √ |
| | xxxxx | | |
| | NXF1 | | |
| | DHX15 | | |
| | CUL2 | | √ |
| | CUL3 | | |
| | xxxxxxxxxx | | |
| RUNX1-CBFA2T3 | KMT2A | √ | |
| | SMARCC1 | | √ |
| | SMARCA4 | √ | √ |
| | CREBBP | | √ |
| | xxxxx | | |
| | EP300 | | √ |
| | CREBBP | | √ |
| | xxxxxxxxxx | | |
| NUP98-PSIP1 | HDAC1 | | √ |
| | KMT2A | √ | |
| | ESR1 | √ | √ |
| | CTNNB1 | √ | |
| | EP300 | | √ |
| | CREBBP | | √ |
| | xxxxx | | |
| | NXF1 | | |
| | EIF4A3 | | |
| | SON | | |
| | xxxxxxxxxx | | |
| NUP98-HOXC13 | HDAC1 | | √ |
| | CTNNB1 | √ | |
| | MAPK8 | √ | |
| | EP300 | | √ |
| | CREBBP | | √ |
| | xxxxx | | |
| | NXF1 | | |
| | HNRNPAB | | |
| | EED | | √ |
| | KPNB1 | | |
| | HNRNPUL1 | | |
| | xxxxx | | |
| | HDAC1 | | √ |
| | CREBBP | | √ |
| | APC | √ | √ |
| | CSNK2A1 | √ | |
| | xxxxx | | |
| | CTNNB1 | √ | |
| | APC | √ | √ |
| | CREBBP | | √ |
| | xxxxxxxxxx | | |
| NUP98-HOXC11 | HDAC1 | | √ |
| | STAT3 | √ | √ |
| | SP1 | | |
| | SMAD3 | √ | |
| | CTNNB1 | √ | |
| | MAPK8 | √ | |

| | | | |
|---|---|---|---|
| | EP300 | | √ |
| | CREBBP | | √ |
| | xxxxx | | |
| | CTNNB1 | √ | |
| | APC | √ | √ |
| | CREBBP | | √ |
| | xxxxxxxxxx | | |
| PAX5-ETV6 | UBE2I | | |
| | HDAC3 | √ | √ |
| | TBP | | |
| | KAT5 | √ | √ |
| | RB1 | √ | √ |
| | PAX5 | √ | √ |
| | RUNX1 | √ | √ |
| | PIN1 | | √ |
| | EP300 | | √ |
| | NCOR1 | √ | |
| | xxxxx | | |
| | MAPK1 | √ | |
| | EP300 | | √ |
| | HDAC6 | √ | |
| | xxxxxxxxxx | | |
| NUP98-HOXA11 | HDAC1 | | √ |
| | HDAC2 | | √ |
| | YY1 | | |
| | CTNNB1 | √ | |
| | CSNK2A1 | √ | |
| | MAPK8 | √ | |
| | EP300 | | √ |
| | CREBBP | | √ |
| | xxxxx | | |
| | CTNNB1 | √ | |
| | APC | √ | √ |
| | CREBBP | | √ |
| | xxxxxxxxxx | | |
| BCR-PDGFRA | TGFBR2 | √ | √ |
| | PDGFRA | √ | |
| | SHC1 | √ | |
| | CRKL | √ | |
| | EGFR | √ | |
| | PLCG1 | √ | |
| | FES | √ | |
| | CRK | √ | |
| | ABL1 | √ | |
| | HCK | √ | |
| | PTPN6 | √ | √ |
| | BCR | √ | √ |
| | UBASH3B | | |
| | INPP5D | | |
| | SOS1 | √ | |
| | GRB2 | √ | |
| | NTRK1 | √ | |
| | CBL | √ | √ |
| | KIT | √ | |

| | | | |
|---|---|---|---|
| | DOK1 | | √ |
| | PIK3R2 | √ | |
| | PIK3R1 | √ | |
| | xxxxx | | |
| | ABL1 | √ | |
| | TP53 | | √ |
| | RB1 | √ | √ |
| | BCR | √ | √ |
| | xxxxxxxxxx | | |
| BCR-FGFR1 | SRC | √ | |
| | ITK | √ | |
| | SOS1 | √ | |
| | ERBB3 | √ | |
| | VAV1 | √ | |
| | CBL | √ | √ |
| | PLCG1 | √ | |
| | ABL1 | √ | |
| | xxxxx | | |
| | ABL1 | √ | |
| | HCK | √ | |
| | BCR | √ | √ |
| | CBL | √ | √ |
| | xxxxxxxxxx | | |
| NPM1-RARA | | | |
| KMT2A-CBL | | | |
| IGH-BCL6 | HDAC1 | | √ |
| | TP53 | | √ |
| | CTBP1 | | √ |
| | EP300 | | √ |
| | NCOR2 | √ | |
| | CREBBP | | √ |
| | xxxxx | | |
| | HDAC2 | | √ |
| | SMARCA4 | √ | √ |
| | CREBBP | | √ |
| | xxxxxxxxxx | | |
| LCP1-BCL6 | HDAC1 | | √ |
| | TP53 | | √ |
| | CTBP1 | | √ |
| | EP300 | | √ |
| | NCOR2 | √ | |
| | CREBBP | | √ |
| | xxxxx | | |
| | HDAC2 | | √ |
| | SMARCA4 | √ | √ |
| | CREBBP | | √ |
| | xxxxxxxxxx | | |
| CREBBP-KAT6A | SMARCC2 | | |
| | KMT2A | √ | |
| | HDAC2 | | √ |
| | CHD3 | √ | |
| | SMARCC1 | | √ |
| | POLR2A | | |

| | | | |
|---|---|---|---|
| | SMARCA2 | √ | |
| | CREBBP | | √ |
| | SIN3A | √ | |
| | xxxxx | | |
| | CTBP1 | | √ |
| | KMT2A | √ | |
| | CREBBP | | √ |
| | xxxxxxxxxx | | |
| KMT2A-ARHGAP26 | SMARCC2 | | |
| | KMT2A | √ | |
| | HDAC2 | | √ |
| | CHD3 | √ | |
| | SMARCC1 | | √ |
| | POLR2A | | |
| | SMARCA2 | √ | |
| | CREBBP | | √ |
| | SIN3A | √ | |
| | xxxxx | | |
| | CTBP1 | | √ |
| | KMT2A | √ | |
| | CREBBP | | √ |
| | xxxxxxxxxx | | |
| FOXO3-KMT2A | SMARCC2 | | |
| | KMT2A | √ | |
| | SMARCC1 | | √ |
| | CHD3 | √ | |
| | SMARCA2 | √ | |
| | CREBBP | | √ |
| | xxxxx | | |
| | SMARCA2 | √ | |
| | EP300 | | √ |
| | CREBBP | | √ |
| | xxxxxxxxxx | | |
| KMT2A-DCPS | SMARCC2 | | |
| | KMT2A | √ | |
| | HDAC2 | | √ |
| | CHD3 | √ | |
| | SMARCC1 | | √ |
| | POLR2A | | |
| | SMARCA2 | √ | |
| | CREBBP | | √ |
| | SIN3A | √ | |
| | xxxxx | | |
| | CTBP1 | | √ |
| | KMT2A | √ | |
| | CREBBP | | √ |
| | xxxxxxxxxx | | |
| KMT2A-EP300 | | | |
| | xxxxxxxxxx | | |
| IGH-CEBPE | UBE2I | | |
| | BATF | √ | |
| | DDIT3 | √ | |
| | CEBPG | | |
| | CEBPE | | |

| | | | |
|---|---|---|---|
| | FOS | √ | |
| | JUN | √ | |
| | STAT6 | √ | |
| | RB1 | √ | √ |
| | PIAS1 | | √ |
| | FOSL1 | √ | |
| | BATF3 | | |
| | BATF2 | | √ |
| | ATF4 | √ | |
| | MYB | √ | |
| | ATF3 | √ | √ |
| | xxxxxxxxxx | | |
| HSP90AA1-BCL6 | | | |

Table S4: SC - OG, TS

| | Proteins | OG | TS |
|---|---|---|---|
| ASPSCR1-TFE3 | | | |
| | xxxxxxxxxxxxxxxxxx | | |
| ASTN2-CNOT2 | CNOT6L | | |
| | AURKA | √ | |
| | CNOT8 | | |
| | TNRC6C | | |
| | TNRC6B | | |
| | CNOT3 | | √ |
| | CNOT2 | | |
| | CNOT1 | | |
| | CNOT7 | | |
| | AGO2 | | |
| | xxxxx | | |
| | HDAC3 | √ | √ |
| | GPS2 | √ | |
| | CNOT2 | | |
| | NCOR2 | √ | |
| | NCOR1 | √ | |
| | xxxxxxxxxxxxxxxxxx | | |
| BCOR-ZC3H7B | HDAC3 | √ | √ |
| | HDAC4 | √ | |
| | SP1 | | |
| | CTBP1 | | √ |
| | NACC1 | | |
| | NCOR2 | √ | |
| | xxxxx | | |
| | HDAC1 | | √ |
| | CTBP1 | | √ |
| | NCOR2 | √ | |
| | xxxxxxxxxxxxxxxxxx | | |
| BCOR-CCNB3 | HDAC3 | √ | √ |
| | HDAC4 | √ | |
| | SP1 | | |
| | CTBP1 | | √ |
| | NACC1 | | |
| | NCOR2 | √ | |

| | | | |
|---|---|---|---|
| | HDAC1 | | √ |
| | CTBP1 | | √ |
| | NCOR2 | √ | |
| | xxxxxxxxxxxxxxxxx | | |
| CDX1-IRF2BP2 | ELAVL1 | √ | |
| | NTRK1 | √ | |
| | IRF2BPL | | |
| | IRF2BP2 | √ | |
| | RBM39 | √ | |
| | xxxxxxxxxxxxxxxxx | | |
| CIC-DUX4 | | | |
| | xxxxxxxxxxxxxxxxx | | |
| CREB1-EWSR1 | BRCA1 | | √ |
| | EPAS1 | | √ |
| | MYOD1 | √ | |
| | ESR1 | √ | √ |
| | JUN | √ | |
| | POLR2A | | |
| | EP300 | | √ |
| | SMARCA4 | √ | √ |
| | EWSR1 | √ | |
| | CREBBP | | √ |
| | xxxxx | | |
| | NR3C1 | | |
| | EP300 | | √ |
| | SMARCA4 | √ | √ |
| | CREBBP | | √ |
| | xxxxx | | |
| | NONO | √ | |
| | SMARCA4 | √ | √ |
| | CUL3 | | |
| | xxxxxxxxxxxxxxxxx | | |
| CTDSP2-FAM19A2 | SETD1A | | |
| | POLR2A | | |
| | INTS6 | √ | √ |
| | CTDSP1 | | |
| | CTDSP2 | | |
| | xxxxxxxxxxxxxxxxx | | |
| CXorf67-MBTD1 | | | |
| | xxxxxxxxxxxxxxxxx | | |
| EPC1-PHF1 | HDAC1 | | √ |
| | DHX9 | √ | |
| | TP53 | | √ |
| | ELAVL1 | √ | |
| | E2F6 | √ | |
| | RBBP7 | | √ |
| | RBBP4 | √ | |
| | EZH1 | | √ |
| | EZH2 | | √ |
| | EED | | √ |
| | XRCC6 | | √ |
| | XRCC5 | | √ |
| | PHF1 | | |
| | xxxxx | | |

| | | | |
|---|---|---|---|
| | HDAC1 | | √ |
| | TP53 | | √ |
| | YEATS4 | | |
| | TRIM27 | √ | |
| | KAT5 | √ | √ |
| | TRIM23 | | |
| | HIST1H2BA | | |
| | DMAP1 | | |
| | MYC | √ | |
| | XRCC6 | | √ |
| | MORF4L1 | | |
| | ING3 | √ | √ |
| | xxxxxxxxxxxxxxxxxx | | |
| ERG-EWSR1 | TP53 | | √ |
| | ESR1 | √ | √ |
| | PARP1 | | √ |
| | EP300 | | √ |
| | CREBBP | | √ |
| | XRCC6 | | √ |
| | xxxxx | | |
| | EPAS1 | | √ |
| | EP300 | | √ |
| | CREBBP | | √ |
| | xxxxxxxxxxxxxxxxxx | | |
| ETV6-NTRK3 | PDGFRB | √ | |
| | SHC1 | √ | |
| | CRKL | √ | |
| | GAB2 | √ | |
| | NTRK1 | √ | |
| | PLCG1 | √ | |
| | GRB2 | √ | |
| | xxxxx | | |
| | HDAC3 | √ | √ |
| | PIN1 | | √ |
| | ETV6 | √ | √ |
| | xxxxxxxxxxxxxxxxxx | | |
| EWSR1-ATF1 | PDGFRB | √ | |
| | SHC1 | √ | |
| | CRKL | √ | |
| | GAB2 | √ | |
| | NTRK1 | √ | |
| | PLCG1 | √ | |
| | GRB2 | √ | |
| | xxxxx | | |
| | HDAC3 | √ | √ |
| | PIN1 | | √ |
| | ETV6 | √ | √ |
| | xxxxxxxxxxxxxxxxxx | | |
| EWSR1-FLI1 | BRCA1 | | √ |
| | POLR2A | | |
| | ESR1 | √ | √ |
| | EP300 | | √ |
| | KAT2B | | |
| | CREBBP | | √ |

| | | | |
|---|---|---|---|
| | xxxxx | | |
| | EPAS1 | | √ |
| | EP300 | | √ |
| | CREBBP | | √ |
| | xxxxxxxxxxxxxxxxxx | | |
| EWSR1-NR4A3 | TSG101 | √ | √ |
| | DHX9 | √ | |
| | HDAC3 | √ | √ |
| | TRIM28 | √ | |
| | FUS | √ | √ |
| | RAD23A | √ | |
| | JUN | √ | |
| | PRMT1 | | |
| | ILK | | √ |
| | BMI1 | √ | |
| | ELK1 | √ | |
| | POLR2A | | |
| | CHERP | | |
| | ATXN3 | | |
| | EP300 | | √ |
| | IRF3 | | √ |
| | EPAS1 | | √ |
| | TP53 | | √ |
| | ESR1 | √ | √ |
| | NONO | √ | |
| | RPA1 | | √ |
| | NTRK1 | √ | |
| | RPA2 | √ | |
| | CUL4A | | |
| | CUL4B | √ | |
| | FASN | | |
| | CREBBP | | √ |
| | HBP1 | √ | √ |
| | CUL5 | | √ |
| | HLTF | √ | √ |
| | EWSR1 | √ | |
| | YBX1 | √ | |
| | CUL1 | √ | √ |
| | CUL2 | | √ |
| | CUL3 | | |
| | xxxxx | | |
| | HDAC2 | | √ |
| | CREBBP | | √ |
| | ESR1 | √ | √ |
| | xxxxx | | |
| | NONO | √ | |
| | FXR2 | | |
| | CUL3 | | |
| | xxxxxxxxxxxxxxxxxx | | |
| EWSR1-ETV4 | TSG101 | √ | √ |
| | DHX9 | √ | |
| | HDAC3 | √ | √ |
| | RFWD2 | | √ |
| | FUS | √ | √ |

| | | | |
|---|---|---|---|
| | RAD23A | √ | |
| | JUN | √ | |
| | PRMT1 | | |
| | ILK | | √ |
| | SMAD2 | √ | √ |
| | BMI1 | √ | |
| | ELK1 | √ | |
| | POLR2A | | |
| | CHERP | | |
| | ATXN3 | | |
| | EP300 | | √ |
| | IRF3 | | √ |
| | EPAS1 | | √ |
| | TP53 | | √ |
| | ESR1 | √ | √ |
| | NONO | √ | |
| | RPA1 | | √ |
| | NTRK1 | √ | |
| | RPA2 | √ | |
| | CUL4A | | |
| | CUL4B | √ | |
| | FASN | | |
| | CREBBP | | √ |
| | HBP1 | √ | √ |
| | CUL5 | | √ |
| | HLTF | √ | √ |
| | EWSR1 | √ | |
| | YBX1 | √ | |
| | CUL1 | √ | √ |
| | CUL2 | | √ |
| | CUL3 | | |
| | xxxxx | | |
| | HDAC2 | | √ |
| | CREBBP | | √ |
| | ESR1 | √ | √ |
| | xxxxx | | |
| | NONO | √ | |
| | FXR2 | | |
| | CUL3 | | |
| | xxxxxxxxxxxxxxxxx | | |
| EWSR1-PATZ1 | TSG101 | √ | √ |
| | DHX9 | √ | |
| | HDAC3 | √ | √ |
| | RFWD2 | | √ |
| | FUS | √ | √ |
| | RAD23A | √ | |
| | JUN | √ | |
| | PRMT1 | | |
| | ILK | | √ |
| | SMAD2 | √ | √ |
| | BMI1 | √ | |
| | ELK1 | √ | |
| | POLR2A | | |
| | CHERP | | |

| | | | |
|---|---|---|---|
| | ATXN3 | | |
| | EP300 | | √ |
| | IRF3 | | √ |
| | EPAS1 | | √ |
| | TP53 | | √ |
| | ESR1 | √ | √ |
| | NONO | √ | |
| | RPA1 | | √ |
| | NTRK1 | √ | |
| | RPA2 | √ | |
| | CUL4A | | |
| | CUL4B | √ | |
| | FASN | | |
| | CREBBP | | √ |
| | HBP1 | √ | √ |
| | CUL5 | | √ |
| | HLTF | √ | √ |
| | EWSR1 | √ | |
| | YBX1 | √ | |
| | CUL1 | √ | √ |
| | CUL2 | | √ |
| | CUL3 | | |
| | xxxxx | | |
| | HDAC2 | | √ |
| | CREBBP | | √ |
| | ESR1 | √ | √ |
| | xxxxx | | |
| | NONO | √ | |
| | FXR2 | | |
| | CUL3 | | |
| | xxxxxxxxxxxxxxxxx | | |
| EWSR1-DDIT3 | HDAC1 | | √ |
| | EPAS1 | | √ |
| | HDAC3 | √ | √ |
| | DDIT3 | √ | |
| | CEBPB | √ | |
| | ESR1 | √ | √ |
| | FOS | √ | |
| | JUN | √ | |
| | HBP1 | √ | √ |
| | POLR2A | | |
| | IRF3 | | √ |
| | TP53 | | √ |
| | EP300 | | √ |
| | EWSR1 | √ | |
| | CREBBP | | √ |
| | xxxxx | | |
| | DHX9 | √ | |
| | CUL4A | | |
| | CUL4B | √ | |
| | CUL5 | | √ |
| | CUL1 | √ | √ |
| | CUL2 | | √ |
| | CUL3 | | |

| | | | |
|---|---|---|---|
| | xxxxxxxxxxxxxxxxxx | | |
| EWSR1-POU5F1 | IRF3 | | √ |
| | EPAS1 | | √ |
| | HDAC3 | √ | √ |
| | TP53 | | √ |
| | ESR1 | √ | √ |
| | JUN | √ | |
| | HBP1 | √ | √ |
| | ETS2 | | |
| | CTNNB1 | √ | |
| | POLR2A | | |
| | EP300 | | √ |
| | EWSR1 | √ | |
| | CREBBP | | √ |
| | xxxxx | | |
| | NONO | √ | |
| | CUL2 | | √ |
| | CUL3 | | |
| | xxxxxxxxxxxxxxxxxx | | |
| EWSR1-SP3 | HDAC1 | | √ |
| | EPAS1 | | √ |
| | HDAC3 | √ | √ |
| | CEBPB | √ | |
| | ESR1 | √ | √ |
| | JUN | √ | |
| | HBP1 | √ | √ |
| | POLR2A | | |
| | IRF3 | | √ |
| | TP53 | | √ |
| | RELA | √ | |
| | EP300 | | √ |
| | EWSR1 | √ | |
| | CREBBP | | √ |
| | xxxxx | | |
| | DHX9 | √ | |
| | CUL4A | | |
| | CUL4B | √ | |
| | CUL5 | | √ |
| | CUL1 | √ | √ |
| | CUL2 | | √ |
| | CUL3 | | |
| | xxxxxxxxxxxxxxxxxx | | |
| FOXO4-CIC | XPO1 | | |
| | CTNNB1 | √ | |
| | VDR | | √ |
| | ESR1 | √ | √ |
| | SMAD4 | √ | √ |
| | SMAD3 | √ | |
| | SFN | | √ |
| | AKT1 | √ | |
| | FOXO4 | √ | √ |
| | MDM2 | √ | |
| | NLK | | |
| | CREBBP | | √ |

| | | | |
|---|---|---|---|
| | xxxxxxxxxxxxxxxxx | | |
| FUS-ERG | RPA1 | | √ |
| | SF3B2 | | |
| | PRKDC | | |
| | PRPF8 | √ | |
| | SF3A2 | | |
| | DHX15 | | |
| | RPA2 | √ | |
| | CUL3 | | |
| | xxxxx | | |
| | ABL1 | √ | |
| | PARP1 | | √ |
| | PRKDC | | |
| | xxxxxxxxxxxxxxxxx | | |
| FUS-CREB3L1 | NONO | √ | |
| | CUL4A | | |
| | CUL4B | √ | |
| | DHX15 | | |
| | CUL5 | | √ |
| | CUL1 | √ | √ |
| | CUL2 | | √ |
| | CUL3 | | |
| | xxxxx | | |
| | VCP | | |
| | FBXW11 | | |
| | CUL1 | √ | √ |
| | xxxxxxxxxxxxxxxxx | | |
| FUS-DDIT3 | HDAC1 | | √ |
| | DDX17 | | |
| | EPAS1 | | √ |
| | RELA | √ | |
| | ESR1 | √ | √ |
| | JUN | √ | |
| | TP73 | √ | √ |
| | EWSR1 | √ | |
| | CTNNB1 | √ | |
| | DDX5 | √ | |
| | CDK2 | √ | √ |
| | DDIT3 | √ | |
| | MDM2 | √ | |
| | EP300 | | √ |
| | TRIP4 | | |
| | CREBBP | | √ |
| | xxxxx | | |
| | VCP | | |
| | FBXW11 | | |
| | CUL1 | √ | √ |
| | xxxxxxxxxxxxxxxxx | | |
| FUS-ATF1 | HDAC1 | | √ |
| | DDX17 | | |
| | EPAS1 | | √ |
| | RELA | √ | |
| | ESR1 | √ | √ |
| | JUN | √ | |

| | | | |
|---|---|---|---|
| | TP73 | √ | √ |
| | EWSR1 | √ | |
| | CTNNB1 | √ | |
| | DDX5 | √ | |
| | CDK2 | √ | √ |
| | DDIT3 | √ | |
| | MDM2 | √ | |
| | EP300 | | √ |
| | TRIP4 | | |
| | CREBBP | | √ |
| | xxxxx | | |
| | VCP | | |
| | FBXW11 | | |
| | CUL1 | √ | √ |
| | xxxxxxxxxxxxxxxxxx | | |
| FUS-CREB3L2 | NONO | √ | |
| | CUL4A | | |
| | CUL4B | √ | |
| | DHX15 | | |
| | CUL5 | | √ |
| | CUL1 | √ | √ |
| | CUL2 | | √ |
| | CUL3 | | |
| | xxxxx | | |
| | VCP | | |
| | FBXW11 | | |
| | CUL1 | √ | √ |
| | xxxxxxxxxxxxxxxxxx | | |
| HEY1-NCOA2 | BRCA1 | | √ |
| | RARA | √ | |
| | NR3C1 | | |
| | VDR | | √ |
| | STAT6 | √ | |
| | HNF4A | | √ |
| | PRMT1 | | |
| | CARM1 | √ | |
| | RXRA | √ | |
| | PPARG | √ | √ |
| | PPARD | √ | |
| | AR | | |
| | PPARA | | √ |
| | ESR2 | √ | √ |
| | EP300 | | √ |
| | NCOA2 | √ | |
| | NCOA3 | √ | |
| | NCOA1 | √ | |
| | TP53 | | √ |
| | ESR1 | √ | √ |
| | AHR | | √ |
| | ARNT | | √ |
| | THRB | | √ |
| | THRA | √ | √ |
| | NR1I3 | | |
| | NR1I2 | | √ |

| | | | |
|---|---|---|---|
| | PGR | | √ |
| | CREBBP | | √ |
| | PIAS3 | √ | |
| | xxxxx | | |
| | NCOA2 | √ | |
| | NCOA1 | √ | |
| | UBR5 | | |
| | xxxxxxxxxxxxxxxxxx | | |
| IRX2-TERT | YWHAZ | | |
| | AKT1 | √ | |
| | RPS6KB1 | | |
| | ENO1 | √ | |
| | MTOR | √ | |
| | MDM2 | √ | |
| | TERT | √ | |
| | XRCC6 | | √ |
| | xxxxx | | |
| | TERF1 | | |
| | STUB1 | | √ |
| | TERT | √ | |
| | POT1 | | |
| | xxxxx | | |
| | TERT | √ | |
| | MTOR | √ | |
| | YWHAQ | | |
| | RUVBL2 | √ | |
| | xxxxx | | |
| | TPP1 | | |
| | TERT | √ | |
| | POT1 | | |
| | xxxxxxxxxxxxxxxxxx | | |
| JAZF1-SUZ12 | DHX9 | √ | |
| | RBM5 | √ | √ |
| | FBXW11 | | |
| | DDX3X | | √ |
| | NXF1 | | |
| | SF3B4 | | |
| | PRMT1 | | |
| | SF3B1 | √ | |
| | SF3B2 | | |
| | PRPF8 | √ | |
| | EED | | √ |
| | CRNKL1 | | |
| | RNPS1 | | |
| | UBE2I | | |
| | SNRNP200 | | |
| | SRSF7 | | |
| | SNRPD3 | | |
| | RALY | | |
| | DDX5 | √ | |
| | PRPF19 | | |
| | RNF2 | | |
| | SNRPA1 | | √ |
| | SON | | |

| | | | |
|---|---|---|---|
| | EFTUD2 | | |
| | U2AF1 | √ | |
| | SF3A1 | | |
| | EPRS | | |
| | ILF2 | | |
| | EIF4A3 | | |
| | CDC40 | | |
| | ILF3 | √ | |
| | RANBP2 | √ | |
| | xxxxx | | |
| | DNMT3B | | √ |
| | HDAC1 | | √ |
| | HDAC2 | | √ |
| | TRIM28 | √ | |
| | CHD4 | √ | |
| | UHRF1 | | |
| | DNMT1 | √ | |
| | MTA1 | √ | |
| | NR2C2 | | √ |
| | EZH2 | | √ |
| | GATAD2B | | |
| | EED | | √ |
| | CBX5 | | √ |
| | RBBP4 | √ | |
| | CBX3 | | |
| | SETDB1 | | |
| | xxxxx | | |
| | VCP | | |
| | BRCA1 | | √ |
| | MTOR | √ | |
| | NXF1 | | |
| | RUVBL2 | √ | |
| | xxxxx | | |
| | VCP | | |
| | FBXW11 | | |
| | SKP1 | √ | |
| | BTRC | √ | |
| | EZH2 | | √ |
| | xxxxx | | |
| | DHX9 | √ | |
| | ADAR | | |
| | EZH2 | | √ |
| | FBXW11 | | |
| | xxxxx | | |
| | HDAC2 | | √ |
| | NXF1 | | |
| | JARID2 | | |
| | SETDB1 | | |
| | xxxxx | | |
| | RELA | √ | |
| | FBXW11 | | |
| | BTRC | √ | |
| | xxxxx | | |
| | BTRC | √ | |

| | | | |
|---|---|---|---|
| | CSNK2B | | |
| | NXF1 | | |
| | xxxxxxxxxxxxxxxxxx | | |
| JAZF1-PHF1 | DHX9 | √ | |
| | EZH1 | | √ |
| | PPARG | √ | √ |
| | EZH2 | | √ |
| | EED | | √ |
| | XRCC6 | | √ |
| | XRCC5 | | √ |
| | PHF1 | | |
| | xxxxx | | |
| | HDAC1 | | √ |
| | PPARG | √ | √ |
| | EZH2 | | √ |
| | PHF1 | | |
| | xxxxx | | |
| | PHF1 | | |
| | TP53 | | √ |
| | XRCC6 | | √ |
| | xxxxxxxxxxxxxxxxxx | | |
| LMNA-NTRK1 | | | |
| | xxxxxxxxxxxxxxxxxx | | |
| MEAF6-TRERF1 | HDAC1 | | √ |
| | TRERF1 | | |
| | KAT5 | √ | √ |
| | ING3 | √ | √ |
| | CREBBP | | √ |
| | YEATS4 | | |
| | EP300 | | √ |
| | MORF4L1 | | |
| | HIST1H2BA | | |
| | xxxxx | | |
| | ELAVL1 | √ | |
| | TRERF1 | | |
| | KAT6A | | |
| | CREBBP | | √ |
| | NR5A1 | | |
| | xxxxx | | |
| | SOX2 | √ | |
| | HDAC1 | | √ |
| | TRERF1 | | |
| | xxxxxxxxxxxxxxxxxx | | |
| MEAF6-PHF1 | DHX9 | √ | |
| | EZH1 | | √ |
| | EZH2 | | √ |
| | EED | | √ |
| | XRCC6 | | √ |
| | XRCC5 | | √ |
| | PHF1 | | |
| | xxxxx | | |
| | PHF1 | | |
| | TP53 | | √ |
| | XRCC6 | | √ |

| | | | |
|---|---|---|---|
| | xxxxx | | |
| | KAT6A | | |
| | TP53 | | √ |
| | ELAVL1 | √ | |
| | xxxxx | | |
| | HDAC1 | | √ |
| | EZH2 | | √ |
| | PHF1 | | |
| | xxxxxxxxxxxxxxxxx | | |
| NR4A3-TAF15 | TRIM28 | √ | |
| | FUS | √ | √ |
| | PRMT1 | | |
| | COPS6 | | |
| | COPS5 | | |
| | POLR2C | | |
| | POLR2A | | |
| | TAF15 | √ | |
| | SF1 | | |
| | NEDD8 | | |
| | POLR2E | | |
| | RPA1 | | √ |
| | RPA2 | √ | |
| | CUL4A | | |
| | CUL4B | √ | |
| | CUL5 | | √ |
| | CUL1 | √ | √ |
| | CUL2 | | √ |
| | CUL3 | | |
| | xxxxx | | |
| | EZH2 | | √ |
| | TRIM28 | √ | |
| | CUL1 | √ | √ |
| | xxxxxxxxxxxxxxxxx | | |
| NR4A3-TFG | CUL4A | | |
| | CUL4B | √ | |
| | CUL5 | | √ |
| | CUL1 | √ | √ |
| | CUL2 | | √ |
| | CUL3 | | |
| | xxxxx | | |
| | TRIM28 | √ | |
| | CUL1 | √ | √ |
| | CUL3 | | |
| | xxxxxxxxxxxxxxxxx | | |
| NR6A1-TRHDE | | | |
| | xxxxxxxxxxxxxxxxx | | |
| NUP107-LGR5 | NUP153 | | |
| | KPNB1 | | |
| | NTRK1 | √ | |
| | CUL3 | | |
| | xxxxx | | |
| | NUP153 | | |
| | EIF4B | | |
| | CUL3 | | |

| | | | |
|---|---|---|---|
| | xxxxx | | |
| | EED | | √ |
| | KPNB1 | | |
| | TP53BP1 | | √ |
| | xxxxxxxxxxxxxxxxxx | | |
| PAPPA-NUP107 | NUP153 | | |
| | ELAVL1 | √ | |
| | KPNB1 | | |
| | NTRK1 | √ | |
| | SMAD3 | √ | |
| | VCP | | |
| | CUL3 | | |
| | EIF4B | | |
| | xxxxx | | |
| | SMAD9 | | |
| | SKIL | √ | √ |
| | PAPPA | | |
| | SMAD2 | √ | √ |
| | SMAD3 | √ | |
| | xxxxx | | |
| | TP53BP1 | | √ |
| | ELAVL1 | √ | |
| | EED | | √ |
| | KPNB1 | | |
| | xxxxx | | |
| | NUP214 | √ | |
| | SMAD2 | √ | √ |
| | SMAD3 | √ | |
| | xxxxxxxxxxxxxxxxxx | | |
| PAX3-FOXO1 | NCOA1 | √ | |
| | ESR1 | √ | √ |
| | PARP1 | | √ |
| | AR | | |
| | EP300 | | √ |
| | CREBBP | | √ |
| | xxxxx | | |
| | TRIM28 | √ | |
| | PARP1 | | √ |
| | CREBBP | | √ |
| | xxxxxxxxxxxxxxxxxx | | |
| PAX7-FOXO1 | RARA | √ | |
| | NCOA1 | √ | |
| | MYOD1 | √ | |
| | ESR1 | √ | √ |
| | HNF4A | | √ |
| | PARP1 | | √ |
| | SMAD3 | √ | |
| | FOXO1 | √ | √ |
| | AR | | |
| | MDM2 | √ | |
| | EP300 | | √ |
| | CREBBP | | √ |
| | xxxxx | | |
| | AKT1 | √ | |

| | | | |
|---|---|---|---|
| | EP300 | | √ |
| | CREBBP | | √ |
| | xxxxxxxxxxxxxxxxxx | | |
| SS18-SSX2 | DPF2 | | |
| | SMARCC2 | | |
| | SMARCC1 | | |
| | PHF10 | | |
| | ELAVL1 | √ | |
| | DPF3 | | |
| | ARID2 | √ | √ |
| | DPF1 | | |
| | SMARCD3 | | |
| | SMARCE1 | | |
| | SMARCD1 | | |
| | EED | | √ |
| | SMARCA2 | √ | √ |
| | EP300 | | √ |
| | SMARCA4 | √ | √ |
| | HDAC1 | | √ |
| | ARID1B | √ | |
| | ARID1A | √ | √ |
| | ACTL6A | | |
| | HDAC2 | | √ |
| | CUL3 | | |
| | RNF2 | | |
| | SMARCD2 | | |
| | xxxxx | | |
| | GRB2 | √ | |
| | YWHAG | | |
| | CUL3 | | |
| | xxxxxxxxxxxxxxxxxx | | |
| SS18-SSX1 | SMARCC2 | | |
| | DPF2 | | |
| | SMARCC1 | | |
| | PHF10 | | |
| | ELAVL1 | √ | |
| | DPF3 | | |
| | DPF1 | | |
| | ARID2 | √ | √ |
| | HDAC2 | | √ |
| | SMARCD3 | | |
| | SMARCE1 | | |
| | SMARCD1 | | |
| | EED | | √ |
| | SMARCA2 | √ | √ |
| | EP300 | | √ |
| | SMARCA4 | √ | √ |
| | HDAC1 | | √ |
| | ARID1B | √ | |
| | ARID1A | √ | √ |
| | ACTL6A | | |
| | CUL3 | | |
| | SMARCD2 | | |
| | xxxxx | | |

| | | | |
|---|---|---|---|
| | GRB2 | √ | |
| | YWHAG | | |
| | CUL3 | | |
| | xxxxxxxxxxxxxxxxx | | |
| SS18L1-SSX1 | SMARCC1 | | |
| | STAT3 | √ | √ |
| | BMI1 | √ | |
| | WHSC1L1 | √ | √ |
| | SMAD3 | √ | |
| | HDAC2 | | √ |
| | SMAD1 | | |
| | EP300 | | √ |
| | SMARCA4 | √ | √ |
| | CREBBP | | √ |
| | xxxxx | | |
| | DPF2 | | |
| | SMARCC1 | | |
| | SMARCE1 | | |
| | SMARCA4 | √ | √ |
| | CUL3 | | |
| | xxxxxxxxxxxxxxxxx | | |
| SSX1-SYT4 | | | |
| | xxxxxxxxxxxxxxxxx | | |
| TGFBR3-MGEA5 | MAST1 | | |
| | RNF32 | | |
| | PAXIP1 | | |
| | xxxxx | | |
| | CBX8 | | |
| | CSNK2B | | |
| | PAXIP1 | | |
| | xxxxxxxxxxxxxxxxx | | |
| TRIO-TERT | YWHAZ | | |
| | AKT1 | √ | |
| | RPS6KB1 | | |
| | ENO1 | √ | |
| | MTOR | √ | |
| | MDM2 | √ | |
| | TERT | √ | |
| | XRCC6 | | √ |
| | xxxxx | | |
| | TERF1 | | |
| | STUB1 | | √ |
| | TERT | √ | |
| | POT1 | | |
| | xxxxx | | |
| | TERT | √ | |
| | MTOR | √ | |
| | YWHAQ | | |
| | RUVBL2 | √ | |
| | xxxxxxxxxxxxxxxxx | | |
| WDR70-RCOR1 | SMARCC2 | | |
| | HDAC1 | | √ |
| | HDAC3 | √ | √ |
| | HDAC2 | | √ |

| | | | |
|---|---|---|---|
| | KDM1A | | |
| | RCOR1 | | |
| | NR2C1 | | |
| | SMARCE1 | | |
| | CTBP1 | | √ |
| | NR2E1 | | |
| | SMARCA4 | √ | √ |
| | CTBP2 | | |
| | KDM5B | | |
| | MTA3 | √ | |
| | xxxxx | | |
| | KDM1A | | |
| | HDAC3 | √ | √ |
| | CTBP1 | | √ |
| | xxxxxxxxxxxxxxxxx | | |
| YWHAE-NUTM2B | LARP1 | | |
| | NOS2 | | |
| | YWHAQ | | |
| | YWHAG | | |
| | HUWE1 | √ | |
| | MAST2 | √ | |
| | NTRK1 | √ | |
| | VCP | | |
| | AKT1 | √ | |
| | MAP2K1 | √ | |
| | FBXW11 | | |
| | ARAF | √ | |
| | CUL3 | | |
| | PARK2 | | √ |
| | RAF1 | √ | |
| | YWHAZ | | |
| | UBXN1 | | |
| | TP53 | | √ |
| | RUVBL2 | √ | |
| | BTRC | √ | |
| | TUBB | | |
| | CDC37 | | |
| | YWHAH | | |
| | MAST3 | | |
| | BRAF | √ | |
| | YWHAB | | |
| | KSR1 | √ | |
| | YWHAE | √ | |
| | CUL1 | √ | √ |
| | MAPK7 | √ | |
| | xxxxx | | |
| | YWHAZ | | |
| | IGF1R | √ | |
| | IRS1 | | |
| | YWHAQ | | |
| | MST1R | √ | √ |
| | NTRK1 | √ | |
| | CBL | √ | √ |
| | TUBB | | |

| | | | |
|---|---|---|---|
| | YWHAB | | |
| | YWHAH | | |
| | GRB2 | √ | |
| | ABL1 | √ | |
| | SORBS2 | √ | |
| | BCAR1 | √ | |
| | YWHAE | √ | |
| | xxxxx | | |
| | VCP | | |
| | CDK2 | √ | √ |
| | CUL1 | √ | √ |
| | xxxxx | | |
| | CDC37 | | |
| | LRRK2 | √ | |
| | YWHAE | √ | |
| | xxxxxxxxxxxxxxxxx | | |
| YWHAE-NUTM2A-AS1 | LARP1 | | |
| | NOS2 | | |
| | YWHAQ | | |
| | YWHAG | | |
| | HUWE1 | √ | |
| | MAST2 | √ | |
| | NTRK1 | √ | |
| | VCP | | |
| | AKT1 | √ | |
| | MAP2K1 | √ | |
| | FBXW11 | | |
| | ARAF | √ | |
| | CUL3 | | |
| | PARK2 | | √ |
| | RAF1 | √ | |
| | YWHAZ | | |
| | UBXN1 | | |
| | TP53 | | √ |
| | RUVBL2 | √ | |
| | BTRC | √ | |
| | TUBB | | |
| | CDC37 | | |
| | YWHAH | | |
| | MAST3 | | |
| | BRAF | √ | |
| | YWHAB | | |
| | KSR1 | √ | |
| | YWHAE | √ | |
| | CUL1 | √ | √ |
| | MAPK7 | √ | |
| | xxxxx | | |
| | YWHAZ | | |
| | IGF1R | √ | |
| | IRS1 | | |
| | YWHAQ | | |
| | MST1R | √ | √ |
| | NTRK1 | √ | |
| | CBL | √ | √ |

| | | | |
|---|---|---|---|
| | TUBB | | |
| | YWHAB | | |
| | YWHAH | | |
| | GRB2 | √ | |
| | ABL1 | √ | |
| | SORBS2 | √ | |
| | BCAR1 | √ | |
| | YWHAE | √ | |
| | xxxxx | | |
| | VCP | | |
| | CDK2 | √ | √ |
| | CUL1 | √ | √ |
| | xxxxx | | |
| | CDC37 | | |
| | LRRK2 | √ | |
| | YWHAE | √ | |
| | xxxxxxxxxxxxxxxxx | | |
| YWHAE-NUTM2A | LARP1 | | |
| | NOS2 | | |
| | YWHAQ | | |
| | YWHAG | | |
| | HUWE1 | √ | |
| | MAST2 | √ | |
| | NTRK1 | √ | |
| | VCP | | |
| | AKT1 | √ | |
| | MAP2K1 | √ | |
| | FBXW11 | | |
| | ARAF | √ | |
| | CUL3 | | |
| | PARK2 | | √ |
| | RAF1 | √ | |
| | YWHAZ | | |
| | UBXN1 | | |
| | TP53 | | √ |
| | RUVBL2 | √ | |
| | BTRC | √ | |
| | TUBB | | |
| | CDC37 | | |
| | YWHAH | | |
| | MAST3 | | |
| | BRAF | √ | |
| | YWHAB | | |
| | KSR1 | √ | |
| | YWHAE | √ | |
| | CUL1 | √ | √ |
| | MAPK7 | √ | |
| | xxxxx | | |
| | YWHAZ | | |
| | IGF1R | √ | |
| | IRS1 | | |
| | YWHAQ | | |
| | MST1R | √ | √ |
| | NTRK1 | √ | |

| | | | |
|---|---|---|---|
| | CBL | √ | √ |
| | TUBB | | |
| | YWHAB | | |
| | YWHAH | | |
| | GRB2 | √ | |
| | ABL1 | √ | |
| | SORBS2 | √ | |
| | BCAR1 | √ | |
| | YWHAE | √ | |
| | xxxxx | | |
| | VCP | | |
| | CDK2 | √ | √ |
| | CUL1 | √ | √ |
| | xxxxx | | |
| | CDC37 | | |
| | LRRK2 | √ | |
| | YWHAE | √ | |
| | xxxxxxxxxxxxxxxxx | | |

Table S5: CA - OG, TS

| | Proteins | OG | TS |
|---|---|---|---|
| ARGLU1-CXCR4 | APP | √ | |
| | CHERP | √ | |
| | SNRNP70 | √ | |
| | SRPK1 | √ | |
| | SRPK2 | | √ |
| | PTK2 | √ | |
| | JAK2 | √ | |
| | SOCS3 | √ | |
| | PTPN11 | √ | |
| | NTRK1 | √ | |
| | PTK2 | √ | |
| | ELAVL1 | √ | |
| | SRPK1 | √ | |
| | NTRK1 | √ | |
| | PTK2 | √ | |
| | JAK2 | √ | |
| | JAK3 | √ | |
| | SOCS3 | √ | |
| | PTPN11 | √ | |
| | STAM | √ | |
| ATXN10-FBLN1 | FN1 | | √ |
| | ATXN10 | √ | |
| | EGFR | √ | |
| | GSTK1 | √ | |
| | ATXN10 | √ | |
| | VCP | | √ |
| | BSG | √ | √ |
| | CUL3 | √ | |
| | VCP | | √ |
| | ABCE1 | √ | √ |
| | ATXN10 | √ | |

| | | | |
|---|---|---|---|
| | APP | √ | |
| | YWHAQ | √ | |
| | CUL3 | √ | |
| BCAS3-NFS1 | CTBP1 | | √ |
| | CTBP2 | √ | |
| | BCAS3 | √ | |
| | KAT2B | √ | |
| | CTBP1 | | √ |
| | CTBP2 | √ | |
| | CDC23 | √ | |
| | KAT2B | √ | |
| | BCAS3 | √ | |
| BCAS4-BCAS3 | CTBP1 | | √ |
| | CTBP2 | √ | |
| | BCAS3 | √ | |
| | KAT2B | √ | |
| | CTBP1 | | √ |
| | CTBP2 | √ | |
| | CDC23 | √ | |
| | KAT2B | √ | |
| | BCAS3 | √ | |
| BCL2L12-PRMT1 | NCOA2 | √ | |
| | NCOA3 | √ | |
| | NCOA1 | √ | |
| | TP53 | | √ |
| | ESR1 | √ | √ |
| | BRCA1 | | √ |
| | PRMT1 | √ | |
| | THRB | √ | |
| | CARM1 | √ | |
| | NR1I2 | √ | |
| | AR | √ | |
| | PPARA | √ | |
| | EP300 | | √ |
| | NCOA2 | √ | |
| | NCOA3 | √ | |
| | NCOA1 | √ | |
| | EP300 | | √ |
| | PARP1 | | √ |
| CAPNS1-WDR62 | YWHAZ | √ | |
| | FBXW11 | √ | |
| | FN1 | | √ |
| | HUWE1 | √ | |
| | YWHAQ | √ | |
| | GAPDH | √ | |
| | FERMT2 | √ | |
| | YWHAH | √ | |
| | VCAM1 | | √ |
| | YWHAB | √ | |
| | PAFAH1B1 | √ | |
| | YWHAG | √ | |
| | PAK2 | √ | |
| | YWHAE | √ | |
| | OGFOD1 | √ | |

| | | | |
|---|---|---|---|
| | MYO1E | √ | |
| | ASNS | √ | |
| | TBCB | √ | |
| | CAPN2 | √ | |
| | PROSC | √ | |
| | YWHAE | √ | |
| CCDC6-ANK3 | HDAC1 | | √ |
| | NR3C1 | √ | |
| | TRIM28 | √ | |
| | SF3A1 | √ | |
| | HNRNPR | √ | |
| | BRCC3 | √ | |
| | HDAC1 | | √ |
| | NR3C1 | √ | |
| | TRIM28 | √ | |
| | ELAVL1 | √ | |
| | SKP1 | √ | |
| | HNRNPR | √ | |
| | PPP1CA | √ | |
| | BRCC3 | √ | |
| | NTRK1 | √ | |
| | SF3A1 | √ | |
| | CUL1 | √ | √ |
| | FBXW7 | √ | √ |
| CCDC9-DHX34 | EIF4A3 | √ | |
| | SNIP1 | √ | |
| | CCDC9 | √ | |
| | PRPF40A | √ | |
| CDC27-ST7L | CDC16 | √ | |
| | CDC27 | √ | |
| | MDC1 | √ | |
| | CDC20 | √ | |
| | CREBBP | | √ |
| | ANAPC2 | √ | |
| | ANAPC7 | √ | |
| | CREBBP | | √ |
| | E2F1 | √ | |
| | RB1 | √ | |
| | TFDP1 | √ | |
| | CDC16 | √ | |
| | CDC27 | √ | |
| | MDC1 | √ | |
| | SMAD2 | √ | √ |
| | TP53BP1 | | √ |
| | CREBBP | | √ |
| | UBE2S | √ | |
| | ANAPC2 | √ | |
| | ANAPC7 | √ | |
| CDK7-RIN3 | BRCA1 | | √ |
| | TP53 | | √ |
| | RUVBL2 | √ | |
| | SUPT5H | √ | |
| | ESR1 | √ | √ |
| | RPA1 | | √ |

| | | | |
|---|---|---|---|
| | HNRNPU | | √ |
| | RPA2 | √ | √ |
| | CDK2 | √ | √ |
| | POLR2A | | √ |
| | GTF2H1 | √ | |
| | RPA1 | | √ |
| | RPA2 | √ | √ |
| | POLR2A | | √ |
| | CCNH | √ | |
| | HDAC2 | | √ |
| | TP53 | | √ |
| | ESR1 | √ | √ |
| | MTA1 | √ | |
| | GTF2H1 | √ | |
| | ERCC3 | √ | |
| | POLR2A | | √ |
| | ERCC5 | √ | |
| | BRCA1 | | √ |
| | POLR2A | | √ |
| | RUVBL2 | √ | |
| | GTF2H1 | √ | |
| | RPA1 | | √ |
| | RPA2 | √ | √ |
| | PRKCI | √ | |
| | APP | √ | |
| | CDK7 | √ | |
| | CDC37 | √ | |
| CHERP-CPAMD8 | DHX8 | √ | |
| | U2AF1 | √ | |
| | RPA1 | | √ |
| | PRPF40A | √ | |
| | RPA2 | √ | √ |
| | CHERP | √ | |
| | SF3A2 | √ | |
| | RBM39 | √ | |
| | EWSR1 | √ | |
| | DHX8 | √ | |
| | AGGF1 | √ | |
| | RNPS1 | √ | |
| | SNIP1 | √ | |
| | SF3B4 | √ | |
| | NTRK1 | √ | |
| | CHERP | √ | |
| | SRPK1 | √ | |
| | SRPK2 | | √ |
| | U2AF1 | √ | |
| | U2AF2 | √ | |
| | RPA1 | | √ |
| | APBB1 | √ | |
| | PRPF40A | √ | |
| | RPA2 | √ | √ |
| | TTC14 | √ | |
| | RBM23 | √ | |
| | SF3A2 | √ | |

| | | | |
|---|---|---|---|
| | SNRNP70 | √ | |
| | RBM39 | √ | |
| | EWSR1 | √ | |
| | WBP4 | √ | |
| CYTH1-PRPSAP1 | DDX17 | √ | |
| | DDX5 | √ | |
| | ILK | | √ |
| | COPS5 | | √ |
| | CYTH1 | √ | |
| | ARRB2 | √ | |
| | ARF6 | √ | |
| | ARRB1 | √ | |
| | DDX5 | √ | |
| | FBXW11 | √ | |
| | DDX17 | √ | |
| | ILK | | √ |
| | ITGB2 | √ | |
| | COPS5 | | √ |
| DLG1-CRYBG3 | DLG1 | √ | |
| | LIN7A | √ | |
| | LIN7C | √ | |
| | APBA1 | √ | √ |
| | CASK | √ | |
| | DLG1 | √ | |
| | NTRK1 | √ | |
| | CASK | √ | |
| | EPB41 | √ | |
| | DLG1 | √ | |
| | KHDRBS1 | √ | |
| | LCK | √ | |
| | NTRK1 | √ | |
| | MAPK1 | √ | |
| | ARRB2 | √ | |
| | ARRB1 | √ | |
| DTX4-CCDC102B | MCM7 | √ | |
| | CDK18 | √ | |
| | LENG1 | √ | |
| | TRIM54 | √ | |
| | TRIM27 | √ | |
| | KIFC3 | √ | |
| | SFN | √ | |
| | MARK1 | √ | |
| | CCDC102B | √ | |
| EHD4-FSIP1 | EHD4 | √ | |
| | CTPS2 | √ | |
| | EHD1 | √ | |
| | EGFR | √ | |
| | NTRK1 | √ | |
| | WARS | √ | |
| | PLCG1 | √ | |
| | UBA2 | √ | |
| | ADSL | √ | |
| | UQCRC2 | √ | |
| | PLCG1 | √ | |

| | | | |
|---|---|---|---|
| | EHD4 | √ | |
| | EGFR | √ | |
| | NTRK1 | √ | |
| ELK4-SLC26A9 | BRCA1 | | √ |
| | MAPK3 | √ | √ |
| | MAPK1 | √ | |
| | ELK4 | √ | |
| | BRCA1 | | √ |
| | MAPK3 | √ | √ |
| | MAPK1 | √ | |
| | ELK4 | √ | |
| | BLM | √ | |
| ERAL1-DIDO1 | HNRNPDL | √ | |
| | RPA1 | | √ |
| | RPA2 | √ | √ |
| | HNRNPK | √ | |
| | RBM15 | √ | |
| | CUL3 | √ | |
| | DIDO1 | √ | |
| | FUS | √ | √ |
| | FUS | √ | √ |
| | APP | √ | |
| | RPA1 | | √ |
| | RPA2 | √ | √ |
| | RBM15 | √ | |
| | CUL3 | √ | |
| | DIDO1 | √ | |
| | SRPK2 | | √ |
| GMDS-CCND3 | PCNA | √ | |
| | PPP1CC | √ | √ |
| | RBL2 | √ | |
| | PPP1CA | √ | |
| | CCND3 | √ | |
| | RB1 | √ | |
| | POLD1 | √ | |
| | CDK2 | √ | √ |
| | CDK4 | √ | |
| | CDK6 | √ | |
| | CREBBP | | √ |
| | GMDS | √ | |
| | NSFL1C | √ | |
| | CTH | √ | |
| | CAPN2 | √ | |
| | ATIC | √ | |
| | RARA | √ | |
| | NCOA2 | √ | |
| | VDR | | √ |
| | CCND3 | √ | |
| | CREBBP | | √ |
| | MCM10 | √ | |
| | RBX1 | √ | |
| | CCND3 | √ | |
| | APP | √ | |
| | NCOA2 | √ | |

| | | | |
|---|---|---|---|
| | RARA | √ | |
| | VDR | | √ |
| | CCND3 | √ | |
| | CREBBP | | √ |
| HJURP-EIF4E2 | FBXW11 | √ | |
| | TP53 | | √ |
| | GIGYF2 | √ | |
| | APP | √ | |
| | HUWE1 | √ | |
| | EIF4E2 | √ | |
| | YWHAB | √ | |
| | SHMT2 | | √ |
| | YWHAE | √ | |
| INTS4-GAB2 | SRC | √ | |
| | PLCG1 | √ | |
| | GRB2 | √ | |
| | ZAP70 | √ | |
| | NTRK1 | √ | |
| | SHC1 | √ | |
| | PIK3CB | √ | |
| | PIK3R2 | √ | |
| | PIK3R1 | √ | |
| KDM5A-ANO2 | HDAC1 | | √ |
| | HDAC2 | | √ |
| | RBL1 | √ | |
| | TBP | √ | |
| | RB1 | √ | |
| | VDR | | √ |
| | KDM5A | √ | |
| | MORF4L1 | √ | |
| | HDAC2 | | √ |
| | EZH2 | | √ |
| | KDM5A | √ | |
| | ESR1 | √ | √ |
| MAPK10-FAM13A | HDAC1 | | √ |
| | TP53 | | √ |
| | HDAC9 | √ | |
| | JUN | √ | |
| | DDX5 | √ | |
| | ELK1 | √ | |
| | MAPK10 | √ | |
| | RELA | √ | |
| | CREBBP | | √ |
| | ATF2 | √ | |
| | APP | √ | |
| | MAPK10 | √ | |
| | MAP2K4 | √ | |
| MAPRE1-TM9SF4 | YWHAZ | √ | |
| | FN1 | | √ |
| | APP | √ | |
| | TUBB | √ | |
| | NTRK1 | √ | |
| | VCAM1 | | √ |
| | UNK | √ | √ |

| | | | |
|---|---|---|---|
| | COPS5 | | √ |
| | CDK5RAP2 | √ | |
| | PRKACA | √ | |
| | AKAP9 | √ | |
| | PRKACB | √ | |
| | CLIP1 | √ | |
| | TUBB | √ | |
| | TUBA1A | √ | |
| | HDAC6 | √ | |
| | PDE4DIP | √ | |
| | PRKACA | √ | |
| | PRKACB | √ | |
| | CDK5RAP2 | √ | |
| | AKAP9 | √ | |
| | TERF1 | √ | √ |
| | SPTAN1 | √ | |
| | DST | √ | |
| | MAPRE1 | √ | |
| NUMB-ALDH6A1 | TP53 | | √ |
| | NUMB | √ | |
| | ITCH | √ | |
| | MDM2 | √ | |
| | EGFR | √ | |
| | EPS15 | √ | |
| | EGFR | √ | |
| | AP2A1 | √ | |
| | NUMB | √ | |
| | PRKCZ | √ | |
| | NUMB | √ | |
| | APP | √ | |
| | EGFR | √ | |
| PARD6B-CD48 | PRKCI | √ | |
| | RASSF8 | √ | |
| | PARD3 | √ | |
| | PARD6G | √ | |
| | APP | √ | |
| | PARD6B | √ | |
| | PARD6A | √ | |
| | YWHAH | √ | |
| | PRKCZ | √ | |
| | WWC1 | √ | |
| | PRKCI | √ | |
| | PARD3 | √ | |
| | PARD6G | √ | |
| | APP | √ | |
| | PARD6B | √ | |
| | PARD6A | √ | |
| | YWHAH | √ | |
| | PRKCZ | √ | |
| | RAC1 | √ | √ |
| | PARD6G | √ | |
| | PARD6B | √ | |
| | PARD6A | √ | |
| PPP1R12A-MGAT4C | KDM1A | √ | √ |

| | | | |
|---|---|---|---|
| | ELAVL1 | √ | |
| | RPA1 | | √ |
| | RPA2 | √ | √ |
| | PPP1R12A | √ | |
| | CUL1 | √ | √ |
| | KDM1A | √ | √ |
| | NUDT5 | √ | |
| | TP53 | | √ |
| | ELAVL1 | √ | |
| | PUS1 | √ | |
| | NUAK1 | √ | |
| | AARSD1 | √ | |
| | RPA1 | | √ |
| | NTRK1 | √ | |
| | RPA2 | √ | √ |
| | RPRD1B | √ | |
| | PPP1R12A | √ | |
| | TRIM47 | √ | |
| | PAXIP1 | √ | √ |
| | ACTR3 | √ | |
| | CUL1 | √ | √ |
| RNF11-C8A | CBLB | √ | |
| | RNF11 | √ | |
| | ITCH | √ | |
| | SMAD4 | √ | √ |
| | EPN1 | √ | |
| | RABGEF1 | √ | |
| | UBE2E1 | √ | √ |
| | UBE2D3 | √ | √ |
| | UBE2E3 | √ | |
| | HGS | √ | √ |
| | GGA1 | √ | |
| | AKT1 | √ | |
| | GGA3 | √ | |
| | GGA2 | √ | |
| | AP2A1 | √ | |
| | EPN3 | √ | |
| | UBE2D1 | √ | √ |
| | AP2B1 | √ | |
| | CSNK2A1 | | √ |
| | SMURF1 | √ | |
| | EPS15 | √ | |
| | SMURF2 | √ | |
| | UBQLN2 | √ | |
| | STAM2 | √ | |
| | NEDD4 | √ | √ |
| | UBQLN4 | √ | √ |
| | NEDD4L | √ | √ |
| | APP | √ | |
| | RNF11 | √ | |
| | PSMD4 | √ | |
| | PSMD7 | √ | |
| | PSMD6 | √ | |
| | PSMD11 | √ | |

| | | | |
|---|---|---|---|
| | PSMD10 | √ | |
| | PSMD3 | √ | |
| | PSMD12 | √ | |
| | PSMD13 | √ | |
| | USP14 | √ | |
| | PSMD14 | √ | |
| | PSMD1 | √ | |
| | PSMD2 | √ | |
| | APP | √ | |
| | GGA1 | √ | |
| | GGA3 | √ | |
| | GGA2 | √ | |
| | APP | √ | |
| | AKT1 | √ | |
| | RNF11 | √ | |
| | TBK1 | √ | |
| | APP | √ | |
| | NEDD4 | √ | √ |
| SIPA1L3-WDR62 | MAPK10 | √ | |
| | WDR62 | √ | |
| | MAPK8 | √ | |
| | MAPK9 | √ | |
| | SFN | √ | |
| | YWHAB | √ | |
| | SIPA1L3 | √ | |
| | YWHAQ | √ | |
| | YWHAB | √ | |
| | FBXW11 | √ | |
| | MAPK10 | √ | |
| | ELAVL1 | √ | |
| | WDR62 | √ | |
| | TBP | √ | |
| | MAPK8 | √ | |
| | MAPK9 | √ | |
| SLC26A6-PRKAR2A | AKAP7 | √ | |
| | AKAP9 | √ | |
| | PRKAR2A | √ | |
| | PRKAR2B | √ | |
| | PRKACA | √ | |
| | PRKACB | √ | |
| | PRKAR2A | √ | |
| | AKAP7 | √ | |
| | PRKACA | √ | |
| | PRKACB | √ | |
| | PRKAR2B | √ | |
| | GCH1 | √ | |
| ST14-APLP2 | BRCA1 | | √ |
| | APLP2 | √ | |
| | ETS1 | √ | |
| | JUN | √ | |
| | SFN | √ | |
| | HDAC5 | √ | |
| | APLP2 | √ | |
| | RPL26 | √ | |

| | | | |
|---|---|---|---|
| | BRCA1 | | √ |
| | JUNB | √ | |
| | JUN | √ | |
| | APBB1 | √ | |
| | APBB2 | √ | |
| | KAT5 | √ | √ |
| | ETS1 | √ | |
| | APLP2 | √ | |
| | MAPK8 | √ | |
| STRADB-NOP58 | NIFK | √ | |
| | NOP56 | √ | |
| | HNRNPU | | √ |
| | RUVBL2 | √ | |
| | NOLC1 | √ | |
| | SNU13 | √ | |
| | NTRK1 | √ | |
| | PUM3 | √ | |
| | RPS15A | √ | |
| | RSL1D1 | √ | |
| | KRR1 | √ | |
| | DDX18 | √ | |
| | EIF6 | √ | |
| | DHX15 | √ | |
| | RPS4X | √ | |
| | EED | | √ |
| | PRPF3 | √ | |
| | TARDBP | √ | |
| | RPL11 | √ | |
| | EIF2S2 | √ | |
| | DDX27 | √ | |
| | NOP58 | √ | |
| | FN1 | | √ |
| | DDX24 | √ | |
| | U2AF1 | √ | |
| | RPL30 | √ | |
| | ESR1 | √ | √ |
| | DDX56 | √ | |
| | FTSJ3 | √ | |
| | DDX47 | √ | |
| | KPNA6 | √ | |
| | GTPBP4 | √ | |
| | WDR36 | √ | |
| | KPNA1 | √ | |
| | DKC1 | √ | |
| | RRP12 | √ | |
| | BOP1 | √ | |
| | FBL | √ | |
| | TBL3 | √ | |
| | WDR36 | √ | |
| | NOP58 | √ | |
| | DHX15 | √ | |
| | NOP56 | √ | |
| STX16-RAE1 | FBXW11 | √ | |
| | NXF1 | √ | |

| | | | |
|---|---|---|---|
| | FAF1 | √ | |
| | RAE1 | √ | |
| | ILF3 | √ | |
| | CUL1 | √ | √ |
| | HNRNPUL1 | √ | |
| | CUL3 | √ | |
| | NXF1 | √ | |
| | CUL1 | √ | √ |
| | ILF3 | √ | |
| | CUL3 | √ | |
| TANC2-CHD6 | ZFYVE9 | √ | |
| | PPP1CC | √ | √ |
| | PPP1CA | √ | |
| | TANC2 | √ | |
| | ZFYVE9 | √ | |
| | PPP1CC | √ | √ |
| | PPP1CA | √ | |
| | TANC2 | √ | |
| | MAEA | √ | |
| | RANBP9 | √ | |
| | MKLN1 | √ | |
| | MMP7 | √ | |
| | RMND5A | √ | |
| | MAEA | √ | |
| | RANBP9 | √ | |
| | MKLN1 | √ | |
| | RANBP10 | √ | |
| | MMP7 | √ | |
| | RMND5A | √ | |
| TMPRSS2-ERG | CDC5L | | √ |
| | DDX3X | √ | |
| | ELAVL1 | √ | |
| | CAD | √ | |
| | NEDD4 | √ | √ |
| | SF3B1 | √ | |
| | PARP1 | | √ |
| | PRKDC | √ | |
| | PRPF8 | √ | |
| | SFPQ | √ | |
| | ERG | √ | |
| | POLR2A | | √ |
| | TOP1 | √ | |
| | CLTC | √ | |
| | SF3B2 | √ | |
| | XRCC5 | | √ |
| | XRCC6 | | √ |
| | DDX23 | √ | |
| | SNRNP200 | √ | |
| | DDX21 | √ | |
| | TUBB | √ | |
| | NONO | √ | |
| | JUN | √ | |
| | HNRNPU | | √ |
| | PRPF40A | √ | |

| | | | |
|---|---|---|---|
| | AR | √ | |
| | NCL | √ | |
| | HNRNPM | √ | |
| | HNRNPC | √ | |
| | TOP2B | √ | |
| | ILF3 | √ | |
| | ILF2 | √ | |
| | PRPF8 | √ | |
| | ERG | √ | |
| | SF3B2 | √ | |
| | SF3B1 | √ | |
| | PARP1 | | √ |
| | PRKDC | √ | |

Table S6: LK,LY,ME,GL –DR/PA

| | UN | OG | TS |
|---|---|---|---|
| BCR-ABL1 | | | |
| RUNX1-RUNX1T1 | | | √ |
| KMT2A-MLLT10 | SMARCC2 | | √ |
| | POLR2A | | √ |
| IGH-BCL2 | CASP3 | | √ |
| KMT2A-AFF1 | SMARCC2 | | √ |
| | POLR2A | | √ |
| PICALM-MLLT10 | DNM2 | √ | |
| | ILVBL | √ | |
| | SEC24D | √ | |
| | SEC24C | √ | |
| PML-RARA | NR3C1 | √ | |
| | KAT2B | √ | |
| | TRIP4 | √ | |
| KMT2A-MLLT3 | SMARCC2 | | √ |
| | POLR2A | | √ |
| KMT2A-AFDN | SMARCC2 | | √ |
| | POLR2A | | √ |
| CBFB-MYH11 | ACTA2 | √ | |
| | MYO1E | √ | |
| IGH-MYC | | | |
| NUP98-DDX10 | SIRT7 | √ | |
| | APP | √ | |
| | DDX56 | √ | |
| | DDX54 | √ | |
| | PUM3 | √ | |
| | PWP1 | √ | |
| | PUM3 | √ | |
| | NXF1 | √ | |
| | KPNB1 | √ | |
| | HNRNPUL1 | √ | |
| | CDC37 | √ | |
| PCM1-JAK2 | TEC | √ | |
| | INSR | √ | |
| KMT2A-SEPT9 | SMARCC2 | | √ |
| | POLR2A | | √ |
| FUS-ERG | SF3B2 | √ | |

| | | | |
|---|---|---|---|
| | PRKDC | √ | |
| | SF3A2 | √ | |
| | DHX15 | √ | |
| | CUL3 | √ | |
| | PRKDC | √ | |
| NUP98-HOXA9 | | | |
| ETV6-ABL1 | UBASH3B | √ | |
| SET-NUP214 | NXF1 | √ | |
| | FAF1 | √ | |
| | SUPT5H | √ | |
| | GART | √ | |
| | CUL3 | √ | |
| MNX1-ETV6 | HDAC9 | √ | |
| | L3MBTL1 | √ | |
| | ETV7 | √ | |
| KMT2A-MLLT1 | SMARCC2 | | √ |
| | POLR2A | | √ |
| KMT2A-MLLT6 | SMARCC2 | | √ |
| | POLR2A | | √ |
| ETV6-ACSL6 | HDAC9 | √ | |
| | L3MBTL1 | √ | |
| | ETV7 | √ | |
| ETV6-MECOM | UBE2I | √ | |
| | EHMT2 | √ | |
| | SMAD1 | √ | |
| | KAT2B | √ | |
| | SUV39H1 | √ | |
| | SUV39H1 | √ | |
| KMT2A-EPS15 | SMARCC2 | | √ |
| | POLR2A | | √ |
| KMT2A-GAS7 | SMARCC2 | | √ |
| | POLR2A | | √ |
| KMT2A-ABL1 | SMARCC2 | | √ |
| | POLR2A | | √ |
| KMT2A-MLLT11 | SMARCC2 | | √ |
| | POLR2A | | √ |
| MN1-ETV6 | HDAC9 | √ | |
| KMT2A-MAML2 | SMARCC2 | | √ |
| KMT2A-FOXO4 | SMARCC2 | | √ |
| | POLR2A | | √ |
| DEK-NUP214 | KAT2B | √ | |
| | NXF1 | √ | |
| | DHX15 | √ | |
| | CUL3 | √ | |
| RUNX1-CBFA2T3 | | | |
| NUP98-PSIP1 | NXF1 | √ | |
| | EIF4A3 | | |
| | SON | | |
| NUP98-HOXC13 | NXF1 | √ | |
| | HNRNPAB | | |
| | KPNB1 | | |
| | HNRNPUL1 | | |
| NUP98-HOXC11 | SP1 | √ | |
| PAX5-ETV6 | UBE2I | √ | |

| | | | |
|---|---|---|---|
| | TBP | √ | |
| NUP98-HOXA11 | YY1 | √ | |
| BCR-PDGFRA | UBASH3B | √ | |
| | INPP5D | √ | |
| BCR-FGFR1 | | | |
| NPM1-RARA | | | |
| KMT2A-CBL | | | |
| IGH-BCL6 | | | √ |
| LCP1-BCL6 | | | √ |
| CREBBP-KAT6A | SMARCC2 | | √ |
| | POLR2A | | √ |
| KMT2A-ARHGAP26 | SMARCC2 | | √ |
| | POLR2A | | √ |
| FOXO3-KMT2A | SMARCC2 | | √ |
| KMT2A-DCPS | SMARCC2 | | √ |
| | POLR2A | | √ |
| KMT2A-EP300 | | | |
| IGH-CEBPE | UBE2I | √ | |
| | CEBPG | √ | |
| | CEBPE | √ | |
| | BATF3 | √ | |
| HSP90AA1-BCL6 | | | |

Table S7: SC -DR/PA

| | UN | OG | TS |
|---|---|---|---|
| ASPSCR1-TFE3 | | | |
| ASTN2-CNOT2 | CNOT6L | √ | |
| | CNOT8 | √ | |
| | TNRC6C | √ | |
| | TNRC6B | √ | |
| | CNOT2 | √ | |
| | CNOT1 | √ | |
| | CNOT7 | √ | |
| | AGO2 | √ | |
| | CNOT2 | √ | |
| BCOR-ZC3H7B | SP1 | √ | |
| | NACC1 | √ | |
| BCOR-CCNB3 | SP1 | √ | |
| | NACC1 | √ | |
| CDX1-IRF2BP2 | IRF2BPL | √ | |
| CIC-DUX4 | | | |
| CREB1-EWSR1 | POLR2A | | √ |
| | NR3C1 | √ | |
| | CUL3 | √ | |
| CTDSP2-FAM19A2 | SETD1A | | √ |
| | POLR2A | | √ |
| | CTDSP1 | | √ |
| | CTDSP2 | | √ |
| CXorf67-MBTD1 | | | |
| EPC1-PHF1 | PHF1 | | √ |
| | YEATS4 | | √ |
| | TRIM23 | | √ |
| | HIST1H2BA | | √ |
| | DMAP1 | | √ |

| | | | |
|---|---|---|---|
| | MORF4L1 | | √ |
| ERG-EWSR1 | | | |
| ETV6-NTRK3 | | | |
| EWSR1-ATF1 | | | |
| EWSR1-FLI1 | POLR2A | | √ |
| | KAT2B | √ | |
| EWSR1-NR4A3 | PRMT1 | √ | |
| | POLR2A | | √ |
| | CHERP | √ | |
| | ATXN3 | √ | |
| | CUL4A | √ | |
| | FASN | √ | |
| | CUL3 | √ | |
| | FXR2 | √ | |
| | CUL3 | √ | |
| EWSR1-ETV4 | PRMT1 | √ | |
| | POLR2A | | √ |
| | CHERP | √ | |
| | ATXN3 | √ | |
| | CUL4A | √ | |
| | FASN | √ | |
| | CUL3 | √ | |
| | FXR2 | √ | |
| | CUL3 | √ | |
| EWSR1-PATZ1 | PRMT1 | √ | |
| | POLR2A | | √ |
| | CHERP | √ | |
| | ATXN3 | √ | |
| | CUL4A | √ | |
| | FASN | √ | |
| | CUL3 | √ | |
| | FXR2 | √ | |
| | CUL3 | √ | |
| EWSR1-DDIT3 | POLR2A | | √ |
| | CUL4A | √ | |
| | CUL3 | √ | |
| EWSR1-POU5F1 | ETS2 | √ | √ |
| | POLR2A | | √ |
| | CUL3 | √ | |
| EWSR1-SP3 | POLR2A | | √ |
| | CUL4A | √ | |
| | CUL3 | √ | |
| FOXO4-CIC | XPO1 | √ | |
| | NLK | √ | |
| FUS-ERG | SF3B2 | √ | |
| | PRKDC | √ | |
| | SF3A2 | √ | |
| | DHX15 | | |
| | CUL3 | √ | |
| | PRKDC | √ | |
| FUS-CREB3L1 | CUL4A | √ | |
| | DHX15 | √ | |
| | CUL3 | √ | |
| | VCP | √ | |

| | | |
|---|---|---|
| | FBXW11 | √ |
| FUS-DDIT3 | DDX17 | √ |
| | TRIP4 | √ |
| | VCP | √ |
| | FBXW11 | √ |
| FUS-ATF1 | DDX17 | √ |
| | TRIP4 | √ |
| | VCP | √ |
| | FBXW11 | √ |
| FUS-CREB3L2 | CUL4A | √ |
| | DHX15 | √ |
| | CUL3 | √ |
| | VCP | √ |
| | FBXW11 | √ |
| HEY1-NCOA2 | NR3C1 | √ |
| | PRMT1 | √ |
| | AR | √ |
| | NR1I3 | √ |
| | UBR5 | √ |
| IRX2-TERT | YWHAZ | √ |
| | RPS6KB1 | √ |
| | TERF1 | √ |
| | POT1 | √ |
| | YWHAQ | √ |
| | TPP1 | √ |
| | POT1 | √ |
| JAZF1-SUZ12 | FBXW11 | √ |
| | NXF1 | √ |
| | SF3B4 | √ |
| | PRMT1 | √ |
| | SF3B2 | √ |
| | CRNKL1 | √ |
| | RNPS1 | √ |
| | UBE2I | √ |
| | SNRNP200 | √ |
| | SRSF7 | √ |
| | SNRPD3 | √ |
| | RALY | √ |
| | PRPF19 | √ |
| | RNF2 | √ |
| | SON | √ |
| | EFTUD2 | √ |
| | SF3A1 | √ |
| | EPRS | √ |
| | ILF2 | √ |
| | EIF4A3 | √ |
| | CDC40 | √ |
| | UHRF1 | √ |
| | GATAD2B | √ |
| | CBX3 | √ |
| | SETDB1 | √ |
| | VCP | √ |
| | NXF1 | √ |
| | VCP | √ |

| | | | |
|---|---|---|---|
| | FBXW11 | √ | |
| | ADAR | √ | |
| | FBXW11 | √ | |
| | NXF1 | √ | |
| | JARID2 | √ | |
| | SETDB1 | √ | |
| | FBXW11 | √ | |
| | CSNK2B | √ | |
| | NXF1 | √ | |
| JAZF1-PHF1 | PHF1 | | √ |
| | PHF1 | | √ |
| | PHF1 | | √ |
| LMNA-NTRK1 | | | |
| MEAF6-TRERF1 | TRERF1 | | √ |
| | YEATS4 | | √ |
| | MORF4L1 | | √ |
| | HIST1H2BA | | √ |
| | TRERF1 | | √ |
| | KAT6A | | √ |
| | NR5A1 | | √ |
| | TRERF1 | | √ |
| MEAF6-PHF1 | PHF1 | | √ |
| | PHF1 | | √ |
| | KAT6A | | √ |
| | PHF1 | | √ |
| NR4A3-TAF15 | PRMT1 | √ | |
| | COPS6 | √ | |
| | COPS5 | √ | |
| | POLR2C | √ | |
| | POLR2A | √ | |
| | SF1 | √ | |
| | NEDD8 | √ | |
| | POLR2E | √ | |
| | CUL4A | √ | |
| | CUL3 | √ | |
| NR4A3-TFG | CUL4A | √ | |
| | CUL3 | √ | |
| | CUL3 | √ | |
| | NUP153 | √ | |
| | KPNB1 | √ | |
| NR6A1-TRHDE | CUL3 | √ | |
| NUP107-LGR5 | NUP153 | √ | |
| | EIF4B | √ | |
| | CUL3 | √ | |
| | KPNB1 | √ | |
| | NUP153 | √ | |
| | KPNB1 | √ | |
| PAPPA-NUP107 | VCP | √ | |
| | CUL3 | √ | |
| | EIF4B | √ | |
| | SMAD9 | √ | |
| | PAPPA | √ | |
| | KPNB1 | √ | |
| | AR | √ | |

| | | | |
|---|---|---|---|
| PAX3-FOXO1 | | | |
| PAX7-FOXO1 | AR | √ | |
| | DPF2 | √ | |
| | SMARCC2 | √ | |
| | SMARCC1 | √ | |
| | PHF10 | √ | |
| SS18-SSX2 | DPF3 | √ | |
| | DPF1 | √ | |
| | SMARCD3 | √ | |
| | SMARCE1 | √ | |
| | SMARCD1 | √ | |
| | ACTL6A | √ | |
| | CUL3 | √ | |
| | RNF2 | √ | |
| | SMARCD2 | √ | |
| | YWHAG | √ | |
| | CUL3 | √ | |
| | SMARCC2 | √ | |
| | DPF2 | √ | |
| | SMARCC1 | √ | |
| | PHF10 | √ | |
| SS18-SSX1 | DPF3 | √ | |
| | DPF1 | √ | |
| | SMARCD3 | √ | |
| | SMARCE1 | √ | |
| | SMARCD1 | √ | |
| | ACTL6A | √ | |
| | CUL3 | √ | |
| | SMARCD2 | √ | |
| | YWHAG | √ | |
| | CUL3 | √ | |
| | SMARCC1 | √ | |
| SS18L1-SSX1 | SMAD1 | √ | |
| | DPF2 | √ | |
| | SMARCC1 | √ | |
| | SMARCE1 | √ | |
| | CUL3 | √ | |
| | MAST1 | √ | |
| | RNF32 | √ | |
| SSX1-SYT4 | PAXIP1 | √ | |
| TGFBR3-MGEA5 | CSNK2B | √ | |
| | PAXIP1 | √ | |
| | YWHAZ | √ | |
| | RPS6KB1 | √ | |
| TRIO-TERT | TERF1 | √ | |
| | POT1 | √ | |
| | YWHAQ | √ | |
| | SMARCC2 | | √ |
| WDR70-RCOR1 | KDM1A | √ | |
| | RCOR1 | √ | |
| | NR2C1 | √ | |
| | SMARCE1 | √ | |
| | NR2E1 | √ | |
| | CTBP2 | √ | |

| | | |
|---|---|---|
| | KDM5B | √ |
| | KDM1A | √ |
| | LARP1 | √ |
| | NOS2 | √ |
| | YWHAQ | √ |
| | YWHAG | √ |
| YWHAE-NUTM2B | VCP | √ |
| | FBXW11 | √ |
| | CUL3 | √ |
| | YWHAZ | √ |
| | UBXN1 | √ |
| | TUBB | √ |
| | CDC37 | √ |
| | YWHAH | √ |
| | MAST3 | √ |
| | YWHAB | √ |
| | YWHAZ | √ |
| | IRS1 | √ |
| | YWHAQ | √ |
| | TUBB | √ |
| | YWHAB | √ |
| | YWHAH | √ |
| | VCP | √ |
| | CDC37 | √ |
| | LARP1 | √ |
| | NOS2 | √ |
| | YWHAQ | √ |
| | YWHAG | √ |
| YWHAE-NUTM2A-AS1 | VCP | √ |
| | FBXW11 | √ |
| | CUL3 | √ |
| | YWHAZ | √ |
| | UBXN1 | √ |
| | TUBB | √ |
| | CDC37 | √ |
| | YWHAH | √ |
| | MAST3 | √ |
| | YWHAB | √ |
| | YWHAZ | √ |
| | IRS1 | √ |
| | YWHAQ | √ |
| | TUBB | √ |
| | YWHAB | √ |
| | YWHAH | √ |
| | VCP | √ |
| | CDC37 | √ |
| | LARP1 | √ |
| | NOS2 | √ |
| | YWHAQ | √ |
| | YWHAG | √ |
| YWHAE-NUTM2A | VCP | √ |
| | FBXW11 | √ |
| | CUL3 | √ |
| | YWHAZ | √ |

| | | |
|---|---|---|
| | UBXN1 | √ |
| | TUBB | √ |
| | CDC37 | √ |
| | YWHAH | √ |
| | MAST3 | √ |
| | YWHAB | √ |
| | YWHAZ | √ |
| | IRS1 | √ |
| | YWHAQ | √ |
| | TUBB | √ |
| | YWHAB | √ |
| | YWHAH | √ |
| | VCP | √ |
| | CDC37 | √ |

Table S8: CA -DR/PA

| | Essential Community Vertices | OG | TS |
|---|---|---|---|
| ARGLU1-CXCR4 | APP | √ | |
| | CHERP | √ | |
| | SNRNP70 | √ | |
| | SRPK1 | √ | |
| | PTK2 | √ | |
| | PTK2 | √ | |
| | SRPK1 | √ | |
| | PTK2 | √ | |
| | STAM | √ | |
| ATXN10-FBLN1 | ATXN10 | √ | |
| | GSTK1 | √ | |
| | ATXN10 | √ | |
| | VCP | | √ |
| | BSG | √ | √ |
| | CUL3 | √ | |
| | VCP | | √ |
| | ABCE1 | √ | √ |
| | ATXN10 | √ | |
| | APP | √ | |
| | YWHAQ | √ | |
| | CUL3 | √ | |
| BCAS3-NFS1 | CTBP2 | √ | |
| | KAT2B | √ | |
| | CTBP2 | √ | |
| | CDC23 | √ | |
| | KAT2B | √ | |
| BCAS4-BCAS3 | CTBP2 | √ | |
| | KAT2B | √ | |
| | CTBP2 | √ | |
| | CDC23 | √ | |
| | KAT2B | √ | |
| BCL2L12-PRMT1 | PRMT1 | √ | |
| | AR | √ | |
| CAPNS1-WDR62 | YWHAZ | √ | |
| | FBXW11 | √ | |

| | | | |
|---|---|---|---|
| | YWHAQ | √ | |
| | FERMT2 | √ | |
| | YWHAH | √ | |
| | VCAM1 | | √ |
| | YWHAB | √ | |
| | PAFAH1B1 | √ | |
| | YWHAG | √ | |
| | PAK2 | √ | |
| | OGFOD1 | √ | |
| | MYO1E | √ | |
| | TBCB | √ | |
| | CAPN2 | √ | |
| | PROSC | √ | |
| CCDC6-ANK3 | NR3C1 | √ | |
| | TRIM28 | √ | |
| | SF3A1 | √ | |
| | HNRNPR | √ | |
| | BRCC3 | √ | |
| | NR3C1 | √ | |
| | TRIM28 | √ | |
| | HNRNPR | √ | |
| | BRCC3 | √ | |
| | SF3A1 | √ | |
| CCDC9-DHX34 | EIF4A3 | √ | |
| | SNIP1 | √ | |
| | CCDC9 | √ | |
| | PRPF40A | √ | |
| CDC27-ST7L | ANAPC2 | √ | |
| | ANAPC7 | √ | |
| | TFDP1 | √ | |
| | UBE2S | √ | |
| | ANAPC2 | √ | |
| | ANAPC7 | √ | |
| CDK7-RIN3 | SUPT5H | √ | |
| | HNRNPU | | √ |
| | POLR2A | | √ |
| | POLR2A | | √ |
| | POLR2A | | √ |
| | POLR2A | | √ |
| | PRKCI | √ | |
| | APP | √ | |
| | CDC37 | √ | |
| CHERP-CPAMD8 | DHX8 | √ | |
| | PRPF40A | √ | |
| | CHERP | √ | |
| | SF3A2 | √ | |
| | DHX8 | √ | |
| | AGGF1 | √ | |
| | RNPS1 | √ | |
| | SNIP1 | √ | |
| | SF3B4 | √ | |
| | CHERP | √ | |
| | SRPK1 | √ | |
| | U2AF2 | √ | |

| | | | |
|---|---|---|---|
| | APBB1 | √ | |
| | PRPF40A | √ | |
| | TTC14 | √ | |
| | RBM23 | √ | |
| | SF3A2 | √ | |
| | SNRNP70 | √ | |
| | WBP4 | √ | |
| CYTH1-PRPSAP1 | DDX17 | √ | |
| | COPS5 | | √ |
| | CYTH1 | √ | |
| | ARRB2 | √ | |
| | ARRB1 | √ | |
| | FBXW11 | √ | |
| | DDX17 | √ | |
| | ITGB2 | √ | |
| | COPS5 | | √ |
| DLG1-CRYBG3 | LIN7A | √ | |
| | LIN7C | √ | |
| | APBA1 | √ | √ |
| | CASK | √ | |
| | CASK | √ | |
| | KHDRBS1 | √ | |
| | ARRB2 | √ | |
| | ARRB1 | √ | |
| DTX4-CCDC102B | MCM7 | √ | |
| | LENG1 | √ | |
| | TRIM54 | √ | |
| | MARK1 | √ | |
| | CCDC102B | √ | |
| EHD4-FSIP1 | EHD4 | √ | |
| | CTPS2 | √ | |
| | EHD1 | √ | |
| | WARS | √ | |
| | UBA2 | √ | |
| | ADSL | √ | |
| | UQCRC2 | √ | |
| | EHD4 | √ | |
| ELK4-SLC26A9 | | | |
| ERAL1-DIDO1 | HNRNPDL | √ | |
| | HNRNPK | √ | |
| | CUL3 | √ | |
| | APP | √ | |
| | CUL3 | √ | |
| GMDS-CCND3 | PPP1CC | √ | √ |
| | POLD1 | √ | |
| | GMDS | √ | |
| | NSFL1C | √ | |
| | CTH | √ | |
| | CAPN2 | √ | |
| | MCM10 | √ | |
| | APP | √ | |
| HJURP-EIF4E2 | FBXW11 | √ | |
| | GIGYF2 | √ | |
| | APP | √ | |

| | | | |
|---|---|---|---|
| | EIF4E2 | √ | |
| | YWHAB | √ | |
| | SHMT2 | | √ |
| INTS4-GAB2 | ZAP70 | √ | |
| KDM5A-ANO2 | TBP | √ | |
| | MORF4L1 | √ | |
| MAPK10-FAM13A | HDAC9 | √ | |
| | RELA | √ | |
| | APP | √ | |
| MAPRE1-TM9SF4 | YWHAZ | √ | |
| | APP | √ | |
| | TUBB | √ | |
| | VCAM1 | | √ |
| | UNK | √ | √ |
| | COPS5 | | √ |
| | CDK5RAP2 | √ | |
| | PRKACA | √ | |
| | PRKACB | √ | |
| | TUBB | √ | |
| | TUBA1A | √ | |
| | PRKACA | √ | |
| | PRKACB | √ | |
| | CDK5RAP2 | √ | |
| | TERF1 | √ | √ |
| | SPTAN1 | √ | |
| NUMB-ALDH6A1 | ITCH | √ | |
| | AP2A1 | √ | |
| | PRKCZ | √ | |
| | APP | √ | |
| PARD6B-CD48 | PRKCI | √ | |
| | PARD3 | √ | |
| | PARD6G | √ | |
| | APP | √ | |
| | PARD6B | √ | |
| | PARD6A | √ | |
| | YWHAH | √ | |
| | PRKCZ | √ | |
| | WWC1 | √ | |
| | PRKCI | √ | |
| | PARD3 | √ | |
| | PARD6G | √ | |
| | APP | √ | |
| | PARD6B | √ | |
| | PARD6A | √ | |
| | YWHAH | √ | |
| | PRKCZ | √ | |
| | PARD6G | √ | |
| | PARD6B | √ | |
| | PARD6A | √ | |
| PPP1R12A-MGAT4C | KDM1A | √ | √ |
| | PPP1R12A | √ | |
| | KDM1A | √ | √ |
| | PUS1 | √ | |
| | AARSD1 | √ | |

| | | | |
|---|---|---|---|
| | RPRD1B | √ | |
| | PPP1R12A | √ | |
| | TRIM47 | √ | |
| | PAXIP1 | √ | √ |
| | ACTR3 | √ | |
| RNF11-C8A | ITCH | √ | |
| | EPN1 | √ | |
| | RABGEF1 | √ | |
| | UBE2E1 | √ | √ |
| | UBE2D3 | √ | √ |
| | UBE2E3 | √ | |
| | HGS | √ | √ |
| | GGA3 | √ | |
| | GGA2 | √ | |
| | AP2A1 | √ | |
| | EPN3 | √ | |
| | UBE2D1 | √ | √ |
| | AP2B1 | √ | |
| | SMURF1 | √ | |
| | SMURF2 | √ | |
| | UBQLN2 | √ | |
| | STAM2 | √ | |
| | UBQLN4 | √ | √ |
| | APP | √ | |
| | PSMD4 | √ | |
| | PSMD7 | √ | |
| | PSMD6 | √ | |
| | PSMD11 | √ | |
| | PSMD10 | √ | |
| | PSMD3 | √ | |
| | PSMD12 | √ | |
| | PSMD13 | √ | |
| | USP14 | √ | |
| | PSMD14 | √ | |
| | PSMD1 | √ | |
| | APP | √ | |
| | GGA3 | √ | |
| | GGA2 | √ | |
| | APP | √ | |
| | TBK1 | √ | |
| | APP | √ | |
| SIPA1L3-WDR62 | WDR62 | √ | |
| | YWHAB | √ | |
| | SIPA1L3 | √ | |
| | YWHAQ | √ | |
| | YWHAB | √ | |
| | FBXW11 | √ | |
| | WDR62 | √ | |
| | TBP | √ | |
| SLC26A6-PRKAR2A | AKAP7 | √ | |
| | PRKAR2A | √ | |
| | PRKAR2B | √ | |
| | PRKACA | √ | |
| | PRKACB | √ | |

| | | | |
|---|---|---|---|
| | PRKAR2A | √ | |
| | AKAP7 | √ | |
| | PRKACA | √ | |
| | PRKACB | √ | |
| | PRKAR2B | √ | |
| | GCH1 | √ | |
| ST14-APLP2 | APLP2 | √ | |
| | HDAC5 | √ | |
| | APLP2 | √ | |
| | APBB1 | √ | |
| | APBB2 | √ | |
| | APLP2 | √ | |
| STRADB-NOP58 | NIFK | √ | |
| | NOP56 | √ | |
| | HNRNPU | | √ |
| | SNU13 | √ | |
| | PUM3 | √ | |
| | RPS15A | √ | |
| | RSL1D1 | √ | |
| | KRR1 | √ | |
| | DDX18 | √ | |
| | EIF6 | √ | |
| | DHX15 | √ | |
| | RPS4X | √ | |
| | PRPF3 | √ | |
| | TARDBP | √ | |
| | EIF2S2 | √ | |
| | DDX27 | √ | |
| | NOP58 | √ | |
| | DDX24 | √ | |
| | RPL30 | √ | |
| | DDX56 | √ | |
| | FTSJ3 | √ | |
| | DDX47 | √ | |
| | KPNA6 | √ | |
| | WDR36 | √ | |
| | KPNA1 | √ | |
| | RRP12 | √ | |
| | FBL | √ | |
| | TBL3 | √ | |
| | WDR36 | √ | |
| | NOP58 | √ | |
| | DHX15 | √ | |
| | NOP56 | √ | |
| STX16-RAE1 | FBXW11 | √ | |
| | NXF1 | √ | |
| | FAF1 | √ | |
| | RAE1 | √ | |
| | HNRNPUL1 | √ | |
| | CUL3 | √ | |
| | NXF1 | √ | |
| | CUL3 | √ | |
| TANC2-CHD6 | ZFYVE9 | √ | |
| | PPP1CC | √ | √ |

| | | | |
|---|---|---|---|
| | TANC2 | √ | |
| | ZFYVE9 | √ | |
| | PPP1CC | √ | √ |
| | TANC2 | √ | |
| | MAEA | √ | |
| | MKLN1 | √ | |
| | MMP7 | √ | |
| | RMND5A | √ | |
| | MAEA | √ | |
| | MKLN1 | √ | |
| | RANBP10 | √ | |
| | MMP7 | √ | |
| | RMND5A | √ | |
| TMPRSS2-ERG | CDC5L | | √ |
| | DDX3X | √ | |
| | PRKDC | √ | |
| | POLR2A | | √ |
| | SF3B2 | √ | |
| | XRCC5 | | √ |
| | XRCC6 | | √ |
| | DDX23 | √ | |
| | SNRNP200 | √ | |
| | DDX21 | √ | |
| | TUBB | √ | |
| | HNRNPU | | √ |
| | PRPF40A | √ | |
| | AR | √ | |
| | NCL | √ | |
| | HNRNPM | √ | |
| | HNRNPC | √ | |
| | ILF2 | √ | |
| | SF3B2 | √ | |
| | PRKDC | √ | |

Table S9: List of fusions and parental proteins

| LK,LY,M,GL | SC | CA |
|---|---|---|
| BCR | ASPSCR1 | ANK3 |
| ABL1 | TFE3 | USP9Y |
| BCR-ABL1 | ASPSCR1-TFE3 | ANK3-USP9Y |
| RUNX1 | ASTN2 | ARGLU1 |
| RUNX1T1 | CNOT2 | CXCR4 |
| RUNX1-RUNX1T1 | ASTN2-CNOT2 | ARGLU1-CXCR4 |
| KMT2A | BCOR | ATXN10 |
| MLLT10 | ZC3H7B | FBLN1 |
| KMT2A-MLLT10 | BCOR-ZC3H7B | ATXN10-FBLN1 |
| IGH | BCOR | BCAS3 |
| BCL2 | CCNB3 | NFS1 |
| IGH-BCL2 | BCOR-CCNB3 | BCAS3-NFS1 |

| | | |
|---|---|---|
| KMT2A | CDX1 | BCAS4 |
| AFF1 | IRF2BP2 | BCAS3 |
| KMT2A-AFF1 | CDX1-IRF2BP2 | BCAS4-BCAS3 |
| PICALM | CIC | BCL2L12 |
| MLLT10 | DUX4 | PRMT1 |
| PICALM-MLLT10 | CIC-DUX4 | BCL2L12-PRMT1 |
| PML | CREB1 | CAPNS1 |
| RARA | EWSR1 | WDR62 |
| PML-RARA | CREB1-EWSR1 | CAPNS1-WDR62 |
| KMT2A | CTDSP2 | CCDC6 |
| MLLT3 | FAM19A2 | ANK3 |
| KMT2A-MLLT3 | CTDSP2-FAM19A2 | CCDC6-ANK3 |
| KMT2A | CXorf67 | CCDC9 |
| AFDN | MBTD1 | DHX34 |
| KMT2A-AFDN | CXorf67-MBTD1 | CCDC9-DHX34 |
| CBFB | EPC1 | CDC27 |
| MYH11 | PHF1 | ST7L |
| CBFB-MYH11 | EPC1-PHF1 | CDC27-ST7L |
| IGH | ERG | CDK7 |
| MYC | EWSR1 | RIN3 |
| IGH-MYC | ERG-EWSR1 | CDK7-RIN3 |
| NUP98 | ETV6 | CHERP |
| DDX10 | NTRK3 | CPAMD8 |
| NUP98-DDX10 | ETV6-NTRK3 | CHERP-CPAMD8 |
| PCM1 | EWSR1 | CPD |
| JAK2 | ATF1 | PIGW |
| PCM1-JAK2 | EWSR1-ATF1 | CPD-PIGW |
| KMT2A | EWSR1 | CYTH1 |
| SEPT9 | FLI1 | PRPSAP1 |
| KMT2A-SEPT9 | EWSR1-FLI1 | CYTH1-PRPSAP1 |
| FUS | EWSR1 | DLG1 |
| ERG | NR4A3 | CRYBG3 |
| FUS-ERG | EWSR1-NR4A3 | DLG1-CRYBG3 |
| NUP98 | EWSR1 | DTX4 |
| HOXA9 | ETV4 | CCDC102B |
| NUP98-HOXA9 | EWSR1-ETV4 | DTX4-CCDC102B |
| ETV6 | EWSR1 | EHD4 |
| ABL1 | PATZ1 | FSIP1 |
| ETV6-ABL1 | EWSR1-PATZ1 | EHD4-FSIP1 |
| SET | EWSR1 | ELK4 |
| NUP214 | DDIT3 | SLC26A9 |
| SET-NUP214 | EWSR1-DDIT3 | ELK4-SLC26A9 |
| MNX1 | EWSR1 | EMID1 |
| ETV6 | POU5F1 | CBY1 |
| MNX1-ETV6 | EWSR1-POU5F1 | EMID1-CBY1 |
| KMT2A | EWSR1 | EPHA6 |
| MLLT1 | SP3 | CNTN6 |
| KMT2A-MLLT1 | EWSR1-SP3 | EPHA6-CNTN6 |
| KMT2A | FOXO4 | ERAL1 |
| MLLT6 | CIC | DIDO1 |

| | | |
|---|---|---|
| KMT2A-MLLT6 | FOXO4-CIC | ERAL1-DIDO1 |
| ETV6 | FUS | GMDS |
| ACSL6 | ERG | CCND3 |
| ETV6-ACSL6 | FUS-ERG | GMDS-CCND3 |
| ETV6 | FUS | HJURP |
| MECOM | CREB3L1 | EIF4E2 |
| ETV6-MECOM | FUS-CREB3L1 | HJURP-EIF4E2 |
| KMT2A | FUS | INTS4 |
| EPS15 | DDIT3 | GAB2 |
| KMT2A-EPS15 | FUS-DDIT3 | INTS4-GAB2 |
| KMT2A | FUS | KCNQ5 |
| GAS7 | ATF1 | RIMS1 |
| KMT2A-GAS7 | FUS-ATF1 | KCNQ5-RIMS1 |
| KMT2A | FUS | KDM5A |
| ABI1 | CREB3L2 | ANO2 |
| KMT2A-ABI1 | FUS-CREB3L2 | KDM5A-ANO2 |
| KMT2A | HEY1 | LAMA5 |
| MLLT11 | NCOA2 | C12orf28 |
| KMT2A-MLLT11 | HEY1-NCOA2 | LAMA5-C12orf28 |
| MN1 | IRX2 | MAPK10 |
| ETV6 | TERT | FAM13A |
| MN1-ETV6 | IRX2-TERT | MAPK10-FAM13A |
| KMT2A | JAZF1 | MAPRE1 |
| MAML2 | SUZ12 | TM9SF4 |
| KMT2A-MAML2 | JAZF1-SUZ12 | MAPRE1-TM9SF4 |
| KMT2A | JAZF1 | NCKAP5 |
| FOXO4 | PHF1 | MZT2A |
| KMT2A-FOXO4 | JAZF1-PHF1 | NCKAP5-MZT2A |
| DEK | LMNA | NUMB |
| NUP214 | NTRK1 | ALDH6A1 |
| DEK-NUP214 | LMNA-NTRK1 | NUMB-ALDH6A1 |
| RUNX1 | MEAF6 | PARD6B |
| CBFA2T3 | TRERF1 | CD48 |
| RUNX1-CBFA2T3 | MEAF6-TRERF1 | PARD6B-CD48 |
| NUP98 | MEAF6 | PPP1R12A |
| PSIP1 | PHF1 | MGAT4C |
| NUP98-PSIP1 | MEAF6-PHF1 | PPP1R12A-MGAT4C |
| NUP98 | NR4A3 | RNF11 |
| HOXC13 | TAF15 | C8A |
| NUP98-HOXC13 | NR4A3-TAF15 | RNF11-C8A |
| NUP98 | NR4A3 | SIPA1L3 |
| HOXC11 | TFG | WDR62 |
| NUP98-HOXC11 | NR4A3-TFG | SIPA1L3-WDR62 |
| PAX5 | NR6A1 | SLC26A6 |
| ETV6 | TRHDE | PRKAR2A |
| PAX5-ETV6 | NR6A1-TRHDE | SLC26A6-PRKAR2A |
| NUP98 | NUP107 | ST14 |
| HOXA11 | LGR5 | APLP2 |
| NUP98-HOXA11 | NUP107-LGR5 | ST14-APLP2 |
| BCR | PAPPA | STRADB |

| | | |
|---|---|---|
| PDGFRA | NUP107 | NOP58 |
| BCR-PDGFRA | PAPPA-NUP107 | STRADB-NOP58 |
| BCR | PAX3 | STX16 |
| FGFR1 | FOXO1 | RAE1 |
| BCR-FGFR1 | PAX3-FOXO1 | STX16-RAE1 |
| NPM1 | PAX7 | TANC2 |
| RARA | FOXO1 | CHD6 |
| NPM1-RARA | PAX7-FOXO1 | TANC2-CHD6 |
| KMT2A | SS18 | THSD7B |
| CBL | SSX2 | DARS |
| KMT2A-CBL | SS18-SSX2 | THSD7B-DARS |
| IGH | SS18 | TMEM123 |
| BCL6 | SSX1 | MMP7 |
| IGH-BCL6 | SS18-SSX1 | TMEM123-MMP7 |
| LCP1 | SS18L1 | TMPRSS2 |
| BCL6 | SSX1 | ERG |
| LCP1-BCL6 | SS18L1-SSX1 | TMPRSS2-ERG |
| CREBBP | SSX1 | TOX3 |
| KAT6A | SYT4 | CNTN5 |
| CREBBP-KAT6A | SSX1-SYT4 | TOX3-CNTN5 |
| KMT2A | TGFBR3 | UBR2 |
| ARHGAP26 | MGEA5 | XPO5 |
| KMT2A-ARHGAP26 | TGFBR3-MGEA5 | UBR2-XPO5 |
| FOXO3 | TRIO | UVRAG |
| KMT2A | TERT | INTS4 |
| FOXO3-KMT2A | TRIO-TERT | UVRAG-INTS4 |
| KMT2A | WDR70 | WNT11 |
| DCPS | RCOR1 | TSPAN8 |
| KMT2A-DCPS | WDR70-RCOR1 | WNT11-TSPAN8 |
| KMT2A | YWHAE | XRCC5 |
| EP300 | NUTM2B | ACADL |
| KMT2A-EP300 | YWHAE-NUTM2B | XRCC5-ACADL |
| IGH | YWHAE | ZCCHC7 |
| CEBPE | NUTM2A-AS1 | PRSS3 |
| IGH-CEBPE | YWHAE-NUTM2A-AS1 | ZCCHC7-PRSS3 |
| HSP90AA1 | YWHAE | ZFP91 |
| BCL6 | NUTM2A | NOX4 |
| HSP90AA1-BCL6 | YWHAE-NUTM2A | ZFP91-NOX4 |

Table S10: List of PAS

| **LK,LY,ME,GL** | Proteins | PAS | PAS/D | D/PAS |
|---|---|---|---|---|
| RUNX1-RUNX1T1 | HDAC1 | 119.739 | 0.2505 | 3.992 |
| | BRCA1 | 94.4444 | 5.55556 | 0.18 |
| | KMT2A | 9.00695 | 0.02286 | 43.744 |
| | SMARCC1 | 103.757 | 0.89445 | 1.118 |
| | HDAC2 | 80.3095 | 0.26681 | 3.748 |
| | CREBBP | 139.491 | 0.47125 | 2.122 |
| | CTBP1 | 98.5353 | 0.66578 | 1.502 |
| | EP300 | 113.477 | 0.2505 | 3.992 |

| | | | | |
|---|---|---|---|---|
| | SMARCA4 | 28.4738 | 0.14237 | 7.024 |
| | NCOR2 | 3.28906 | 0.03163 | 31.62 |
| | NCOR1 | 4.69557 | 0.0311 | 32.158 |
| | VDR | 119.946 | 1.34771 | 0.742 |
| | SMARCA4 | 28.4738 | 0.14237 | 7.024 |
| KMT2A-MLLT10 | SMARCC2 | 103.531 | 0.80257 | 1.246 |
| | KMT2A | 9.00695 | 0.02286 | 43.744 |
| | HDAC2 | 80.3095 | 0.26681 | 3.748 |
| | CHD3 | 5.7671 | 0.0395 | 25.316 |
| | SMARCC1 | 103.757 | 0.89445 | 1.118 |
| | POLR2A | 101.291 | 0.40355 | 2.478 |
| | SMARCA2 | 3.16923 | 0.03077 | 32.5 |
| | CREBBP | 139.491 | 0.47125 | 2.122 |
| | SIN3A | 6.12284 | 0.03189 | 31.358 |
| | CTBP1 | 98.5353 | 0.66578 | 1.502 |
| | KMT2A | 9.00695 | 0.02286 | 43.744 |
| | CREBBP | 139.491 | 0.47125 | 2.122 |
| IGH-BCL2 | TP53 | 240.49 | 0.25025 | 3.996 |
| | CASP3 | 92.4453 | 0.99404 | 1.006 |
| | PARP1 | 88.8078 | 0.40552 | 2.466 |
| | CASP8 | 172.68 | 1.28866 | 0.776 |
| | HIF1A | 97.973 | 0.67568 | 1.48 |
| | BCL2 | 5.70804 | 0.06072 | 16.468 |
| | BAG3 | 7.46766 | 0.01667 | 59.992 |
| | PARP1 | 88.8078 | 0.40552 | 2.466 |
| | BCL2 | 5.70804 | 0.06072 | 16.468 |
| KMT2A-AFF1 | SMARCC2 | 103.531 | 0.80257 | 1.246 |
| | KMT2A | 9.00695 | 0.02286 | 43.744 |
| | HDAC2 | 80.3095 | 0.26681 | 3.748 |
| | CHD3 | 5.7671 | 0.0395 | 25.316 |
| | SMARCC1 | 103.757 | 0.89445 | 1.118 |
| | POLR2A | 101.291 | 0.40355 | 2.478 |
| | SMARCA2 | 3.16923 | 0.03077 | 32.5 |
| | CREBBP | 139.491 | 0.47125 | 2.122 |
| | SIN3A | 6.12284 | 0.03189 | 31.358 |
| | CTBP1 | 98.5353 | 0.66578 | 1.502 |
| | KMT2A | 9.00695 | 0.02286 | 43.744 |
| | CREBBP | 139.491 | 0.47125 | 2.122 |
| PICALM-MLLT10 | FN1 | 105.505 | 4.58716 | 0.218 |
| | EEF1A1 | 158.526 | 0.42845 | 2.334 |
| | EGFR | 8.00992 | 0.00962 | 103.996 |
| | NTRK1 | 8.72582 | 0.05257 | 19.024 |
| | PLCG1 | 5.71137 | 0.05099 | 19.61 |
| | DNM2 | 5.98802 | 0.06805 | 14.696 |
| | ILVBL | 2.86994 | 0.07 | 14.286 |
| | PICALM | 3.01572 | 0.03388 | 29.512 |
| | HNRNPD | 6.06047 | 0.02253 | 44.386 |
| | FUS | 19.9375 | 0.0625 | 16 |
| | FN1 | 105.505 | 4.58716 | 0.218 |
| | DDX1 | 89.9433 | 0.70822 | 1.412 |
| | SEC24D | 2.43765 | 0.09376 | 10.666 |
| | PICALM | 3.01572 | 0.03388 | 29.512 |
| | NTRK1 | 8.72582 | 0.05257 | 19.024 |

| | | | | |
|---|---|---|---|---|
| | SEC24C | 3.28071 | 0.05126 | 19.508 |
| PML-RARA | NCOA2 | 3.38768 | 0.05293 | 18.892 |
| | NR3C1 | 6.22472 | 0.03965 | 25.222 |
| | NR4A1 | 185.771 | 1.97628 | 0.506 |
| | KAT2B | 6.53815 | 0.03736 | 26.766 |
| | RXRA | 4.57676 | 0.04199 | 23.816 |
| | PPARG | 5.04776 | 0.03883 | 25.754 |
| | TP53 | 240.49 | 0.25025 | 3.996 |
| | MDM2 | 9.93482 | 0.05257 | 19.024 |
| | EP300 | 113.477 | 0.2505 | 3.992 |
| | SMARCA4 | 28.4738 | 0.14237 | 7.024 |
| | RELA | 9.3566 | 0.05257 | 19.024 |
| | RARA | 5.00185 | 0.04631 | 21.592 |
| | NCOA3 | 3.96588 | 0.03741 | 26.728 |
| | NCOA1 | 3.6746 | 0.03638 | 27.486 |
| | STAT3 | 19.766 | 0.08864 | 11.282 |
| | ARNT | 4.42664 | 0.1054 | 9.488 |
| | NPAS2 | 83.3333 | 3.20513 | 0.312 |
| | PARP1 | 88.8078 | 0.40552 | 2.466 |
| | NFKB1 | 23.3366 | 0.16551 | 6.042 |
| | TRIP4 | 1.76435 | 0.04303 | 23.238 |
| | CREBBP | 139.491 | 0.47125 | 2.122 |
| | NFKB1 | 23.3366 | 0.16551 | 6.042 |
| | EP300 | 113.477 | 0.2505 | 3.992 |
| | PARP1 | 88.8078 | 0.40552 | 2.466 |
| KMT2A-MLLT3 | SMARCC2 | 103.531 | 0.80257 | 1.246 |
| | KMT2A | 9.00695 | 0.02286 | 43.744 |
| | HDAC2 | 80.3095 | 0.26681 | 3.748 |
| | CHD3 | 5.7671 | 0.0395 | 25.316 |
| | SMARCC1 | 103.757 | 0.89445 | 1.118 |
| | POLR2A | 101.291 | 0.40355 | 2.478 |
| | SMARCA2 | 3.16923 | 0.03077 | 32.5 |
| | CREBBP | 139.491 | 0.47125 | 2.122 |
| | SIN3A | 6.12284 | 0.03189 | 31.358 |
| | CTBP1 | 98.5353 | 0.66578 | 1.502 |
| | KMT2A | 9.00695 | 0.02286 | 43.744 |
| | CREBBP | 139.491 | 0.47125 | 2.122 |
| KMT2A-AFDN | SMARCC2 | 103.531 | 0.80257 | 1.246 |
| | KMT2A | 9.00695 | 0.02286 | 43.744 |
| | HDAC2 | 80.3095 | 0.26681 | 3.748 |
| | CHD3 | 5.7671 | 0.0395 | 25.316 |
| | SMARCC1 | 103.757 | 0.89445 | 1.118 |
| | POLR2A | 101.291 | 0.40355 | 2.478 |
| | SMARCA2 | 3.16923 | 0.03077 | 32.5 |
| | CREBBP | 139.491 | 0.47125 | 2.122 |
| | SIN3A | 6.12284 | 0.03189 | 31.358 |
| | CTBP1 | 98.5353 | 0.66578 | 1.502 |
| | KMT2A | 9.00695 | 0.02286 | 43.744 |
| | CREBBP | 139.491 | 0.47125 | 2.122 |
| CBFB-MYH11 | ACTA2 | 4.39678 | 0.05362 | 18.65 |
| | MYH11 | 4.45976 | 0.10136 | 9.866 |
| | MYO1E | 2.45997 | 0.04473 | 22.358 |
| | RPA1 | 115.982 | 0.2505 | 3.992 |
| | RPA2 | 11.8797 | 0.05257 | 19.024 |

| | | | | |
|---|---|---|---|---|
| | ACTB | 5.97407 | 0.02679 | 37.328 |
| | RPA1 | 115.982 | 0.2505 | 3.992 |
| | ELAVL1 | 5.70617 | 0.02679 | 37.328 |
| | RPA2 | 11.8797 | 0.05257 | 19.024 |
| NUP98-DDX10 | SIRT7 | 9.04121 | 0.05257 | 19.024 |
| | DDX10 | 2.75895 | 0.02172 | 46.032 |
| | APP | 6.2537 | 0.037 | 27.024 |
| | DDX56 | 1.94682 | 0.02374 | 42.12 |
| | NTRK1 | 8.72582 | 0.05257 | 19.024 |
| | DDX54 | 3.01289 | 0.07007 | 14.272 |
| | PUM3 | 1.32572 | 0.0204 | 49.03 |
| | PWP1 | 1.48662 | 0.0413 | 24.216 |
| | CSNK2A1 | 8.68379 | 0.02273 | 43.99 |
| | HDAC1 | 119.739 | 0.2505 | 3.992 |
| | CTNNB1 | 7.85024 | 0.02745 | 36.432 |
| | MAPK8 | 9.04121 | 0.05257 | 19.024 |
| | EP300 | 113.477 | 0.2505 | 3.992 |
| | CREBBP | 139.491 | 0.47125 | 2.122 |
| | PUM3 | 1.32572 | 0.0204 | 49.03 |
| | NXF1 | 1.24414 | 0.0204 | 49.03 |
| | EED | 197.605 | 2.99401 | 0.334 |
| | KPNB1 | 6.2426 | 0.02959 | 33.8 |
| | HNRNPUL1 | 3.24977 | 0.02902 | 34.464 |
| | HDAC1 | 119.739 | 0.2505 | 3.992 |
| | CREBBP | 139.491 | 0.47125 | 2.122 |
| | APC | 12.2289 | 0.07279 | 13.738 |
| | CSNK2A1 | 8.68379 | 0.02273 | 43.99 |
| | CTNNB1 | 7.85024 | 0.02745 | 36.432 |
| | APC | 12.2289 | 0.07279 | 13.738 |
| | CREBBP | 139.491 | 0.47125 | 2.122 |
| | USP7 | 1.76435 | 0.04303 | 23.238 |
| | NTRK1 | 8.72582 | 0.05257 | 19.024 |
| | CDC37 | 4.10142 | 0.09321 | 10.728 |
| PCM1-JAK2 | ERBB2 | 2.49512 | 0.10848 | 9.218 |
| | ERBB3 | 1.33581 | 0.05808 | 17.218 |
| | VAV1 | 2.1312 | 0.05198 | 19.238 |
| | TEC | 6.57728 | 0.04216 | 23.718 |
| | EGFR | 8.00992 | 0.00962 | 103.996 |
| | INSR | 2.24215 | 0.05469 | 18.286 |
| | PLCG1 | 5.71137 | 0.05099 | 19.61 |
| | JAK2 | 1.44948 | 0.03535 | 28.286 |
| | STAT5A | 16.7896 | 0.07529 | 13.282 |
| | JAK2 | 1.44948 | 0.03535 | 28.286 |
| | INSR | 2.24215 | 0.05469 | 18.286 |
| KMT2A-SEPT9 | SMARCC2 | 103.531 | 0.80257 | 1.246 |
| | KMT2A | 9.00695 | 0.02286 | 43.744 |
| | HDAC2 | 80.3095 | 0.26681 | 3.748 |
| | CHD3 | 5.7671 | 0.0395 | 25.316 |
| | SMARCC1 | 103.757 | 0.89445 | 1.118 |
| | POLR2A | 101.291 | 0.40355 | 2.478 |
| | SMARCA2 | 3.16923 | 0.03077 | 32.5 |
| | CREBBP | 139.491 | 0.47125 | 2.122 |
| | SIN3A | 6.12284 | 0.03189 | 31.358 |
| | CTBP1 | 98.5353 | 0.66578 | 1.502 |

| | | | | |
|---|---|---|---|---|
| | KMT2A | 9.00695 | 0.02286 | 43.744 |
| | CREBBP | 139.491 | 0.47125 | 2.122 |
| FUS-ERG | RPA1 | 115.982 | 0.2505 | 3.992 |
| | SF3B2 | 1.77281 | 0.06818 | 14.666 |
| | PRKDC | 1.01978 | 0.0204 | 49.03 |
| | PRPF8 | 4.5895 | 0.05099 | 19.61 |
| | SF3A2 | 2.05274 | 0.07895 | 12.666 |
| | DHX15 | 1.94682 | 0.02374 | 42.12 |
| | RPA2 | 11.8797 | 0.05257 | 19.024 |
| | CUL3 | 3.3748 | 0.04821 | 20.742 |
| | ABL1 | 5.80459 | 0.03355 | 29.804 |
| | PARP1 | 88.8078 | 0.40552 | 2.466 |
| | PRKDC | 1.01978 | 0.0204 | 49.03 |
| NUP98-HOXA9 | HDAC1 | 119.739 | 0.2505 | 3.992 |
| | TP53 | 240.49 | 0.25025 | 3.996 |
| | SMAD4 | 25.6219 | 0.24876 | 4.02 |
| | CTNNB1 | 7.85024 | 0.02745 | 36.432 |
| | CSNK2A1 | 8.68379 | 0.02273 | 43.99 |
| | MAPK8 | 9.04121 | 0.05257 | 19.024 |
| | EP300 | 113.477 | 0.2505 | 3.992 |
| | CREBBP | 139.491 | 0.47125 | 2.122 |
| | CTNNB1 | 7.85024 | 0.02745 | 36.432 |
| | APC | 12.2289 | 0.07279 | 13.738 |
| | CREBBP | 139.491 | 0.47125 | 2.122 |
| ETV6-ABL1 | ERBB2 | 2.49512 | 0.10848 | 9.218 |
| | CBLB | 3.45695 | 0.07857 | 12.728 |
| | UBASH3B | 4.44644 | 0.04079 | 24.514 |
| | SOS1 | 2.36839 | 0.05383 | 18.578 |
| | ERBB4 | 18.8834 | 0.82102 | 1.218 |
| | SRC | 4.15958 | 0.09454 | 10.578 |
| | VAV1 | 2.1312 | 0.05198 | 19.238 |
| | EGFR | 8.00992 | 0.00962 | 103.996 |
| | CBL | 19.2785 | 0.08274 | 12.086 |
| | PLCG1 | 5.71137 | 0.05099 | 19.61 |
| | PIK3R2 | 3.9234 | 0.03599 | 27.782 |
| | PIK3R1 | 5.51541 | 0.03804 | 26.29 |
| | ABL1 | 5.80459 | 0.03355 | 29.804 |
| | ABL1 | 5.80459 | 0.03355 | 29.804 |
| | ABL2 | 5.13354 | 0.03355 | 29.804 |
| | JAK1 | 1.44948 | 0.03535 | 28.286 |
| SET-NUP214 | NXF1 | 1.24414 | 0.0204 | 49.03 |
| | FAF1 | 4.40782 | 0.19164 | 5.218 |
| | SUPT5H | 5.13901 | 0.08425 | 11.87 |
| | GART | 1.93343 | 0.06905 | 14.482 |
| | CUL2 | 138.926 | 0.16678 | 5.996 |
| | CUL3 | 3.3748 | 0.04821 | 20.742 |
| | NXF1 | 1.24414 | 0.0204 | 49.03 |
| | CUL2 | 138.926 | 0.16678 | 5.996 |
| | RANBP2 | 4.07955 | 0.05099 | 19.61 |
| MNX1-ETV6 | HDAC3 | 22.5933 | 0.98232 | 1.018 |
| | ETV6 | 37.2168 | 1.61812 | 0.618 |
| | HDAC9 | 2.52262 | 0.0174 | 57.48 |
| | PIN1 | 102.388 | 0.45914 | 2.178 |
| | NCOR1 | 4.69557 | 0.0311 | 32.158 |

| | | | | |
|---|---|---|---|---|
| | SIN3A | 6.12284 | 0.03189 | 31.358 |
| | L3MBTL1 | 3.92193 | 0.01859 | 53.8 |
| | ETV7 | 3.18648 | 0.13854 | 7.218 |
| | ETV6 | 37.2168 | 1.61812 | 0.618 |
| KMT2A-MLLT1 | SMARCC2 | 103.531 | 0.80257 | 1.246 |
| | KMT2A | 9.00695 | 0.02286 | 43.744 |
| | HDAC2 | 80.3095 | 0.26681 | 3.748 |
| | CHD3 | 5.7671 | 0.0395 | 25.316 |
| | SMARCC1 | 103.757 | 0.89445 | 1.118 |
| | POLR2A | 101.291 | 0.40355 | 2.478 |
| | SMARCA2 | 3.16923 | 0.03077 | 32.5 |
| | CREBBP | 139.491 | 0.47125 | 2.122 |
| | SIN3A | 6.12284 | 0.03189 | 31.358 |
| | CTBP1 | 98.5353 | 0.66578 | 1.502 |
| | KMT2A | 9.00695 | 0.02286 | 43.744 |
| | CREBBP | 139.491 | 0.47125 | 2.122 |
| KMT2A-MLLT6 | SMARCC2 | 103.531 | 0.80257 | 1.246 |
| | KMT2A | 9.00695 | 0.02286 | 43.744 |
| | HDAC2 | 80.3095 | 0.26681 | 3.748 |
| | CHD3 | 5.7671 | 0.0395 | 25.316 |
| | SMARCC1 | 103.757 | 0.89445 | 1.118 |
| | POLR2A | 101.291 | 0.40355 | 2.478 |
| | SMARCA2 | 3.16923 | 0.03077 | 32.5 |
| | CREBBP | 139.491 | 0.47125 | 2.122 |
| | SIN3A | 6.12284 | 0.03189 | 31.358 |
| | CTBP1 | 98.5353 | 0.66578 | 1.502 |
| | KMT2A | 9.00695 | 0.02286 | 43.744 |
| | CREBBP | 139.491 | 0.47125 | 2.122 |
| ETV6-ACSL6 | HDAC3 | 22.5933 | 0.98232 | 1.018 |
| | ETV6 | 37.2168 | 1.61812 | 0.618 |
| | HDAC9 | 2.52262 | 0.0174 | 57.48 |
| | PIN1 | 102.388 | 0.45914 | 2.178 |
| | NCOR1 | 4.69557 | 0.0311 | 32.158 |
| | SIN3A | 6.12284 | 0.03189 | 31.358 |
| | L3MBTL1 | 3.92193 | 0.01859 | 53.8 |
| | ETV7 | 3.18648 | 0.13854 | 7.218 |
| | ETV6 | 37.2168 | 1.61812 | 0.618 |
| ETV6-MECOM | HDAC1 | 119.739 | 0.2505 | 3.992 |
| | UBE2I | 107.519 | 0.25063 | 3.99 |
| | HDAC3 | 22.5933 | 0.98232 | 1.018 |
| | EHMT2 | 9.91714 | 0.01191 | 83.996 |
| | ELAVL1 | 5.70617 | 0.02679 | 37.328 |
| | MECOM | 7.55275 | 0.03996 | 25.024 |
| | SMAD1 | 8.21988 | 0.04281 | 23.358 |
| | SMAD2 | 22.8889 | 0.22222 | 4.5 |
| | SMAD3 | 1.9619 | 0.01905 | 52.5 |
| | KAT2B | 6.53815 | 0.03736 | 26.766 |
| | SUV39H1 | 7.75095 | 0.12706 | 7.87 |
| | NCOR1 | 4.69557 | 0.0311 | 32.158 |
| | CTBP1 | 98.5353 | 0.66578 | 1.502 |
| | CREBBP | 139.491 | 0.47125 | 2.122 |
| | MECOM | 7.55275 | 0.03996 | 25.024 |
| | SUV39H1 | 7.75095 | 0.12706 | 7.87 |

| | | | | |
|---|---|---|---|---|
| | HDAC4 | 4.33094 | 0.02987 | 33.48 |
| KMT2A-EPS15 | SMARCC2 | 103.531 | 0.80257 | 1.246 |
| | KMT2A | 9.00695 | 0.02286 | 43.744 |
| | HDAC2 | 80.3095 | 0.26681 | 3.748 |
| | CHD3 | 5.7671 | 0.0395 | 25.316 |
| | SMARCC1 | 103.757 | 0.89445 | 1.118 |
| | POLR2A | 101.291 | 0.40355 | 2.478 |
| | SMARCA2 | 3.16923 | 0.03077 | 32.5 |
| | CREBBP | 139.491 | 0.47125 | 2.122 |
| | SIN3A | 6.12284 | 0.03189 | 31.358 |
| | CTBP1 | 98.5353 | 0.66578 | 1.502 |
| | KMT2A | 9.00695 | 0.02286 | 43.744 |
| | CREBBP | 139.491 | 0.47125 | 2.122 |
| KMT2A-GAS7 | SMARCC2 | 103.531 | 0.80257 | 1.246 |
| | KMT2A | 9.00695 | 0.02286 | 43.744 |
| | HDAC2 | 80.3095 | 0.26681 | 3.748 |
| | CHD3 | 5.7671 | 0.0395 | 25.316 |
| | SMARCC1 | 103.757 | 0.89445 | 1.118 |
| | POLR2A | 101.291 | 0.40355 | 2.478 |
| | SMARCA2 | 3.16923 | 0.03077 | 32.5 |
| | CREBBP | 139.491 | 0.47125 | 2.122 |
| | SIN3A | 6.12284 | 0.03189 | 31.358 |
| | CTBP1 | 98.5353 | 0.66578 | 1.502 |
| | KMT2A | 9.00695 | 0.02286 | 43.744 |
| | CREBBP | 139.491 | 0.47125 | 2.122 |
| KMT2A-ABL1 | SMARCC2 | 103.531 | 0.80257 | 1.246 |
| | KMT2A | 9.00695 | 0.02286 | 43.744 |
| | SMARCC1 | 103.757 | 0.89445 | 1.118 |
| | CHD3 | 5.7671 | 0.0395 | 25.316 |
| | POLR2A | 101.291 | 0.40355 | 2.478 |
| | SMARCA2 | 3.16923 | 0.03077 | 32.5 |
| | CREBBP | 139.491 | 0.47125 | 2.122 |
| | ABL1 | 5.80459 | 0.03355 | 29.804 |
| | CBLB | 3.45695 | 0.07857 | 12.728 |
| | CBL | 19.2785 | 0.08274 | 12.086 |
| KMT2A-MLLT11 | SMARCC2 | 103.531 | 0.80257 | 1.246 |
| | KMT2A | 9.00695 | 0.02286 | 43.744 |
| | HDAC2 | 80.3095 | 0.26681 | 3.748 |
| | CHD3 | 5.7671 | 0.0395 | 25.316 |
| | SMARCC1 | 103.757 | 0.89445 | 1.118 |
| | POLR2A | 101.291 | 0.40355 | 2.478 |
| | SMARCA2 | 3.16923 | 0.03077 | 32.5 |
| | CREBBP | 139.491 | 0.47125 | 2.122 |
| | SIN3A | 6.12284 | 0.03189 | 31.358 |
| | CTBP1 | 98.5353 | 0.66578 | 1.502 |
| | KMT2A | 9.00695 | 0.02286 | 43.744 |
| | CREBBP | 139.491 | 0.47125 | 2.122 |
| MN1-ETV6 | HDAC3 | 22.5933 | 0.98232 | 1.018 |
| | ETV6 | 37.2168 | 1.61812 | 0.618 |
| | HDAC9 | 2.52262 | 0.0174 | 57.48 |
| | PIN1 | 102.388 | 0.45914 | 2.178 |
| | EP300 | 113.477 | 0.2505 | 3.992 |
| | NCOR1 | 4.69557 | 0.0311 | 32.158 |
| | SIN3A | 6.12284 | 0.03189 | 31.358 |

| | | | | |
|---|---|---|---|---|
| | HDAC3 | 22.5933 | 0.98232 | 1.018 |
| | EP300 | 113.477 | 0.2505 | 3.992 |
| | KAT5 | 32.4274 | 0.15442 | 6.476 |
| KMT2A-MAML2 | SMARCC2 | 103.531 | 0.80257 | 1.246 |
| | KMT2A | 9.00695 | 0.02286 | 43.744 |
| | SMARCC1 | 103.757 | 0.89445 | 1.118 |
| | CHD3 | 5.7671 | 0.0395 | 25.316 |
| | SMARCA2 | 3.16923 | 0.03077 | 32.5 |
| | CREBBP | 139.491 | 0.47125 | 2.122 |
| | SMARCA2 | 3.16923 | 0.03077 | 32.5 |
| | EP300 | 113.477 | 0.2505 | 3.992 |
| | CREBBP | 139.491 | 0.47125 | 2.122 |
| KMT2A-FOXO4 | SMARCC2 | 103.531 | 0.80257 | 1.246 |
| | KMT2A | 9.00695 | 0.02286 | 43.744 |
| | HDAC2 | 80.3095 | 0.26681 | 3.748 |
| | CHD3 | 5.7671 | 0.0395 | 25.316 |
| | SMARCC1 | 103.757 | 0.89445 | 1.118 |
| | POLR2A | 101.291 | 0.40355 | 2.478 |
| | SMARCA2 | 3.16923 | 0.03077 | 32.5 |
| | CREBBP | 139.491 | 0.47125 | 2.122 |
| | SIN3A | 6.12284 | 0.03189 | 31.358 |
| | CTBP1 | 98.5353 | 0.66578 | 1.502 |
| | KMT2A | 9.00695 | 0.02286 | 43.744 |
| | CREBBP | 139.491 | 0.47125 | 2.122 |
| DEK-NUP214 | ESR1 | 28.1174 | 1.22249 | 0.818 |
| | KAT2B | 6.53815 | 0.03736 | 26.766 |
| | SMAD2 | 22.8889 | 0.22222 | 4.5 |
| | SMAD3 | 1.9619 | 0.01905 | 52.5 |
| | CDK2 | 14.4846 | 0.06217 | 16.086 |
| | DEK | 3.70705 | 0.04521 | 22.12 |
| | EP300 | 113.477 | 0.2505 | 3.992 |
| | CREBBP | 139.491 | 0.47125 | 2.122 |
| | NXF1 | 1.24414 | 0.0204 | 49.03 |
| | DHX15 | 1.94682 | 0.02374 | 42.12 |
| | CUL2 | 138.926 | 0.16678 | 5.996 |
| | CUL3 | 3.3748 | 0.04821 | 20.742 |
| RUNX1-CBFA2T3 | KMT2A | 9.00695 | 0.02286 | 43.744 |
| | SMARCC1 | 103.757 | 0.89445 | 1.118 |
| | SMARCA4 | 28.4738 | 0.14237 | 7.024 |
| | CREBBP | 139.491 | 0.47125 | 2.122 |
| | EP300 | 113.477 | 0.2505 | 3.992 |
| | CREBBP | 139.491 | 0.47125 | 2.122 |
| NUP98-PSIP1 | HDAC1 | 119.739 | 0.2505 | 3.992 |
| | KMT2A | 9.00695 | 0.02286 | 43.744 |
| | ESR1 | 28.1174 | 1.22249 | 0.818 |
| | CTNNB1 | 7.85024 | 0.02745 | 36.432 |
| | EP300 | 113.477 | 0.2505 | 3.992 |
| | CREBBP | 139.491 | 0.47125 | 2.122 |
| | NXF1 | 1.24414 | 0.0204 | 49.03 |
| | EIF4A3 | 6.71796 | 0.00806 | 123.996 |
| | SON | 3.01825 | 0.0686 | 14.578 |
| NUP98- | HDAC1 | 119.739 | 0.2505 | 3.992 |

| | | | | |
|---|---|---|---|---|
| HOXC13 | | | | |
| | CTNNB1 | 7.85024 | 0.02745 | 36.432 |
| | MAPK8 | 9.04121 | 0.05257 | 19.024 |
| | EP300 | 113.477 | 0.2505 | 3.992 |
| | CREBBP | 139.491 | 0.47125 | 2.122 |
| | NXF1 | 1.24414 | 0.0204 | 49.03 |
| | HNRNPAB | 2.52262 | 0.0174 | 57.48 |
| | EED | 197.605 | 2.99401 | 0.334 |
| | KPNB1 | 6.2426 | 0.02959 | 33.8 |
| | HNRNPUL1 | 3.24977 | 0.02902 | 34.464 |
| | HDAC1 | 119.739 | 0.2505 | 3.992 |
| | CREBBP | 139.491 | 0.47125 | 2.122 |
| | APC | 12.2289 | 0.07279 | 13.738 |
| | CSNK2A1 | 8.68379 | 0.02273 | 43.99 |
| | CTNNB1 | 7.85024 | 0.02745 | 36.432 |
| | APC | 12.2289 | 0.07279 | 13.738 |
| | CREBBP | 139.491 | 0.47125 | 2.122 |
| NUP98-HOXC11 | HDAC1 | 119.739 | 0.2505 | 3.992 |
| | STAT3 | 19.766 | 0.08864 | 11.282 |
| | SP1 | 3.01825 | 0.0686 | 14.578 |
| | SMAD3 | 1.9619 | 0.01905 | 52.5 |
| | CTNNB1 | 7.85024 | 0.02745 | 36.432 |
| | MAPK8 | 9.04121 | 0.05257 | 19.024 |
| | EP300 | 113.477 | 0.2505 | 3.992 |
| | CREBBP | 139.491 | 0.47125 | 2.122 |
| | CTNNB1 | 7.85024 | 0.02745 | 36.432 |
| | APC | 12.2289 | 0.07279 | 13.738 |
| | CREBBP | 139.491 | 0.47125 | 2.122 |
| PAX5-ETV6 | UBE2I | 107.519 | 0.25063 | 3.99 |
| | HDAC3 | 22.5933 | 0.98232 | 1.018 |
| | TBP | 6.06579 | 0.03888 | 25.718 |
| | KAT5 | 32.4274 | 0.15442 | 6.476 |
| | RB1 | 7.40059 | 0.0341 | 29.322 |
| | PAX5 | 22.5 | 2.5 | 0.4 |
| | RUNX1 | 2.91921 | 0.04293 | 23.294 |
| | PIN1 | 102.388 | 0.45914 | 2.178 |
| | EP300 | 113.477 | 0.2505 | 3.992 |
| | NCOR1 | 4.69557 | 0.0311 | 32.158 |
| | MAPK1 | 5.32418 | 0.02488 | 40.194 |
| | EP300 | 113.477 | 0.2505 | 3.992 |
| | HDAC6 | 3.86873 | 0.02668 | 37.48 |
| NUP98-HOXA11 | HDAC1 | 119.739 | 0.2505 | 3.992 |
| | HDAC2 | 80.3095 | 0.26681 | 3.748 |
| | YY1 | 4.46363 | 0.03689 | 27.108 |
| | CTNNB1 | 7.85024 | 0.02745 | 36.432 |
| | CSNK2A1 | 8.68379 | 0.02273 | 43.99 |
| | MAPK8 | 9.04121 | 0.05257 | 19.024 |
| | EP300 | 113.477 | 0.2505 | 3.992 |
| | CREBBP | 139.491 | 0.47125 | 2.122 |
| | CTNNB1 | 7.85024 | 0.02745 | 36.432 |
| | APC | 12.2289 | 0.07279 | 13.738 |
| | CREBBP | 139.491 | 0.47125 | 2.122 |
| BCR-PDGFRA | TGFBR2 | 7.20943 | 0.08792 | 11.374 |

| | | | | |
|---|---|---|---|---|
| | PDGFRA | 3.84 | 0.08 | 12.5 |
| | SHC1 | 7.13762 | 0.03335 | 29.982 |
| | CRKL | 6.10583 | 0.05654 | 17.688 |
| | EGFR | 8.00992 | 0.00962 | 103.996 |
| | PLCG1 | 5.71137 | 0.05099 | 19.61 |
| | FES | 2.49512 | 0.10848 | 9.218 |
| | CRK | 6.51925 | 0.04405 | 22.702 |
| | ABL1 | 5.80459 | 0.03355 | 29.804 |
| | HCK | 4.13571 | 0.0701 | 14.266 |
| | PTPN6 | 4.41336 | 0.0679 | 14.728 |
| | BCR | 15.6512 | 0.27949 | 3.578 |
| | UBASH3B | 4.44644 | 0.04079 | 24.514 |
| | INPP5D | 2.63514 | 0.06757 | 14.8 |
| | SOS1 | 2.36839 | 0.05383 | 18.578 |
| | GRB2 | 9.90537 | 0.01923 | 51.992 |
| | NTRK1 | 8.72582 | 0.05257 | 19.024 |
| | CBL | 19.2785 | 0.08274 | 12.086 |
| | KIT | 2.56536 | 0.06108 | 16.372 |
| | DOK1 | 98.8024 | 2.99401 | 0.334 |
| | PIK3R2 | 3.9234 | 0.03599 | 27.782 |
| | PIK3R1 | 5.51541 | 0.03804 | 26.29 |
| | ABL1 | 5.80459 | 0.03355 | 29.804 |
| | TP53 | 240.49 | 0.25025 | 3.996 |
| | RB1 | 7.40059 | 0.0341 | 29.322 |
| | BCR | 15.6512 | 0.27949 | 3.578 |
| BCR-FGFR1 | SRC | 4.15958 | 0.09454 | 10.578 |
| | ITK | 1.83972 | 0.04487 | 22.286 |
| | SOS1 | 2.36839 | 0.05383 | 18.578 |
| | ERBB3 | 1.33581 | 0.05808 | 17.218 |
| | VAV1 | 2.1312 | 0.05198 | 19.238 |
| | CBL | 19.2785 | 0.08274 | 12.086 |
| | PLCG1 | 5.71137 | 0.05099 | 19.61 |
| | ABL1 | 5.80459 | 0.03355 | 29.804 |
| | ABL1 | 5.80459 | 0.03355 | 29.804 |
| | HCK | 4.13571 | 0.0701 | 14.266 |
| | BCR | 15.6512 | 0.27949 | 3.578 |
| | CBL | 19.2785 | 0.08274 | 12.086 |
| IGH-BCL6 | HDAC1 | 119.739 | 0.2505 | 3.992 |
| | TP53 | 240.49 | 0.25025 | 3.996 |
| | CTBP1 | 98.5353 | 0.66578 | 1.502 |
| | EP300 | 113.477 | 0.2505 | 3.992 |
| | NCOR2 | 3.28906 | 0.03163 | 31.62 |
| | CREBBP | 139.491 | 0.47125 | 2.122 |
| | HDAC2 | 80.3095 | 0.26681 | 3.748 |
| | SMARCA4 | 28.4738 | 0.14237 | 7.024 |
| | CREBBP | 139.491 | 0.47125 | 2.122 |
| LCP1-BCL6 | HDAC1 | 119.739 | 0.2505 | 3.992 |
| | TP53 | 240.49 | 0.25025 | 3.996 |
| | CTBP1 | 98.5353 | 0.66578 | 1.502 |
| | EP300 | 113.477 | 0.2505 | 3.992 |
| | NCOR2 | 3.28906 | 0.03163 | 31.62 |
| | CREBBP | 139.491 | 0.47125 | 2.122 |
| | HDAC2 | 80.3095 | 0.26681 | 3.748 |
| | SMARCA4 | 28.4738 | 0.14237 | 7.024 |

| | | | | |
|---|---|---|---|---|
| | CREBBP | 139.491 | 0.47125 | 2.122 |
| CREBBP-KAT6A | SMARCC2 | 103.531 | 0.80257 | 1.246 |
| | KMT2A | 9.00695 | 0.02286 | 43.744 |
| | HDAC2 | 80.3095 | 0.26681 | 3.748 |
| | CHD3 | 5.7671 | 0.0395 | 25.316 |
| | SMARCC1 | 103.757 | 0.89445 | 1.118 |
| | POLR2A | 101.291 | 0.40355 | 2.478 |
| | SMARCA2 | 3.16923 | 0.03077 | 32.5 |
| | CREBBP | 139.491 | 0.47125 | 2.122 |
| | SIN3A | 6.12284 | 0.03189 | 31.358 |
| | CTBP1 | 98.5353 | 0.66578 | 1.502 |
| | KMT2A | 9.00695 | 0.02286 | 43.744 |
| | CREBBP | 139.491 | 0.47125 | 2.122 |
| KMT2A-ARHGAP26 | SMARCC2 | 103.531 | 0.80257 | 1.246 |
| | KMT2A | 9.00695 | 0.02286 | 43.744 |
| | HDAC2 | 80.3095 | 0.26681 | 3.748 |
| | CHD3 | 5.7671 | 0.0395 | 25.316 |
| | SMARCC1 | 103.757 | 0.89445 | 1.118 |
| | POLR2A | 101.291 | 0.40355 | 2.478 |
| | SMARCA2 | 3.16923 | 0.03077 | 32.5 |
| | CREBBP | 139.491 | 0.47125 | 2.122 |
| | SIN3A | 6.12284 | 0.03189 | 31.358 |
| | CTBP1 | 98.5353 | 0.66578 | 1.502 |
| | KMT2A | 9.00695 | 0.02286 | 43.744 |
| | CREBBP | 139.491 | 0.47125 | 2.122 |
| FOXO3-KMT2A | SMARCC2 | 103.531 | 0.80257 | 1.246 |
| | KMT2A | 9.00695 | 0.02286 | 43.744 |
| | SMARCC1 | 103.757 | 0.89445 | 1.118 |
| | CHD3 | 5.7671 | 0.0395 | 25.316 |
| | SMARCA2 | 3.16923 | 0.03077 | 32.5 |
| | CREBBP | 139.491 | 0.47125 | 2.122 |
| | SMARCA2 | 3.16923 | 0.03077 | 32.5 |
| | EP300 | 113.477 | 0.2505 | 3.992 |
| | CREBBP | 139.491 | 0.47125 | 2.122 |
| KMT2A-DCPS | SMARCC2 | 103.531 | 0.80257 | 1.246 |
| | KMT2A | 9.00695 | 0.02286 | 43.744 |
| | HDAC2 | 80.3095 | 0.26681 | 3.748 |
| | CHD3 | 5.7671 | 0.0395 | 25.316 |
| | SMARCC1 | 103.757 | 0.89445 | 1.118 |
| | POLR2A | 101.291 | 0.40355 | 2.478 |
| | SMARCA2 | 3.16923 | 0.03077 | 32.5 |
| | CREBBP | 139.491 | 0.47125 | 2.122 |
| | SIN3A | 6.12284 | 0.03189 | 31.358 |
| | CTBP1 | 98.5353 | 0.66578 | 1.502 |
| | KMT2A | 9.00695 | 0.02286 | 43.744 |
| | CREBBP | 139.491 | 0.47125 | 2.122 |
| IGH-CEBPE | UBE2I | 107.519 | 0.25063 | 3.99 |
| | BATF | 1.28936 | 0.06786 | 14.736 |
| | DDIT3 | 4.74834 | 0.06783 | 14.742 |
| | CEBPG | 1.22222 | 0.05556 | 18 |
| | CEBPE | 4.10142 | 0.09321 | 10.728 |
| | FOS | 7.41248 | 0.05702 | 17.538 |

| | | | | |
|---|---|---|---|---|
| | JUN | 6.18406 | 0.02931 | 34.12 |
| | STAT6 | 6.18034 | 0.10132 | 9.87 |
| | RB1 | 7.40059 | 0.0341 | 29.322 |
| | PIAS1 | 30.7789 | 0.31407 | 3.184 |
| | FOSL1 | 1.93343 | 0.06905 | 14.482 |
| | BATF3 | 1.39553 | 0.06645 | 15.048 |
| | BATF2 | 109.254 | 6.42674 | 0.1556 |
| | ATF4 | 5.16412 | 0.07074 | 14.136 |
| | MYB | 2.9401 | 0.06125 | 16.326 |
| | ATF3 | 11.6832 | 0.19802 | 5.05 |
| SC | | | | |
| ASTN2-CNOT2 | CNOT6L | 1.171 | 0.065 | 15.37 |
| | AURKA | 4.163 | 0.051 | 19.46 |
| | CNOT8 | 2.06 | 0.09 | 11.17 |
| | TNRC6C | 1.942 | 0.067 | 14.93 |
| | TNRC6B | 3.5 | 0.05 | 20 |
| | CNOT3 | 86.83 | 2.994 | 0.334 |
| | CNOT2 | 3.106 | 0.068 | 14.81 |
| | CNOT1 | 4.661 | 0.059 | 16.95 |
| | CNOT7 | 2.513 | 0.057 | 17.51 |
| | AGO2 | 4.73 | 0.054 | 18.61 |
| | HDAC3 | 22.59 | 0.982 | 1.018 |
| | GPS2 | 0.696 | 0.054 | 18.67 |
| | CNOT2 | 3.106 | 0.068 | 14.81 |
| | NCOR2 | 3.289 | 0.032 | 31.62 |
| | NCOR1 | 4.696 | 0.031 | 32.16 |
| BCOR-ZC3H7B | HDAC3 | 22.59 | 0.982 | 1.018 |
| | HDAC4 | 4.331 | 0.03 | 33.48 |
| | SP1 | 3.018 | 0.069 | 14.58 |
| | CTBP1 | 98.54 | 0.666 | 1.502 |
| | NACC1 | 3.947 | 0.132 | 7.6 |
| | NCOR2 | 3.289 | 0.032 | 31.62 |
| | HDAC1 | 119.7 | 0.251 | 3.992 |
| | CTBP1 | 98.54 | 0.666 | 1.502 |
| | NCOR2 | 3.289 | 0.032 | 31.62 |
| BCOR-CCNB3 | HDAC3 | 22.59 | 0.982 | 1.018 |
| | HDAC4 | 4.331 | 0.03 | 33.48 |
| | SP1 | 3.018 | 0.069 | 14.58 |
| | CTBP1 | 98.54 | 0.666 | 1.502 |
| | NACC1 | 3.947 | 0.132 | 7.6 |
| | NCOR2 | 3.289 | 0.032 | 31.62 |
| | HDAC1 | 119.7 | 0.251 | 3.992 |
| | CTBP1 | 98.54 | 0.666 | 1.502 |
| | NCOR2 | 3.289 | 0.032 | 31.62 |
| CDX1-IRF2BP2 | ELAVL1 | 5.706 | 0.027 | 37.33 |
| | NTRK1 | 8.726 | 0.053 | 19.02 |
| | IRF2BPL | 1.378 | 0.024 | 42.1 |
| | IRF2BP2 | 2.624 | 0.045 | 22.1 |
| | RBM39 | 3.092 | 0.021 | 46.57 |
| CREB1-EWSR1 | BRCA1 | 94.44 | 5.556 | 0.18 |
| | EPAS1 | 252.5 | 2.475 | 0.404 |
| | MYOD1 | 3.693 | 0.054 | 18.41 |
| | ESR1 | 28.12 | 1.222 | 0.818 |
| | JUN | 6.184 | 0.029 | 34.12 |

| | | | | |
|---|---|---|---|---|
| | POLR2A | 101.3 | 0.404 | 2.478 |
| | EP300 | 113.5 | 0.251 | 3.992 |
| | SMARCA4 | 28.47 | 0.142 | 7.024 |
| | EWSR1 | 10.05 | 0.016 | 63.99 |
| | CREBBP | 139.5 | 0.471 | 2.122 |
| | NR3C1 | 6.225 | 0.04 | 25.22 |
| | EP300 | 113.5 | 0.251 | 3.992 |
| | SMARCA4 | 28.47 | 0.142 | 7.024 |
| | CREBBP | 139.5 | 0.471 | 2.122 |
| | NONO | 4.27 | 0.03 | 33.72 |
| | SMARCA4 | 28.47 | 0.142 | 7.024 |
| | CUL3 | 3.375 | 0.048 | 20.74 |
| CTDSP2-FAM19A2 | SETD1A | 1.835 | 0.068 | 14.71 |
| | POLR2A | 101.3 | 0.404 | 2.478 |
| | INTS6 | 27.59 | 0.476 | 2.102 |
| | CTDSP1 | 2.625 | 0.187 | 5.334 |
| | CTDSP2 | 4.199 | 0.3 | 3.334 |
| EPC1-PHF1 | HDAC1 | 119.7 | 0.251 | 3.992 |
| | DHX9 | 3.391 | 0.045 | 22.12 |
| | TP53 | 240.5 | 0.25 | 3.996 |
| | ELAVL1 | 5.706 | 0.027 | 37.33 |
| | E2F6 | 2.239 | 0.061 | 16.53 |
| | RBBP7 | 4.723 | 0.031 | 31.97 |
| | RBBP4 | 4.303 | 0.025 | 39.51 |
| | EZH1 | 135.1 | 15.02 | 0.067 |
| | EZH2 | 111.5 | 0.406 | 2.466 |
| | EED | 197.6 | 2.994 | 0.334 |
| | XRCC6 | 232.4 | 0.88 | 1.136 |
| | XRCC5 | 152.1 | 0.576 | 1.736 |
| | PHF1 | 1.917 | 0.083 | 12 |
| | HDAC1 | 119.7 | 0.251 | 3.992 |
| | TP53 | 240.5 | 0.25 | 3.996 |
| | YEATS4 | 6.25 | 0.063 | 15.84 |
| | TRIM27 | 1.32 | 0.12 | 8.334 |
| | KAT5 | 32.43 | 0.154 | 6.476 |
| | TRIM23 | 1.32 | 0.12 | 8.334 |
| | HIST1H2BA | 4.442 | 0.049 | 20.26 |
| | DMAP1 | 2.383 | 0.041 | 24.34 |
| | MYC | 151 | 0.25 | 3.994 |
| | XRCC6 | 232.4 | 0.88 | 1.136 |
| | MORF4L1 | 4.827 | 0.046 | 21.96 |
| | ING3 | 27.59 | 0.476 | 2.102 |
| ERG-EWSR1 | TP53 | 240.5 | 0.25 | 3.996 |
| | ESR1 | 28.12 | 1.222 | 0.818 |
| | PARP1 | 88.81 | 0.406 | 2.466 |
| | EP300 | 113.5 | 0.251 | 3.992 |
| | CREBBP | 139.5 | 0.471 | 2.122 |
| | XRCC6 | 232.4 | 0.88 | 1.136 |
| | EPAS1 | 252.5 | 2.475 | 0.404 |
| | EP300 | 113.5 | 0.251 | 3.992 |
| | CREBBP | 139.5 | 0.471 | 2.122 |
| ETV6-NTRK3 | PDGFRB | 3.317 | 0.06 | 16.58 |
| | SHC1 | 7.138 | 0.033 | 29.98 |
| | CRKL | 6.106 | 0.057 | 17.69 |

| | | | | |
|---|---|---|---|---|
| | GAB2 | 1.768 | 0.054 | 18.67 |
| | NTRK1 | 8.726 | 0.053 | 19.02 |
| | PLCG1 | 5.711 | 0.051 | 19.61 |
| | GRB2 | 9.905 | 0.019 | 51.99 |
| | HDAC3 | 22.59 | 0.982 | 1.018 |
| | PIN1 | 102.4 | 0.459 | 2.178 |
| | ETV6 | 27.36 | 0.829 | 1.206 |
| EWSR1-ATF1 | PDGFRB | 3.317 | 0.06 | 16.58 |
| | SHC1 | 7.138 | 0.033 | 29.98 |
| | CRKL | 6.106 | 0.057 | 17.69 |
| | GAB2 | 1.768 | 0.054 | 18.67 |
| | NTRK1 | 8.726 | 0.053 | 19.02 |
| | PLCG1 | 5.711 | 0.051 | 19.61 |
| | GRB2 | 9.905 | 0.019 | 51.99 |
| | HDAC3 | 22.59 | 0.982 | 1.018 |
| | PIN1 | 102.4 | 0.459 | 2.178 |
| | ETV6 | 27.36 | 0.829 | 1.206 |
| EWSR1-FLI1 | BRCA1 | 94.44 | 5.556 | 0.18 |
| | POLR2A | 101.3 | 0.404 | 2.478 |
| | ESR1 | 28.12 | 1.222 | 0.818 |
| | EP300 | 113.5 | 0.251 | 3.992 |
| | KAT2B | 6.538 | 0.037 | 26.77 |
| | CREBBP | 139.5 | 0.471 | 2.122 |
| | EPAS1 | 252.5 | 2.475 | 0.404 |
| | EP300 | 113.5 | 0.251 | 3.992 |
| | CREBBP | 139.5 | 0.471 | 2.122 |
| EWSR1-NR4A3 | TSG101 | 17.85 | 0.123 | 8.124 |
| | DHX9 | 3.391 | 0.045 | 22.12 |
| | HDAC3 | 22.59 | 0.982 | 1.018 |
| | TRIM28 | 6.961 | 0.031 | 32.04 |
| | FUS | 19.94 | 0.063 | 16 |
| | RAD23A | 4.032 | 0.027 | 36.46 |
| | JUN | 6.184 | 0.029 | 34.12 |
| | PRMT1 | 5.148 | 0.036 | 28.17 |
| | ILK | 200.6 | 0.951 | 1.052 |
| | BMI1 | 2.558 | 0.041 | 24.62 |
| | ELK1 | 2.68 | 0.099 | 10.07 |
| | POLR2A | 101.3 | 0.404 | 2.478 |
| | CHERP | 2.649 | 0.046 | 21.52 |
| | ATXN3 | 5.241 | 0.065 | 15.46 |
| | EP300 | 113.5 | 0.251 | 3.992 |
| | IRF3 | 84.6 | 1.263 | 0.792 |
| | EPAS1 | 252.5 | 2.475 | 0.404 |
| | TP53 | 240.5 | 0.25 | 3.996 |
| | ESR1 | 28.12 | 1.222 | 0.818 |
| | NONO | 4.27 | 0.03 | 33.72 |
| | RPA1 | 116 | 0.251 | 3.992 |
| | NTRK1 | 8.726 | 0.053 | 19.02 |
| | RPA2 | 11.88 | 0.053 | 19.02 |
| | CUL4A | 6.399 | 0.023 | 43.6 |
| | CUL4B | 8.304 | 0.03 | 33.6 |
| | FASN | 3.59 | 0.042 | 23.95 |
| | CREBBP | 139.5 | 0.471 | 2.122 |
| | HBP1 | 17.21 | 0.82 | 1.22 |

| | | | | |
|---|---|---|---|---|
| | CUL5 | 77.54 | 0.278 | 3.598 |
| | HLTF | 18.24 | 0.397 | 2.522 |
| | EWSR1 | 10.05 | 0.016 | 63.99 |
| | YBX1 | 3.744 | 0.021 | 47.54 |
| | CUL1 | 27.92 | 0.042 | 23.99 |
| | CUL2 | 138.9 | 0.167 | 5.996 |
| | CUL3 | 3.375 | 0.048 | 20.74 |
| | HDAC2 | 80.31 | 0.267 | 3.748 |
| | CREBBP | 139.5 | 0.471 | 2.122 |
| | ESR1 | 28.12 | 1.222 | 0.818 |
| | NONO | 4.27 | 0.03 | 33.72 |
| | FXR2 | 8.166 | 0.07 | 14.21 |
| | CUL3 | 3.375 | 0.048 | 20.74 |
| EWSR1-ETV4 | TSG101 | 17.85 | 0.123 | 8.124 |
| | DHX9 | 3.391 | 0.045 | 22.12 |
| | HDAC3 | 22.59 | 0.982 | 1.018 |
| | RFWD2 | 2.935 | 0.039 | 25.89 |
| | FUS | 19.94 | 0.063 | 16 |
| | RAD23A | 4.032 | 0.027 | 36.46 |
| | JUN | 6.184 | 0.029 | 34.12 |
| | PRMT1 | 5.148 | 0.036 | 28.17 |
| | ILK | 200.6 | 0.951 | 1.052 |
| | SMAD2 | 22.89 | 0.222 | 4.5 |
| | BMI1 | 2.558 | 0.041 | 24.62 |
| | ELK1 | 2.68 | 0.099 | 10.07 |
| | POLR2A | 101.3 | 0.404 | 2.478 |
| | CHERP | 2.649 | 0.046 | 21.52 |
| | ATXN3 | 5.241 | 0.065 | 15.46 |
| | EP300 | 113.5 | 0.251 | 3.992 |
| | IRF3 | 84.6 | 1.263 | 0.792 |
| | EPAS1 | 252.5 | 2.475 | 0.404 |
| | TP53 | 240.5 | 0.25 | 3.996 |
| | ESR1 | 28.12 | 1.222 | 0.818 |
| | NONO | 4.27 | 0.03 | 33.72 |
| | RPA1 | 116 | 0.251 | 3.992 |
| | NTRK1 | 8.726 | 0.053 | 19.02 |
| | RPA2 | 11.88 | 0.053 | 19.02 |
| | CUL4A | 6.399 | 0.023 | 43.6 |
| | CUL4B | 8.304 | 0.03 | 33.6 |
| | FASN | 3.59 | 0.042 | 23.95 |
| | CREBBP | 139.5 | 0.471 | 2.122 |
| | HBP1 | 17.21 | 0.82 | 1.22 |
| | CUL5 | 77.54 | 0.278 | 3.598 |
| | HLTF | 18.24 | 0.397 | 2.522 |
| | EWSR1 | 10.05 | 0.016 | 63.99 |
| | YBX1 | 3.744 | 0.021 | 47.54 |
| | CUL1 | 27.92 | 0.042 | 23.99 |
| | CUL2 | 138.9 | 0.167 | 5.996 |
| | CUL3 | 3.375 | 0.048 | 20.74 |
| | HDAC2 | 80.31 | 0.267 | 3.748 |
| | CREBBP | 139.5 | 0.471 | 2.122 |
| | ESR1 | 28.12 | 1.222 | 0.818 |
| | NONO | 4.27 | 0.03 | 33.72 |
| | FXR2 | 8.166 | 0.07 | 14.21 |

| | | | | |
|---|---|---|---|---|
| | CUL3 | 3.375 | 0.048 | 20.74 |
| EWSR1-PATZ1 | TSG101 | 17.85 | 0.123 | 8.124 |
| | DHX9 | 3.391 | 0.045 | 22.12 |
| | HDAC3 | 22.59 | 0.982 | 1.018 |
| | RFWD2 | 2.935 | 0.039 | 25.89 |
| | FUS | 19.94 | 0.063 | 16 |
| | RAD23A | 4.032 | 0.027 | 36.46 |
| | JUN | 6.184 | 0.029 | 34.12 |
| | PRMT1 | 5.148 | 0.036 | 28.17 |
| | ILK | 200.6 | 0.951 | 1.052 |
| | SMAD2 | 22.89 | 0.222 | 4.5 |
| | BMI1 | 2.558 | 0.041 | 24.62 |
| | ELK1 | 2.68 | 0.099 | 10.07 |
| | POLR2A | 101.3 | 0.404 | 2.478 |
| | CHERP | 2.649 | 0.046 | 21.52 |
| | ATXN3 | 5.241 | 0.065 | 15.46 |
| | EP300 | 113.5 | 0.251 | 3.992 |
| | IRF3 | 84.6 | 1.263 | 0.792 |
| | EPAS1 | 252.5 | 2.475 | 0.404 |
| | TP53 | 240.5 | 0.25 | 3.996 |
| | ESR1 | 28.12 | 1.222 | 0.818 |
| | NONO | 4.27 | 0.03 | 33.72 |
| | RPA1 | 116 | 0.251 | 3.992 |
| | NTRK1 | 8.726 | 0.053 | 19.02 |
| | RPA2 | 11.88 | 0.053 | 19.02 |
| | CUL4A | 6.399 | 0.023 | 43.6 |
| | CUL4B | 8.304 | 0.03 | 33.6 |
| | FASN | 3.59 | 0.042 | 23.95 |
| | CREBBP | 139.5 | 0.471 | 2.122 |
| | HBP1 | 17.21 | 0.82 | 1.22 |
| | CUL5 | 77.54 | 0.278 | 3.598 |
| | HLTF | 18.24 | 0.397 | 2.522 |
| | EWSR1 | 10.05 | 0.016 | 63.99 |
| | YBX1 | 3.744 | 0.021 | 47.54 |
| | CUL1 | 27.92 | 0.042 | 23.99 |
| | CUL2 | 138.9 | 0.167 | 5.996 |
| | CUL3 | 3.375 | 0.048 | 20.74 |
| | HDAC2 | 80.31 | 0.267 | 3.748 |
| | CREBBP | 139.5 | 0.471 | 2.122 |
| | ESR1 | 28.12 | 1.222 | 0.818 |
| | NONO | 4.27 | 0.03 | 33.72 |
| | FXR2 | 8.166 | 0.07 | 14.21 |
| | CUL3 | 3.375 | 0.048 | 20.74 |
| EWSR1-DDIT3 | HDAC1 | 119.7 | 0.251 | 3.992 |
| | EPAS1 | 252.5 | 2.475 | 0.404 |
| | HDAC3 | 22.59 | 0.982 | 1.018 |
| | DDIT3 | 4.748 | 0.068 | 14.74 |
| | CEBPB | 2.123 | 0.032 | 31.09 |
| | ESR1 | 28.12 | 1.222 | 0.818 |
| | FOS | 7.412 | 0.057 | 17.54 |
| | JUN | 6.184 | 0.029 | 34.12 |
| | HBP1 | 17.21 | 0.82 | 1.22 |
| | POLR2A | 101.3 | 0.404 | 2.478 |
| | IRF3 | 84.6 | 1.263 | 0.792 |

| | | | | |
|---|---|---|---|---|
| | TP53 | 240.5 | 0.25 | 3.996 |
| | EP300 | 113.5 | 0.251 | 3.992 |
| | EWSR1 | 10.05 | 0.016 | 63.99 |
| | CREBBP | 139.5 | 0.471 | 2.122 |
| | DHX9 | 3.391 | 0.045 | 22.12 |
| | CUL4A | 6.399 | 0.023 | 43.6 |
| | CUL4B | 8.304 | 0.03 | 33.6 |
| | CUL5 | 77.54 | 0.278 | 3.598 |
| | CUL1 | 27.92 | 0.042 | 23.99 |
| | CUL2 | 138.9 | 0.167 | 5.996 |
| | CUL3 | 3.375 | 0.048 | 20.74 |
| EWSR1-POU5F1 | IRF3 | 84.6 | 1.263 | 0.792 |
| | EPAS1 | 252.5 | 2.475 | 0.404 |
| | HDAC3 | 22.59 | 0.982 | 1.018 |
| | TP53 | 240.5 | 0.25 | 3.996 |
| | ESR1 | 28.12 | 1.222 | 0.818 |
| | JUN | 6.184 | 0.029 | 34.12 |
| | HBP1 | 17.21 | 0.82 | 1.22 |
| | ETS2 | 2.266 | 0.078 | 12.8 |
| | CTNNB1 | 7.85 | 0.027 | 36.43 |
| | POLR2A | 101.3 | 0.404 | 2.478 |
| | EP300 | 113.5 | 0.251 | 3.992 |
| | EWSR1 | 10.05 | 0.016 | 63.99 |
| | CREBBP | 139.5 | 0.471 | 2.122 |
| | NONO | 4.27 | 0.03 | 33.72 |
| | CUL2 | 138.9 | 0.167 | 5.996 |
| | CUL3 | 3.375 | 0.048 | 20.74 |
| EWSR1-SP3 | HDAC1 | 119.7 | 0.251 | 3.992 |
| | EPAS1 | 252.5 | 2.475 | 0.404 |
| | HDAC3 | 22.59 | 0.982 | 1.018 |
| | CEBPB | 2.123 | 0.032 | 31.09 |
| | ESR1 | 28.12 | 1.222 | 0.818 |
| | JUN | 6.184 | 0.029 | 34.12 |
| | HBP1 | 17.21 | 0.82 | 1.22 |
| | POLR2A | 101.3 | 0.404 | 2.478 |
| | IRF3 | 84.6 | 1.263 | 0.792 |
| | TP53 | 240.5 | 0.25 | 3.996 |
| | RELA | 9.357 | 0.053 | 19.02 |
| | EP300 | 113.5 | 0.251 | 3.992 |
| | EWSR1 | 10.05 | 0.016 | 63.99 |
| | CREBBP | 139.5 | 0.471 | 2.122 |
| | DHX9 | 3.391 | 0.045 | 22.12 |
| | CUL4A | 6.399 | 0.023 | 43.6 |
| | CUL4B | 8.304 | 0.03 | 33.6 |
| | CUL5 | 77.54 | 0.278 | 3.598 |
| | CUL1 | 27.92 | 0.042 | 23.99 |
| | CUL2 | 138.9 | 0.167 | 5.996 |
| | CUL3 | 3.375 | 0.048 | 20.74 |
| FOXO4-CIC | XPO1 | 4.291 | 0.048 | 20.74 |
| | CTNNB1 | 7.85 | 0.027 | 36.43 |
| | VDR | 119.9 | 1.348 | 0.742 |
| | ESR1 | 28.12 | 1.222 | 0.818 |
| | SMAD4 | 25.62 | 0.249 | 4.02 |
| | SMAD3 | 1.962 | 0.019 | 52.5 |

| | | | | |
|---|---|---|---|---|
| | SFN | 8.389 | 0.034 | 29.8 |
| | AKT1 | 2.106 | 0.035 | 28.49 |
| | FOXO4 | 20.83 | 1.042 | 0.96 |
| | MDM2 | 9.935 | 0.053 | 19.02 |
| | NLK | 5.058 | 0.09 | 11.07 |
| | CREBBP | 139.5 | 0.471 | 2.122 |
| FUS-ERG | RPA1 | 116 | 0.251 | 3.992 |
| | SF3B2 | 1.773 | 0.068 | 14.67 |
| | PRKDC | 5.478 | 0.023 | 42.72 |
| | PRPF8 | 2.684 | 0.018 | 56.99 |
| | SF3A2 | 2.053 | 0.079 | 12.67 |
| | DHX15 | 1.947 | 0.024 | 42.12 |
| | RPA2 | 11.88 | 0.053 | 19.02 |
| | CUL3 | 3.375 | 0.048 | 20.74 |
| | ABL1 | 5.805 | 0.034 | 29.8 |
| | PARP1 | 88.81 | 0.406 | 2.466 |
| | PRKDC | 5.478 | 0.023 | 42.72 |
| FUS-CREB3L1 | NONO | 4.27 | 0.03 | 33.72 |
| | CUL4A | 6.399 | 0.023 | 43.6 |
| | CUL4B | 8.304 | 0.03 | 33.6 |
| | DHX15 | 1.947 | 0.024 | 42.12 |
| | CUL5 | 77.54 | 0.278 | 3.598 |
| | CUL1 | 27.92 | 0.042 | 23.99 |
| | CUL2 | 138.9 | 0.167 | 5.996 |
| | CUL3 | 3.375 | 0.048 | 20.74 |
| | VCP | 146.7 | 0.25 | 3.994 |
| | FBXW11 | 3.809 | 0.05 | 19.95 |
| | CUL1 | 27.92 | 0.042 | 23.99 |
| FUS-DDIT3 | HDAC1 | 119.7 | 0.251 | 3.992 |
| | DDX17 | 2.559 | 0.018 | 55.49 |
| | EPAS1 | 252.5 | 2.475 | 0.404 |
| | RELA | 9.357 | 0.053 | 19.02 |
| | ESR1 | 28.12 | 1.222 | 0.818 |
| | JUN | 6.184 | 0.029 | 34.12 |
| | TP73 | 30.45 | 0.272 | 3.678 |
| | EWSR1 | 10.05 | 0.016 | 63.99 |
| | CTNNB1 | 7.85 | 0.027 | 36.43 |
| | DDX5 | 3.278 | 0.016 | 61.93 |
| | CDK2 | 28.82 | 0.124 | 8.086 |
| | DDIT3 | 4.748 | 0.068 | 14.74 |
| | MDM2 | 9.935 | 0.053 | 19.02 |
| | EP300 | 113.5 | 0.251 | 3.992 |
| | TRIP4 | 1.764 | 0.043 | 23.24 |
| | CREBBP | 139.5 | 0.471 | 2.122 |
| | VCP | 146.7 | 0.25 | 3.994 |
| | FBXW11 | 3.809 | 0.05 | 19.95 |
| | CUL1 | 27.92 | 0.042 | 23.99 |
| FUS-ATF1 | HDAC1 | 119.7 | 0.251 | 3.992 |
| | DDX17 | 2.559 | 0.018 | 55.49 |
| | EPAS1 | 252.5 | 2.475 | 0.404 |
| | RELA | 9.357 | 0.053 | 19.02 |
| | ESR1 | 28.12 | 1.222 | 0.818 |
| | JUN | 6.184 | 0.029 | 34.12 |
| | TP73 | 30.45 | 0.272 | 3.678 |

| | | | | |
|---|---|---|---|---|
| | EWSR1 | 10.05 | 0.016 | 63.99 |
| | CTNNB1 | 7.85 | 0.027 | 36.43 |
| | DDX5 | 3.278 | 0.016 | 61.93 |
| | CDK2 | 28.82 | 0.124 | 8.086 |
| | DDIT3 | 4.748 | 0.068 | 14.74 |
| | MDM2 | 9.935 | 0.053 | 19.02 |
| | EP300 | 113.5 | 0.251 | 3.992 |
| | TRIP4 | 1.764 | 0.043 | 23.24 |
| | CREBBP | 139.5 | 0.471 | 2.122 |
| | VCP | 146.7 | 0.25 | 3.994 |
| | FBXW11 | 3.809 | 0.05 | 19.95 |
| | CUL1 | 27.92 | 0.042 | 23.99 |
| FUS-CREB3L2 | NONO | 4.27 | 0.03 | 33.72 |
| | CUL4A | 6.399 | 0.023 | 43.6 |
| | CUL4B | 8.304 | 0.03 | 33.6 |
| | DHX15 | 1.947 | 0.024 | 42.12 |
| | CUL5 | 77.54 | 0.278 | 3.598 |
| | CUL1 | 27.92 | 0.042 | 23.99 |
| | CUL2 | 138.9 | 0.167 | 5.996 |
| | CUL3 | 3.375 | 0.048 | 20.74 |
| | VCP | 146.7 | 0.25 | 3.994 |
| | FBXW11 | 3.809 | 0.05 | 19.95 |
| | CUL1 | 27.92 | 0.042 | 23.99 |
| HEY1-NCOA2 | BRCA1 | 94.44 | 5.556 | 0.18 |
| | RARA | 5.002 | 0.046 | 21.59 |
| | NR3C1 | 6.225 | 0.04 | 25.22 |
| | VDR | 119.9 | 1.348 | 0.742 |
| | STAT6 | 6.18 | 0.101 | 9.87 |
| | HNF4A | 128 | 2 | 0.5 |
| | PRMT1 | 5.148 | 0.036 | 28.17 |
| | CARM1 | 3.802 | 0.043 | 23.15 |
| | RXRA | 4.577 | 0.042 | 23.82 |
| | PPARG | 5.048 | 0.039 | 25.75 |
| | PPARD | 3.533 | 0.08 | 12.45 |
| | AR | 2.106 | 0.035 | 28.49 |
| | PPARA | 3.641 | 0.087 | 11.53 |
| | ESR2 | 41.67 | 0.417 | 2.4 |
| | EP300 | 113.5 | 0.251 | 3.992 |
| | NCOA2 | 3.388 | 0.053 | 18.89 |
| | NCOA3 | 3.966 | 0.037 | 26.73 |
| | NCOA1 | 3.675 | 0.036 | 27.49 |
| | TP53 | 240.5 | 0.25 | 3.996 |
| | ESR1 | 28.12 | 1.222 | 0.818 |
| | AHR | 150.8 | 3.968 | 0.252 |
| | ARNT | 222.5 | 5.297 | 0.189 |
| | THRB | 3.31 | 0.069 | 14.5 |
| | THRA | 4.344 | 0.063 | 15.88 |
| | NR1I3 | 2.242 | 0.077 | 12.93 |
| | NR1I2 | 3.385 | 0.106 | 9.454 |
| | PGR | 3.769 | 0.061 | 16.45 |
| | CREBBP | 139.5 | 0.471 | 2.122 |
| | PIAS3 | 4.01 | 0.08 | 12.47 |
| | NCOA2 | 3.388 | 0.053 | 18.89 |
| | NCOA1 | 3.675 | 0.036 | 27.49 |

| | | | | |
|---|---|---|---|---|
| | UBR5 | 4.334 | 0.02 | 50.07 |
| IRX2-TERT | YWHAZ | 9.094 | 0.027 | 36.84 |
| | AKT1 | 2.106 | 0.035 | 28.49 |
| | RPS6KB1 | 3.456 | 0.061 | 16.49 |
| | ENO1 | 6.627 | 0.245 | 4.074 |
| | MTOR | 6.768 | 0.048 | 20.69 |
| | MDM2 | 9.935 | 0.053 | 19.02 |
| | TERT | 4.892 | 0.074 | 13.49 |
| | XRCC6 | 232.4 | 0.88 | 1.136 |
| | TERF1 | 15.75 | 0.051 | 19.49 |
| | STUB1 | 7.969 | 0.033 | 29.99 |
| | TERT | 4.892 | 0.074 | 13.49 |
| | POT1 | 15.33 | 0.073 | 13.76 |
| | TERT | 4.892 | 0.074 | 13.49 |
| | MTOR | 6.768 | 0.048 | 20.69 |
| | YWHAQ | 9.32 | 0.023 | 43.99 |
| | RUVBL2 | 5.408 | 0.024 | 41.24 |
| | TPP1 | 4.73 | 0.042 | 23.68 |
| | TERT | 4.892 | 0.074 | 13.49 |
| | POT1 | 15.33 | 0.073 | 13.76 |
| JAZF1-SUZ12 | DHX9 | 3.391 | 0.045 | 22.12 |
| | RBM5 | 2.34 | 0.075 | 13.25 |
| | FBXW11 | 3.809 | 0.05 | 19.95 |
| | DDX3X | 97.86 | 0.67 | 1.492 |
| | NXF1 | 1.244 | 0.02 | 49.03 |
| | SF3B4 | 3.44 | 0.034 | 29.07 |
| | PRMT1 | 5.148 | 0.036 | 28.17 |
| | SF3B1 | 3.067 | 0.017 | 57.71 |
| | SF3B2 | 1.773 | 0.068 | 14.67 |
| | PRPF8 | 2.684 | 0.018 | 56.99 |
| | EED | 197.6 | 2.994 | 0.334 |
| | CRNKL1 | 2.125 | 0.051 | 19.77 |
| | RNPS1 | 3.812 | 0.023 | 44.07 |
| | UBE2I | 4.334 | 0.02 | 50.07 |
| | SNRNP200 | 2.819 | 0.019 | 53.92 |
| | SRSF7 | 2.019 | 0.019 | 53.5 |
| | SNRPD3 | 2.323 | 0.022 | 44.76 |
| | RALY | 2.018 | 0.027 | 37.66 |
| | DDX5 | 3.278 | 0.016 | 61.93 |
| | PRPF19 | 2.593 | 0.019 | 53.61 |
| | RNF2 | 177.8 | 0.25 | 3.994 |
| | SNRPA1 | 3.076 | 0.023 | 43.56 |
| | SON | 3.018 | 0.069 | 14.58 |
| | EFTUD2 | 2.743 | 0.02 | 50.68 |
| | U2AF1 | 4.334 | 0.02 | 50.07 |
| | SF3A1 | 2.895 | 0.019 | 52.86 |
| | EPRS | 3.12 | 0.03 | 33.33 |
| | ILF2 | 2.535 | 0.039 | 25.64 |
| | EIF4A3 | 6.718 | 0.008 | 124 |
| | CDC40 | 1.125 | 0.043 | 23.11 |
| | ILF3 | 1.727 | 0.027 | 37.64 |
| | RANBP2 | 3.654 | 0.029 | 33.94 |
| | DNMT3B | 293.1 | 4.31 | 0.232 |
| | HDAC1 | 119.7 | 0.251 | 3.992 |

| | | | | |
|---|---|---|---|---|
| | HDAC2 | 80.31 | 0.267 | 3.748 |
| | TRIM28 | 6.961 | 0.031 | 32.04 |
| | CHD4 | 3.583 | 0.03 | 33.21 |
| | UHRF1 | 5.639 | 0.042 | 23.76 |
| | DNMT1 | 3.02 | 0.036 | 27.81 |
| | MTA1 | 3.367 | 0.036 | 27.62 |
| | NR2C2 | 0.959 | 0.074 | 13.56 |
| | EZH2 | 111.5 | 0.406 | 2.466 |
| | GATAD2B | 1.071 | 0.054 | 18.67 |
| | EED | 197.6 | 2.994 | 0.334 |
| | CBX5 | 60.85 | 0.378 | 2.646 |
| | RBBP4 | 4.303 | 0.025 | 39.51 |
| | CBX3 | 4.562 | 0.045 | 22.14 |
| | SETDB1 | 9.781 | 0.078 | 12.88 |
| | VCP | 146.7 | 0.25 | 3.994 |
| | BRCA1 | 94.44 | 5.556 | 0.18 |
| | MTOR | 6.768 | 0.048 | 20.69 |
| | NXF1 | 1.244 | 0.02 | 49.03 |
| | RUVBL2 | 5.408 | 0.024 | 41.24 |
| | VCP | 146.7 | 0.25 | 3.994 |
| | FBXW11 | 3.809 | 0.05 | 19.95 |
| | SKP1 | 7.259 | 0.039 | 25.48 |
| | BTRC | 1.723 | 0.041 | 24.38 |
| | EZH2 | 111.5 | 0.406 | 2.466 |
| | DHX9 | 3.391 | 0.045 | 22.12 |
| | ADAR | 2.668 | 0.044 | 22.49 |
| | EZH2 | 111.5 | 0.406 | 2.466 |
| | FBXW11 | 3.809 | 0.05 | 19.95 |
| | HDAC2 | 80.31 | 0.267 | 3.748 |
| | NXF1 | 1.244 | 0.02 | 49.03 |
| | JARID2 | 1.378 | 0.024 | 42.1 |
| | SETDB1 | 9.781 | 0.078 | 12.88 |
| | RELA | 9.357 | 0.053 | 19.02 |
| | FBXW11 | 3.809 | 0.05 | 19.95 |
| | BTRC | 1.723 | 0.041 | 24.38 |
| | BTRC | 1.723 | 0.041 | 24.38 |
| | CSNK2B | 3.051 | 0.073 | 13.77 |
| | NXF1 | 1.244 | 0.02 | 49.03 |
| JAZF1-PHF1 | DHX9 | 3.391 | 0.045 | 22.12 |
| | EZH1 | 135.1 | 15.02 | 0.067 |
| | PPARG | 5.048 | 0.039 | 25.75 |
| | EZH2 | 111.5 | 0.406 | 2.466 |
| | EED | 197.6 | 2.994 | 0.334 |
| | XRCC6 | 232.4 | 0.88 | 1.136 |
| | XRCC5 | 152.1 | 0.576 | 1.736 |
| | PHF1 | 1.917 | 0.083 | 12 |
| | HDAC1 | 119.7 | 0.251 | 3.992 |
| | PPARG | 5.048 | 0.039 | 25.75 |
| | EZH2 | 111.5 | 0.406 | 2.466 |
| | PHF1 | 1.917 | 0.083 | 12 |
| | PHF1 | 1.917 | 0.083 | 12 |
| | TP53 | 240.5 | 0.25 | 3.996 |
| | XRCC6 | 232.4 | 0.88 | 1.136 |
| MEAF6-TRERF1 | HDAC1 | 119.7 | 0.251 | 3.992 |

| | | | | |
|---|---|---|---|---|
| | TRERF1 | 1.32 | 0.12 | 8.334 |
| | KAT5 | 32.43 | 0.154 | 6.476 |
| | ING3 | 27.59 | 0.476 | 2.102 |
| | CREBBP | 139.5 | 0.471 | 2.122 |
| | YEATS4 | 6.25 | 0.063 | 15.84 |
| | EP300 | 113.5 | 0.251 | 3.992 |
| | MORF4L1 | 4.827 | 0.046 | 21.96 |
| | HIST1H2BA | 4.442 | 0.049 | 20.26 |
| | ELAVL1 | 5.706 | 0.027 | 37.33 |
| | TRERF1 | 1.32 | 0.12 | 8.334 |
| | KAT6A | 308.3 | 8.333 | 0.12 |
| | CREBBP | 139.5 | 0.471 | 2.122 |
| | NR5A1 | 2.7 | 0.104 | 9.63 |
| | SOX2 | 19.59 | 0.057 | 17.66 |
| | HDAC1 | 119.7 | 0.251 | 3.992 |
| | TRERF1 | 1.32 | 0.12 | 8.334 |
| MEAF6-PHF1 | DHX9 | 3.391 | 0.045 | 22.12 |
| | EZH1 | 135.1 | 15.02 | 0.067 |
| | EZH2 | 111.5 | 0.406 | 2.466 |
| | EED | 197.6 | 2.994 | 0.334 |
| | XRCC6 | 232.4 | 0.88 | 1.136 |
| | XRCC5 | 152.1 | 0.576 | 1.736 |
| | PHF1 | 1.917 | 0.083 | 12 |
| | PHF1 | 1.917 | 0.083 | 12 |
| | TP53 | 240.5 | 0.25 | 3.996 |
| | XRCC6 | 232.4 | 0.88 | 1.136 |
| | KAT6A | 308.3 | 8.333 | 0.12 |
| | TP53 | 240.5 | 0.25 | 3.996 |
| | ELAVL1 | 5.706 | 0.027 | 37.33 |
| | HDAC1 | 119.7 | 0.251 | 3.992 |
| | EZH2 | 111.5 | 0.406 | 2.466 |
| | PHF1 | 1.917 | 0.083 | 12 |
| NR4A3-TAF15 | TRIM28 | 6.961 | 0.031 | 32.04 |
| | FUS | 19.94 | 0.063 | 16 |
| | PRMT1 | 5.148 | 0.036 | 28.17 |
| | COPS6 | 199 | 0.25 | 3.994 |
| | COPS5 | 199 | 0.25 | 3.994 |
| | POLR2C | 3.4 | 0.034 | 29.7 |
| | POLR2A | 101.3 | 0.404 | 2.478 |
| | TAF15 | 2.424 | 0.042 | 23.93 |
| | SF1 | 5.091 | 0.041 | 24.16 |
| | NEDD8 | 5.227 | 0.02 | 49.17 |
| | POLR2E | 2.95 | 0.029 | 34.91 |
| | RPA1 | 116 | 0.251 | 3.992 |
| | RPA2 | 11.88 | 0.053 | 19.02 |
| | CUL4A | 6.399 | 0.023 | 43.6 |
| | CUL4B | 8.304 | 0.03 | 33.6 |
| | CUL5 | 77.54 | 0.278 | 3.598 |
| | CUL1 | 27.92 | 0.042 | 23.99 |
| | CUL2 | 138.9 | 0.167 | 5.996 |
| | CUL3 | 3.375 | 0.048 | 20.74 |
| | EZH2 | 111.5 | 0.406 | 2.466 |
| | TRIM28 | 6.961 | 0.031 | 32.04 |
| | CUL1 | 27.92 | 0.042 | 23.99 |

| | | | | |
|---|---|---|---|---|
| NR4A3-TFG | CUL4A | 6.399 | 0.023 | 43.6 |
| | CUL4B | 8.304 | 0.03 | 33.6 |
| | CUL5 | 77.54 | 0.278 | 3.598 |
| | CUL1 | 27.92 | 0.042 | 23.99 |
| | CUL2 | 138.9 | 0.167 | 5.996 |
| | CUL3 | 3.375 | 0.048 | 20.74 |
| | TRIM28 | 6.961 | 0.031 | 32.04 |
| | CUL1 | 27.92 | 0.042 | 23.99 |
| | CUL3 | 3.375 | 0.048 | 20.74 |
| NUP107-LGR5 | NUP153 | 3.7 | 0.035 | 28.38 |
| | KPNB1 | 4.058 | 0.072 | 13.8 |
| | NTRK1 | 8.726 | 0.053 | 19.02 |
| | CUL3 | 3.375 | 0.048 | 20.74 |
| | NUP153 | 3.7 | 0.035 | 28.38 |
| | EIF4B | 3.678 | 0.048 | 20.94 |
| | CUL3 | 3.375 | 0.048 | 20.74 |
| | EED | 197.6 | 2.994 | 0.334 |
| | KPNB1 | 4.058 | 0.072 | 13.8 |
| | TP53BP1 | 240.5 | 0.25 | 3.996 |
| PAPPA-NUP107 | NUP153 | 3.7 | 0.035 | 28.38 |
| | ELAVL1 | 5.706 | 0.027 | 37.33 |
| | KPNB1 | 4.058 | 0.072 | 13.8 |
| | NTRK1 | 8.726 | 0.053 | 19.02 |
| | SMAD3 | 1.962 | 0.019 | 52.5 |
| | VCP | 146.7 | 0.25 | 3.994 |
| | CUL3 | 3.375 | 0.048 | 20.74 |
| | EIF4B | 3.678 | 0.048 | 20.94 |
| | SMAD9 | 13.88 | 0.125 | 8 |
| | SKIL | 7.834 | 0.082 | 12.13 |
| | PAPPA | 1.31 | 0.131 | 7.636 |
| | SMAD2 | 22.89 | 0.222 | 4.5 |
| | SMAD3 | 1.962 | 0.019 | 52.5 |
| | TP53BP1 | 240.5 | 0.25 | 3.996 |
| | ELAVL1 | 5.706 | 0.027 | 37.33 |
| | EED | 197.6 | 2.994 | 0.334 |
| | KPNB1 | 4.058 | 0.072 | 13.8 |
| | NUP214 | 2.745 | 0.047 | 21.49 |
| | SMAD2 | 22.89 | 0.222 | 4.5 |
| | SMAD3 | 1.962 | 0.019 | 52.5 |
| PAX3-FOXO1 | NCOA1 | 3.675 | 0.036 | 27.49 |
| | ESR1 | 28.12 | 1.222 | 0.818 |
| | PARP1 | 88.81 | 0.406 | 2.466 |
| | AR | 2.106 | 0.035 | 28.49 |
| | EP300 | 113.5 | 0.251 | 3.992 |
| | CREBBP | 139.5 | 0.471 | 2.122 |
| | TRIM28 | 6.961 | 0.031 | 32.04 |
| | PARP1 | 88.81 | 0.406 | 2.466 |
| | CREBBP | 139.5 | 0.471 | 2.122 |
| PAX7-FOXO1 | RARA | 5.002 | 0.046 | 21.59 |
| | NCOA1 | 3.675 | 0.036 | 27.49 |
| | MYOD1 | 3.693 | 0.054 | 18.41 |
| | ESR1 | 28.12 | 1.222 | 0.818 |
| | HNF4A | 128 | 2 | 0.5 |
| | PARP1 | 88.81 | 0.406 | 2.466 |

| | | | | |
|---|---|---|---|---|
| | SMAD3 | 1.962 | 0.019 | 52.5 |
| | FOXO1 | 28.04 | 0.519 | 1.926 |
| | AR | 2.106 | 0.035 | 28.49 |
| | MDM2 | 9.935 | 0.053 | 19.02 |
| | EP300 | 113.5 | 0.251 | 3.992 |
| | CREBBP | 139.5 | 0.471 | 2.122 |
| | AKT1 | 2.106 | 0.035 | 28.49 |
| | EP300 | 113.5 | 0.251 | 3.992 |
| | CREBBP | 139.5 | 0.471 | 2.122 |
| SS18-SSX2 | DPF2 | 2.095 | 0.047 | 21.48 |
| | SMARCC2 | 103.5 | 0.803 | 1.246 |
| | SMARCC1 | 103.8 | 0.894 | 1.118 |
| | PHF10 | 1.917 | 0.083 | 12 |
| | ELAVL1 | 5.706 | 0.027 | 37.33 |
| | DPF3 | 0.95 | 0.038 | 26.31 |
| | ARID2 | 30 | 1.25 | 0.8 |
| | DPF1 | 0.703 | 0.078 | 12.8 |
| | SMARCD3 | 1.971 | 0.049 | 20.29 |
| | SMARCE1 | 4.706 | 0.04 | 25.08 |
| | SMARCD1 | 4.956 | 0.046 | 21.59 |
| | EED | 197.6 | 2.994 | 0.334 |
| | SMARCA2 | 3.169 | 0.031 | 32.5 |
| | EP300 | 113.5 | 0.251 | 3.992 |
| | SMARCA4 | 28.47 | 0.142 | 7.024 |
| | HDAC1 | 119.7 | 0.251 | 3.992 |
| | ARID1B | 1.658 | 0.045 | 22.32 |
| | ARID1A | 1.812 | 0.033 | 30.36 |
| | ACTL6A | 3.533 | 0.034 | 29.15 |
| | HDAC2 | 80.31 | 0.267 | 3.748 |
| | CUL3 | 3.375 | 0.048 | 20.74 |
| | RNF2 | 177.8 | 0.25 | 3.994 |
| | SMARCD2 | 1.937 | 0.042 | 23.74 |
| | GRB2 | 9.905 | 0.019 | 51.99 |
| | YWHAG | 8.003 | 0.029 | 33.99 |
| | CUL3 | 3.375 | 0.048 | 20.74 |
| SS18-SSX1 | SMARCC2 | 103.5 | 0.803 | 1.246 |
| | DPF2 | 2.095 | 0.047 | 21.48 |
| | SMARCC1 | 103.8 | 0.894 | 1.118 |
| | PHF10 | 5.155 | 0.224 | 4.462 |
| | ELAVL1 | 5.706 | 0.027 | 37.33 |
| | DPF3 | 0.95 | 0.038 | 26.31 |
| | DPF1 | 0.703 | 0.078 | 12.8 |
| | ARID2 | 30 | 1.25 | 0.8 |
| | HDAC2 | 80.31 | 0.267 | 3.748 |
| | SMARCD3 | 1.971 | 0.049 | 20.29 |
| | SMARCE1 | 4.706 | 0.04 | 25.08 |
| | SMARCD1 | 4.956 | 0.046 | 21.59 |
| | EED | 197.6 | 2.994 | 0.334 |
| | SMARCA2 | 3.169 | 0.031 | 32.5 |
| | EP300 | 113.5 | 0.251 | 3.992 |
| | SMARCA4 | 28.47 | 0.142 | 7.024 |
| | HDAC1 | 119.7 | 0.251 | 3.992 |
| | ARID1B | 1.658 | 0.045 | 22.32 |
| | ARID1A | 1.812 | 0.033 | 30.36 |

| | | | | |
|---|---|---|---|---|
| | ACTL6A | 3.533 | 0.034 | 29.15 |
| | CUL3 | 3.375 | 0.048 | 20.74 |
| | SMARCD2 | 1.937 | 0.042 | 23.74 |
| | GRB2 | 9.905 | 0.019 | 51.99 |
| | YWHAG | 8.003 | 0.029 | 33.99 |
| | CUL3 | 3.375 | 0.048 | 20.74 |
| SS18L1-SSX1 | SMARCC1 | 103.8 | 0.894 | 1.118 |
| | STAT3 | 9.578 | 0.043 | 23.28 |
| | BMI1 | 2.558 | 0.041 | 24.62 |
| | WHSC1L1 | 119.9 | 1.348 | 0.742 |
| | SMAD3 | 1.962 | 0.019 | 52.5 |
| | HDAC2 | 80.31 | 0.267 | 3.748 |
| | SMAD1 | 8.22 | 0.043 | 23.36 |
| | EP300 | 113.5 | 0.251 | 3.992 |
| | SMARCA4 | 28.47 | 0.142 | 7.024 |
| | CREBBP | 139.5 | 0.471 | 2.122 |
| | DPF2 | 2.095 | 0.047 | 21.48 |
| | SMARCC1 | 103.8 | 0.894 | 1.118 |
| | SMARCE1 | 4.706 | 0.04 | 25.08 |
| | SMARCA4 | 28.47 | 0.142 | 7.024 |
| | CUL3 | 3.375 | 0.048 | 20.74 |
| TGFBR3-MGEA5 | MAST1 | 2.031 | 0.052 | 19.2 |
| | RNF32 | 2.1 | 0.05 | 20 |
| | PAXIP1 | 13.28 | 0.052 | 19.35 |
| | CBX8 | 5.876 | 0.046 | 21.78 |
| | CSNK2B | 3.051 | 0.073 | 13.77 |
| | PAXIP1 | 13.28 | 0.052 | 19.35 |
| TRIO-TERT | YWHAZ | 9.094 | 0.027 | 36.84 |
| | AKT1 | 2.106 | 0.035 | 28.49 |
| | RPS6KB1 | 3.456 | 0.061 | 16.49 |
| | ENO1 | 6.627 | 0.245 | 4.074 |
| | MTOR | 6.768 | 0.048 | 20.69 |
| | MDM2 | 9.935 | 0.053 | 19.02 |
| | TERT | 4.892 | 0.074 | 13.49 |
| | XRCC6 | 232.4 | 0.88 | 1.136 |
| | TERF1 | 15.75 | 0.051 | 19.49 |
| | STUB1 | 7.969 | 0.033 | 29.99 |
| | TERT | 4.892 | 0.074 | 13.49 |
| | POT1 | 15.33 | 0.073 | 13.76 |
| | TERT | 4.892 | 0.074 | 13.49 |
| | MTOR | 6.768 | 0.048 | 20.69 |
| | YWHAQ | 9.32 | 0.023 | 43.99 |
| | RUVBL2 | 5.408 | 0.024 | 41.24 |
| WDR70-RCOR1 | SMARCC2 | 103.5 | 0.803 | 1.246 |
| | HDAC1 | 119.7 | 0.251 | 3.992 |
| | HDAC3 | 22.59 | 0.982 | 1.018 |
| | HDAC2 | 80.31 | 0.267 | 3.748 |
| | KDM1A | 32.43 | 0.154 | 6.476 |
| | RCOR1 | 3.204 | 0.062 | 16.23 |
| | NR2C1 | 1.132 | 0.042 | 23.86 |
| | SMARCE1 | 4.706 | 0.04 | 25.08 |
| | CTBP1 | 98.54 | 0.666 | 1.502 |
| | NR2E1 | 4 | 0.129 | 7.75 |
| | SMARCA4 | 28.47 | 0.142 | 7.024 |

| | | | | |
|---|---|---|---|---|
| | CTBP2 | 9.352 | 0.094 | 10.59 |
| | KDM5B | 9.343 | 0.044 | 22.48 |
| | MTA3 | 2.232 | 0.041 | 24.64 |
| | KDM1A | 32.43 | 0.154 | 6.476 |
| | HDAC3 | 22.59 | 0.982 | 1.018 |
| | CTBP1 | 98.54 | 0.666 | 1.502 |
| YWHAE-NUTM2B | LARP1 | 2.17 | 0.026 | 37.78 |
| | NOS2 | 4.671 | 0.032 | 31.26 |
| | YWHAQ | 9.32 | 0.023 | 43.99 |
| | YWHAG | 8.003 | 0.029 | 33.99 |
| | HUWE1 | 7.079 | 0.016 | 63.99 |
| | MAST2 | 2.442 | 0.122 | 8.19 |
| | NTRK1 | 8.726 | 0.053 | 19.02 |
| | VCP | 146.7 | 0.25 | 3.994 |
| | AKT1 | 2.106 | 0.035 | 28.49 |
| | MAP2K1 | 3.938 | 0.067 | 14.98 |
| | FBXW11 | 3.809 | 0.05 | 19.95 |
| | ARAF | 6.317 | 0.073 | 13.77 |
| | CUL3 | 3.375 | 0.048 | 20.74 |
| | PARK2 | 102 | 0.251 | 3.99 |
| | RAF1 | 6.385 | 0.043 | 23.49 |
| | YWHAZ | 9.094 | 0.027 | 36.84 |
| | UBXN1 | 5.639 | 0.042 | 23.76 |
| | TP53 | 240.5 | 0.25 | 3.996 |
| | RUVBL2 | 5.408 | 0.024 | 41.24 |
| | BTRC | 1.723 | 0.041 | 24.38 |
| | TUBB | 4.334 | 0.02 | 50.07 |
| | CDC37 | 4.101 | 0.093 | 10.73 |
| | YWHAH | 6.807 | 0.038 | 26.44 |
| | MAST3 | 4.683 | 0.109 | 9.182 |
| | BRAF | 2.061 | 0.049 | 20.38 |
| | YWHAB | 5.302 | 0.017 | 58.65 |
| | KSR1 | 1.73 | 0.108 | 9.25 |
| | YWHAE | 9.029 | 0.026 | 38.65 |
| | CUL1 | 27.92 | 0.042 | 23.99 |
| | MAPK7 | 3.783 | 0.061 | 16.39 |
| | YWHAZ | 9.094 | 0.027 | 36.84 |
| | IGF1R | 3.686 | 0.057 | 17.64 |
| | IRS1 | 2.624 | 0.045 | 22.1 |
| | YWHAQ | 9.32 | 0.023 | 43.99 |
| | MST1R | 1.235 | 0.062 | 16.19 |
| | NTRK1 | 8.726 | 0.053 | 19.02 |
| | CBL | 19.28 | 0.083 | 12.09 |
| | TUBB | 4.334 | 0.02 | 50.07 |
| | YWHAB | 5.302 | 0.017 | 58.65 |
| | YWHAH | 6.807 | 0.038 | 26.44 |
| | GRB2 | 9.905 | 0.019 | 51.99 |
| | ABL1 | 5.805 | 0.034 | 29.8 |
| | SORBS2 | 2.606 | 0.057 | 17.65 |
| | BCAR1 | 3.383 | 0.054 | 18.62 |
| | YWHAE | 9.029 | 0.026 | 38.65 |
| | VCP | 146.7 | 0.25 | 3.994 |
| | CDK2 | 28.82 | 0.124 | 8.086 |
| | CUL1 | 27.92 | 0.042 | 23.99 |

| | | | | |
|---|---|---|---|---|
| | CDC37 | 4.101 | 0.093 | 10.73 |
| | LRRK2 | 7.159 | 0.052 | 19.28 |
| | YWHAE | 9.029 | 0.026 | 38.65 |
| YWHAE-NUTM2A-AS1 | LARP1 | 2.17 | 0.026 | 37.78 |
| | NOS2 | 4.671 | 0.032 | 31.26 |
| | YWHAQ | 9.32 | 0.023 | 43.99 |
| | YWHAG | 8.003 | 0.029 | 33.99 |
| | HUWE1 | 7.079 | 0.016 | 63.99 |
| | MAST2 | 2.442 | 0.122 | 8.19 |
| | NTRK1 | 8.726 | 0.053 | 19.02 |
| | VCP | 146.7 | 0.25 | 3.994 |
| | AKT1 | 2.106 | 0.035 | 28.49 |
| | MAP2K1 | 3.938 | 0.067 | 14.98 |
| | FBXW11 | 3.809 | 0.05 | 19.95 |
| | ARAF | 6.317 | 0.073 | 13.77 |
| | CUL3 | 3.375 | 0.048 | 20.74 |
| | PARK2 | 102 | 0.251 | 3.99 |
| | RAF1 | 6.385 | 0.043 | 23.49 |
| | YWHAZ | 9.094 | 0.027 | 36.84 |
| | UBXN1 | 5.639 | 0.042 | 23.76 |
| | TP53 | 240.5 | 0.25 | 3.996 |
| | RUVBL2 | 5.408 | 0.024 | 41.24 |
| | BTRC | 1.723 | 0.041 | 24.38 |
| | TUBB | 4.334 | 0.02 | 50.07 |
| | CDC37 | 4.101 | 0.093 | 10.73 |
| | YWHAH | 6.807 | 0.038 | 26.44 |
| | MAST3 | 4.683 | 0.109 | 9.182 |
| | BRAF | 2.061 | 0.049 | 20.38 |
| | YWHAB | 5.302 | 0.017 | 58.65 |
| | KSR1 | 1.73 | 0.108 | 9.25 |
| | YWHAE | 9.029 | 0.026 | 38.65 |
| | CUL1 | 27.92 | 0.042 | 23.99 |
| | MAPK7 | 3.783 | 0.061 | 16.39 |
| | YWHAZ | 9.094 | 0.027 | 36.84 |
| | IGF1R | 3.686 | 0.057 | 17.64 |
| | IRS1 | 2.624 | 0.045 | 22.1 |
| | YWHAQ | 9.32 | 0.023 | 43.99 |
| | MST1R | 1.235 | 0.062 | 16.19 |
| | NTRK1 | 8.726 | 0.053 | 19.02 |
| | CBL | 19.28 | 0.083 | 12.09 |
| | TUBB | 4.334 | 0.02 | 50.07 |
| | YWHAB | 5.302 | 0.017 | 58.65 |
| | YWHAH | 6.807 | 0.038 | 26.44 |
| | GRB2 | 9.905 | 0.019 | 51.99 |
| | ABL1 | 5.805 | 0.034 | 29.8 |
| | SORBS2 | 2.606 | 0.057 | 17.65 |
| | BCAR1 | 3.383 | 0.054 | 18.62 |
| | YWHAE | 9.029 | 0.026 | 38.65 |
| | VCP | 146.7 | 0.25 | 3.994 |
| | CDK2 | 28.82 | 0.124 | 8.086 |
| | CUL1 | 27.92 | 0.042 | 23.99 |
| | CDC37 | 4.101 | 0.093 | 10.73 |
| | LRRK2 | 7.159 | 0.052 | 19.28 |
| | YWHAE | 9.029 | 0.026 | 38.65 |

| | | | | |
|---|---|---|---|---|
| YWHAE-NUTM2A | LARP1 | 2.17 | 0.026 | 37.78 |
| | NOS2 | 4.671 | 0.032 | 31.26 |
| | YWHAQ | 9.32 | 0.023 | 43.99 |
| | YWHAG | 8.003 | 0.029 | 33.99 |
| | HUWE1 | 7.079 | 0.016 | 63.99 |
| | MAST2 | 2.442 | 0.122 | 8.19 |
| | NTRK1 | 8.726 | 0.053 | 19.02 |
| | VCP | 146.7 | 0.25 | 3.994 |
| | AKT1 | 2.106 | 0.035 | 28.49 |
| | MAP2K1 | 3.938 | 0.067 | 14.98 |
| | FBXW11 | 3.809 | 0.05 | 19.95 |
| | ARAF | 6.317 | 0.073 | 13.77 |
| | CUL3 | 3.375 | 0.048 | 20.74 |
| | PARK2 | 102 | 0.251 | 3.99 |
| | RAF1 | 6.385 | 0.043 | 23.49 |
| | YWHAZ | 9.094 | 0.027 | 36.84 |
| | UBXN1 | 5.639 | 0.042 | 23.76 |
| | TP53 | 240.5 | 0.25 | 3.996 |
| | RUVBL2 | 5.408 | 0.024 | 41.24 |
| | BTRC | 1.723 | 0.041 | 24.38 |
| | TUBB | 4.334 | 0.02 | 50.07 |
| | CDC37 | 4.101 | 0.093 | 10.73 |
| | YWHAH | 6.807 | 0.038 | 26.44 |
| | MAST3 | 4.683 | 0.109 | 9.182 |
| | BRAF | 2.061 | 0.049 | 20.38 |
| | YWHAB | 5.302 | 0.017 | 58.65 |
| | KSR1 | 1.73 | 0.108 | 9.25 |
| | YWHAE | 9.029 | 0.026 | 38.65 |
| | CUL1 | 27.92 | 0.042 | 23.99 |
| | MAPK7 | 3.783 | 0.061 | 16.39 |
| | YWHAZ | 9.094 | 0.027 | 36.84 |
| | IGF1R | 3.686 | 0.057 | 17.64 |
| | IRS1 | 2.624 | 0.045 | 22.1 |
| | YWHAQ | 9.32 | 0.023 | 43.99 |
| | MST1R | 1.235 | 0.062 | 16.19 |
| | NTRK1 | 8.726 | 0.053 | 19.02 |
| | CBL | 19.28 | 0.083 | 12.09 |
| | TUBB | 4.334 | 0.02 | 50.07 |
| | YWHAB | 5.302 | 0.017 | 58.65 |
| | YWHAH | 6.807 | 0.038 | 26.44 |
| | GRB2 | 9.905 | 0.019 | 51.99 |
| | ABL1 | 5.805 | 0.034 | 29.8 |
| | SORBS2 | 2.606 | 0.057 | 17.65 |
| | BCAR1 | 3.383 | 0.054 | 18.62 |
| | YWHAE | 9.029 | 0.026 | 38.65 |
| | VCP | 146.7 | 0.25 | 3.994 |
| | CDK2 | 28.82 | 0.124 | 8.086 |
| | CUL1 | 27.92 | 0.042 | 23.99 |
| | CDC37 | 4.101 | 0.093 | 10.73 |
| | LRRK2 | 7.159 | 0.052 | 19.28 |
| | YWHAE | 9.029 | 0.026 | 38.65 |
| **CA** | | | | |

| | | | | |
|---|---|---|---|---|
| ARGLU1-CXCR4 | APP | 1.25 | 0.139 | 7.2 |
| | CHERP | 2.649 | 0.046 | 21.5 |
| | SNRNP70 | 2.622 | 0.021 | 47.7 |
| | SRPK1 | 8.282 | 0.037 | 26.7 |
| | SRPK2 | 112.5 | 0.251 | 3.99 |
| | PTK2 | 5.022 | 0.044 | 22.7 |
| | JAK2 | 1.449 | 0.035 | 28.3 |
| | SOCS3 | 5.134 | 0.073 | 13.6 |
| | PTPN11 | 5.675 | 0.054 | 18.7 |
| | NTRK1 | 8.726 | 0.053 | 19 |
| | PTK2 | 5.022 | 0.044 | 22.7 |
| | ELAVL1 | 5.706 | 0.027 | 37.3 |
| | SRPK1 | 8.282 | 0.037 | 26.7 |
| | NTRK1 | 8.726 | 0.053 | 19 |
| | PTK2 | 5.022 | 0.044 | 22.7 |
| | JAK2 | 1.449 | 0.035 | 28.3 |
| | JAK3 | 6.989 | 0.097 | 10.3 |
| | SOCS3 | 5.134 | 0.073 | 13.6 |
| | PTPN11 | 5.675 | 0.054 | 18.7 |
| | STAM | 3.871 | 0.067 | 15 |
| ATXN10-FBLN1 | FN1 | 105.5 | 4.587 | 0.22 |
| | ATXN10 | 3.474 | 0.109 | 9.21 |
| | EGFR | 8.01 | 0.01 | 104 |
| | GSTK1 | 4.688 | 0.104 | 9.6 |
| | ATXN10 | 3.474 | 0.109 | 9.21 |
| | VCP | 146.7 | 0.25 | 3.99 |
| | BSG | 12.04 | 0.096 | 10.4 |
| | CUL3 | 3.375 | 0.048 | 20.7 |
| | VCP | 146.7 | 0.25 | 3.99 |
| | ABCE1 | 11.03 | 0.047 | 21.4 |
| | ATXN10 | 3.474 | 0.109 | 9.21 |
| | APP | 1.25 | 0.139 | 7.2 |
| | YWHAQ | 6.807 | 0.038 | 26.4 |
| | CUL3 | 3.375 | 0.048 | 20.7 |
| BCAS3-NFS1 | CTBP1 | 98.54 | 0.666 | 1.5 |
| | CTBP2 | 9.352 | 0.094 | 10.6 |
| | BCAS3 | 3.52 | 0.11 | 9.09 |
| | KAT2B | 6.538 | 0.037 | 26.8 |
| | CTBP1 | 98.54 | 0.666 | 1.5 |
| | CTBP2 | 9.352 | 0.094 | 10.6 |
| | CDC23 | 5.183 | 0.051 | 19.5 |
| | KAT2B | 6.538 | 0.037 | 26.8 |
| | BCAS3 | 3.52 | 0.11 | 9.09 |
| BCAS4-BCAS3 | CTBP1 | 98.54 | 0.666 | 1.5 |
| | CTBP2 | 9.352 | 0.094 | 10.6 |
| | BCAS3 | 3.52 | 0.11 | 9.09 |
| | KAT2B | 6.538 | 0.037 | 26.8 |
| | CTBP1 | 98.54 | 0.666 | 1.5 |
| | CTBP2 | 9.352 | 0.094 | 10.6 |
| | CDC23 | 5.183 | 0.051 | 19.5 |
| | KAT2B | 6.538 | 0.037 | 26.8 |
| | BCAS3 | 3.52 | 0.11 | 9.09 |
| BCL2L12- | NCOA2 | 3.388 | 0.053 | 18.9 |

| | | | | |
|---|---|---|---|---|
| PRMT1 | | | | |
| | NCOA3 | 3.966 | 0.037 | 26.7 |
| | NCOA1 | 3.675 | 0.036 | 27.5 |
| | TP53 | 240.5 | 0.25 | 4 |
| | ESR1 | 28.12 | 1.222 | 0.82 |
| | BRCA1 | 94.44 | 5.556 | 0.18 |
| | PRMT1 | 5.148 | 0.036 | 28.2 |
| | THRB | 3.31 | 0.069 | 14.5 |
| | CARM1 | 3.802 | 0.043 | 23.1 |
| | NR1I2 | 3.385 | 0.106 | 9.45 |
| | AR | 2.106 | 0.035 | 28.5 |
| | PPARA | 3.641 | 0.087 | 11.5 |
| | EP300 | 113.5 | 0.251 | 3.99 |
| | NCOA2 | 3.388 | 0.053 | 18.9 |
| | NCOA3 | 3.966 | 0.037 | 26.7 |
| | NCOA1 | 3.675 | 0.036 | 27.5 |
| | EP300 | 113.5 | 0.251 | 3.99 |
| | PARP1 | 88.81 | 0.406 | 2.47 |
| CAPNS1-WDR62 | YWHAZ | 9.094 | 0.027 | 36.8 |
| | FBXW11 | 3.809 | 0.05 | 20 |
| | FN1 | 105.5 | 4.587 | 0.22 |
| | HUWE1 | 7.079 | 0.016 | 64 |
| | YWHAQ | 6.807 | 0.038 | 26.4 |
| | GAPDH | 4.678 | 0.024 | 41.9 |
| | FERMT2 | 3.036 | 0.061 | 16.5 |
| | YWHAH | 6.807 | 0.038 | 26.4 |
| | VCAM1 | 110.8 | 0.251 | 3.99 |
| | YWHAB | 5.302 | 0.017 | 58.7 |
| | PAFAH1B1 | 3.751 | 0.064 | 15.7 |
| | YWHAG | 8.003 | 0.029 | 34 |
| | PAK2 | 4.364 | 0.061 | 16.5 |
| | YWHAE | 9.029 | 0.026 | 38.7 |
| | OGFOD1 | 1.833 | 0.057 | 17.5 |
| | MYO1E | 2.46 | 0.045 | 22.4 |
| | ASNS | 4.051 | 0.049 | 20.2 |
| | TBCB | 2.353 | 0.055 | 18.3 |
| | CAPN2 | 2.922 | 0.064 | 15.7 |
| | PROSC | 3.906 | 0.067 | 14.8 |
| | YWHAE | 9.029 | 0.026 | 38.7 |
| CCDC6-ANK3 | HDAC1 | 119.7 | 0.251 | 3.99 |
| | NR3C1 | 6.225 | 0.04 | 25.2 |
| | TRIM28 | 6.961 | 0.031 | 32 |
| | SF3A1 | 2.895 | 0.019 | 52.9 |
| | HNRNPR | 2.866 | 0.015 | 66.6 |
| | BRCC3 | 2.59 | 0.05 | 20.1 |
| | HDAC1 | 119.7 | 0.251 | 3.99 |
| | NR3C1 | 6.225 | 0.04 | 25.2 |
| | TRIM28 | 6.961 | 0.031 | 32 |
| | ELAVL1 | 5.706 | 0.027 | 37.3 |
| | SKP1 | 7.259 | 0.039 | 25.5 |
| | HNRNPR | 2.866 | 0.015 | 66.6 |
| | PPP1CA | 9.653 | 0.034 | 29.5 |
| | BRCC3 | 2.59 | 0.05 | 20.1 |
| | NTRK1 | 8.726 | 0.053 | 19 |

| | | | | |
|---|---|---|---|---|
| | SF3A1 | 2.895 | 0.019 | 52.9 |
| | CUL1 | 27.92 | 0.042 | 24 |
| | FBXW7 | 17.98 | 0.062 | 16 |
| CCDC9-DHX34 | EIF4A3 | 6.718 | 0.008 | 124 |
| | SNIP1 | 3.521 | 0.053 | 18.7 |
| | CCDC9 | 1.838 | 0.115 | 8.71 |
| | PRPF40A | 4.985 | 0.028 | 35.7 |
| CDC27-ST7L | CDC16 | 2.219 | 0.036 | 27.5 |
| | CDC27 | 3.771 | 0.038 | 26.5 |
| | MDC1 | 6.512 | 0.035 | 28.9 |
| | CDC20 | 4.633 | 0.031 | 31.9 |
| | CREBBP | 139.5 | 0.471 | 2.12 |
| | ANAPC2 | 1.684 | 0.038 | 26.1 |
| | ANAPC7 | 1.91 | 0.035 | 28.8 |
| | CREBBP | 139.5 | 0.471 | 2.12 |
| | E2F1 | 4.257 | 0.039 | 25.8 |
| | RB1 | 7.401 | 0.034 | 29.3 |
| | TFDP1 | 2.172 | 0.062 | 16.1 |
| | CDC16 | 2.219 | 0.036 | 27.5 |
| | CDC27 | 3.771 | 0.038 | 26.5 |
| | MDC1 | 6.512 | 0.035 | 28.9 |
| | SMAD2 | 22.89 | 0.222 | 4.5 |
| | TP53BP1 | 240.5 | 0.25 | 4 |
| | CREBBP | 139.5 | 0.471 | 2.12 |
| | UBE2S | 4.152 | 0.066 | 15.2 |
| | ANAPC2 | 1.684 | 0.038 | 26.1 |
| | ANAPC7 | 1.91 | 0.035 | 28.8 |
| CDK7-RIN3 | BRCA1 | 94.44 | 5.556 | 0.18 |
| | TP53 | 240.5 | 0.25 | 4 |
| | RUVBL2 | 5.408 | 0.024 | 41.2 |
| | SUPT5H | 4.864 | 0.04 | 25.3 |
| | ESR1 | 28.12 | 1.222 | 0.82 |
| | RPA1 | 116 | 0.251 | 3.99 |
| | HNRNPU | 126 | 0.251 | 3.99 |
| | RPA2 | 11.88 | 0.053 | 19 |
| | CDK2 | 28.82 | 0.124 | 8.09 |
| | POLR2A | 101.3 | 0.404 | 2.48 |
| | GTF2H1 | 3.282 | 0.062 | 16.1 |
| | RPA1 | 116 | 0.251 | 3.99 |
| | RPA2 | 11.88 | 0.053 | 19 |
| | POLR2A | 101.3 | 0.404 | 2.48 |
| | CCNH | 2.845 | 0.056 | 17.9 |
| | HDAC2 | 80.31 | 0.267 | 3.75 |
| | TP53 | 240.5 | 0.25 | 4 |
| | ESR1 | 28.12 | 1.222 | 0.82 |
| | MTA1 | 3.367 | 0.036 | 27.6 |
| | GTF2H1 | 3.282 | 0.062 | 16.1 |
| | ERCC3 | 2.875 | 0.063 | 16 |
| | POLR2A | 101.3 | 0.404 | 2.48 |
| | ERCC5 | 1.781 | 0.099 | 10.1 |
| | BRCA1 | 94.44 | 5.556 | 0.18 |
| | POLR2A | 101.3 | 0.404 | 2.48 |
| | RUVBL2 | 5.408 | 0.024 | 41.2 |
| | GTF2H1 | 3.282 | 0.062 | 16.1 |

| | | | | |
|---|---|---|---|---|
| | RPA1 | 116 | 0.251 | 3.99 |
| | RPA2 | 11.88 | 0.053 | 19 |
| | PRKCI | 4.328 | 0.063 | 15.9 |
| | APP | 1.25 | 0.139 | 7.2 |
| | CDK7 | 3.61 | 0.048 | 21.1 |
| | CDC37 | 4.101 | 0.093 | 10.7 |
| CHERP-CPAMD8 | DHX8 | 3.14 | 0.05 | 20.1 |
| | U2AF1 | 2.213 | 0.019 | 51.5 |
| | RPA1 | 116 | 0.251 | 3.99 |
| | PRPF40A | 4.985 | 0.028 | 35.7 |
| | RPA2 | 11.88 | 0.053 | 19 |
| | CHERP | 2.649 | 0.046 | 21.5 |
| | SF3A2 | 2.053 | 0.079 | 12.7 |
| | RBM39 | 3.092 | 0.021 | 46.6 |
| | EWSR1 | 10.05 | 0.016 | 64 |
| | DHX8 | 3.14 | 0.05 | 20.1 |
| | AGGF1 | 3.937 | 0.187 | 5.33 |
| | RNPS1 | 3.812 | 0.023 | 44.1 |
| | SNIP1 | 3.521 | 0.053 | 18.7 |
| | SF3B4 | 3.44 | 0.034 | 29.1 |
| | NTRK1 | 8.726 | 0.053 | 19 |
| | CHERP | 2.649 | 0.046 | 21.5 |
| | SRPK1 | 8.282 | 0.037 | 26.7 |
| | SRPK2 | 112.5 | 0.251 | 3.99 |
| | U2AF1 | 2.213 | 0.019 | 51.5 |
| | U2AF2 | 5.046 | 0.014 | 69.2 |
| | RPA1 | 116 | 0.251 | 3.99 |
| | APBB1 | 5.343 | 0.066 | 15.2 |
| | PRPF40A | 4.985 | 0.028 | 35.7 |
| | RPA2 | 11.88 | 0.053 | 19 |
| | TTC14 | 1.1 | 0.1 | 10 |
| | RBM23 | 4.659 | 0.116 | 8.59 |
| | SF3A2 | 2.053 | 0.079 | 12.7 |
| | SNRNP70 | 2.622 | 0.021 | 47.7 |
| | RBM39 | 3.092 | 0.021 | 46.6 |
| | EWSR1 | 10.05 | 0.016 | 64 |
| | WBP4 | 2.368 | 0.054 | 18.6 |
| CYTH1-PRPSAP1 | DDX17 | 2.559 | 0.018 | 55.5 |
| | DDX5 | 3.278 | 0.016 | 61.9 |
| | ILK | 200.6 | 0.951 | 1.05 |
| | COPS5 | 199 | 0.25 | 3.99 |
| | CYTH1 | 2.062 | 0.187 | 5.33 |
| | ARRB2 | 0.238 | 0.024 | 42 |
| | ARF6 | 4.542 | 0.103 | 9.69 |
| | ARRB1 | 0.282 | 0.04 | 24.9 |
| | DDX5 | 3.278 | 0.016 | 61.9 |
| | FBXW11 | 3.809 | 0.05 | 20 |
| | DDX17 | 2.559 | 0.018 | 55.5 |
| | ILK | 200.6 | 0.951 | 1.05 |
| | ITGB2 | 3.25 | 0.125 | 8 |
| | COPS5 | 199 | 0.25 | 3.99 |
| DLG1-CRYBG3 | DLG1 | 5.592 | 0.112 | 8.94 |
| | LIN7A | 3.508 | 0.121 | 8.27 |

| | | | | |
|---|---|---|---|---|
| | LIN7C | 2.864 | 0.082 | 12.2 |
| | APBA1 | 12.22 | 0.13 | 7.69 |
| | CASK | 6.782 | 0.109 | 9.14 |
| | DLG1 | 5.592 | 0.112 | 8.94 |
| | NTRK1 | 8.726 | 0.053 | 19 |
| | CASK | 6.782 | 0.109 | 9.14 |
| | EPB41 | 2.719 | 0.094 | 10.7 |
| | DLG1 | 5.592 | 0.112 | 8.94 |
| | KHDRBS1 | 6.273 | 0.039 | 25.7 |
| | LCK | 6.021 | 0.073 | 13.8 |
| | NTRK1 | 8.726 | 0.053 | 19 |
| | MAPK1 | 5.324 | 0.025 | 40.2 |
| | ARRB2 | 0.238 | 0.024 | 42 |
| | ARRB1 | 0.282 | 0.04 | 24.9 |
| DTX4-CCDC102B | MCM7 | 5.206 | 0.034 | 29 |
| | CDK18 | 5.579 | 0.101 | 9.86 |
| | LENG1 | 4.548 | 0.106 | 9.45 |
| | TRIM54 | 10.87 | 0.086 | 11.7 |
| | TRIM27 | 1.32 | 0.12 | 8.33 |
| | KIFC3 | 6.577 | 0.058 | 17.3 |
| | SFN | 8.356 | 0.033 | 29.9 |
| | MARK1 | 1.222 | 0.111 | 9 |
| | CCDC102B | 5.772 | 0.105 | 9.53 |
| EHD4-FSIP1 | EHD4 | 3.141 | 0.087 | 11.5 |
| | CTPS2 | 2.97 | 0.106 | 9.43 |
| | EHD1 | 5.281 | 0.081 | 12.3 |
| | EGFR | 8.01 | 0.01 | 104 |
| | NTRK1 | 8.726 | 0.053 | 19 |
| | WARS | 4.256 | 0.071 | 14.1 |
| | PLCG1 | 5.711 | 0.051 | 19.6 |
| | UBA2 | 4.141 | 0.053 | 18.8 |
| | ADSL | 4.562 | 0.071 | 14 |
| | UQCRC2 | 3.585 | 0.027 | 36.5 |
| | PLCG1 | 5.711 | 0.051 | 19.6 |
| | EHD4 | 3.141 | 0.087 | 11.5 |
| | EGFR | 8.01 | 0.01 | 104 |
| | NTRK1 | 8.726 | 0.053 | 19 |
| ELK4-SLC26A9 | BRCA1 | 94.44 | 5.556 | 0.18 |
| | MAPK3 | 13.19 | 0.075 | 13.3 |
| | MAPK1 | 5.324 | 0.025 | 40.2 |
| | ELK4 | 1.5 | 0.187 | 5.33 |
| | BRCA1 | 94.44 | 5.556 | 0.18 |
| | MAPK3 | 13.19 | 0.075 | 13.3 |
| | MAPK1 | 5.324 | 0.025 | 40.2 |
| | ELK4 | 1.5 | 0.187 | 5.33 |
| | BLM | 5.025 | 0.045 | 22.1 |
| ERAL1-DIDO1 | HNRNPDL | 2.214 | 0.02 | 49.2 |
| | RPA1 | 116 | 0.251 | 3.99 |
| | RPA2 | 11.88 | 0.053 | 19 |
| | HNRNPK | 3.744 | 0.015 | 64.9 |
| | RBM15 | 2.61 | 0.062 | 16.1 |
| | CUL3 | 3.375 | 0.048 | 20.7 |
| | DIDO1 | 2.75 | 0.086 | 11.6 |
| | FUS | 19.94 | 0.063 | 16 |

| | | | | |
|---|---|---|---|---|
| | FUS | 19.94 | 0.063 | 16 |
| | APP | 1.25 | 0.139 | 7.2 |
| | RPA1 | 116 | 0.251 | 3.99 |
| | RPA2 | 11.88 | 0.053 | 19 |
| | RBM15 | 2.61 | 0.062 | 16.1 |
| | CUL3 | 3.375 | 0.048 | 20.7 |
| | DIDO1 | 2.75 | 0.086 | 11.6 |
| | SRPK2 | 112.5 | 0.251 | 3.99 |
| GMDS-CCND3 | PCNA | 8.779 | 0.032 | 31.2 |
| | PPP1CC | 11.62 | 0.042 | 23.6 |
| | RBL2 | 3.341 | 0.055 | 18.3 |
| | PPP1CA | 9.653 | 0.034 | 29.5 |
| | CCND3 | 4.471 | 0.091 | 11 |
| | RB1 | 7.401 | 0.034 | 29.3 |
| | POLD1 | 3.859 | 0.063 | 15.8 |
| | CDK2 | 28.82 | 0.124 | 8.09 |
| | CDK4 | 7.033 | 0.049 | 20.3 |
| | CDK6 | 7.725 | 0.067 | 14.9 |
| | CREBBP | 139.5 | 0.471 | 2.12 |
| | GMDS | 3.025 | 0.138 | 7.27 |
| | NSFL1C | 4.352 | 0.057 | 17.7 |
| | CTH | 4.861 | 0.139 | 7.2 |
| | CAPN2 | 2.922 | 0.064 | 15.7 |
| | ATIC | 4.422 | 0.063 | 15.8 |
| | RARA | 5.002 | 0.046 | 21.6 |
| | NCOA2 | 3.388 | 0.053 | 18.9 |
| | VDR | 119.9 | 1.348 | 0.74 |
| | CCND3 | 4.471 | 0.091 | 11 |
| | CREBBP | 139.5 | 0.471 | 2.12 |
| | MCM10 | 3.093 | 0.063 | 15.8 |
| | RBX1 | 5.513 | 0.035 | 28.7 |
| | CCND3 | 4.471 | 0.091 | 11 |
| | APP | 1.25 | 0.139 | 7.2 |
| | NCOA2 | 3.388 | 0.053 | 18.9 |
| | RARA | 5.002 | 0.046 | 21.6 |
| | VDR | 119.9 | 1.348 | 0.74 |
| | CCND3 | 4.471 | 0.091 | 11 |
| | CREBBP | 139.5 | 0.471 | 2.12 |
| HJURP-EIF4E2 | FBXW11 | 3.809 | 0.05 | 20 |
| | TP53 | 240.5 | 0.25 | 4 |
| | GIGYF2 | 2.64 | 0.05 | 20.1 |
| | APP | 1.25 | 0.139 | 7.2 |
| | HUWE1 | 7.079 | 0.016 | 64 |
| | EIF4E2 | 5.17 | 0.086 | 11.6 |
| | YWHAB | 5.302 | 0.017 | 58.7 |
| | SHMT2 | 107 | 0.251 | 3.99 |
| | YWHAE | 9.029 | 0.026 | 38.7 |
| INTS4-GAB2 | SRC | 7.782 | 0.035 | 28.4 |
| | PLCG1 | 5.711 | 0.051 | 19.6 |
| | GRB2 | 9.905 | 0.019 | 52 |
| | ZAP70 | 2.313 | 0.048 | 20.8 |
| | NTRK1 | 8.726 | 0.053 | 19 |
| | SHC1 | 7.138 | 0.033 | 30 |
| | PIK3CB | 1.729 | 0.069 | 14.5 |

| | | | | |
|---|---|---|---|---|
| | PIK3R2 | 3.923 | 0.036 | 27.8 |
| | PIK3R1 | 5.515 | 0.038 | 26.3 |
| KDM5A-ANO2 | HDAC1 | 119.7 | 0.251 | 3.99 |
| | HDAC2 | 80.31 | 0.267 | 3.75 |
| | RBL1 | 3.75 | 0.054 | 18.4 |
| | TBP | 6.066 | 0.039 | 25.7 |
| | RB1 | 7.401 | 0.034 | 29.3 |
| | VDR | 119.9 | 1.348 | 0.74 |
| | KDM5A | 1.438 | 0.058 | 17.4 |
| | MORF4L1 | 4.827 | 0.046 | 22 |
| | HDAC2 | 80.31 | 0.267 | 3.75 |
| | EZH2 | 111.5 | 0.406 | 2.47 |
| | KDM5A | 1.438 | 0.058 | 17.4 |
| | ESR1 | 28.12 | 1.222 | 0.82 |
| MAPK10-FAM13A | HDAC1 | 119.7 | 0.251 | 3.99 |
| | TP53 | 240.5 | 0.25 | 4 |
| | HDAC9 | 2.523 | 0.017 | 57.5 |
| | JUN | 6.184 | 0.029 | 34.1 |
| | DDX5 | 3.278 | 0.016 | 61.9 |
| | ELK1 | 2.68 | 0.099 | 10.1 |
| | MAPK10 | 4.019 | 0.093 | 10.7 |
| | RELA | 7.966 | 0.03 | 33.3 |
| | CREBBP | 139.5 | 0.471 | 2.12 |
| | ATF2 | 9.133 | 0.045 | 22.1 |
| | APP | 1.25 | 0.139 | 7.2 |
| | MAPK10 | 4.019 | 0.093 | 10.7 |
| | MAP2K4 | 2.703 | 0.068 | 14.8 |
| MAPRE1-TM9SF4 | YWHAZ | 9.094 | 0.027 | 36.8 |
| | FN1 | 105.5 | 4.587 | 0.22 |
| | APP | 1.25 | 0.139 | 7.2 |
| | TUBB | 4.334 | 0.02 | 50.1 |
| | NTRK1 | 8.726 | 0.053 | 19 |
| | VCAM1 | 110.8 | 0.251 | 3.99 |
| | UNK | 14.91 | 0.051 | 19.6 |
| | COPS5 | 199 | 0.25 | 3.99 |
| | CDK5RAP2 | 5.3 | 0.1 | 10 |
| | PRKACA | 8.583 | 0.067 | 14.9 |
| | AKAP9 | 5.377 | 0.094 | 10.6 |
| | PRKACB | 3.854 | 0.066 | 15.1 |
| | CLIP1 | 2.941 | 0.123 | 8.16 |
| | TUBB | 4.334 | 0.02 | 50.1 |
| | TUBA1A | 7.605 | 0.037 | 27.4 |
| | HDAC6 | 3.869 | 0.027 | 37.5 |
| | PDE4DIP | 7.234 | 0.096 | 10.4 |
| | PRKACA | 8.583 | 0.067 | 14.9 |
| | PRKACB | 3.854 | 0.066 | 15.1 |
| | CDK5RAP2 | 5.3 | 0.1 | 10 |
| | AKAP9 | 5.377 | 0.094 | 10.6 |
| | TERF1 | 15.75 | 0.051 | 19.5 |
| | SPTAN1 | 4.962 | 0.033 | 30 |
| | DST | 4.014 | 0.074 | 13.5 |
| | MAPRE1 | 6.158 | 0.039 | 25.3 |
| NUMB- | TP53 | 240.5 | 0.25 | 4 |

| | | | | |
|---|---|---|---|---|
| ALDH6A1 | | | | |
| | NUMB | 2.828 | 0.074 | 13.4 |
| | ITCH | 7.348 | 0.045 | 22.3 |
| | MDM2 | 9.935 | 0.053 | 19 |
| | EGFR | 8.01 | 0.01 | 104 |
| | EPS15 | 4.535 | 0.035 | 28.9 |
| | EGFR | 8.01 | 0.01 | 104 |
| | AP2A1 | 2.521 | 0.03 | 33.7 |
| | NUMB | 2.828 | 0.074 | 13.4 |
| | PRKCZ | 4.7 | 0.059 | 16.8 |
| | NUMB | 2.828 | 0.074 | 13.4 |
| | APP | 1.25 | 0.139 | 7.2 |
| | EGFR | 8.01 | 0.01 | 104 |
| PARD6B-CD48 | PRKCI | 4.328 | 0.063 | 15.9 |
| | RASSF8 | 4.091 | 0.093 | 10.8 |
| | PARD3 | 3.66 | 0.061 | 16.4 |
| | PARD6G | 1.663 | 0.098 | 10.2 |
| | APP | 1.25 | 0.139 | 7.2 |
| | PARD6B | 8.532 | 0.114 | 8.79 |
| | PARD6A | 4.967 | 0.09 | 11.1 |
| | YWHAH | 6.807 | 0.038 | 26.4 |
| | PRKCZ | 4.7 | 0.059 | 16.8 |
| | WWC1 | 2.441 | 0.066 | 15.2 |
| | PRKCI | 4.328 | 0.063 | 15.9 |
| | PARD3 | 3.66 | 0.061 | 16.4 |
| | PARD6G | 1.663 | 0.098 | 10.2 |
| | APP | 1.25 | 0.139 | 7.2 |
| | PARD6B | 8.532 | 0.114 | 8.79 |
| | PARD6A | 4.967 | 0.09 | 11.1 |
| | YWHAH | 6.807 | 0.038 | 26.4 |
| | PRKCZ | 4.7 | 0.059 | 16.8 |
| | RAC1 | 13.31 | 0.088 | 11.4 |
| | PARD6G | 1.663 | 0.098 | 10.2 |
| | PARD6B | 8.532 | 0.114 | 8.79 |
| | PARD6A | 4.967 | 0.09 | 11.1 |
| PPP1R12A-MGAT4C | KDM1A | 32.43 | 0.154 | 6.48 |
| | ELAVL1 | 5.706 | 0.027 | 37.3 |
| | RPA1 | 116 | 0.251 | 3.99 |
| | RPA2 | 11.88 | 0.053 | 19 |
| | PPP1R12A | 2.913 | 0.046 | 22 |
| | CUL1 | 27.92 | 0.042 | 24 |
| | KDM1A | 32.43 | 0.154 | 6.48 |
| | NUDT5 | 2.125 | 0.133 | 7.53 |
| | TP53 | 240.5 | 0.25 | 4 |
| | ELAVL1 | 5.706 | 0.027 | 37.3 |
| | PUS1 | 3.795 | 0.108 | 9.22 |
| | NUAK1 | 2.19 | 0.095 | 10.5 |
| | AARSD1 | 2.984 | 0.063 | 15.8 |
| | RPA1 | 116 | 0.251 | 3.99 |
| | NTRK1 | 8.726 | 0.053 | 19 |
| | RPA2 | 11.88 | 0.053 | 19 |
| | RPRD1B | 3.023 | 0.044 | 22.5 |
| | PPP1R12A | 2.913 | 0.046 | 22 |
| | TRIM47 | 2.143 | 0.143 | 7 |

| | | | | |
|---|---|---|---|---|
| | PAXIP1 | 13.28 | 0.052 | 19.3 |
| | ACTR3 | 2.425 | 0.03 | 33 |
| | CUL1 | 27.92 | 0.042 | 24 |
| RNF11-C8A | CBLB | 3.457 | 0.079 | 12.7 |
| | RNF11 | 4.744 | 0.052 | 19.4 |
| | ITCH | 7.348 | 0.045 | 22.3 |
| | SMAD4 | 25.62 | 0.249 | 4.02 |
| | EPN1 | 4 | 0.063 | 15.8 |
| | RABGEF1 | 4.564 | 0.127 | 7.89 |
| | UBE2E1 | 11.81 | 0.104 | 9.65 |
| | UBE2D3 | 12.18 | 0.06 | 16.7 |
| | UBE2E3 | 8.567 | 0.097 | 10.3 |
| | HGS | 11.27 | 0.056 | 17.8 |
| | GGA1 | 4.433 | 0.089 | 11.3 |
| | AKT1 | 2.106 | 0.035 | 28.5 |
| | GGA3 | 2.646 | 0.08 | 12.5 |
| | GGA2 | 3.152 | 0.096 | 10.5 |
| | AP2A1 | 2.521 | 0.03 | 33.7 |
| | EPN3 | 4.833 | 0.173 | 5.79 |
| | UBE2D1 | 11.54 | 0.047 | 21.1 |
| | AP2B1 | 3.636 | 0.039 | 25.9 |
| | CSNK2A1 | 95.74 | 0.251 | 3.99 |
| | SMURF1 | 7.662 | 0.03 | 32.9 |
| | EPS15 | 4.535 | 0.035 | 28.9 |
| | SMURF2 | 5.487 | 0.064 | 15.7 |
| | UBQLN2 | 4.183 | 0.053 | 18.9 |
| | STAM2 | 4.296 | 0.072 | 14 |
| | NEDD4 | 18.77 | 0.067 | 15 |
| | UBQLN4 | 16.78 | 0.091 | 11 |
| | NEDD4L | 15.54 | 0.089 | 11.2 |
| | APP | 1.25 | 0.139 | 7.2 |
| | RNF11 | 4.744 | 0.052 | 19.4 |
| | PSMD4 | 3.411 | 0.022 | 46.3 |
| | PSMD7 | 2.016 | 0.019 | 51.6 |
| | PSMD6 | 1.459 | 0.017 | 58.9 |
| | PSMD11 | 2.596 | 0.02 | 49.7 |
| | PSMD10 | 2.101 | 0.033 | 30.5 |
| | PSMD3 | 2.201 | 0.018 | 55 |
| | PSMD12 | 1.493 | 0.016 | 61.6 |
| | PSMD13 | 1.499 | 0.017 | 59.4 |
| | USP14 | 1.978 | 0.03 | 33.4 |
| | PSMD14 | 1.925 | 0.018 | 55.1 |
| | PSMD1 | 1.91 | 0.016 | 61.3 |
| | PSMD2 | 3.15 | 0.02 | 51.1 |
| | APP | 1.25 | 0.139 | 7.2 |
| | GGA1 | 4.433 | 0.089 | 11.3 |
| | GGA3 | 2.646 | 0.08 | 12.5 |
| | GGA2 | 3.152 | 0.096 | 10.5 |
| | APP | 1.25 | 0.139 | 7.2 |
| | AKT1 | 2.106 | 0.035 | 28.5 |
| | RNF11 | 4.744 | 0.052 | 19.4 |
| | TBK1 | 4.968 | 0.045 | 22.3 |
| | APP | 1.25 | 0.139 | 7.2 |
| | NEDD4 | 18.77 | 0.067 | 15 |

| | | | | |
|---|---|---|---|---|
| SIPA1L3-WDR62 | MAPK10 | 4.019 | 0.093 | 10.7 |
| | WDR62 | 3.977 | 0.114 | 8.8 |
| | MAPK8 | 9.041 | 0.053 | 19 |
| | MAPK9 | 8.342 | 0.086 | 11.6 |
| | SFN | 8.356 | 0.033 | 29.9 |
| | YWHAB | 5.302 | 0.017 | 58.7 |
| | SIPA1L3 | 1.757 | 0.08 | 12.5 |
| | YWHAQ | 6.807 | 0.038 | 26.4 |
| | YWHAB | 5.302 | 0.017 | 58.7 |
| | FBXW11 | 3.809 | 0.05 | 20 |
| | MAPK10 | 4.019 | 0.093 | 10.7 |
| | ELAVL1 | 5.706 | 0.027 | 37.3 |
| | WDR62 | 3.977 | 0.114 | 8.8 |
| | TBP | 6.066 | 0.039 | 25.7 |
| | MAPK8 | 9.041 | 0.053 | 19 |
| | MAPK9 | 8.342 | 0.086 | 11.6 |
| SLC26A6-PRKAR2A | AKAP7 | 1.25 | 0.139 | 7.2 |
| | AKAP9 | 5.377 | 0.094 | 10.6 |
| | PRKAR2A | 3.894 | 0.081 | 12.3 |
| | PRKAR2B | 4.793 | 0.096 | 10.4 |
| | PRKACA | 8.583 | 0.067 | 14.9 |
| | PRKACB | 3.854 | 0.066 | 15.1 |
| | PRKAR2A | 3.894 | 0.081 | 12.3 |
| | AKAP7 | 1.25 | 0.139 | 7.2 |
| | PRKACA | 8.583 | 0.067 | 14.9 |
| | PRKACB | 3.854 | 0.066 | 15.1 |
| | PRKAR2B | 4.793 | 0.096 | 10.4 |
| | GCH1 | 6.08 | 0.203 | 4.93 |
| ST14-APLP2 | BRCA1 | 94.44 | 5.556 | 0.18 |
| | APLP2 | 3.615 | 0.164 | 6.09 |
| | ETS1 | 2.767 | 0.053 | 18.8 |
| | JUN | 6.184 | 0.029 | 34.1 |
| | SFN | 8.356 | 0.033 | 29.9 |
| | HDAC5 | 7.535 | 0.022 | 45 |
| | APLP2 | 3.615 | 0.164 | 6.09 |
| | RPL26 | 1.144 | 0.012 | 80.4 |
| | BRCA1 | 94.44 | 5.556 | 0.18 |
| | JUNB | 3.213 | 0.078 | 12.8 |
| | JUN | 6.184 | 0.029 | 34.1 |
| | APBB1 | 5.343 | 0.066 | 15.2 |
| | APBB2 | 3.937 | 0.146 | 6.86 |
| | KAT5 | 32.43 | 0.154 | 6.48 |
| | ETS1 | 2.767 | 0.053 | 18.8 |
| | APLP2 | 3.615 | 0.164 | 6.09 |
| | MAPK8 | 9.041 | 0.053 | 19 |
| STRADB-NOP58 | NIFK | 3.255 | 0.023 | 43 |
| | NOP56 | 3.419 | 0.014 | 72.8 |
| | HNRNPU | 126 | 0.251 | 3.99 |
| | RUVBL2 | 5.408 | 0.024 | 41.2 |
| | NOLC1 | 3.131 | 0.032 | 31 |
| | SNU13 | 3.398 | 0.021 | 48.3 |
| | NTRK1 | 8.726 | 0.053 | 19 |

| | | | | |
|---|---|---|---|---|
| | PUM3 | 1.326 | 0.02 | 49 |
| | RPS15A | 1.32 | 0.008 | 127 |
| | RSL1D1 | 1.545 | 0.027 | 36.9 |
| | KRR1 | 1.636 | 0.026 | 38.5 |
| | DDX18 | 1.737 | 0.018 | 56.4 |
| | EIF6 | 4.407 | 0.031 | 32.2 |
| | DHX15 | 1.947 | 0.024 | 42.1 |
| | RPS4X | 1.488 | 0.007 | 134 |
| | EED | 197.6 | 2.994 | 0.33 |
| | PRPF3 | 2.727 | 0.028 | 35.2 |
| | TARDBP | 3.859 | 0.013 | 77.5 |
| | RPL11 | 1.378 | 0.008 | 127 |
| | EIF2S2 | 3.9 | 0.06 | 16.7 |
| | DDX27 | 1.525 | 0.026 | 38 |
| | NOP58 | 1.809 | 0.018 | 55.8 |
| | FN1 | 105.5 | 4.587 | 0.22 |
| | DDX24 | 2.62 | 0.023 | 43.9 |
| | U2AF1 | 2.213 | 0.019 | 51.5 |
| | RPL30 | 1.104 | 0.008 | 119 |
| | ESR1 | 28.12 | 1.222 | 0.82 |
| | DDX56 | 1.947 | 0.024 | 42.1 |
| | FTSJ3 | 2.13 | 0.022 | 46.5 |
| | DDX47 | 2.772 | 0.06 | 16.6 |
| | KPNA6 | 3.831 | 0.07 | 14.4 |
| | GTPBP4 | 1.916 | 0.022 | 45.4 |
| | WDR36 | 2.158 | 0.038 | 26.4 |
| | KPNA1 | 4.85 | 0.048 | 20.8 |
| | DKC1 | 2.607 | 0.027 | 37.6 |
| | RRP12 | 1.834 | 0.031 | 32.7 |
| | BOP1 | 2.56 | 0.053 | 18.8 |
| | FBL | 4.035 | 0.018 | 56.3 |
| | TBL3 | 1.752 | 0.037 | 26.8 |
| | WDR36 | 2.158 | 0.038 | 26.4 |
| | NOP58 | 1.809 | 0.018 | 55.8 |
| | DHX15 | 1.947 | 0.024 | 42.1 |
| | NOP56 | 3.419 | 0.014 | 72.8 |
| STX16-RAE1 | FBXW11 | 3.809 | 0.05 | 20 |
| | NXF1 | 1.244 | 0.02 | 49 |
| | FAF1 | 5.636 | 0.045 | 22 |
| | RAE1 | 6.302 | 0.053 | 19 |
| | ILF3 | 1.727 | 0.027 | 37.6 |
| | CUL1 | 27.92 | 0.042 | 24 |
| | HNRNPUL1 | 3.25 | 0.029 | 34.5 |
| | CUL3 | 3.375 | 0.048 | 20.7 |
| | NXF1 | 1.244 | 0.02 | 49 |
| | CUL1 | 27.92 | 0.042 | 24 |
| | ILF3 | 1.727 | 0.027 | 37.6 |
| | CUL3 | 3.375 | 0.048 | 20.7 |
| TANC2-CHD6 | ZFYVE9 | 4.194 | 0.108 | 9.3 |
| | PPP1CC | 11.62 | 0.042 | 23.6 |
| | PPP1CA | 9.653 | 0.034 | 29.5 |
| | TANC2 | 1.637 | 0.205 | 4.89 |
| | ZFYVE9 | 4.194 | 0.108 | 9.3 |
| | PPP1CC | 11.62 | 0.042 | 23.6 |

| | | | | |
|---|---|---|---|---|
| | PPP1CA | 9.653 | 0.034 | 29.5 |
| | TANC2 | 1.637 | 0.205 | 4.89 |
| | MAEA | 1.425 | 0.079 | 12.6 |
| | RANBP9 | 6.682 | 0.06 | 16.6 |
| | MKLN1 | 2.003 | 0.08 | 12.5 |
| | MMP7 | 1.84 | 0.08 | 12.5 |
| | RMND5A | 2.158 | 0.065 | 15.3 |
| | MAEA | 1.425 | 0.079 | 12.6 |
| | RANBP9 | 6.682 | 0.06 | 16.6 |
| | MKLN1 | 2.003 | 0.08 | 12.5 |
| | RANBP10 | 3.066 | 0.123 | 8.15 |
| | MMP7 | 1.84 | 0.08 | 12.5 |
| | RMND5A | 2.158 | 0.065 | 15.3 |
| TMPRSS2-ERG | CDC5L | 145.2 | 0.25 | 3.99 |
| | DDX3X | 3.219 | 0.022 | 45.4 |
| | ELAVL1 | 5.706 | 0.027 | 37.3 |
| | CAD | 2.517 | 0.025 | 39.7 |
| | NEDD4 | 18.77 | 0.067 | 15 |
| | SF3B1 | 3.067 | 0.017 | 57.7 |
| | PARP1 | 88.81 | 0.406 | 2.47 |
| | PRKDC | 5.478 | 0.023 | 42.7 |
| | PRPF8 | 2.684 | 0.018 | 57 |
| | SFPQ | 3.346 | 0.021 | 48.4 |
| | ERG | 2.976 | 0.038 | 26.6 |
| | POLR2A | 101.3 | 0.404 | 2.48 |
| | TOP1 | 3.021 | 0.021 | 48.3 |
| | CLTC | 6.885 | 0.025 | 40.2 |
| | SF3B2 | 1.773 | 0.068 | 14.7 |
| | XRCC5 | 152.1 | 0.576 | 1.74 |
| | XRCC6 | 232.4 | 0.88 | 1.14 |
| | DDX23 | 2.608 | 0.038 | 26.5 |
| | SNRNP200 | 2.819 | 0.019 | 53.9 |
| | DDX21 | 2.554 | 0.023 | 43.5 |
| | TUBB | 4.334 | 0.02 | 50.1 |
| | NONO | 4.27 | 0.03 | 33.7 |
| | JUN | 6.184 | 0.029 | 34.1 |
| | HNRNPU | 126 | 0.251 | 3.99 |
| | PRPF40A | 4.985 | 0.028 | 35.7 |
| | AR | 2.106 | 0.035 | 28.5 |
| | NCL | 3.53 | 0.015 | 64.6 |
| | HNRNPM | 3.043 | 0.013 | 78.9 |
| | HNRNPC | 3.27 | 0.02 | 50.1 |
| | TOP2B | 2.37 | 0.038 | 26.2 |
| | ILF3 | 1.727 | 0.027 | 37.6 |
| | ILF2 | 2.535 | 0.039 | 25.6 |
| | PRPF8 | 2.684 | 0.018 | 57 |
| | ERG | 2.976 | 0.038 | 26.6 |
| | SF3B2 | 1.773 | 0.068 | 14.7 |
| | SF3B1 | 3.067 | 0.017 | 57.7 |
| | PARP1 | 88.81 | 0.406 | 2.47 |
| | PRKDC | 5.478 | 0.023 | 42.7 |

Table S11: Communities as predicted using scores

| LK,LY,ME,GL | |
|---|---|
| | Community |
| BCR-ABL1 | |
| RUNX1-RUNX1T1 | HDAC1 BRCA1 KMT2A SMARCC1 HDAC2 CREBBP CTBP1 |
| | EP300 SMARCA4 NCOR2 NCOR1 |
| | VDR EP300 SMARCA4 CREBBP |
| KMT2A-MLLT10 | SMARCC2 KMT2A HDAC2 CHD3 SMARCC1 POLR2A |
| | SMARCA2 CREBBP SIN3A |
| | CTBP1 KMT2A CREBBP |
| IGH-BCL2 | TP53 CASP3 PARP1 CASP8 HIF1A BCL2 |
| | BAG3 PARP1 BCL2 |
| KMT2A-AFF1 | SMARCC2 KMT2A HDAC2 CHD3 SMARCC1 POLR2A |
| | SMARCA2 CREBBP SIN3A |
| | CTBP1 KMT2A CREBBP |
| PICALM-MLLT10 | FN1 EEF1A1 EGFR NTRK1 PLCG1 DNM2 ILVBL PICALM |
| | HNRNPD FUS FN1 DDX1 |
| | SEC24D PICALM NTRK1 SEC24C |
| PML-RARA | NCOA2 NR3C1 NR4A1 KAT2B RXRA PPARG TP53 MDM2 EP300 |
| | SMARCA4 RELA RARA NCOA3 NCOA1 STAT3 ARNT NPAS2 |
| | PARP1 NFKB1 TRIP4 CREBBP NFKB1 EP300 PARP1 |
| KMT2A-MLLT3 | SMARCC2 KMT2A HDAC2 CHD3 SMARCC1 POLR2A SMARCA2 |
| | CREBBP SIN3A CTBP1 KMT2A CREBBP |
| KMT2A-AFDN | SMARCC2 KMT2A HDAC2 CHD3 SMARCC1 POLR2A SMARCA2 |
| | CREBBP SIN3A |
| | CTBP1 KMT2A CREBBP |
| CBFB-MYH11 | ACTA2 MYH11 MYO1E RPA1 RPA2 ACTB |
| | RPA1 ELAVL1 RPA2 |
| IGH-MYC | |
| NUP98-DDX10 | SIRT7 DDX10 APP DDX56 NTRK1 DDX54 PUM3 PWP1 CSNK2A1 |
| | HDAC1 CTNNB1 MAPK8 EP300 CREBBP |
| | PUM3 NXF1 EED KPNB1 HNRNPUL1 |
| | HDAC1 CREBBP APC CSNK2A1 |
| | CTNNB1 APC CREBBP |
| | USP7 NTRK1 CDC37 |
| PCM1-JAK2 | ERBB2 ERBB3 VAV1 TEC EGFR INSR PLCG1 JAK2 |
| | STAT5A JAK2 INSR |
| KMT2A-SEPT9 | SMARCC2 KMT2A HDAC2 CHD3 SMARCC1 POLR2A SMARCA2 |
| | CREBBP SIN3A |
| | CTBP1 KMT2A CREBBP |
| FUS-ERG | RPA1 SF3B2 PRKDC PRPF8 SF3A2 DHX15 RPA2 CUL3 |
| | ABL1 PARP1 PRKDC |
| NUP98-HOXA9 | HDAC1 TP53 SMAD4 CTNNB1 CSNK2A1 MAPK8 EP300 CREBBP |
| | CTNNB1 APC CREBBP |
| ETV6-ABL1 | ERBB2 CBLB UBASH3B SOS1 ERBB4 SRC VAV1 EGFR CBL |
| | PLCG1 PIK3R2 PIK3R1 ABL1 |
| | ABL1 ABL2 JAK1 |

| | |
|---|---|
| SET-NUP214 | NXF1 FAF1 SUPT5H GART CUL2 CUL3 |
| | NXF1 CUL2 RANBP2 |
| MNX1-ETV6 | HDAC3 ETV6 HDAC9 PIN1 NCOR1 SIN3A |
| | L3MBTL1 ETV7 ETV6 |
| KMT2A-MLLT1 | SMARCC2 KMT2A HDAC2 CHD3 SMARCC1 POLR2A |
| | SMARCA2 CREBBP SIN3A |
| KMT2A-MLLT6 | SMARCC2 KMT2A HDAC2 CHD3 SMARCC1 POLR2A |
| | SMARCA2 CREBBP SIN3A |
| | CTBP1 KMT2A CREBBP |
| ETV6-ACSL6 | HDAC3 ETV6 HDAC9 PIN1 NCOR1 SIN3A |
| | L3MBTL1 ETV7 ETV6 |
| ETV6-MECOM | HDAC1 UBE2I HDAC3 EHMT2 ELAVL1 |
| | SMAD3 KAT2B SUV39H1 NCOR1 CTBP1 |
| | MECOM SMAD1 SMAD2 |
| | CREBBP MECOM SUV39H1 HDAC4 |
| KMT2A-EPS15 | SMARCC2 KMT2A HDAC2 CHD3 SMARCC1 |
| | POLR2A SMARCA2 CREBBP SIN3A |
| | CTBP1 KMT2A CREBBP |
| KMT2A-GAS7 | SMARCC2 KMT2A HDAC2 CHD3 SMARCC1 |
| | POLR2A SMARCA2 CREBBP SIN3A |
| | CTBP1 KMT2A CREBBP |
| KMT2A-ABL1 | SMARCC2 KMT2A SMARCC1 CHD3 POLR2A |
| | SMARCA2 CREBBP ABL1 CBLB CBL |
| KMT2A-MLLT11 | SMARCC2 KMT2A HDAC2 CHD3 SMARCC1 |
| | POLR2A SMARCA2 CREBBP SIN3A |
| | CTBP1 KMT2A CREBBP |
| MN1-ETV6 | HDAC3 ETV6 HDAC9 PIN1 EP300 NCOR1 SIN3A |
| | HDAC3 EP300 KAT5 |
| KMT2A-MAML2 | SMARCC2 KMT2A SMARCC1 CHD3 SMARCA2 CREBBP |
| | SMARCA2 EP300 CREBBP |
| KMT2A-FOXO4 | SMARCC2 KMT2A HDAC2 CHD3 SMARCC1 |
| | POLR2A SMARCA2 CREBBP SIN3A |
| | CTBP1 KMT2A CREBBP |
| DEK-NUP214 | ESR1 KAT2B SMAD2 SMAD3 CDK2 DEK EP300 CREBBP |
| | NXF1 DHX15 CUL2 CUL3 |
| RUNX1-CBFA2T3 | KMT2A SMARCC1 SMARCA4 CREBBP |
| | EP300 CREBBP |
| NUP98-PSIP1 | HDAC1 KMT2A ESR1 CTNNB1 EP300 CREBBP |
| | NXF1 EIF4A3 SON |
| NUP98-HOXC13 | HDAC1 CTNNB1 MAPK8 EP300 CREBBP |
| | NXF1 HNRNPAB EED KPNB1 HNRNPUL1 |
| | HDAC1 CREBBP APC CSNK2A1 |
| | CTNNB1 APC CREBBP |
| NUP98-HOXC11 | HDAC1 STAT3 SP1 SMAD3 CTNNB1 MAPK8 EP300 CREBBP |
| | CTNNB1 APC CREBBP |
| PAX5-ETV6 | UBE2I HDAC3 TBP KAT5 RB1 PAX5 RUNX1 PIN1 EP300 NCOR1 |
| | MAPK1 EP300 HDAC6 |
| NUP98-HOXA11 | HDAC1 HDAC2 YY1 CTNNB1 CSNK2A1 MAPK8 EP300 CREBBP |

| | |
|---|---|
| | CTNNB1 APC CREBBP |
| BCR-PDGFRA | TGFBR2 PDGFRA SHC1 CRKL EGFR PLCG1 FES CRK ABL1 |
| | HCK PTPN6 BCR UBASH3B INPP5D SOS1 GRB2 NTRK1 |
| | CBL KIT DOK1 PIK3R2 PIK3R1 |
| | ABL1 TP53 RB1 BCR |
| BCR-FGFR1 | SRC ITK SOS1 ERBB3 VAV1 CBL PLCG1 ABL1 |
| | ABL1 HCK BCR CBL |
| NPM1-RARA | |
| KMT2A-CBL | |
| IGH-BCL6 | HDAC1 TP53 CTBP1 EP300 NCOR2 CREBBP |
| | HDAC2 SMARCA4 CREBBP |
| LCP1-BCL6 | HDAC1 TP53 CTBP1 EP300 NCOR2 CREBBP |
| | HDAC2 SMARCA4 CREBBP |
| CREBBP-KAT6A | SMARCC2 KMT2A HDAC2 CHD3 SMARCC1 POLR2A SMARCA2 |
| | CREBBP SIN3A |
| | CTBP1 KMT2A CREBBP |
| KMT2A-ARHGAP26 | SMARCC2 KMT2A HDAC2 CHD3 SMARCC1 POLR2A SMARCA2 |
| | CREBBP SIN3A |
| | CTBP1 KMT2A CREBBP |
| FOXO3-KMT2A | SMARCC2 KMT2A SMARCC1 CHD3 SMARCA2 CREBBP |
| | SMARCA2 EP300 CREBBP |
| KMT2A-DCPS | SMARCC2 KMT2A HDAC2 CHD3 SMARCC1 POLR2A SMARCA2 |
| | CREBBP SIN3A |
| | CTBP1 KMT2A CREBBP |
| KMT2A-EP300 | |
| IGH-CEBPE | UBE2I BATF DDIT3 CEBPG CEBPE FOS JUN STAT6 |
| | RB1 PIAS1 FOSL1 BATF3 BATF2 ATF4 MYB ATF3 |
| HSP90AA1-BCL6 | |
| **SC** | |
| ASPSCR1-TFE3 | |
| | xxxxxxxxxxxxxxxxxx |
| ASTN2-CNOT2 | CNOT6L AURKA CNOT8 TNRC6C TNRC6B CNOT3 CNOT2 CNOT1 CNOT7 AGO2 |
| | xxxxx |
| | HDAC3 GPS2 CNOT2 NCOR2 NCOR1 |
| | xxxxxxxxxxxxxxxxxx |
| BCOR-ZC3H7B | HDAC3 HDAC4 SP1 CTBP1 NACC1 NCOR2 |
| | xxxxx |
| | HDAC1 CTBP1 NCOR2 |
| | xxxxxxxxxxxxxxxxxx |
| BCOR-CCNB3 | HDAC3 HDAC4 SP1 CTBP1 NACC1 NCOR2 |
| | HDAC1 CTBP1 NCOR2 |

| | |
|---|---|
| | xxxxxxxxxxxxxxxxx |
| CDX1-IRF2BP2 | ELAVL1 NTRK1 IRF2BPL IRF2BP2 RBM39 |
| | xxxxxxxxxxxxxxxxx |
| CIC-DUX4 | |
| | xxxxxxxxxxxxxxxxx |
| CREB1-EWSR1 | BRCA1 EPAS1 MYOD1 ESR1 JUN POLR2A EP300 SMARCA4 EWSR1 CREBBP |
| | xxxxx |
| | NR3C1 EP300 SMARCA4 CREBBP |
| | xxxxx |
| | NONO SMARCA4 CUL3 |
| | xxxxxxxxxxxxxxxxx |
| CTDSP2-FAM19A2 | SETD1A POLR2A INTS6 CTDSP1 CTDSP2 |
| | xxxxxxxxxxxxxxxxx |
| CXorf67-MBTD1 | |
| | xxxxxxxxxxxxxxxxx |
| EPC1-PHF1 | HDAC1 DHX9 TP53 ELAVL1 E2F6 RBBP7 RBBP4 EZH1 EZH2 EED XRCC6 XRCC5 PHF1 |
| | xxxxx |
| | HDAC1 TP53 YEATS4 TRIM27 KAT5 TRIM23 HIST1H2BA DMAP1 MYC XRCC6 MORF4L1 ING3 |
| | xxxxxxxxxxxxxxxxx |
| ERG-EWSR1 | TP53 ESR1 PARP1 EP300 CREBBP XRCC6 |
| | xxxxx |
| | EPAS1 EP300 CREBBP |
| | xxxxxxxxxxxxxxxxx |
| ETV6-NTRK3 | PDGFRB SHC1 CRKL GAB2 NTRK1 PLCG1 GRB2 |
| | xxxxx |
| | HDAC3 PIN1 ETV6 |
| | xxxxxxxxxxxxxxxxx |
| EWSR1-ATF1 | PDGFRB SHC1 CRKL GAB2 NTRK1 PLCG1 GRB2 |
| | xxxxx |
| | HDAC3 PIN1 ETV6 |
| | xxxxxxxxxxxxxxxxx |
| EWSR1-FLI1 | BRCA1 POLR2A ESR1 EP300 KAT2B CREBBP |
| | xxxxx |
| | EPAS1 EP300 CREBBP |
| | xxxxxxxxxxxxxxxxx |
| EWSR1-NR4A3 | TSG101 DHX9 HDAC3 TRIM28 FUS RAD23A JUN PRMT1 ILK BMI1 ELK1 POLR2A CHERP ATXN3 EP300 IRF3 EPAS1 TP53 ESR1 NONO RPA1 NTRK1 RPA2 CUL4A CUL4B FASN CREBBP HBP1 CUL5 HLTF EWSR1 YBX1 CUL1 CUL2 CUL3 |
| | xxxxx |
| | HDAC2 CREBBP ESR1 |
| | xxxxx |
| | NONO FXR2 CUL3 |
| | xxxxxxxxxxxxxxxxx |
| EWSR1-ETV4 | TSG101 DHX9 HDAC3 RFWD2 FUS RAD23A JUN PRMT1 ILK SMAD2 BMI1 ELK1 POLR2A CHERP ATXN3 EP300 IRF3 EPAS1 TP53 ESR1 NONO RPA1 NTRK1 RPA2 CUL4A CUL4B FASN CREBBP HBP1 CUL5 HLTF EWSR1 YBX1 CUL1 CUL2 CUL3 |
| | xxxxx |

| | |
|---|---|
| | HDAC2 CREBBP ESR1 |
| | xxxxx |
| | NONO FXR2 CUL3 |
| | xxxxxxxxxxxxxxxxxx |
| EWSR1-PATZ1 | TSG101 DHX9 HDAC3 RFWD2 FUS RAD23A JUN PRMT1 ILK SMAD2 BMI1 ELK1 POLR2A CHERP ATXN3 EP300 IRF3 EPAS1 TP53 ESR1 NONO RPA1 NTRK1 RPA2 CUL4A CUL4B FASN CREBBP HBP1 CUL5 HLTF EWSR1 YBX1 CUL1 CUL2 CUL3 |
| | xxxxx |
| | HDAC2 CREBBP ESR1 |
| | xxxxx |
| | NONO FXR2 CUL3 |
| | xxxxxxxxxxxxxxxxxx |
| EWSR1-DDIT3 | HDAC1 EPAS1 HDAC3 DDIT3 CEBPB ESR1 FOS JUN HBP1 POLR2A IRF3 TP53 EP300 EWSR1 CREBBP |
| | xxxxx |
| | DHX9 CUL4A CUL4B CUL5 CUL1 CUL2 CUL3 |
| | xxxxxxxxxxxxxxxxxx |
| EWSR1-POU5F1 | IRF3 EPAS1 HDAC3 TP53 ESR1 JUN HBP1 ETS2 CTNNB1 POLR2A EP300 EWSR1 CREBBP |
| | xxxxx |
| | NONO CUL2 CUL3 |
| | xxxxxxxxxxxxxxxxxx |
| EWSR1-SP3 | HDAC1 EPAS1 HDAC3 CEBPB ESR1 JUN HBP1 POLR2A IRF3 TP53 RELA EP300 EWSR1 CREBBP |
| | xxxxx |
| | DHX9 CUL4A CUL4B CUL5 CUL1 CUL2 CUL3 |
| | xxxxxxxxxxxxxxxxxx |
| FOXO4-CIC | XPO1 CTNNB1 VDR ESR1 SMAD4 SMAD3 SFN AKT1 FOXO4 MDM2 NLK CREBBP |
| | xxxxxxxxxxxxxxxxxx |
| FUS-ERG | RPA1 SF3B2 PRKDC PRPF8 SF3A2 DHX15 RPA2 CUL3 |
| | ABL1 PARP1 PRKDC |
| | xxxxxxxxxxxxxxxxxx |
| FUS-CREB3L1 | NONO CUL4A CUL4B DHX15 CUL5 CUL1 CUL2 CUL3 |
| | VCP FBXW11 CUL1 |
| | xxxxxxxxxxxxxxxxxx |
| FUS-DDIT3 | HDAC1 DDX17 EPAS1 RELA ESR1 JUN TP73 EWSR1 CTNNB1 DDX5 CDK2 DDIT3 MDM2 EP300 TRIP4 CREBBP |
| | VCP FBXW11 CUL1 |
| | xxxxxxxxxxxxxxxxxx |
| FUS-ATF1 | HDAC1 DDX17 EPAS1 RELA ESR1 JUN TP73 EWSR1 CTNNB1 DDX5 CDK2 DDIT3 MDM2 EP300 TRIP4 CREBBP |
| | VCP FBXW11 CUL1 |
| | xxxxxxxxxxxxxxxxxx |
| FUS-CREB3L2 | NONO CUL4A CUL4B DHX15 CUL5 CUL1 CUL2 CUL3 |
| | VCP FBXW11 CUL1 |
| | xxxxxxxxxxxxxxxxxx |
| HEY1-NCOA2 | BRCA1 RARA NR3C1 VDR STAT6 HNF4A PRMT1 CARM1 RXRA PPARG PPARD AR PPARA ESR2 EP300 NCOA2 NCOA3 NCOA1 TP53 ESR1 AHR ARNT THRB THRA NR1I3 NR1I2 PGR CREBBP PIAS3 |

| | |
|---|---|
| | NCOA2 NCOA1 UBR5 |
| | xxxxxxxxxxxxxxxxxx |
| IRX2-TERT | YWHAZ AKT1 RPS6KB1 ENO1 MTOR MDM2 TERT XRCC6 |
| | TERF1 STUB1 TERT POT1 |
| | TERT MTOR YWHAQ RUVBL2 |
| | TPP1 TERT POT1 |
| | xxxxxxxxxxxxxxxxxx |
| JAZF1-SUZ12 | DHX9 RBM5 FBXW11 DDX3X NXF1 SF3B4 PRMT1 SF3B1 SF3B2 PRPF8 EED CRNKL1 RNPS1 UBE2I SNRNP200 SRSF7 SNRPD3 RALY DDX5 PRPF19 RNF2 SNRPA1 SON EFTUD2 U2AF1 SF3A1 EPRS ILF2 EIF4A3 CDC40 ILF3 RANBP2 |
| | DNMT3B HDAC1 HDAC2 TRIM28 CHD4 UHRF1 DNMT1 MTA1 NR2C2 EZH2 GATAD2B EED CBX5 RBBP4 CBX3 SETDB1 |
| | VCP BRCA1 MTOR NXF1 RUVBL2 |
| | VCP FBXW11 SKP1 BTRC EZH2 |
| | DHX9 ADAR EZH2 FBXW11 |
| | HDAC2 NXF1 JARID2 SETDB1 |
| | RELA FBXW11 BTRC |
| | BTRC CSNK2B NXF1 |
| | xxxxxxxxxxxxxxxxxx |
| JAZF1-PHF1 | DHX9 EZH1 PPARG EZH2 EED XRCC6 XRCC5 PHF1 |
| | HDAC1 PPARG EZH2 PHF1 |
| | PHF1 TP53 XRCC6 |
| | xxxxxxxxxxxxxxxxxx |
| LMNA-NTRK1 | |
| | xxxxxxxxxxxxxxxxxx |
| MEAF6-TRERF1 | HDAC1 TRERF1 KAT5 ING3 CREBBP YEATS4 EP300 MORF4L1 HIST1H2BA |
| | ELAVL1 TRERF1 KAT6A CREBBP NR5A1 |
| | SOX2 HDAC1 TRERF1 |
| | xxxxxxxxxxxxxxxxxx |
| MEAF6-PHF1 | DHX9 EZH1 EZH2 EED XRCC6 XRCC5 PHF1 |
| | PHF1 TP53 XRCC6 |
| | KAT6A TP53 ELAVL1 |
| | HDAC1 EZH2 PHF1 |
| | xxxxxxxxxxxxxxxxxx |
| NR4A3-TAF15 | TRIM28 FUS PRMT1 COPS6 COPS5 POLR2C POLR2A TAF15 SF1 NEDD8 POLR2E RPA1 RPA2 CUL4A CUL4B CUL5 CUL1 CUL2 CUL3 |
| | EZH2 TRIM28 CUL1 |
| | xxxxxxxxxxxxxxxxxx |
| NR4A3-TFG | CUL4A CUL4B CUL5 CUL1 CUL2 CUL3 |
| | TRIM28 CUL1 CUL3 |
| | xxxxxxxxxxxxxxxxxx |
| NR6A1-TRHDE | |
| | xxxxxxxxxxxxxxxxxx |
| NUP107-LGR5 | NUP153 KPNB1 NTRK1 CUL3 |
| | NUP153 EIF4B CUL3 |
| | EED KPNB1 TP53BP1 |
| | xxxxxxxxxxxxxxxxxx |
| PAPPA-NUP107 | NUP153 ELAVL1 KPNB1 NTRK1 SMAD3 VCP CUL3 EIF4B |
| | SMAD9 SKIL PAPPA SMAD2 SMAD3 |
| | TP53BP1 ELAVL1 EED KPNB1 |

| | |
|---|---|
| | NUP214 SMAD2 SMAD3 |
| | xxxxxxxxxxxxxxxxxx |
| PAX3-FOXO1 | NCOA1 ESR1 PARP1 AR EP300 CREBBP |
| | TRIM28 PARP1 CREBBP |
| | xxxxxxxxxxxxxxxxxx |
| PAX7-FOXO1 | RARA NCOA1 MYOD1 ESR1 HNF4A PARP1 SMAD3 FOXO1 AR MDM2 EP300 CREBBP |
| | AKT1 EP300 CREBBP |
| | xxxxxxxxxxxxxxxxxx |
| SS18-SSX2 | DPF2 SMARCC2 SMARCC1 PHF10 ELAVL1 DPF3 ARID2 DPF1 SMARCD3 SMARCE1 SMARCD1 EED SMARCA2 EP300 SMARCA4 HDAC1 ARID1B ARID1A ACTL6A HDAC2 CUL3 RNF2 SMARCD2 |
| | GRB2 YWHAG CUL3 |
| | xxxxxxxxxxxxxxxxxx |
| SS18-SSX1 | SMARCC2 DPF2 SMARCC1 PHF10 ELAVL1 DPF3 DPF1 ARID2 HDAC2 SMARCD3 SMARCE1 SMARCD1 EED SMARCA2 EP300 SMARCA4 HDAC1 ARID1B ARID1A ACTL6A CUL3 SMARCD2 |
| | GRB2 YWHAG CUL3 |
| | xxxxxxxxxxxxxxxxxx |
| SS18L1-SSX1 | SMARCC1 STAT3 BMI1 WHSC1L1 SMAD3 HDAC2 SMAD1 EP300 SMARCA4 CREBBP |
| | DPF2 SMARCC1 SMARCE1 SMARCA4 CUL3 |
| | xxxxxxxxxxxxxxxxxx |
| SSX1-SYT4 | |
| | xxxxxxxxxxxxxxxxxx |
| TGFBR3-MGEA5 | MAST1 RNF32 PAXIP1 |
| | CBX8 CSNK2B PAXIP1 |
| | xxxxxxxxxxxxxxxxxx |
| TRIO-TERT | YWHAZ AKT1 RPS6KB1 ENO1 MTOR MDM2 TERT XRCC6 |
| | TERF1 STUB1 TERT POT1 |
| | TERT MTOR YWHAQ RUVBL2 |
| | xxxxxxxxxxxxxxxxxx |
| WDR70-RCOR1 | SMARCC2 HDAC1 HDAC3 HDAC2 KDM1A RCOR1 NR2C1 SMARCE1 CTBP1 NR2E1 SMARCA4 CTBP2 KDM5B MTA3 |
| | KDM1A HDAC3 CTBP1 |
| | xxxxxxxxxxxxxxxxxx |
| YWHAE-NUTM2B | LARP1 NOS2 YWHAQ YWHAG HUWE1 MAST2 NTRK1 VCP AKT1 MAP2K1 FBXW11 ARAF CUL3 PARK2 RAF1 YWHAZ UBXN1 TP53 RUVBL2 BTRC TUBB CDC37 YWHAH MAST3 BRAF YWHAB KSR1 YWHAE CUL1 MAPK7 |
| | YWHAZ IGF1R IRS1 YWHAQ MST1R NTRK1 CBL TUBB YWHAB YWHAH GRB2 ABL1 SORBS2 BCAR1 YWHAE |
| | VCP CDK2 CUL1 |
| | CDC37 LRRK2 YWHAE |
| | xxxxxxxxxxxxxxxxxx |
| YWHAE-NUTM2A-AS1 | LARP1 NOS2 YWHAQ YWHAG HUWE1 MAST2 NTRK1 VCP AKT1 MAP2K1 FBXW11 ARAF CUL3 PARK2 RAF1 YWHAZ UBXN1 TP53 RUVBL2 BTRC TUBB CDC37 YWHAH MAST3 BRAF YWHAB KSR1 YWHAE CUL1 MAPK7 |
| | YWHAZ IGF1R IRS1 YWHAQ MST1R NTRK1 CBL TUBB YWHAB YWHAH GRB2 ABL1 SORBS2 BCAR1 YWHAE |

| | |
|---|---|
| | VCP CDK2 CUL1 |
| | CDC37 LRRK2 YWHAE |
| | xxxxxxxxxxxxxxxxxx |
| YWHAE-NUTM2A | LARP1 NOS2 YWHAQ YWHAG HUWE1 MAST2 NTRK1 VCP AKT1 MAP2K1 FBXW11 ARAF CUL3 PARK2 RAF1 YWHAZ UBXN1 TP53 RUVBL2 BTRC TUBB CDC37 YWHAH MAST3 BRAF YWHAB KSR1 YWHAE CUL1 MAPK7 |
| | YWHAZ IGF1R IRS1 YWHAQ MST1R NTRK1 CBL TUBB YWHAB YWHAH GRB2 ABL1 SORBS2 BCAR1 YWHAE |
| | VCP CDK2 CUL1 |
| | CDC37 LRRK2 YWHAE |
| | xxxxxxxxxxxxxxxxxx |
| **CA** | |
| ANK3-USP9Y | |
| ARGLU1-CXCR4 | APP CHERP SNRNP70 SRPK1 SRPK2 |
| | PTK2 JAK2 SOCS3 PTPN11 NTRK1 |
| | PTK2 ELAVL1 SRPK1 NTRK1 |
| | PTK2 JAK2 JAK3 SOCS3 PTPN11 STAM |
| ATXN10-FBLN1 | FN1 ATXN10 EGFR GSTK1 |
| | ATXN10 VCP BSG CUL3 |
| | VCP ABCE1 ATXN10 APP YWHAQ CUL3 |
| BCAS3-NFS1 | CTBP1 CTBP2 BCAS3 KAT2B |
| | CTBP1 CTBP2 CDC23 KAT2B BCAS3 |
| BCAS4-BCAS3 | CTBP1 CTBP2 BCAS3 KAT2B |
| | CTBP1 CTBP2 CDC23 KAT2B BCAS3 |
| BCL2L12-PRMT1 | NCOA2 NCOA3 NCOA1 TP53 ESR1 BRCA1 PRMT1 THRB CARM1 NR1I2 AR PPARA EP300 |
| | NCOA2 NCOA3 NCOA1 EP300 PARP1 |
| CAPNS1-WDR62 | YWHAZ FBXW11 FN1 HUWE1 YWHAQ GAPDH FERMT2 YWHAH VCAM1 YWHAB PAFAH1B1 YWHAG PAK2 YWHAE |
| | OGFOD1 MYO1E ASNS TBCB CAPN2 PROSC YWHAE |
| CCDC6-ANK3 | HDAC1 NR3C1 TRIM28 SF3A1 HNRNPR BRCC3 |
| | HDAC1 NR3C1 TRIM28 ELAVL1 SKP1 HNRNPR PPP1CA BRCC3 NTRK1 SF3A1 CUL1 FBXW7 |
| CCDC9-DHX34 | EIF4A3 SNIP1 CCDC9 PRPF40A |
| CDC27-ST7L | CDC16 CDC27 MDC1 CDC20 CREBBP ANAPC2 ANAPC7 |
| | CREBBP E2F1 RB1 TFDP1 |
| | CDC16 CDC27 MDC1 SMAD2 TP53BP1 CREBBP UBE2S ANAPC2 ANAPC7 |
| CDK7-RIN3 | BRCA1 TP53 RUVBL2 SUPT5H ESR1 RPA1 HNRNPU RPA2 CDK2 POLR2A |
| | GTF2H1 RPA1 RPA2 POLR2A CCNH |
| | HDAC2 TP53 ESR1 MTA1 |
| | GTF2H1 ERCC3 POLR2A ERCC5 |
| | BRCA1 POLR2A RUVBL2 GTF2H1 RPA1 RPA2 |
| | PRKCI APP CDK7 CDC37 |

| | |
|---|---|
| CHERP-CPAMD8 | DHX8 U2AF1 RPA1 PRPF40A RPA2 CHERP SF3A2 RBM39 EWSR1 |
| | DHX8 AGGF1 RNPS1 SNIP1 SF3B4 NTRK1 CHERP SRPK1 SRPK2 U2AF1 U2AF2 RPA1 APBB1 PRPF40A RPA2 TTC14 RBM23 SF3A2 SNRNP70 RBM39 EWSR1 WBP4 |
| CPD-PIGW | |
| CYTH1-PRPSAP1 | DDX17 DDX5 ILK COPS5 |
| | CYTH1 ARRB2 ARF6 ARRB1 |
| | DDX5 FBXW11 DDX17 ILK ITGB2 COPS5 |
| DLG1-CRYBG3 | DLG1 LIN7A LIN7C APBA1 CASK |
| | DLG1 NTRK1 CASK EPB41 |
| | DLG1 KHDRBS1 LCK NTRK1 |
| | MAPK1 ARRB2 ARRB1 |
| DTX4-CCDC102B | MCM7 CDK18 LENG1 TRIM54 TRIM27 KIFC3 |
| | SFN MARK1 CCDC102B |
| EHD4-FSIP1 | EHD4 CTPS2 EHD1 EGFR NTRK1 WARS PLCG1 UBA2 ADSL UQCRC2 |
| | PLCG1 EHD4 EGFR NTRK1 |
| ELK4-SLC26A9 | BRCA1 MAPK3 MAPK1 ELK4 |
| | BRCA1 MAPK3 MAPK1 ELK4 BLM |
| EMID1-CBY1 | xx |
| EPHA6-CNTN6 | xx |
| ERAL1-DIDO1 | HNRNPDL RPA1 RPA2 HNRNPK RBM15 CUL3 DIDO1 FUS |
| | FUS APP RPA1 RPA2 RBM15 CUL3 DIDO1 SRPK2 |
| GMDS-CCND3 | PCNA PPP1CC RBL2 PPP1CA CCND3 RB1 POLD1 CDK2 CDK4 CDK6 CREBBP |
| | GMDS NSFL1C CTH CAPN2 ATIC |
| | RARA NCOA2 VDR CCND3 CREBBP |
| | MCM10 RBX1 CCND3 APP |
| | NCOA2 RARA VDR CCND3 CREBBP |
| HJURP-EIF4E2 | FBXW11 TP53 GIGYF2 APP HUWE1 EIF4E2 YWHAB SHMT2 YWHAE |
| INTS4-GAB2 | SRC PLCG1 GRB2 ZAP70 NTRK1 SHC1 |
| | PIK3CB PIK3R2 PIK3R1 |
| KCNQ5-RIMS1 | |
| KDM5A-ANO2 | HDAC1 HDAC2 RBL1 TBP RB1 VDR KDM5A MORF4L1 |
| | HDAC2 EZH2 KDM5A ESR1 |
| LAMA5-C12orf28 | |
| MAPK10-FAM13A | HDAC1 TP53 HDAC9 JUN DDX5 ELK1 MAPK10 RELA CREBBP ATF2 |
| | APP MAPK10 MAP2K4 |
| MAPRE1-TM9SF4 | YWHAZ FN1 APP TUBB NTRK1 VCAM1DR/PAK COPS5 |
| | CDK5RAP2 PRKACA AKAP9 PRKACB |
| | CLIP1 TUBB TUBA1A HDAC6 |
| | PDE4DIP PRKACA PRKACB CDK5RAP2 AKAP9 |
| | TERF1 SPTAN1 DST MAPRE1 |
| NCKAP5-MZT2A | |
| NUMB-ALDH6A1 | TP53 NUMB ITCH MDM2 EGFR |
| | EPS15 EGFR AP2A1 NUMB |
| | PRKCZ NUMB APP EGFR |
| PARD6B-CD48 | PRKCI RASSF8 PARD3 PARD6G APP PARD6B PARD6A YWHAH |

| | |
|---|---|
| | PRKCZ WWC1 |
| | PRKCI PARD3 PARD6G APP PARD6B PARD6A YWHAH PRKCZ |
| | RAC1 PARD6G PARD6B PARD6A |
| PPP1R12A-MGAT4C | KDM1A ELAVL1 RPA1 RPA2 PPP1R12A CUL1 |
| | KDM1A NUDT5 TP53 ELAVL1 PUS1 NUAK1 AARSD1 RPA1 NTRK1 RPA2 RPRD1B PPP1R12A TRIM47 PAXIP1 ACTR3 CUL1 |
| RNF11-C8A | CBLB RNF11 ITCH SMAD4 EPN1 RABGEF1 UBE2E1 UBE2D3 UBE2E3 HGS GGA1 AKT1 GGA3 GGA2 AP2A1 EPN3 UBE2D1 AP2B1 CSNK2A1 SMURF1 EPS15 SMURF2 UBQLN2 STAM2 NEDD4 UBQLN4 NEDD4L APP |
| | RNF11 PSMD4 PSMD7 PSMD6 PSMD11 PSMD10 PSMD3 PSMD12 PSMD13 USP14 PSMD14 PSMD1 PSMD2 APP |
| | GGA1 GGA3 GGA2 APP |
| | AKT1 RNF11 TBK1 APP NEDD4 |
| SIPA1L3-WDR62 | MAPK10 WDR62 MAPK8 MAPK9 |
| | SFN YWHAB SIPA1L3 YWHAQ |
| | YWHAB FBXW11 MAPK10 ELAVL1 WDR62 TBP MAPK8 MAPK9 |
| SLC26A6-PRKAR2A | AKAP7 AKAP9 PRKAR2A PRKAR2B PRKACA PRKACB |
| | PRKAR2A AKAP7 PRKACA PRKACB PRKAR2B GCH1 |
| ST14-APLP2 | BRCA1 APLP2 ETS1 JUN |
| | SFN HDAC5 APLP2 RPL26 |
| | BRCA1 JUNB JUN APBB1 APBB2 KAT5 ETS1 APLP2 MAPK8 |
| STRADB-NOP58 | NIFK NOP56 HNRNPU RUVBL2 NOLC1 SNU13 NTRK1 PUM3 RPS15A RSL1D1 KRR1 DDX18 EIF6 DHX15 RPS4X EED PRPF3 TARDBP RPL11 EIF2S2 DDX27 NOP58 FN1 DDX24 U2AF1 RPL30 ESR1 DDX56 FTSJ3 DDX47 KPNA6 GTPBP4 WDR36 KPNA1 DKC1 RRP12 BOP1 FBL TBL3 |
| | WDR36 NOP58 DHX15 NOP56 |
| STX16-RAE1 | FBXW11 NXF1 FAF1 RAE1 ILF3 CUL1 HNRNPUL1 CUL3 |
| | NXF1 CUL1 ILF3 CUL3 |
| TANC2-CHD6 | ZFYVE9 PPP1CC PPP1CA TANC2 |
| THSD7B-DARS | ZFYVE9 PPP1CC PPP1CA TANC2 |
| TMEM123-MMP7 | MAEA RANBP9 MKLN1 MMP7 RMND5A |
| | MAEA RANBP9 MKLN1 RANBP10 MMP7 RMND5A |
| TMPRSS2-ERG | CDC5L DDX3X ELAVL1 CAD NEDD4 SF3B1 PARP1 PRKDC PRPF8 SFPQ ERG POLR2A TOP1 CLTC SF3B2 XRCC5 XRCC6 DDX23 SNRNP200 DDX21 TUBB NONO JUN HNRNPU PRPF40A AR NCL HNRNPM HNRNPC TOP2B ILF3 ILF2 |
| | PRPF8 ERG SF3B2 SF3B1 PARP1 PRKDC |
| TOX3-CNTN5 | |
| UBR2-XPO5 | |
| UVRAG-INTS4 | |
| WNT11-TSPAN8 | |

| |
|---|
| XRCC5-ACADL |
| ZCCHC7-PRSS3 |
| ZFP91-NOX4 |

***Table S12: OG and TS predicted by NBC, MutsigCV and 20/20+***

| Tools | Number of predicted driver genes | Percentage in TCGA |
|---|---|---|
| NBC | 2750 | 0.68 |
| RFS | 870 | 0.18 |
| DeepLearning | 560 | 0.2 |
| Wrzeszczynski2011 | 430 | 0.22 |
| MutSigCV | 1080 | 0.38 |
| OncodriveCLUST | 430 | 0.095 |
| OncodriveFM | 210 | 0.06 |
| 20/20+ | 940 | 0.45 |
| ActiveDriver | 230 | 0.08 |
| MuSiC | 580 | 0.06 |
| TUSON | 900 | 0.4 |
| OncodriveFML | 810 | 0.14 |

***Table S13: Prediction rate of unique OG and TS by NBC, MuSiC, OncodriveCLUST, OncodriveFM, 20/20+***

| Tools | Percentage of predicted OG/TS in TSGene,Kumar et al,TissGDB | Actual number of predicted OG/TS |
|---|---|---|
| NBC | 610 | 93.05 |
| RFS | 465 | 70 |
| DeepLearning | 180 | 58 |
| Wrzeszczynski2011 | 210 | 59 |
| MutSigCV | 370 | 68 |
| OncodriveCLUST | 490 | 72.25 |
| OncodriveFM | 501 | 79.75 |
| 20/20+ | 520 | 89 |
| ActiveDriver | 160 | 57 |
| MuSiC | 510 | 87 |
| TUSON | 300 | 61 |
| OncodriveFML | 320 | 63 |

***Table S14. Comparison of NBC with others with respect to p-value***

| Method | Test to TS | Test to OG | DR/PA to TS | DR/PA to OG |
|---|---|---|---|---|
| NBC | 0.925 | 0.942 | 0.912 | 0.943 |
| RFS | 0.91 | 0.94 | 0.88 | 0.93 |
| DeepLearning | 0.6 | 0.58 | 0.67 | 0.62 |
| Wrzeszczynski2011 | 0.63 | 0.65 | 0.68 | 0.69 |
| MutSigCV | 0.71 | 0.88 | 0.61 | 0.71 |
| OncodriveCLUST | 0.72 | 0.75 | 0.81 | 0.61 |
| OncodriveFM | 0.83 | 0.91 | 0.74 | 0.71 |
| 20/20+ | 0.86 | 0.83 | 0.89 | 0.88 |
| ActiveDriver | 0.71 | 0.73 | 0.68 | 0.70 |
| MuSiC | 0.65 | 0.61 | 0.69 | 0.66 |
| TUSON | 0.63 | 0.67 | 0.61 | 0.62 |
| OncodriveFML | 0.82 | 0.88 | 0.76 | 0.75 |
